# Full-length *de novo* protein structure determination from cryo-EM maps using deep learning

**DOI:** 10.1101/2020.08.28.271981

**Authors:** Jiahua He, Sheng-You Huang

## Abstract

Advances in microscopy instruments and image processing algorithms have led to an increasing number of cryo-EM maps. However, building accurate models for the EM maps at 3-5 Å resolution remains a challenging and time-consuming process. With the rapid growth of deposited EM maps, there is an increasing gap between the maps and reconstructed/modeled 3-dimensional (3D) structures. Therefore, automatic reconstruction of atomic-accuracy full-atom structures from EM maps is pressingly needed. Here, we present a semi-automatic *de novo* structure determination method using a deep learning-based framework, named as DeepMM, which builds atomic-accuracy all-atom models from cryo-EM maps at near-atomic resolution. In our method, the main-chain and Cα positions as well as their amino acid and secondary structure types are predicted in the EM map using Densely Connected Convolutional Networks. DeepMM was extensively validated on 40 simulated maps at 5 Å resolution and 30 experimental maps at 2.6-4.8 Å resolution as well as an EMDB-wide data set of 2931 experimental maps at 2.6-4.9 Å resolution, and compared with state-of-the-art algorithms including RosettaES, MAINMAST, and Phenix. Overall, our DeepMM algorithm obtained a significant improvement over existing methods in terms of both accuracy and coverage in building full-length protein structures on all test sets, demonstrating the efficacy and general applicability of DeepMM.

**Availability:** https://github.com/JiahuaHe/DeepMM

**Supplementary information:** Supplementary data are available.

## 1 Introduction

Cryo-electron microscopy (cryo-EM) has now become a widely used technique for structure determination of macromolecular structures in the recent decade^1–4^. Advances in microscopy instruments and image processing algorithms have led to the rapid increase in the number of solved EM maps^1–3^. The ‘resolution revolution’ in cryo-EM has paved a way for the determination of high-resolution structures of previously intractable biological systems^5–16^. According to the statistics of the Electron Microscopy Data Bank (EMDB)^17^, there were 2435 maps deposited in 2019, which are almost 4 times the 640 maps released in 2015.

With the rapid growth of deposited EM maps, there is an increasing gap between the maps and reconstructed/modeled 3-dimensional (3D) structures. As of April 1, 2020, there were 10560 EMDB maps, but only 4805 associated structures were deposited in the Protein Data Bank (PDB)^18^. For those maps determined at near-atomic resolution (3.0∼ 5.0 Å), it is difficult to build high-resolution models with conventional software designed for X-ray crystallography. In view of the fact that near-atomic resolution maps take up the majority of current and henceforth released maps^17^, tools, which can reconstruct structures *de novo* from EM maps without using known structures as templates^19^, are pressingly needed. As such, some algorithms like EM-fold^20^, Gorgon^21^, Rosetta^22,23^, Pathwalking^24–26^, Phenix^27–29^, and MAINMAST^30,31^, have been recently presented for constructing and/or assembling structure fragments from Cryo-EM maps.

Despite the present progress in *de novo* structure building for cryo-EM maps, there are various limitations in current approaches. They can either only build structural fragments^20,21,28^ or have low accuracy in terms coverage and/or sequence reproduction^23,24,30^. It remains challenging to automatically build an accurate all-atom structure from the EM maps at near-atomic resolution. Recently, machine learning has been actively applied in structure determination for EM maps, such as single particle picking^32^, tomogram annotation^33^, secondary structure prediction^34^, and backbone tracing^35^. However, applying deep learning to build full-length protein structures for near-atomic resolution EM maps remains a challenging work.

Here, we have developed a semi-automatic *de novo* atomic-accuracy structure reconstruction method for EM maps at near-atomic resolution through Densely Connected Convolutional Networks (DenseNets) using a deep learning-based framework, named DeepMM. Instead of tracing the protein main-chain on the raw EM density map, DeepMM first predicted the probability of main-chain atoms (N, C, and C*α*) and C*α* positions near each grid point using one DenseNet^36^. Then, the method traced the main-chain according to the predicted main-chain probability map. The amino acid and secondary structure types were predicted by a second DenseNet. Finally, the protein sequence was aligned to the main-chain according to the predicted C*α* probabilities, amino acid types, and secondary structure types for all-atom structure building.

## 2 Methods

### 2.1 Workflow of DeepMM

The workflow of DeepMM is illustrated in Figure 1**a**. Specifically, staring from a cryo-EM map and the target protein sequence, DeepMM first standardizes the order of axis, and interpolates grid interval to 1.0 Å. Then, DeepMM cuts the entire map into small voxels of size 11Å × 11Å × 11Å. Afterwards, one DenseNet (say DenseNet A) is used to predict the main-chain and C*α* probability on each of the voxels. All the predicted probability values form a 3D probability map. Next, possible mainchain paths are generated in the predicted main-chain probability map using a main-chain tracing algorithm^30^. The C*α* probability values of main-chain points are interpolated from the predicted 3D C*α* probability map. Afterwards, the amino acid and secondary structure types are predicted for each main-chain point through the second DenseNet (say DenseNet B). With the predicted C*α* probability, amino acid type, and secondary structure type for each main-chain point, the target protein sequence is then aligned to the main-chain paths based on the Smith-Waterman dynamic programming (DP) algorithm^37^. The resulted multiple C*α* models are ranked by their alignment scores. Finally, the all-atom structures are constructed from the top C*α* models using the *ctrip* program in the Jackal modeling package ^38,39^ and refined by an energy minimization using Amber^40^.

**Figure 1:**
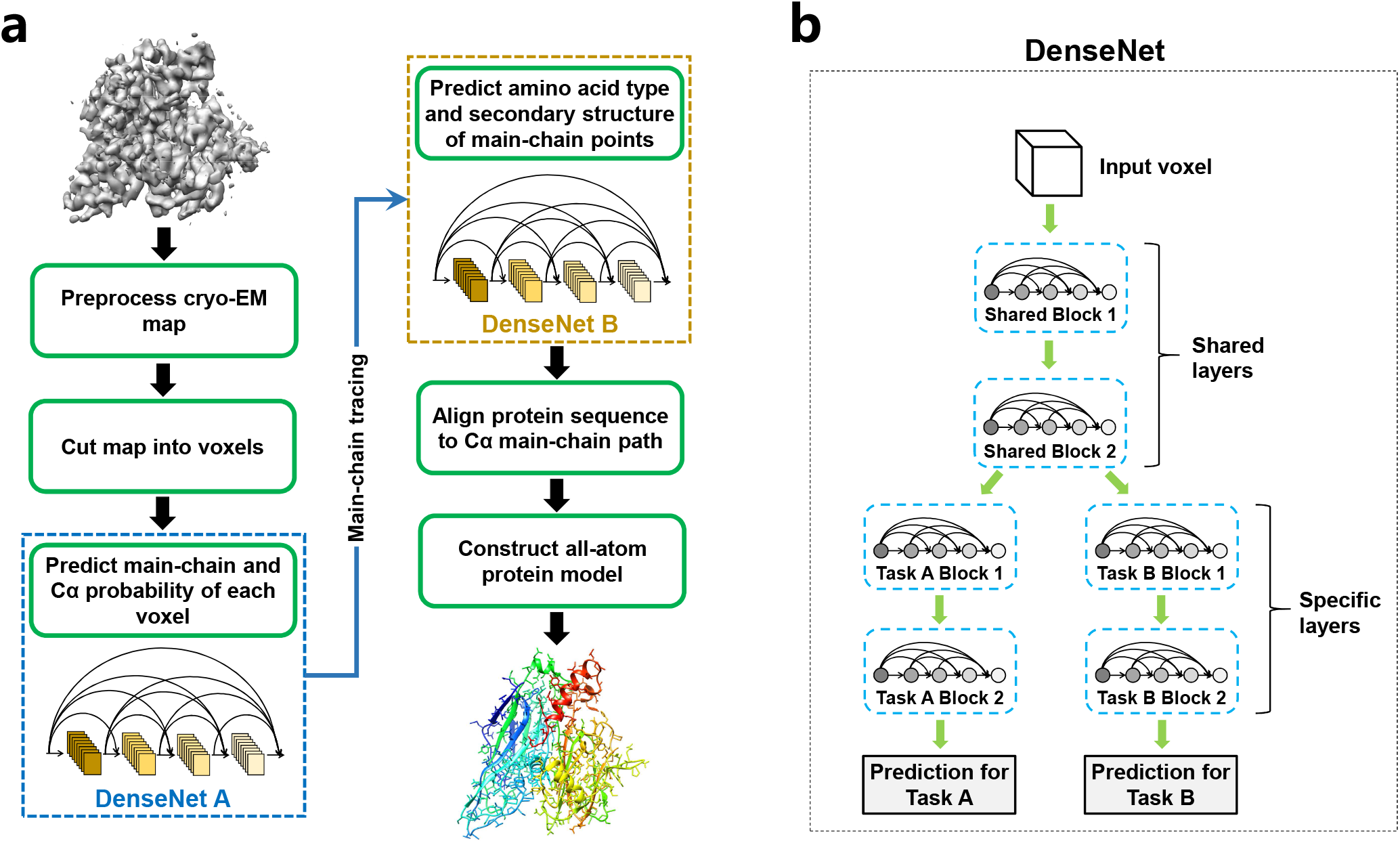
Workflow of our DeepMM method. **(a)** The flowchart of DeepMM. DeepMM first predicts the main-chain and C*α* probability of each density voxel using a Densely Connected Convolutional Network (DenseNet), and then traces the protein’s main-chain path on the predicted main-chain probability map. Next, the amino acid and secondary structure types for each main chain point are predicted by a second DenseNet. The C*α* models are generated by aligning the target sequence to the main-chain paths. Finally, the all-atom structures are constructed from the C*α* models using the ctrip program and refined by an Amber energy minimization. **(b)** The multi-task deep DenseNet architecture used in DeepMM. Starting from an input EM density voxel, two dense blocks are shared by both tasks in DenseNet A, while only one dense block is shared by both tasks in DenseNet B. Each prediction task employs two task-specific dense blocks and gives the final prediction.

### 2.2 Training the DenseNets of DeepMM

Two Densely Connected Convolutional Networks (DenseNets) are embedded into our DeepMM algorithm. Figure 1**b** illustrates the architecture of the networks. DenseNet is a feed-forward multi-layer network which uses additional paths between earlier and later layers in a dense block. DenseNets have several compelling advantages. They alleviate the vanishing-gradient problem, strengthen feature propagation, encourage feature reuse, and substantially reduce the number of parameters^36^. DeepMM also employs a hard parameter-sharing multi-task learning method, which can greatly reduces the risk of overfitting^41^. The first network (i.e. DenseNet A) is used to simultaneously predict the main-chain probability and C*α* probability of a grid point. The second network (i.e. DenseNet B) is used to predict the amino acid type and secondary structure type of a main-chain local dense point (LDP). The input for the DenseNet A are voxels of size 11Å × 11Å × 11Å. The second network (DenseNet B) takes the voxels of size 10Å × 10Å × 10Å as input because main-chain points are not always on the integer grid after mean shift. For each voxel, the density values are normalized to the range of [0, 1] according to the maximum and minimum density values in the voxel. 3D convolutions and 3D pooling layers are used instead of their 2D counterparts used in traditional image processing because the density maps have three dimensions. Several dense blocks are used in both networks, each of which consists of eight densely connected layers. For DenseNet A, the first two dense blocks are shared by both tasks, whereas for DenseNet B, only one shared block is adopted. After the shared blocks, each task employs two task-specific blocks and gives the final prediction. The details of network architecture are provided in Supplementary Table 1.

All the training parameters and procedure used for simulated EM maps are essentially the same to the parameters and procedure used for experimental EM maps unless otherwise specified.

For DenseNet A, all the grid points above a density value *D*_0_ were used for training, where *D*_0_ was set to 1.0 for simulated maps at 5.0 Å resolution. For experimental maps, *D*_0_ was set to 1/2 of its recommended contour level. The labels (main-chain probability and C*α* probability) of a grid point *a* were calculated as follows:

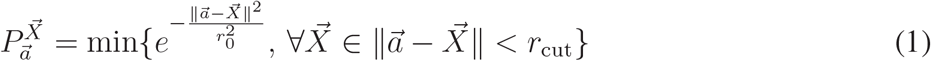

where *X* stands for the N, C, or C*α* atoms. The *r*_0_ is the radius at which the probability drop to 1*/e*. If no atom is within *r*_cut_ of a grid point, the corresponding probability is set to 0. A total of 512 voxels were trained in one batch and 30 epochs were trained for the whole data set. The Adam optimizer with an initial learning rate of 0.001 was used to minimize the mean absolute error (MAE). Learning rate decay was adopted, where the learning rate was reduced to 1/10 of the current value after every 10 epochs. To avoid over-fitting, the *weight decay* parameter of Adam optimizer was set to 1e-6 as the L2 regularization.

For DenseNet B, one point was randomly sampled within 1.0 Å for every main-chain atom in the training set. The corresponding amino acid type and second structure type marked by STRIDE^42^ were assigned to each point. Twenty types of amino acids were grouped into four classes according to their sizes, shapes and distributions in their EM density maps^43^, as illustrated in Figure 2**d**. Specifically, GLY, ALA, SER, CYS, VAL, THR, ILE and PRO are grouped as Class I. LEU, ASP, ASN, GLU, GLN and MET are grouped as Class II. LYS and ARG are grouped as Class III. HIS, PHE, TYR and TRP are grouped as Class IV. Residues that have structure codes of H, G, or I by STRIDE were labelled as “Helix”, those with codes of B/b or E were labelled as “Sheet”, and the other residues were labelled as “Coil”. All the training parameters were identical to those for DenseNet A except for using CrossEntropyLoss as loss function.

**Figure 2:**
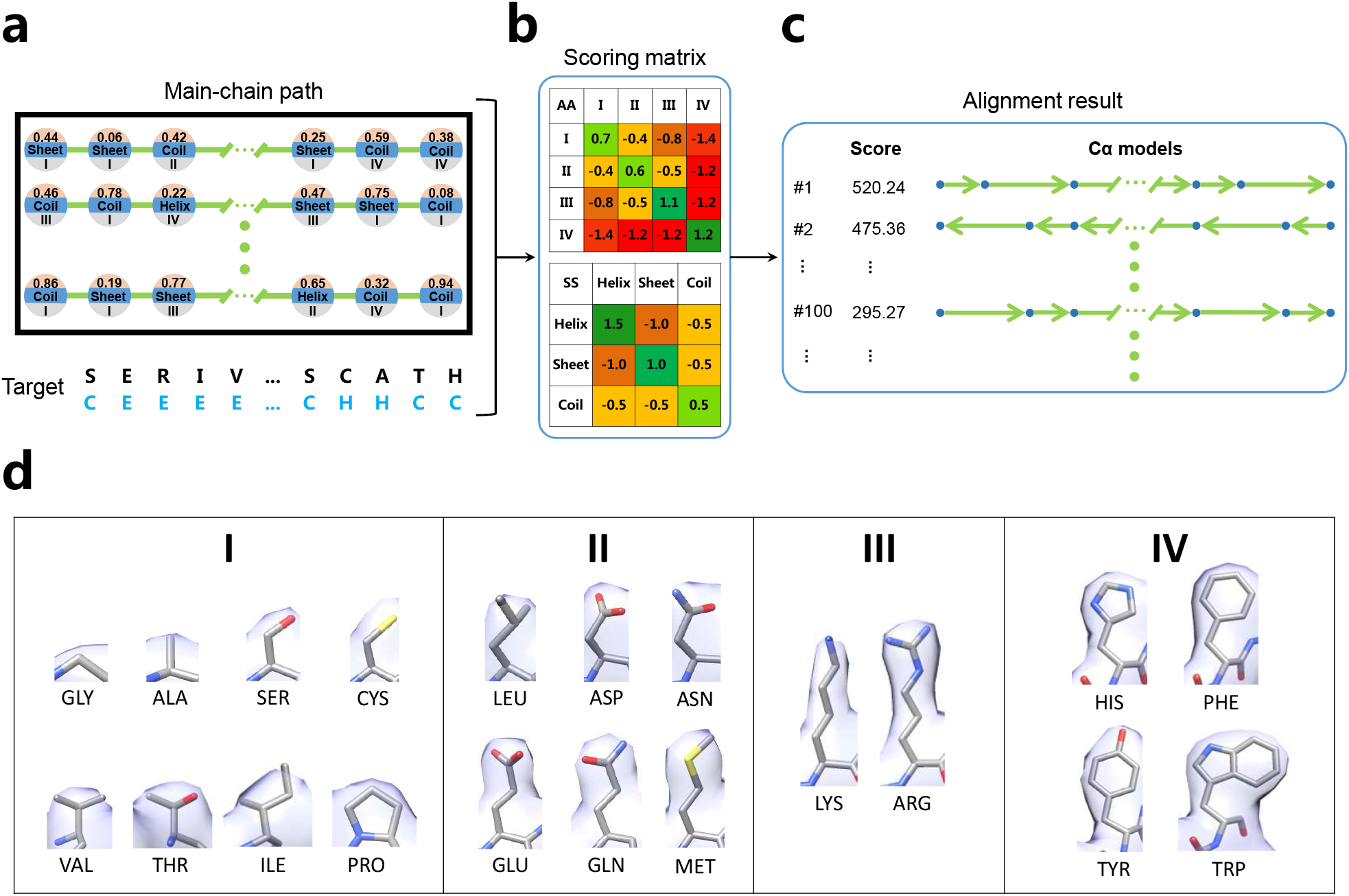
Alignment protocol between the target sequence and the predicted main-chain for DeepMM. **(a)** DeepMM runs alignments of the target sequence of the EM map against each candidate main-chain path. Each sphere represents a predicted local dense point (LDP) on the main-chain path. Predicted information including the C*α* probability (on the top), secondary structure (in the middle) and amino acid class (at the bottom) of LDPs is utilized during alignment. For the target sequence, its secondary structure is predicted by the SPIDER2 program, as illustrated in the sequence colored in azure under the amino acid sequence. **(b)** Scoring matrices for amino acid type matching and secondary structure matching. **(c)** The generated C*α* models are ranked by their alignment score. **(d)** Twenty amino acids are grouped into four classed according to the similarity of their side-chain EM densities.

### 2.3 Tracing the main-chain path

The main-chain tracing algorithm in MAINMAST^30^ was used to trace the main-chain path in our predicted main-chain probability map. In brief, local dense points (LDPs) are first identified using the mean shift algorithm, which iteratively shifts the initial grid points towards the local highest probability by computing the weighted average of probability values. Then, the shifted points that are within a threshold distance of 0.5 Å are clustered, and the point with the highest probability in the cluster is chosen as the representative, called LDP. The next step is to connect LDPs into a minimum spanning tree (MST) and iteratively refine the tree structure with a Tabu search method. After multiple steps of Tabu search, the longest path of the refined tree is traced as the main-chain path. The details of the algorithm can be found in the MAINMAST study^30^.

### 2.4 Aligning target sequence to main-chain path

The Smith-Waterman dynamic programming (DP) algorithm^37^ is used to align the target sequence to the predicted main-chain path. The predicted C*α* probability value, amino acid type, and secondary structure type are assigned to each point of the main-chain. Instead of using 20 amino acid types, amino acids are grouped into four classes according to their sizes, shapes, and distributions in EM density maps (Figure 2**d**). Secondary structures are categorized into three types of Helix, Sheet, and Coil. The match between the target sequence and main-chain path is evaluated by two scoring matrices for amino acid and secondary structure, respectively (Figure 2**b**). Namely, a target residue is more likely to be aligned to a main-chain point with the same amino acid type, the same secondary structure type, and a higher C*α* probability, and vice versa. The detailed alignment protocol is shown in Figures 2**a, b** and **c**. The *n* residues {*A*_i_(*i* = 1, *… n*)} in the protein are aligned to *m* LDPs {*L*_j_(*j* = 1, *… m*)} in the main-chain path. The matching score *M* (*i, j*) for a pair of *A*_i_ and *L*_j_ is computed as follows.

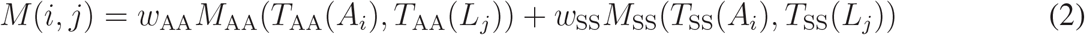

where *M*_AA_ and *M*_AA_ are the scoring matrices for amino acid and secondary structure matching^43^, 44, respectively. For a residue *A*_i_, the amino acid type is one of the four amino acid classes (*T*_AA_(*A*_i_) = 1, 2, 3, 4). The predicted amino acid type for an LDP *L*_j_ is also one of the four amino acid classes (*T*_AA_(*L*_i_) = 1, 2, 3, 4). Similarly, the secondary structure matching score is calculated using the secondary structure type predicted from the sequence (*T*_SS_(*A*_i_) = 1, 2, 3) by SPIDER2^45^ and secondary structure type predicted on LDPs (*T*_SS_(*L*_i_) = 1, 2, 3). The scoring matrices *M*_AA_ and *M*_SS_ used in the alignment are shown in Figure 2**b**. The *w*_AA_ and *w*_SS_ are the weights for corresponding matching scores and set to 1.0 and 0.5, respectively. With the calculated matching score *M* (*i, j*), an alignment is calculated with the follow rule to form a DP matrix, *F*, as follows.

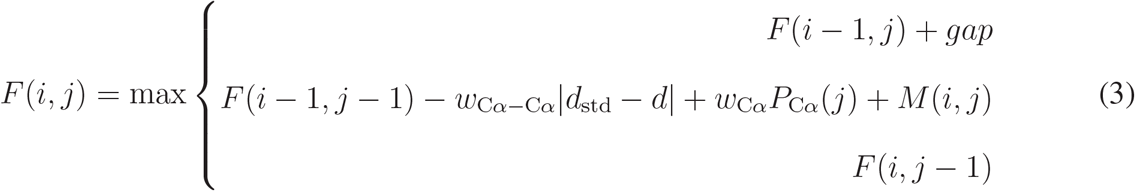

where *gap* is the gap penalty for unassigned residues in the protein sequence. To ensure a full-length structure reconstruction, *gap* is set to −10000.0 so as to forbid skipped residues. The |*d*_std_ − *d*| is the penalty score for C*α*-C*α* distance, where *d*_std_ is the standard C*α*-C*α* distance and *d* is the distance between LDP *L*_j_ and the last aligned LDP. The *P*_Cα_(*j*) is the predicted C*α* probability for LDP *L*_j_. The *w*_Cα−Cα_ and *w*_Cα_ are the weights for the corresponding scores. Here, *w*_Cα_ is set to 1.6, and *w*_Cα−Cα_ is set to 1.0, 0.7, and 0.8 for “Helix”, “Sheet”, and “Coil”, respectively. For each combination of parameters in the main-chain tracing procedure, 160 C*α* models are generated. Finally, all the generated C*α* models are ranked by their alignment scores.

### 2.5 Parameter settings of DeepMM

The parameters of mean-shift, MST construction, and Tabu search are set to be the same to those in MAINMAST^30^, unless otherwise specified. DeepMM employs several parameter combinations to generate multiple C*α* models for one EM map. For each combination of parameters, 10 trajectories of Tabu search are carried out, yielding 10 main-chain paths. Since DeepMM starts from the main-chain probability map, fewer parameter combinations are needed to reconstruct reliable 3D structures. For both simulated and experimental maps, the thresholds of probability (Φ_thr_) and normalized probability (*θ*_thr_) are both set to 0. For the 40 simulated maps, only one parameter combination is adopted. Specifically, the maximum number of Tabu search steps (*N*_roundr_) is set to 100, the sphere radius of local MST (*r*_local_) is set to 5.0 Å, and the constraint for the length (*d*_keep_) is set to 0.5 Å. For the 30 experimental maps, we employ the following 27 combinations of parameters: the sphere radius of local MST (*r*_local_=5.0, 7.5, 10.0 Å), the edge weight threshold (*d*_keep_=0.5, 1.0, 1.5 Å), and the maximum number of the Tabu search steps (*N*_roundr_=2500, 5000, 7500). For the extended EMDB-wide test set of 2931 maps, we employ fewer combinations of parameters so as to save computational cost: the edge weight threshold (*d*_keep_=0.5, 1.0 Å) and the maximum number of the Tabu search steps (*N*_roundr_=2500, 5000). The sphere radius of local MST (*r*_local_) is set to 10 Å. For each of the generated main-chain path, 16 C*α* models are generated using 8 different standard C*α*-C*α* distances (*d*_std_=3.1, 3.2, 3.3, 3.4, 3.5, 3.6, 3.7, 3.8 Å) on two sequence directions. Namely, 160 models (16 models for each of the 10 trajectories) are constructed for each parameter combination. The C*α* models are ranked by their alignment scores and then an RMSD cutoff of 5 Å is used to remove the one with lower alignment score in two similar structures. Finally, the top 10 scored protein C*α* models are selected to build the all-atom structures.

### 2.6 Datasets used

#### 2.6.1 Training sets

Two data sets, simulated EM map set and experimental EM map set, were used to train our DeepMM method for simulated maps and experimental maps, respectively.

For simulated EM maps, 2000 representative structures for different superfamilies in the SCOPe database^46^ were taken from Emap2sec^34^ as training set. Those structures were removed from the training set if they have a TM-score^47^ of over 0.5 with any structure in the test set. To save the computational cost, only 100 randomly selected structures from the training set were retained. Next, we used the *e2pdb2mrc*.*py* program from the EMAN2 package (version 2.11)^48^ to generate the simulated EM maps at 5.0 Å resolution and 1.0 Å grid interval for each structure in training and test set. The training SCOPe entries used in this study were listed in Supplementary Table 5.

For experimental EM maps, all the EM density maps at 2-5 Å resolution that have associated PDB models were downloaded from the EMDB. As of December 26, 2019, 2546 EM maps were collected. Any PDB structure and its corresponding EM map that met the following criteria were removed: (i) including nucleic acids, (ii) missing side-chain atoms, (iii) including “HETATM” residues, (iv) including “UNK” residues, (v) including more than 1 subunit (MODEL), and (vi) including less than 50 or more than 300 residues. Then, 1588 chains from the remaining 361 experimental EM maps were clustered with 50% sequence identity using CD-HIT^49^, yielding a total of 1340 chains. To ensure a valid evaluation, chains were removed from training set if they have over 30% sequence identity with any chain in the test set. Each protein chain was zoned out from the whole map using a distance of 4.0 Å ^30^. For good quality maps, protein chain and its associated map should have sufficient structural agreement. The cross-correlation between the experimental map and the simulated map density at the same resolution with the experimental map generated from the structure was calculated using the UCSF Chimera^50^. Only the chains with a cross-correlation of over 0.65 were kept^34^. The final training set consists of 100 non-redundant protein chains. The grid intervals for experimental maps were unified to 1.0 Å using trilinear interpolation. The training EM maps and their corresponding PDB chains used in this study are listed in Supplementary Table 6.

#### 2.6.2 Test sets

Three test sets were used to evaluate our DeepMM approach for its accuracy and general applicability, including one simulated map set and two experimental maps.

The simulated map set was taken from the test set of 40 simulated maps used by MAINMAST^30^. The maps were generated at 5.0 Å resolution with a grid spacing of 1.0 Å using the *e*2*pdb*2*mrc*.*py* program in the EMAN2 package^48^.

The first experimental test set is the benchmark of 30 EM maps at 2.6-4.8 Å resolution, which have been used to evaluate MAINMAST^30^. The corresponding EM maps were downloaded from the EMDB, For each EM map, a single subunit was zoned out from the whole density map at a distance cutoff of 4.0 Å.

In addition, to evaluate the accuracy and general applicability of DeepMM, we have also constructed a large test set of EMBD-wide experimental maps. The generation procedure of this set was similar to that for the experimental training set. Specifically, for each chain of the EM PDB structure at 2.5-5.0 Å resolution and no more than one subunit (MODEL) from the EMDB, a single density patch was zoned out from the whole density map at a distance cutoff of 4.0 Å. Any protein chain and its corresponding EM map patch that met the following situation were removed: (i) including nucleic acids, (ii) missing side-chain atoms, (iii) including “HETATM” residues, (iv) including “UNK” residues, (v) including less than 50 or equal or more than 300 residues, (vi) having over 30% sequence identity to any chain in the training set. The cross-correlation between the experimental map and the simulated density map at the same resolution generated from the structure should be over 0.65^34^. Each protein chain was zoned out from the whole map using a distance of 4.0 Å ^30^. The finial test set consists of 2931 protein chains, which are listed in Supplementary Table 4.

## 3 Results

### 3.1 Model reconstruction for simulated EM maps

We first evaluated the performance of our DeepMM algorithm on the test set of 40 simulated density maps at 5 Å resolution. DeepMM traced the main-chain of protein on the predicted main-chain probability map rather than the raw EM density map. Thus, the generated C*α* models by our DeepMM are closer to the native structures with fewer search trajectories and steps compared to MAINMAST. For each of the 40 maps, DeepMM built 160 C*α* models, which were ranked by their alignment scores. The top-ranked model was selected as the predicted structure.

Figure 3 shows a comparison of the predicted C*α* models for the protein chains of different lengths by DeepMM and MAINMAST. The detailed results are provided in Supplementary Table 2. It can be seen from the figure that our DeepMM method obtained a much better performance than MAIN-MAST. As shown in Figure 3**a**, DeepMM built significantly more accurate C*α* models, and achieved an average C*α* RMSD of 0.54 Å \ when the top scored model was considered, compared to 1.79 Å for MAINMAST. DeepMM also generated high-quality models with less than 1.0 Å C*α* RMSD for all of the 40 maps, compared with only one such model by MAINMAST. Moreover, DeepMM achieved the high-accuracy models with less than 0.5 Å RMSD for 22 of 40 maps, whereas MAINMAST failed to generate any model with *<* 0.5 Å RMSD (Figure 3a). The program CLICK^51^ was also used to evaluate the accuracy of the C*α* models built by DeepMM and MAINMAST. The corresponding results are shown in Figure 3**b**. Similar to the results of C*α* RMSD comparison, DeepMM generated many more high-quality models according to the CLICK RMSD criterion and achieved an average CLICK RMSD of 0.53 Å when the top model was considered, compared to 2.18 Å for MAINMAST.

**Figure 3:**
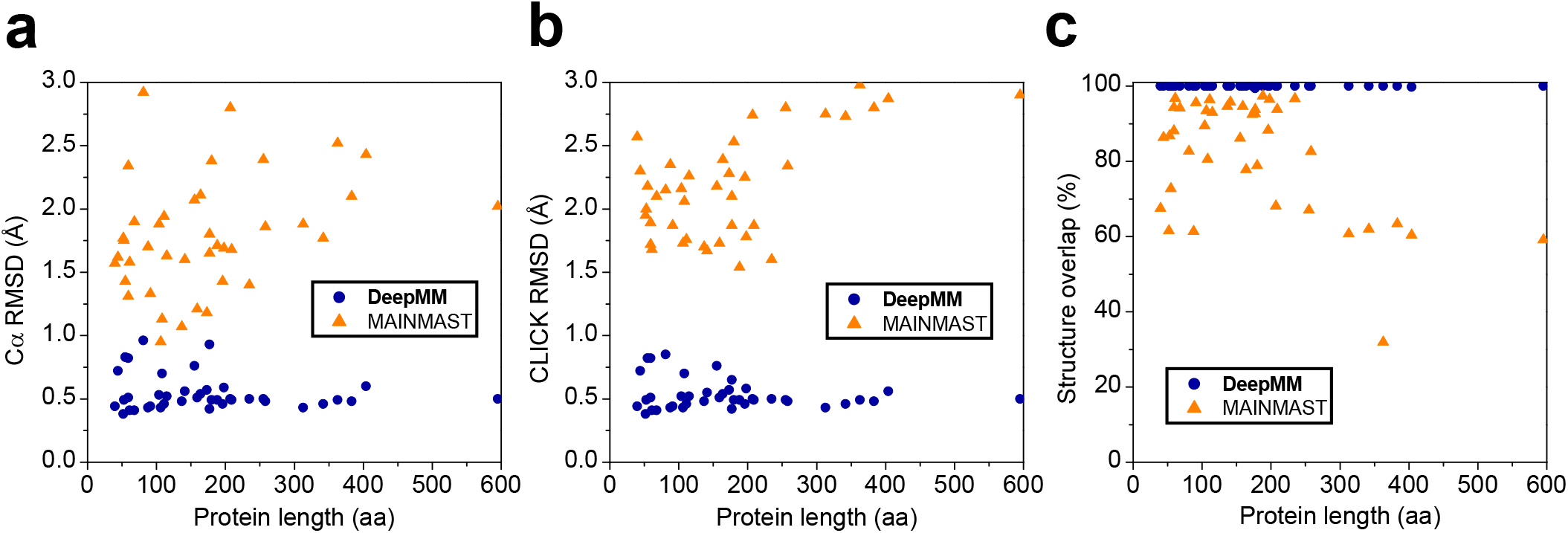
Comparison of the results by DeepMM and MAINMAST for the protein chains with different lengths. **(a)** The C*α* RMSDs of the top predicted models. **(b)** The RMSDs of matched C*α* atoms within 3.5 Å by the structure alignment tool CLICK. **(c)** The structure overlap calculated by CLICK, which is defined as the fraction of matched C*α* atoms.

In addition, DeepMM also achieved a significantly higher structure overlap than MAINMAST (Figure 3**c**). Except for two top scored models with 99.75% and 99.44% structure overlap, the rest 38 top models generated by DeepMM all have a 100% structure overlap. On average, DeepMM obtained a high structure overlap of 99.98%, compare to 81.88% for MAINMAST. Figure 3 also reveals that DeepMM generated consistently high-accuracy models for all the proteins of different lengthes, whereas MAINMAST tended to perform worse with the increasing number of residues in the protein, suggesting the higher robustness of DeepMM than MAINMAST.

### 3.2 Model reconstruction for experimental EM maps

Our DeepMM method was further tested on the benchmark of 30 experimental density maps at 2.6-4.8 Å resolution. For each of the 30 experimental density maps, DeepMM built 4320 protein C*α* models, which were then ranked by their alignment scores.

Figure 4**a** shows a comparison of the C*α* RMSDs for the models built by DeepMM and MAIN-MAST. The corresponding data are provided in Supplementary Table 3. It can be seen from the figure that DeepMM generated significantly more accurate models than MAINMAST. On average, DeepMM obtained a C*α* RMSD of 10.7 Å for the top scored models, which is much better than 22.4 Å by MAINMAST. Moreover, DeepMM predicted a model of *<* 10 Å for 18 out of 30 top scored models, of which 14 models are within 5.0 Å C*α* RMSD. By contrast, only 7 and 4 models are within 10.0 Å and 5.0 Å for MAINMAST, respectively. Figure 4**b** shows a comparison of the results for the models predicted by DeepMM and RosettaES. It can be seen from the figure that DeepMM performed much better and generated many more accurate models than RosettaES. Compared to 18 models within 10 Å RMSD by DeepMM, only six models were predicted within 10.0 Å RMSD by RosettaES for the top predictions. On average, Rosetta obtained an average C*α* RMSD of 27.0 Å, which is much higher than 10.7 Å for DeepMM.

**Figure 4:**
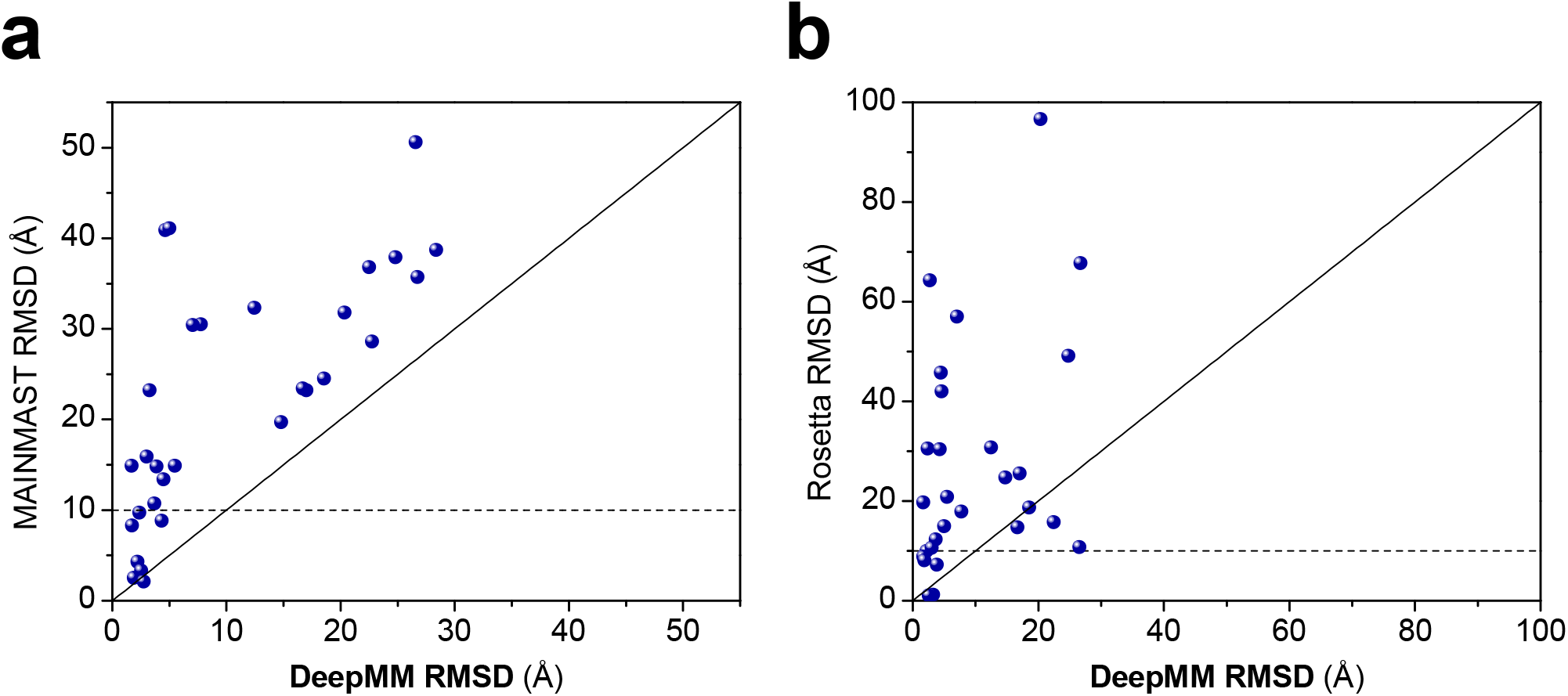
Comparison of the top models for DeepMM and two other approaches on the test set of 30 experimental maps. The solid line in the figure is the plot of *y* = *x*, and the dashed line stands for *y* = 10. **(a)** Comparison of the models by DeepMM and MAINMAST in terms of C*α* RMSD. **(b)** Comparison of the models by DeepMM and Rosetta in terms of C*α* RMSD.

Further examination of the predicted results also reveals that the model accuracy depends more on the quality than on the resolution of a map. Namely, compared to maps with relatively higher resolution but lower quality like EMD-3246A/B (2.8 Å) and EMD-5495 (3.5 Å), maps with relatively lower resolution but higher quality like EMD-2867 (4.3 Å) and EMD-3073 (4.1 Å) are more likely to be successful in reconstructing a correct model (Supplementary Table 3). This phenomenon can be attributed to the fact that resolution is a global estimation and resolvability is not necessarily uniform throughout the whole map^52^.

Figure 5 gives two examples of successfully reconstructed structures by DeepMM. One example, EMD-2867, which is a nucleoprotein at 4.3 Å resolution, was successfully reconstructed by DeepMM, as shown in Figure 5**a**. It can be seen from the figure that the predicted main-chain by DeepMM overlaps well with that of the deposited structure. Accordingly, the predicted model shows an atomic-accuracy with a C*α* RMSD of 3.1 Å. Figure 5**b** shows the results of another example, EMD-6272, which is the bovine rotavirus VP6 at 2.6 Å resolution. Because of its high resolution, DeepMM predicted a very high accurate model with a small C*α* RMSD of 1.7 Å. Correspondingly, the constructed full-atom model by DeepMM shows an excellent overlap with the deposited structure (Figure 5**b**).

**Figure 5:**
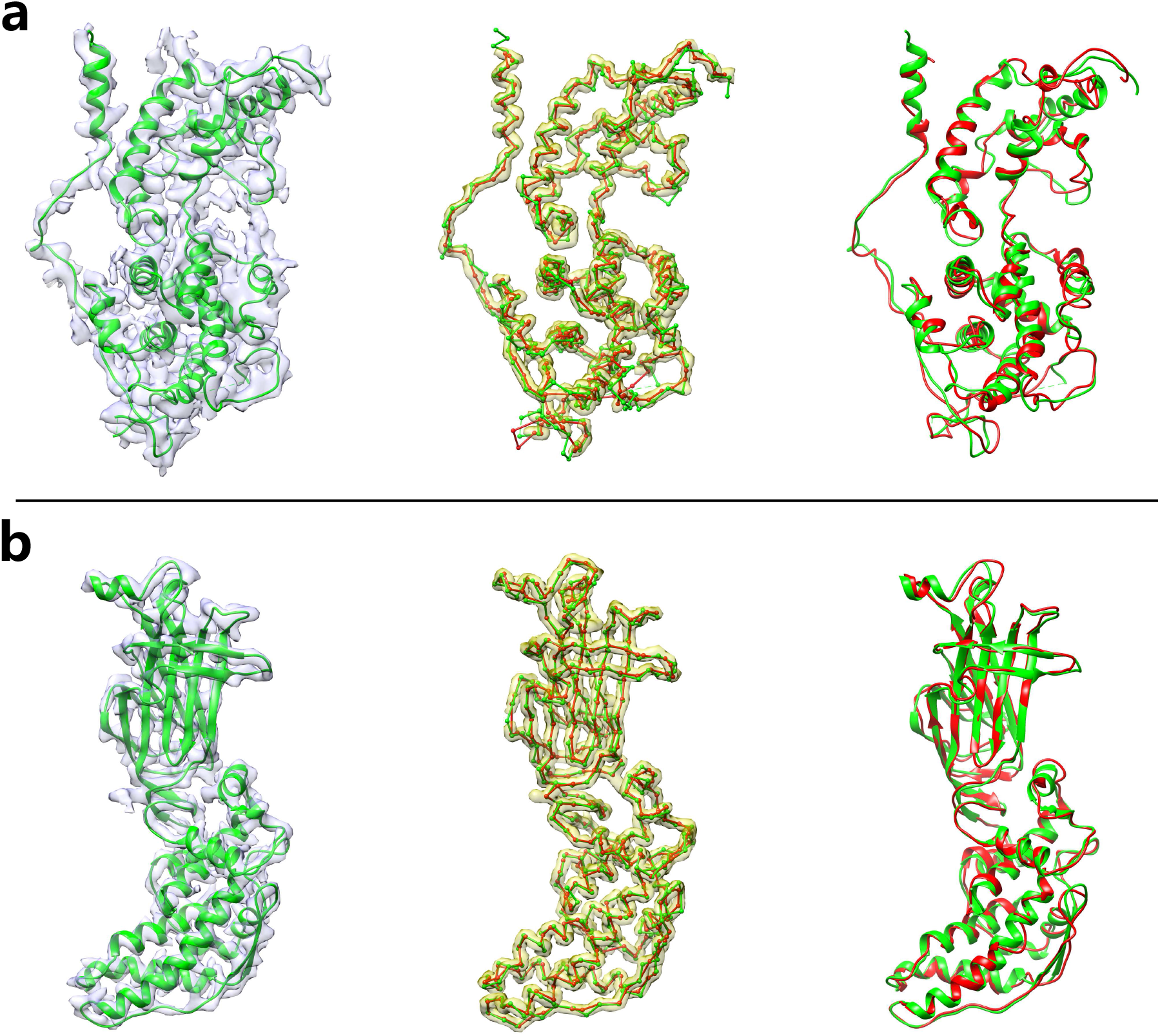
Examples of the models generated by DeepMM for experimental EM maps. The EM density map (transparent grey) and its associated native protein structure (green) are displayed on the left side. The C*α* chains of the DeepMM model (red) and the native structure (green) are shown in ball-and-stick format on the predicted main-chain probability map (transparent yellow) in the middle. The full-atom structure generated by DeepMM (red) and the native protein structure (green) are displayed on the right side. **(a)** The nucleoprotein at 4.3 Å map resolution (EMD-2867). The top ranked model by DeepMM has a C*α* RMSD of 3.1 Å. **(b)** The bovine rotavirus VP6 at 2.6Å map resolution (EMD-6272). The top model by DeepMM has a C*α* RMSD of 1.7 Å.

### 3.3 Evaluation of DeepMM on the EMDB-wide data set

To investigate the accuracy and general applicability of our DeepMM method, we have further evaluated the performance of DeepMM on a large test set of EMDB-wide experimental maps. This large test set consists of 2931 diverse EM maps with 2.6-4.9 Å resolutions from the EMDB that have associated structures in the PDB (See the **Methods** section). For each of the 2931 test cases, our DeepMM method was conducted to reconstruct structures using four combinations of parameters, yielding 640 models for each case. Figure 6 shows a summary of the results predicted by DeepMM. The corresponding data are provided in Supplementary Table 4. Two metrics, RMSD and TMscore, were used to evaluate the overall accuracy of predicted models. On average, DeepMM achieved a C*α* RMSD of 9.8 Å for the top prediction and 8.4 Å for the top 10 predictions on this test set of 2931 maps. The corresponding average TM-scores are 0.648 and 0.694 for top 1 and top 10 predictions, suggesting the high accuracy of our DeepMM approach.

**Figure 6:**
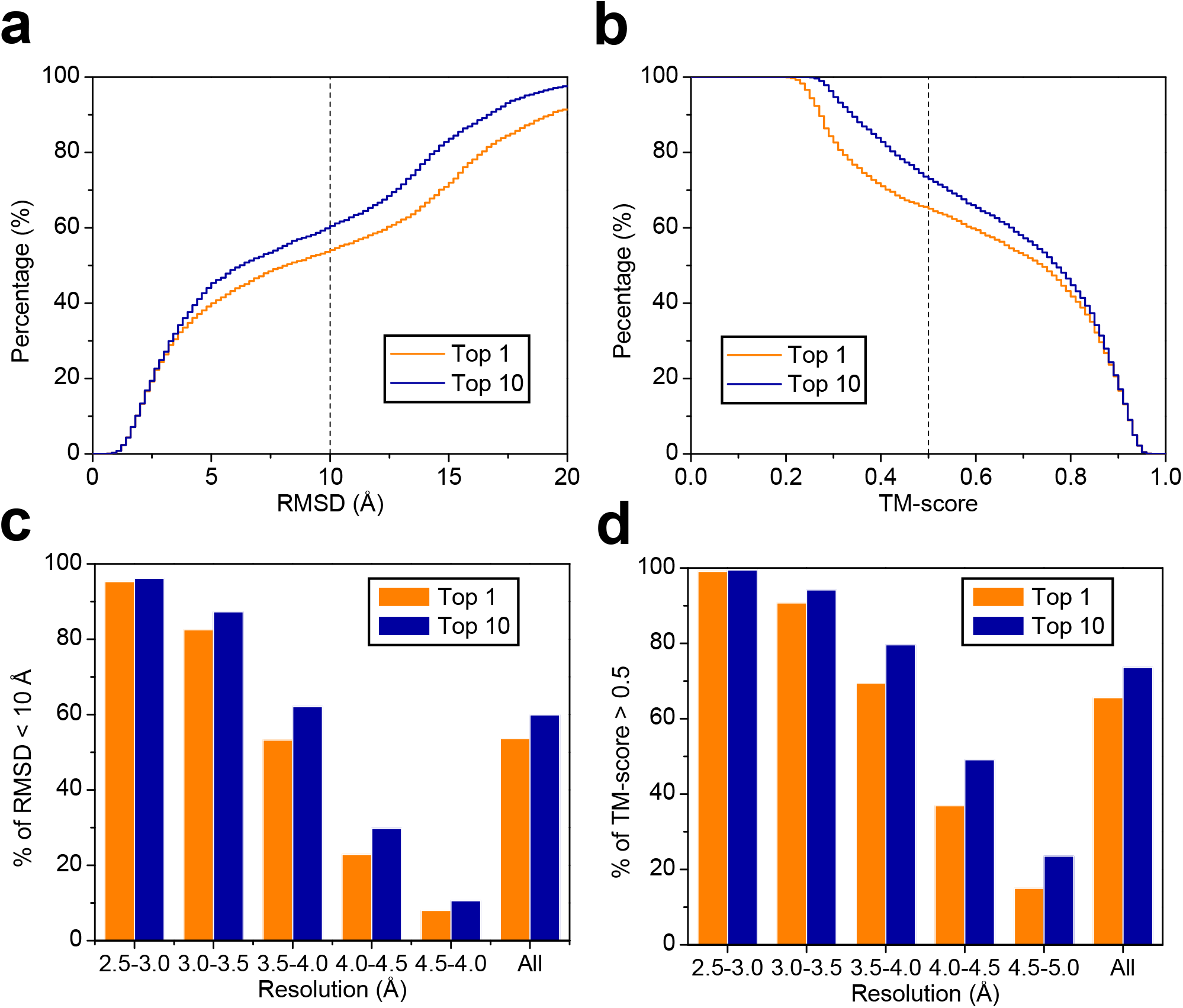
Test results of DeepMM on the 2931 experimental test cases. **(a)** The percentage of the top scored models at different C*α* RMSD cutoffs. **(b)** The percentage of the top scored models at different TM-score cutoffs. **(c)** The percentages of top scored models within 10 Å RMSD in different map resolution ranges. **(d)** The percentages of the top scored models with a TM-score above 0.5 in different map resolution ranges.

Figure 6**a** shows the percentage of the predicted models at different C*α* RMSD cutoffs. It can be seen from the figure that 53.6% of the top models built by DeepMM are within 10 Å C*α* RMSD. For the top 10 scored predictions, 59.9% of the cases have an RMSD of less than 10 Å. The percentage of the models with different TM-score cutoffs are showed in Figure 6**b**. It can be seen from the figure that 65.6% of the top models built by DeepMM have a TM-score of *>* 0.5. When the top 10 models were considered, the corresponding percentage increased to 73.6%. Comparing the results in Figures 6**a** and **b** also reveals that the percentages for TM-score are significantly higher than those for C*α*-RMSD, suggesting that the models built by DeepMM still share the same fold with native structure even if they have a large C*α* RMSD.

Figure 6**c** shows the percentage of correctly predicted top models (i.e. within 10 Å C*α* RMSD) at different resolutions. For EM maps at 2.5-3.0 Å resolution, DeepMM achieved an excellent performance in successfully reconstructing a correct model, and achieved a success rate of 95.3% and 96.2% for the top 1 and 10 scored models, respectively. The performance of DeepMM decreased with the decreasing map resolution. Specifically, for the EM maps with a resolution of 3.0-3.5 Å, 3.5-4.0 Å, and 4.0-4.5 Å, DeepMM obtained a success rate of 82.5%/87.3%, 53.2%/62.1%, and 22.9%/29.8% for the top 1/10 predictions, respectively. For EM maps with a resolution of 4.5 Å or worse, it is challenging for DeepMM to build correct models. On average, for the maps at 3-5 Å resolution, DeepMM gave a success rate of 50.3% and 57.0% in reconstructing a correct model within 10 Å C*α*-RMSD for the top 1 and 10 predictions, respectively. Figure 6**d** shows the percentage of correctly predicted top models using the criterion of TM-score *>* 0.5 in different resolution ranges. Similar trends in Figure 6**c** can be observed in Figure 6**d**. Specifically, for the maps with a resolution of 2.5-3.0 Å, 3.0-3.5 Å, 3.5-4.0 Å, 4.0-4.5 Å, and 4.5-5.0 Å, DeepMM achieved correct models with a TMscore of *>* 0.5 for 99.1%/99.5%, 90.7%/94.2%, 69.5%/79.6%, 36.9%/49.1%, and 15.0%/23.5% of the test cases when the top 1/10 predictions were considered, respectively. On average, for the maps at 3-5 Å resolution, DeepMM obtained a success rate of 63.0% and 71.6% in building a model with TMscore *>* 0.5 for the top 1 and 10 predictions, respectively.

Next, DeepMM was compared with Phenix on this test set, where the Phenix models were generated using the *phenix*.*map to model* tool^28^ in the Phenix package (version 1.18.2-3874). Two metrics calculated by *phenix*.*chain comparison* were used to evaluate the accuracy of a model. One is the fraction of the CA atoms in one model matching the CA atoms in another model within 3.0 Å regardless of their residue names (i.e. coverage or residue match). The other is the percentage of the sequence in the target structure reproduced by the query model (i.e. specificity of sequence match). It should be mentioned that our sequence match is conducted using 20 types of amino acids. A model with a high percentage of residue match may have a very low percentage of sequence match because of mismatching of residue names. Figures 7**a** and **b** show the percentages of protein residues and the sequence reproduced by DeepMM and Phenix at different resolutions. Figures 7**c** and **d** give the histograms of corresponding average values at different resolutions. It can be seen from the figure that DeepMM achieved a significantly better performance than Phenix in both residue match and sequence match, especially for those maps at low resolutions. For the maps at resolutions better than 3.0 Å, 94.2% of protein residues in the deposited structures were reproduced by our DeepMM method, compared to 84.7% by Phenix. The corresponding average sequence match is 78.0% for our DeepMM approach, which is much higher than 59.7% for Phenix. For the maps at 3-5 Å resolution, the average residue match for DeepMM is 80.7%, compared with 65.0% for Phenix. The corresponding average sequence match is 38.1% for DeepMM, which is much higher than 19.2% for Phenix. Given that the prediction of sequence match is much more challenging than that of residue match, the much better performance of DeepMM than Phenix in sequence match demonstrated the atomic-accuracy of the model built by DeepMM.

**Figure 7:**
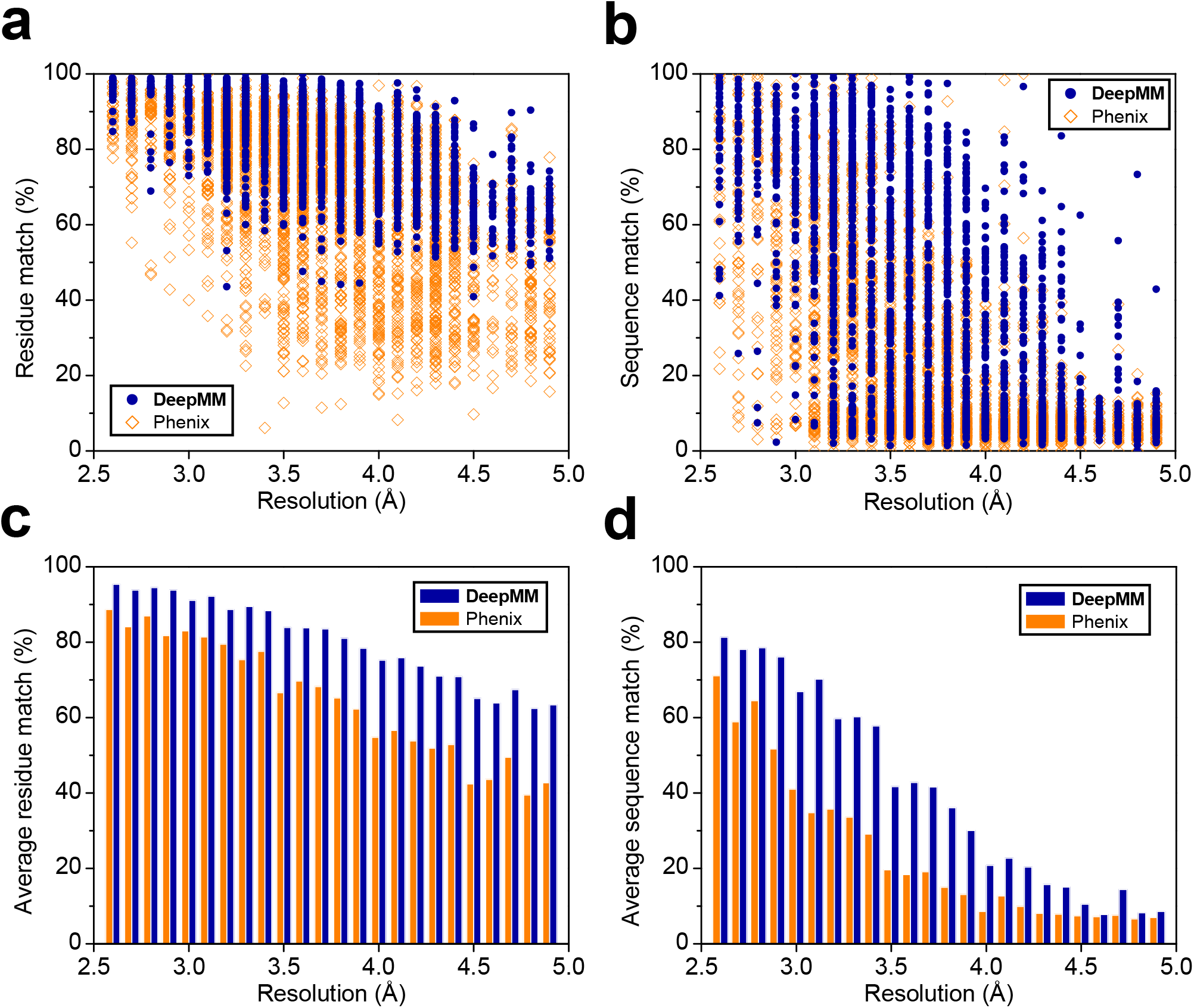
Comparison of the models by DeepMM and Phenix on the large test set of 2931 experimental maps at different resolutions. The results for Phenix are colored in orange, and those for DeepMM are colored in royal blue. **(a)** Percentages of the protein residues in the deposited structures reproduced by DeepMM and Phenix. **(b)** Percentages of the sequence of the deposited structure reproduced by DeepMM and Phenix. **(c)** Average percentage of residue match by DeepMM and Phenix. **(d)** Average percentage of sequence match by DeepMM and Phenix.

It is worth mentioning that DeepMM can build fully-connected, full-length all-atom protein models, whereas Phenix is designed to build initial models of structure fragments. Figure 8 shows the protein models built by DeepMM and Phenix for one example, Chain A of 6DW1, part of a GABAA receptor at 3.1 Å resolution. The deposited structure with its associated EM density map (EMD-8923) is displayed in panel **a**. Figures 8**b** and **c** show the Phenix model and its superimposition with the deposited structure, respectively. It can be seen from the figures that the model built by Phenix consists of multiple fragments without showing any secondary structures, as expected. The predicted model by Phenix for this map had a residue match of 86.7%, but gave a very low sequence match of 9.9%. Therefore, although Phenix recovered most parts of the target protein structure from the EM density map, it assigned wrong residue names for most of the modeled fragments because its low sequence match, as shown in Figure 8**c**. In contrast, DeepMM built an excellent all-atom structure for this map, with a near-perfect residue match of 97.1% and a high sequence match of 86.8%. Therefore, the model predicted by DeepMM reproduced most of the secondary structures and had an almost identical chain trace to the deposited structure(Figure 8**d**). The corresponding amino acid names were also assigned correctly by our DeepMM approach (Figure 8**e**).

**Figure 8:**
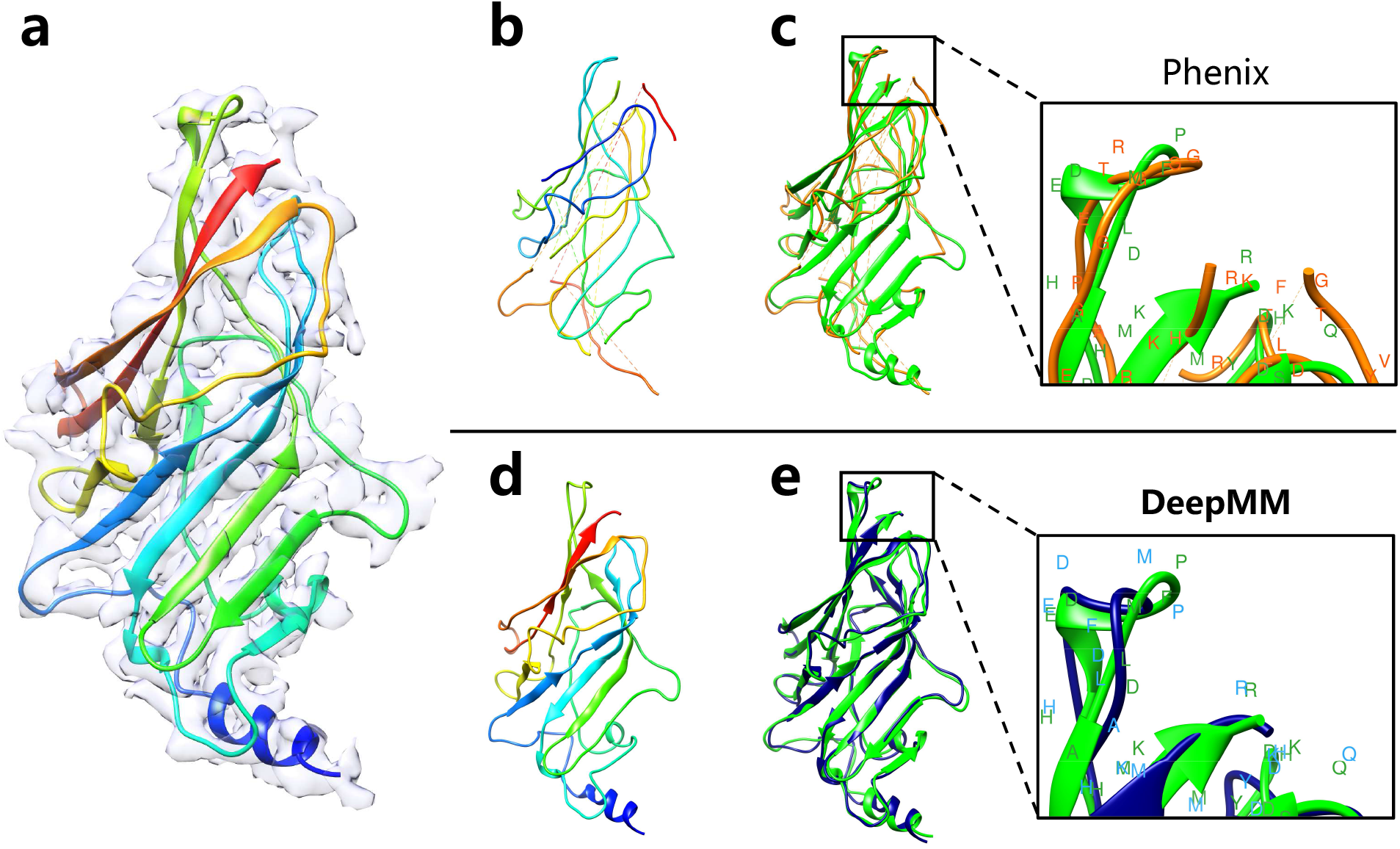
Protein models reconstructed by DeepMM and Phenix for the Chain A of 6DW1 and its associated EM density map at 3.1 Å resolution (EMD-8923). **(a)** The native structure overlapped with its associated EM density map. **(b)** The model predicted by Phenix, which has a residue match of 86.7% and a sequence match of 9.9%. **(c)** The Phenix model (orange) overlapped with the native structure (green). The enlarged box on the right side shows that the residue names assigned by Phenix model are different from those of the native structure. **(d)** The model predicted by Phenix, which has a residue match of 97.1% and a sequence match of 86.8%. **(e)** The DeepMM model (royal blue) overlapped with the native structure (green). The enlarged view of the top region of the protein on the right side shows that the sequence assigned by DeepMM is close to that of the native structure.

## 4 Conclusion

In summary, we have developed a semi-automatic *de novo* structure determination method for nearatomic resolution cryo-EM maps using a deep learning-based framework, named as DeepMM. Our DeepMM approach can reconstruct complete all-atom protein structures for EM maps with atomicaccuracy. DeepMM was extensively validated on diverse benchmarks and compared with state-of-theart approaches including RosettaES, MAINMAST, and Phenix. DeepMM has also been evaluated on an EMDB-wide large test set of 2931 experimental maps at 2.6-4.9 Å resolution. Overall, DeepMM was able reconstruct the protein models with TMscore*>*0.5 for over 60% of the test cases. DeepMM is fast and able to reconstruct an all-atom structure from an EM map within 1 hr on a single-GPU machine for an average-length protein chain of 300 amino acids. Given the high computational efficiency and all-atomic accuracy, it is anticipated that DeepMM will serve as an indispensable tool for semi-automatic atomic-accuracy structure determination for near-atomic-resolution cryo-EM maps.

## Acknowledgements

The authors acknowledge professor Daisuke Kihara and his students Genki Terashi and Sai Raghavendra Maddhuri Venkata Subramaniya from Purdue University for providing their datasets. This work was supported by the National Natural Science Foundation of China (grant Nos. 62072199 and 31670724) and the startup grant of Huazhong University of Science and Technology.

