## Supplementary Table 1 for "Full-length *de novo* protein structure determination from cryo-EM maps using deep learning"

### DenseNet A

growth rate=12, compression=0.5, multiplicative factor=4, number of initial features=32, drop rate=0.2

| Layers | Outout Size | Details | Number of Layers |
|---|---|---|---|
| Convolution | 11×11×11 | 5×5×5 conv, stride 1 | 1 |
| Shared Dense Block 1 | 11×11×11 | BN-ReLU-1×1×1 conv3d-BN-ReLU-3×3×3 conv 3d | 8 |
| Γransition Layer 1 (no pooling | 11×11×11 | BN-ReLU-1×1×1 conv3d | 1 |
| Shared Dense Block 2 | 11×11×11 | BN-ReLU-1×1×1 conv3d-BN-ReLU-3×3×3 conv 3d | 8 |
| Γransition Layer 2 (no pooling | 11×11×11 | BN-ReLU-1×1×1 conv3d | 1 |
| Task Specific Dense Block 1 | 11×11×11 | BN-ReLU-1×1×1 conv3d-BN-ReLU-3×3×3 conv 3d | 8 |
| Transition Layer 1 | 5×5×5 | BN-ReLU-1×1×1 conv3d-2×2 average pool with stride 2 | 1 |
| Task Specific Dense Block 2 | 5×5×5 | BN-ReLU-1×1×1 conv3d-BN-ReLU-3×3×3 conv 3d | 8 |
| Classification Layer | 1 | BN-ReLU- 5×5×5 global pool-FC | 1 |

### DenseNet B

growth rate=12, compression=0.5, multiplicative factor=4, number of initial features=32, drop rate=0.2

| Layers | Outout Size | Details | Number of Layers |
|---|---|---|---|
| Convolution | 11×11×11 | 5×5×5 conv | 1 |
| Shared Dense Block 1 | 11×11×11 | BN-ReLU-1×1×1 conv3d-BN-ReLU-3×3×3 conv 3d | 8 |
| Γransition Layer 1 (no pooling | 11×11×11 | BN-ReLU-1×1×1 conv3d | 1 |
| Task Specific Dense Block 1 | 11×11×11 | BN-ReLU-1×1×1 conv3d-BN-ReLU-3×3×3 conv 3d | 8 |
| Transition Layer 1 | 5×5×5 | BN-ReLU-1×1×1 conv3d-2×2 average pool with stride 2 | 1 |
| Task Specific Dense Block 2 | 5×5×5 | BN-ReLU-1×1×1 conv3d-BN-ReLU-3×3×3 conv 3d | 8 |
| Classification Layer | 3 for AA and 4 for SS | BN-ReLU- 5×5×5 global average pool-FC | 1 |
