## Supplementary Table 2 for "Full-length *de novo* protein structure determination from cryo-EM maps using deep learning"

| PDB id | Number of residues | Cα RMSD (Å) | CLICK-RMSD (Å) | CLICK-SO (%) |
|---|---|---|---|---|
| 1A1X | 106 | 0.43 | 0.43 | 100.00 |
| 1AUU | 55 | 0.83 | 0.82 | 100.00 |
| 1B25 | 209 | 0.49 | 0.49 | 100.00 |
| 1BA3 | 53 | 0.49 | 0.49 | 100.00 |
| 1EJD | 207 | 0.50 | 0.50 | 100.00 |
| 1EWF | 180 | 0.49 | 0.49 | 100.00 |
| 1H12 | 404 | 0.60 | 0.56 | 99.75 |
| 1H70 | 255 | 0.50 | 0.49 | 100.00 |
| 1H8P | 44 | 0.72 | 0.72 | 100.00 |
| 1I5P | 198 | 0.59 | 0.58 | 100.00 |
| 1IGD | 61 | 0.41 | 0.41 | 100.00 |
| 1IWM | 177 | 0.42 | 0.42 | 100.00 |
| 1J0P | 108 | 0.70 | 0.70 | 100.00 |
| 1LKT | 104 | 0.53 | 0.52 | 100.00 |
| 1M3Y | 188 | 0.49 | 0.49 | 100.00 |
| 1MKN | 59 | 0.82 | 0.82 | 100.00 |
| 1N7V | 177 | 0.93 | 0.65 | 99.44 |
| 1OAI | 59 | 0.51 | 0.51 | 100.00 |
| 1OGQ | 313 | 0.43 | 0.43 | 100.00 |
| 1P9H | 159 | 0.51 | 0.51 | 100.00 |
| 1PLQ | 258 | 0.48 | 0.48 | 100.00 |
| 1PPR | 155 | 0.76 | 0.76 | 100.00 |
| 1QSA | 363 | 0.49 | 0.49 | 100.00 |
| 1RG8 | 141 | 0.56 | 0.55 | 100.00 |
| 1TL2 | 235 | 0.50 | 0.50 | 100.00 |
| 1V3W | 173 | 0.57 | 0.57 | 100.00 |
| 1VBW | 68 | 0.41 | 0.41 | 100.00 |
| 1VQ8 | 115 | 0.52 | 0.52 | 100.00 |
| 1W6S | 595 | 0.50 | 0.50 | 100.00 |
| 1WRU | 88 | 0.43 | 0.43 | 100.00 |
| 1YFQ | 342 | 0.46 | 0.46 | 100.00 |
| 2BF6 | 383 | 0.48 | 0.48 | 100.00 |
| 2BMO | 137 | 0.48 | 0.48 | 100.00 |
| 2DPF | 111 | 0.46 | 0.46 | 100.00 |
| 2EIY | 164 | 0.54 | 0.54 | 100.00 |
| 2ERL | 40 | 0.44 | 0.44 | 100.00 |
| 2HBA | 52 | 0.38 | 0.38 | 100.00 |
| 2HNU | 81 | 0.96 | 0.85 | 100.00 |
| 3C7X | 196 | 0.46 | 0.46 | 100.00 |
| 3HMS | 91 | 0.44 | 0.44 | 100.00 |
| | Average | 0.54 | 0.53 | 99.98 |
