## Supplementary Table 3 for "Full-length *de novo* protein structure determination from cryo-EM maps using deep learning"

| EMDB id | Resolution (Å) | Top 1 Cα RMSD (Å) | Top 10 Cα RMSD (Å) | Top 1 all-atom Cα RMSD (Å) | Top 10 all-atom Cα RMSD (Å) |
|---|---|---|---|---|---|
| 1461 | 3.8 | 7.8 | 7.1 | 8.0 | 7.3 |
| 2513A | 3.36 | 2.2 | 2.2 | 2.2 | 2.2 |
| 2513B | 3.36 | 2.4 | 2.4 | 2.5 | 2.5 |
| 2513C | 3.36 | 1.7 | 1.7 | 1.7 | 1.7 |
| 3231 | 3.6 | 5.5 | 5.5 | 5.6 | 5.6 |
| 3246A | 2.8 | 14.8 | 12.9 | 15.2 | 13.3 |
| 3246B | 2.8 | 24.8 | 20.7 | 25.2 | 21.1 |
| 5185 | 3.3 | 3.3 | 3.3 | 3.4 | 3.4 |
| 5495 | 3.5 | 26.8 | 25.5 | 27.0 | 25.7 |
| 5584 | 3.9 | 7.1 | 6.1 | 7.2 | 6.1 |
| 5764 | 3.5 | 12.5 | 12.5 | 12.5 | 12.5 |
| 5778 | 3.275 | 3.9 | 3.9 | 4.1 | 4.1 |
| 5925 | 3.64 | 2.5 | 2.5 | 2.9 | 2.9 |
| 6219 | 4.8 | 17.0 | 15.7 | 17.5 | 16.0 |
| 6272 | 2.6 | 1.7 | 1.7 | 1.7 | 1.7 |
| 6374 | 2.9 | 1.9 | 1.9 | 1.9 | 1.9 |
| 6478 | 2.9 | 4.6 | 4.3 | 4.6 | 4.5 |
| 6551 | 3.8 | 3.7 | 3.7 | 3.8 | 3.8 |
| 6555 | 2.9 | 4.3 | 4.3 | 4.4 | 4.4 |
| 8011 | 3.7 | 4.5 | 4.5 | 4.6 | 4.6 |
| 8015 | 3.1 | 2.8 | 2.8 | 2.8 | 2.8 |
| 8116 | 3.8 | 26.6 | 26.6 | 26.7 | 26.7 |
| 2364 | 4.3 | 28.4 | 28.4 | 28.7 | 28.7 |
| 2850 | 4.3 | 16.7 | 16.7 | 17.1 | 17.1 |
| 2867 | 4.3 | 3.0 | 3.0 | 3.1 | 3.1 |
| 3063 | 4.2 | 22.8 | 18.6 | 23.1 | 18.9 |
| 3073 | 4.1 | 5.0 | 4.0 | 5.3 | 4.2 |
| 3074 | 4.1 | 22.5 | 19.1 | 22.8 | 19.3 |
| 5155 | 4.2 | 20.4 | 20.4 | 20.6 | 20.6 |
| 5376 | 4.1 | 18.6 | 18.4 | 18.8 | 18.7 |
|  | Average | 10.7 | 10.0 | 10.8 | 10.2 |
