## Supplementary Table 4 for "Full-length *de novo* protein structure determination from cryo-EM maps using deep learning"

| PDB id | Chain | Resolution (Å) | EMDB id | Contour level | Pixel spacing (Å) | Number of residues | Top 1 Cα RMSD (Å) | Top 10 C α RMSD (Å) | Top 1 TM-Score | Top 10 TM-Score | DeepMM FOUND (%) | DeepMM SEQ MATCH (%) | Phenix found (%) | Phenix seq match (%) |
|---|---|---|---|---|---|---|---|---|---|---|---|---|---|---|
| 3j7y | 7 | 3.4 | 2762 | 0.0175 | 1.34 | 266 | 7.4 | 4.7 | 0.810 | 0.867 | 87.6 | 42.5 | 83.8 | 15.7 |
| 3j80 | A | 3.8 | 2764 | 0.065 | 1.34 | 206 | 3.6 | 3.6 | 0.861 | 0.861 | 88.3 | 46.7 | 72.3 | 33.6 |
| 3j80 | B | 3.8 | 2764 | 0.065 | 1.34 | 214 | 6.1 | 6.1 | 0.840 | 0.840 | 84.6 | 42 | 68.7 | 20.4 |
| 3j80 | C | 3.8 | 2764 | 0.065 | 1.34 | 217 | 3.9 | 3.9 | 0.881 | 0.881 | 88.5 | 54.7 | 75.6 | 6.1 |
| 3j80 | E | 3.8 | 2764 | 0.065 | 1.34 | 260 | 6.0 | 4.1 | 0.827 | 0.848 | 86.9 | 52.7 | 73.8 | 6.8 |
| 3j80 | H | 3.8 | 2764 | 0.065 | 1.34 | 184 | 10.2 | 5.4 | 0.797 | 0.798 | 81 | 30.2 | 72.8 | 8.2 |
| 3j80 | J | 3.8 | 2764 | 0.065 | 1.34 | 182 | 5.2 | 3.5 | 0.877 | 0.877 | 90.1 | 65.2 | 80.8 | 6.8 |
| 3j80 | K | 3.8 | 2764 | 0.065 | 1.34 | 96 | 14.4 | 12.0 | 0.420 | 0.428 | 65.6 | 12.7 | 63.5 | 13.1 |
| 3j81 | A | 4 | 2763 | 0.065 | 1.34 | 207 | 7.4 | 5.9 | 0.799 | 0.799 | 83.6 | 41.6 | 83.1 | 8.7 |
| 3j81 | B | 4 | 2763 | 0.065 | 1.34 | 215 | 10.9 | 10.1 | 0.665 | 0.690 | 75.3 | 10.5 | 60.5 | 4.6 |
| 3j81 | C | 4 | 2763 | 0.065 | 1.34 | 217 | 8.7 | 8.6 | 0.762 | 0.780 | 81.1 | 30.7 | 77 | 8.4 |
| 3j81 | F | 4 | 2763 | 0.065 | 1.34 | 206 | 5.0 | 5.0 | 0.810 | 0.812 | 82 | 42 | 64.1 | 4.5 |
| 3j81 | H | 4 | 2763 | 0.065 | 1.34 | 184 | 11.9 | 11.9 | 0.588 | 0.601 | 75 | 12.3 | 42.9 | 3.8 |
| 3j81 | K | 4 | 2763 | 0.065 | 1.34 | 96 | 12.8 | 12.6 | 0.365 | 0.373 | 72.9 | 5.7 | 71.9 | 8.7 |
| 3j9i | A | 3.3 | 5623 | 0.25 | 1.2156 | 224 | 1.9 | 1.9 | 0.940 | 0.940 | 95.5 | 91.6 | 76.8 | 40.1 |
| 3j9i | B | 3.3 | 5623 | 0.25 | 1.2156 | 224 | 1.9 | 1.9 | 0.942 | 0.942 | 95.5 | 87.9 | 81.7 | 54.1 |
| 3j9i | C | 3.3 | 5623 | 0.25 | 1.2156 | 224 | 2.6 | 2.6 | 0.924 | 0.924 | 93.8 | 59.5 | 85.3 | 33.5 |
| 3j9i | D | 3.3 | 5623 | 0.25 | 1.2156 | 224 | 7.2 | 2.4 | 0.893 | 0.915 | 91.5 | 62.4 | 87.9 | 51.8 |
| 3j9i | E | 3.3 | 5623 | 0.25 | 1.2156 | 224 | 2.0 | 2.0 | 0.927 | 0.927 | 95.1 | 78.9 | 82.1 | 65.2 |
| 3j9i | F | 3.3 | 5623 | 0.25 | 1.2156 | 224 | 2.8 | 2.8 | 0.912 | 0.912 | 93.8 | 58.1 | 86.2 | 37.8 |
| 3j9i | G | 3.3 | 5623 | 0.25 | 1.2156 | 224 | 2.5 | 2.5 | 0.931 | 0.931 | 95.5 | 71.5 | 82.6 | 38.4 |
| 3j9i | O | 3.3 | 5623 | 0.25 | 1.2156 | 224 | 2.2 | 2.2 | 0.930 | 0.930 | 94.6 | 74.1 | 80.8 | 40.9 |
| 3j9i | P | 3.3 | 5623 | 0.25 | 1.2156 | 224 | 2.1 | 2.1 | 0.935 | 0.935 | 94.6 | 85.8 | 86.2 | 41.5 |
| 3j9i | Q | 3.3 | 5623 | 0.25 | 1.2156 | 224 | 2.1 | 2.1 | 0.929 | 0.929 | 94.6 | 77.8 | 82.1 | 61.4 |
| 3j9i | R | 3.3 | 5623 | 0.25 | 1.2156 | 224 | 3.2 | 3.2 | 0.922 | 0.922 | 93.8 | 53.3 | 81.2 | 45.1 |
| 3j9i | S | 3.3 | 5623 | 0.25 | 1.2156 | 224 | 4.6 | 3.1 | 0.898 | 0.915 | 91.5 | 63.9 | 80.4 | 23.9 |
| 3j9i | T | 3.3 | 5623 | 0.25 | 1.2156 | 224 | 2.9 | 2.9 | 0.923 | 0.923 | 93.8 | 56.2 | 83 | 44.6 |
| 3j9i | U | 3.3 | 5623 | 0.25 | 1.2156 | 224 | 2.4 | 2.4 | 0.931 | 0.931 | 95.5 | 74.3 | 82.1 | 45.1 |
| 3jam | A | 3.5 | 3047 | 0.1 | 1.34 | 208 | 2.1 | 2.1 | 0.941 | 0.941 | 97.1 | 87.1 | 83.7 | 60.9 |
| 3jam | B | 3.5 | 3047 | 0.1 | 1.34 | 223 | 5.2 | 5.2 | 0.904 | 0.904 | 91.9 | 75.6 | 66.8 | 14.8 |
| 3jam | H | 3.5 | 3047 | 0.1 | 1.34 | 184 | 3.3 | 3.3 | 0.885 | 0.893 | 89.7 | 47.3 | 79.9 | 27.9 |
| 3jap | A | 4.9 | 3048 | 0.07 | 1.34 | 208 | 19.8 | 17.3 | 0.260 | 0.461 | 69.2 | 3.5 | 58.2 | 10.7 |
| 3jap | B | 4.9 | 3048 | 0.07 | 1.34 | 222 | 23.0 | 21.3 | 0.331 | 0.359 | 72.1 | 4.4 | 61.3 | 7.4 |

| | | | | | | | | | | | | | | |
|---|---|---|---|---|---|---|---|---|---|---|---|---|---|---|
| 3jap | H | 4.9 | 3048 | 0.07 | 1.34 | 184 | 13.6 | 13.6 | 0.486 | 0.535 | 66.8 | 5.7 | 49.5 | 8.8 |
| 3jb9 | d | 3.6 | 6413 | 0.0203 | 1.32 | 155 | 15.4 | 15.4 | 0.259 | 0.346 | 67.1 | 6.7 | 40 | 6.5 |
| 3jb9 | D | 3.6 | 6413 | 0.0203 | 1.32 | 96 | 2.9 | 2.9 | 0.854 | 0.854 | 91.7 | 77.3 | 72.9 | 2.9 |
| 3jb9 | e | 3.6 | 6413 | 0.0203 | 1.32 | 144 | 3.2 | 3.2 | 0.896 | 0.896 | 93.8 | 58.5 | 74.3 | 9.3 |
| 3jb9 | J | 3.6 | 6413 | 0.0203 | 1.32 | 73 | 13.0 | 9.6 | 0.320 | 0.547 | 75.3 | 27.3 | 65.8 | 22.9 |
| 3jb9 | L | 3.6 | 6413 | 0.0203 | 1.32 | 293 | 12.9 | 11.8 | 0.641 | 0.653 | 71 | 15.4 | 60.8 | 6.2 |
| 3jbr | E | 4.2 | 6475 | 0.035 | 1.32 | 138 | 7.1 | 6.5 | 0.536 | 0.536 | 73.9 | 10.8 | 76.8 | 5.7 |
| 3jc6 | A | 3.7 | 6534 | 0.02 | 1.01 | 208 | 5.1 | 5.1 | 0.828 | 0.828 | 85.6 | 36 | 77.9 | 17.3 |
| 3jca | A | 4.8 | 6441 | 0.5 | 1.31 | 262 | 12.6 | 10.2 | 0.564 | 0.599 | 68.7 | 11.7 | 50 | 4.6 |
| 3jca | B | 4.8 | 6441 | 0.5 | 1.31 | 253 | 19.2 | 15.0 | 0.289 | 0.320 | 59.3 | 4.7 | 44.7 | 2.7 |
| 3jca | E | 4.8 | 6441 | 0.5 | 1.31 | 262 | 21.5 | 17.1 | 0.230 | 0.515 | 65.3 | 4.7 | 58 | 5.9 |
| 3jca | F | 4.8 | 6441 | 0.5 | 1.31 | 253 | 19.4 | 18.4 | 0.286 | 0.368 | 64.4 | 5.5 | 51 | 3.9 |
| 3jck | E | 3.5 | 6479 | 0.0642 | 1.31 | 277 | 5.8 | 5.8 | 0.821 | 0.821 | 85.2 | 42.4 | 80.5 | 23.3 |
| 3jck | G | 3.5 | 6479 | 0.0642 | 1.31 | 257 | 5.3 | 5.3 | 0.864 | 0.864 | 86.4 | 42.8 | 82.1 | 4.3 |
| 3jck | H | 3.5 | 6479 | 0.0642 | 1.31 | 246 | 16.9 | 15.4 | 0.415 | 0.435 | 69.1 | 9.4 | 56.5 | 9.4 |
| 3jcm | L | 3.8 | 6561 | 0.0147 | 1.32 | 139 | 2.3 | 2.3 | 0.889 | 0.889 | 92.1 | 71.1 | 82 | 23.7 |
| 3jcm | M | 3.8 | 6561 | 0.0147 | 1.32 | 126 | 2.8 | 2.8 | 0.921 | 0.921 | 94.4 | 63 | 88.9 | 9.8 |
| 3jcu | n | 3.2 | 6617 | 0.1 | 1.35 | 218 | 3.7 | 3.7 | 0.878 | 0.878 | 89.9 | 51.5 | 59.2 | 42.6 |
| 3jcu | N | 3.2 | 6617 | 0.1 | 1.35 | 218 | 5.8 | 5.2 | 0.852 | 0.852 | 88.5 | 59.1 | 63.3 | 27.5 |
| 3jcu | o | 3.2 | 6617 | 0.1 | 1.35 | 243 | 19.6 | 19.3 | 0.532 | 0.537 | 69.5 | 17.2 | 57.2 | 7.9 |
| 3jcu | O | 3.2 | 6617 | 0.1 | 1.35 | 243 | 23.4 | 19.9 | 0.323 | 0.566 | 68.7 | 16.8 | 56 | 11 |
| 3jcu | y | 3.2 | 6617 | 0.1 | 1.35 | 218 | 2.6 | 2.6 | 0.901 | 0.901 | 94.5 | 76.7 | 75.7 | 34.5 |
| 3jcu | Y | 3.2 | 6617 | 0.1 | 1.35 | 218 | 6.5 | 2.6 | 0.901 | 0.910 | 91.7 | 40 | 74.8 | 56.4 |
| 3jcu | z | 3.2 | 6617 | 0.1 | 1.35 | 61 | 2.1 | 2.1 | 0.803 | 0.803 | 91.8 | 67.9 | 91.8 | 8.9 |
| 3jcu | Z | 3.2 | 6617 | 0.1 | 1.35 | 61 | 3.4 | 3.4 | 0.760 | 0.760 | 90.2 | 56.4 | 88.5 | 42.6 |
| 4v19 | 1 | 3.4 | 2787 | 0.07 | 1.4 | 244 | 2.4 | 2.4 | 0.925 | 0.925 | 93.4 | 75.9 | 79.9 | 16.4 |
| 4v19 | 2 | 3.4 | 2787 | 0.07 | 1.4 | 178 | 1.5 | 1.5 | 0.949 | 0.949 | 97.8 | 89.1 | 86 | 47.7 |
| 4v19 | 3 | 3.4 | 2787 | 0.07 | 1.4 | 118 | 2.3 | 2.3 | 0.902 | 0.902 | 93.2 | 75.5 | 83.1 | 44.9 |
| 4v19 | N | 3.4 | 2787 | 0.07 | 1.4 | 177 | 1.9 | 1.9 | 0.936 | 0.936 | 94.9 | 82.1 | 90.4 | 68.8 |
| 4v19 | O | 3.4 | 2787 | 0.07 | 1.4 | 115 | 12.5 | 6.9 | 0.375 | 0.811 | 84.3 | 21.6 | 80 | 5.4 |
| 4v19 | Q | 3.4 | 2787 | 0.07 | 1.4 | 221 | 4.4 | 4.3 | 0.926 | 0.926 | 95 | 74.3 | 91.4 | 37.1 |
| 4v19 | S | 3.4 | 2787 | 0.07 | 1.4 | 143 | 2.5 | 2.5 | 0.875 | 0.875 | 91.6 | 81.7 | 78.3 | 15.2 |
| 4v19 | T | 3.4 | 2787 | 0.07 | 1.4 | 224 | 2.9 | 2.9 | 0.922 | 0.922 | 92.9 | 80.8 | 67.4 | 32.5 |
| 4v19 | V | 3.4 | 2787 | 0.07 | 1.4 | 155 | 2.1 | 2.1 | 0.932 | 0.932 | 95.5 | 75.7 | 93.5 | 47.6 |
| 4v19 | W | 3.4 | 2787 | 0.07 | 1.4 | 166 | 2.5 | 2.5 | 0.887 | 0.887 | 89.2 | 83.8 | 84.9 | 22 |
| 5a2q | A | 3.9 | 3019 | 0.05 | 1.39 | 216 | 7.3 | 7.3 | 0.813 | 0.826 | 90.3 | 74.9 | 80.6 | 29.9 |
| 5a2q | B | 3.9 | 3019 | 0.05 | 1.39 | 213 | 9.3 | 5.7 | 0.846 | 0.879 | 92 | 68.9 | 75.6 | 14.3 |

| | | | | | | | | | | | | | | |
|---|---|---|---|---|---|---|---|---|---|---|---|---|---|---|
| 5a2q | C | 3.9 | 3019 | 0.05 | 1.39 | 218 | 2.2 | 2.2 | 0.926 | 0.926 | 94.5 | 81.6 | 80.3 | 24.6 |
| 5a2q | E | 3.9 | 3019 | 0.05 | 1.39 | 262 | 2.5 | 2.5 | 0.933 | 0.933 | 94.7 | 76.2 | 79.8 | 7.2 |
| 5a2q | H | 3.9 | 3019 | 0.05 | 1.39 | 186 | 7.6 | 4.1 | 0.852 | 0.864 | 87.6 | 44.8 | 78 | 21.4 |
| 5a2q | J | 3.9 | 3019 | 0.05 | 1.39 | 180 | 1.9 | 1.9 | 0.913 | 0.913 | 95 | 81.9 | 80.6 | 46.2 |
| 5a2q | Y | 3.9 | 3019 | 0.05 | 1.39 | 124 | 2.6 | 2.6 | 0.881 | 0.881 | 93.5 | 69 | 85.5 | 34.9 |
| 5aco | G | 4.4 | 3121 | 0.041 | 1.31 | 132 | 16.3 | 14.9 | 0.256 | 0.375 | 63.6 | 4.8 | 24.2 | 9.4 |
| 5aco | H | 4.4 | 3121 | 0.041 | 1.31 | 132 | 16.8 | 14.2 | 0.411 | 0.432 | 64.4 | 5.9 | 27.3 | 19.4 |
| 5aco | I | 4.4 | 3121 | 0.041 | 1.31 | 132 | 15.9 | 15.0 | 0.373 | 0.417 | 64.4 | 18.8 | 23.5 | 3.2 |
| 5aco | J | 4.4 | 3121 | 0.041 | 1.31 | 103 | 15.2 | 14.0 | 0.323 | 0.436 | 63.1 | 13.8 | 27.2 | 14.3 |
| 5aco | K | 4.4 | 3121 | 0.041 | 1.31 | 103 | 15.8 | 14.0 | 0.259 | 0.287 | 62.1 | 4.7 | 57.3 | 6.8 |
| 5aco | L | 4.4 | 3121 | 0.041 | 1.31 | 103 | 14.0 | 12.7 | 0.373 | 0.373 | 64.1 | 7.6 | 43.7 | 17.8 |
| 5adx | K | 4 | 2856 | 0.088 | 1.34 | 275 | 16.1 | 15.4 | 0.549 | 0.648 | 72.4 | 22.1 | 51.3 | 5 |
| 5adx | L | 4 | 2856 | 0.088 | 1.34 | 270 | 15.4 | 13.2 | 0.734 | 0.743 | 78.1 | 11.4 | 65.6 | 4.5 |
| 5adx | V | 4 | 2856 | 0.088 | 1.34 | 165 | 16.5 | 13.4 | 0.286 | 0.425 | 69.1 | 9.6 | 47.3 | 7.7 |
| 5ady | G | 4.5 | 3133 | 1.4 | 1.1659 | 176 | 10.8 | 10.8 | 0.528 | 0.528 | 64.2 | 12.4 | 65.3 | 8.7 |
| 5ady | V | 4.5 | 3133 | 1.4 | 1.1659 | 94 | 12.5 | 10.9 | 0.385 | 0.464 | 70.2 | 10.6 | 46.8 | 2.3 |
| 5aj3 | B | 3.6 | 2913 | 0.06 | 1.39 | 220 | 8.1 | 8.1 | 0.670 | 0.729 | 90.9 | 49.5 | 87.7 | 53.9 |
| 5aj3 | c | 3.6 | 2913 | 0.06 | 1.39 | 169 | 2.9 | 2.9 | 0.915 | 0.915 | 95.3 | 55.3 | 71.6 | 6.6 |
| 5aj3 | f | 3.6 | 2913 | 0.06 | 1.39 | 99 | 5.2 | 5.2 | 0.798 | 0.798 | 85.9 | 58.8 | 75.8 | 10.7 |
| 5aj3 | F | 3.6 | 2913 | 0.06 | 1.39 | 123 | 5.2 | 3.1 | 0.838 | 0.846 | 86.2 | 51.9 | 78 | 12.5 |
| 5aj3 | G | 3.6 | 2913 | 0.06 | 1.39 | 208 | 3.4 | 3.4 | 0.899 | 0.899 | 92.3 | 59.4 | 73.1 | 47.4 |
| 5aj3 | h | 3.6 | 2913 | 0.06 | 1.39 | 103 | 3.3 | 3.3 | 0.851 | 0.851 | 91.3 | 56.4 | 72.8 | 58.7 |
| 5aj3 | k | 3.6 | 2913 | 0.06 | 1.39 | 275 | 23.1 | 3.5 | 0.666 | 0.866 | 76 | 12.9 | 73.1 | 16.9 |
| 5aj3 | p | 3.6 | 2913 | 0.06 | 1.39 | 188 | 4.5 | 4.5 | 0.875 | 0.882 | 89.4 | 68.5 | 72.9 | 33.6 |
| 5aj3 | Q | 3.6 | 2913 | 0.06 | 1.39 | 109 | 3.4 | 3.4 | 0.853 | 0.853 | 89.9 | 69.4 | 70.6 | 11.7 |
| 5fj8 | E | 3.9 | 3178 | 0.06 | 1.084 | 215 | 21.3 | 20.2 | 0.261 | 0.348 | 73 | 10.2 | 48.4 | 3.8 |
| 5fj8 | H | 3.9 | 3178 | 0.06 | 1.084 | 140 | 18.1 | 15.5 | 0.259 | 0.430 | 81.4 | 7.9 | 43.6 | 9.8 |
| 5fj9 | E | 4.6 | 3179 | 0.055 | 1.084 | 215 | 16.7 | 16.7 | 0.393 | 0.431 | 70.7 | 11.2 | 54 | 5.2 |
| 5fj9 | H | 4.6 | 3179 | 0.055 | 1.084 | 140 | 15.4 | 14.4 | 0.237 | 0.390 | 65.7 | 8.7 | 54.3 | 7.9 |
| 5fja | E | 4.7 | 3180 | 0.055 | 1.084 | 215 | 21.9 | 20.9 | 0.236 | 0.311 | 63.3 | 7.4 | 54.4 | 4.3 |
| 5fja | H | 4.7 | 3180 | 0.055 | 1.084 | 140 | 17.2 | 15.2 | 0.292 | 0.314 | 62.9 | 10.2 | 34.3 | 2.1 |
| 5flm | C | 3.4 | 3218 | 0.07 | 1.35 | 257 | 6.6 | 6.3 | 0.898 | 0.898 | 90.7 | 73.8 | 83.7 | 5.1 |
| 5flm | E | 3.4 | 3218 | 0.07 | 1.35 | 209 | 4.0 | 3.1 | 0.909 | 0.909 | 93.8 | 74.5 | 77 | 17.4 |
| 5flm | H | 3.4 | 3218 | 0.07 | 1.35 | 148 | 2.0 | 2.0 | 0.926 | 0.926 | 95.9 | 85.2 | 77 | 8.8 |
| 5fmg | B | 3.6 | 3231 | 3 | 1.04 | 212 | 17.9 | 16.3 | 0.374 | 0.658 | 73.6 | 3.8 | 75.9 | 9.3 |
| 5fmg | C | 3.6 | 3231 | 3 | 1.04 | 220 | 14.7 | 14.6 | 0.513 | 0.531 | 75 | 23 | 75 | 25.5 |
| 5fmg | E | 3.6 | 3231 | 3 | 1.04 | 217 | 17.8 | 17.1 | 0.315 | 0.500 | 77.9 | 5.9 | 61.8 | 7.5 |

| | | | | | | | | | | | | | | |
|---|---|---|---|---|---|---|---|---|---|---|---|---|---|---|
| 5fmg | F | 3.6 | 3231 | 3 | 1.04 | 219 | 13.3 | 13.1 | 0.412 | 0.422 | 74.9 | 26.8 | 78.1 | 5.3 |
| 5fmg | K | 3.6 | 3231 | 3 | 1.04 | 194 | 5.8 | 5.8 | 0.819 | 0.819 | 83 | 27.3 | 71.1 | 3.6 |
| 5fmg | P | 3.6 | 3231 | 3 | 1.04 | 212 | 18.7 | 16.8 | 0.306 | 0.614 | 74.1 | 11.5 | 82.5 | 14.3 |
| 5fmg | Q | 3.6 | 3231 | 3 | 1.04 | 220 | 16.7 | 10.4 | 0.610 | 0.684 | 81.4 | 15.1 | 74.5 | 17.7 |
| 5fmg | S | 3.6 | 3231 | 3 | 1.04 | 217 | 18.1 | 16.7 | 0.350 | 0.655 | 81.1 | 8.5 | 73.3 | 6.9 |
| 5fmg | T | 3.6 | 3231 | 3 | 1.04 | 219 | 19.2 | 17.6 | 0.366 | 0.371 | 75.8 | 7.8 | 75.3 | 4.8 |
| 5fmg | Y | 3.6 | 3231 | 3 | 1.04 | 194 | 6.0 | 6.0 | 0.773 | 0.777 | 82 | 22 | 74.2 | 9 |
| 5fn3 | D | 4.1 | 3238 | 0.04 | 1.4 | 100 | 5.7 | 3.4 | 0.751 | 0.780 | 82 | 32.9 | 84 | 22.6 |
| 5fn4 | D | 4 | 3239 | 0.04 | 1.4 | 100 | 4.7 | 4.7 | 0.736 | 0.759 | 84 | 41.7 | 71 | 5.6 |
| 5fuu | H | 4.2 | 3308 | 0.034 | 1.31 | 135 | 14.3 | 12.5 | 0.397 | 0.746 | 76.3 | 23.3 | 32.6 | 4.5 |
| 5fuu | L | 4.2 | 3308 | 0.034 | 1.31 | 114 | 16.9 | 13.7 | 0.272 | 0.395 | 74.6 | 11.8 | 43.9 | 24 |
| 5fuu | M | 4.2 | 3308 | 0.034 | 1.31 | 135 | 17.7 | 16.2 | 0.277 | 0.601 | 74.8 | 6.9 | 63 | 8.2 |
| 5fuu | N | 4.2 | 3308 | 0.034 | 1.31 | 112 | 15.0 | 13.9 | 0.287 | 0.398 | 68.8 | 7.8 | 34.8 | 12.8 |
| 5fyw | C | 4.4 | 3378 | 0.0325 | 1.35 | 262 | 28.3 | 23.7 | 0.344 | 0.400 | 75.6 | 12.1 | 72.5 | 6.3 |
| 5fyw | D | 4.4 | 3378 | 0.0325 | 1.35 | 157 | 18.2 | 13.6 | 0.249 | 0.389 | 56.1 | 5.7 | 50.3 | 3.8 |
| 5fyw | E | 4.4 | 3378 | 0.0325 | 1.35 | 213 | 14.1 | 13.9 | 0.491 | 0.545 | 76.1 | 9.9 | 39.4 | 6 |
| 5fyw | F | 4.4 | 3378 | 0.0325 | 1.35 | 83 | 8.1 | 8.1 | 0.612 | 0.627 | 68.7 | 35.1 | 36.1 | 6.7 |
| 5fyw | G | 4.4 | 3378 | 0.0325 | 1.35 | 171 | 17.2 | 14.3 | 0.327 | 0.341 | 66.7 | 5.3 | 29.8 | 7.8 |
| 5fyw | H | 4.4 | 3378 | 0.0325 | 1.35 | 136 | 16.2 | 15.7 | 0.259 | 0.273 | 65.4 | 3.4 | 36.8 | 8 |
| 5fyw | M | 4.4 | 3378 | 0.0325 | 1.35 | 231 | 21.5 | 20.4 | 0.362 | 0.566 | 63.2 | 3.4 | 74.5 | 8.1 |
| 5g5l | E | 4.8 | 3439 | 0.023 | 1.35 | 212 | 20.2 | 13.5 | 0.272 | 0.641 | 74.1 | 7.6 | 31.1 | 9.1 |
| 5g5l | G | 4.8 | 3439 | 0.023 | 1.35 | 193 | 26.1 | 17.8 | 0.287 | 0.311 | 62.2 | 10.8 | 38.9 | 4 |
| 5g5l | H | 4.8 | 3439 | 0.023 | 1.35 | 131 | 17.6 | 14.0 | 0.256 | 0.278 | 62.6 | 8.5 | 44.3 | 3.4 |
| 5gad | F | 3.7 | 8000 | 0.02 | 1.385 | 177 | 8.5 | 4.8 | 0.815 | 0.825 | 84.7 | 18 | 47.5 | 9.5 |
| 5gad | G | 3.7 | 8000 | 0.02 | 1.385 | 176 | 23.2 | 9.1 | 0.258 | 0.618 | 69.9 | 13 | 67 | 5.9 |
| 5gad | K | 3.7 | 8000 | 0.02 | 1.385 | 142 | 5.9 | 4.6 | 0.793 | 0.793 | 84.5 | 42.5 | 75.4 | 15 |
| 5gad | P | 3.7 | 8000 | 0.02 | 1.385 | 117 | 3.8 | 3.8 | 0.812 | 0.813 | 88.9 | 59.6 | 83.8 | 21.4 |
| 5gad | W | 3.7 | 8000 | 0.02 | 1.385 | 94 | 4.4 | 4.4 | 0.812 | 0.828 | 89.4 | 25 | 66 | 14.5 |
| 5gae | a | 3.3 | 8001 | 0.03 | 1.385 | 58 | 2.2 | 2.2 | 0.853 | 0.853 | 96.6 | 75 | 82.8 | 68.8 |
| 5gae | E | 3.3 | 8001 | 0.03 | 1.385 | 201 | 2.5 | 2.5 | 0.919 | 0.919 | 94.5 | 77.4 | 84.6 | 55.3 |
| 5gae | K | 3.3 | 8001 | 0.03 | 1.385 | 142 | 1.5 | 1.5 | 0.933 | 0.933 | 97.2 | 91.3 | 74.6 | 6.6 |
| 5gae | L | 3.3 | 8001 | 0.03 | 1.385 | 123 | 3.3 | 3.3 | 0.886 | 0.886 | 92.7 | 53.5 | 83.7 | 7.8 |
| 5gae | N | 3.3 | 8001 | 0.03 | 1.385 | 136 | 1.6 | 1.6 | 0.929 | 0.929 | 96.3 | 84.7 | 86 | 30.8 |
| 5gae | O | 3.3 | 8001 | 0.03 | 1.385 | 125 | 2.1 | 2.1 | 0.899 | 0.899 | 92 | 78.3 | 84 | 47.6 |
| 5gae | P | 3.3 | 8001 | 0.03 | 1.385 | 117 | 4.3 | 4.3 | 0.808 | 0.829 | 85.5 | 48 | 69.2 | 11.1 |
| 5gae | Q | 3.3 | 8001 | 0.03 | 1.385 | 114 | 2.0 | 2.0 | 0.898 | 0.898 | 93.9 | 77.6 | 72.8 | 21.7 |
| 5gae | S | 3.3 | 8001 | 0.03 | 1.385 | 103 | 3.1 | 3.1 | 0.868 | 0.868 | 93.2 | 42.7 | 76.7 | 7.6 |

| | | | | | | | | | | | | | | |
|---|---|---|---|---|---|---|---|---|---|---|---|---|---|---|
| 5gae | T | 3.3 | 8001 | 0.03 | 1.385 | 110 | 3.5 | 3.5 | 0.886 | 0.886 | 92.7 | 63.7 | 85.5 | 5.3 |
| 5gae | U | 3.3 | 8001 | 0.03 | 1.385 | 95 | 1.7 | 1.7 | 0.884 | 0.884 | 94.7 | 86.7 | 55.8 | 3.8 |
| 5gae | V | 3.3 | 8001 | 0.03 | 1.385 | 102 | 4.1 | 4.1 | 0.775 | 0.775 | 81.4 | 51.8 | 77.5 | 3.8 |
| 5gae | W | 3.3 | 8001 | 0.03 | 1.385 | 94 | 2.8 | 2.8 | 0.878 | 0.878 | 93.6 | 73.9 | 62.8 | 10.2 |
| 5gae | Z | 3.3 | 8001 | 0.03 | 1.385 | 62 | 2.8 | 2.8 | 0.822 | 0.822 | 90.3 | 64.3 | 74.2 | 63 |
| 5gag | F | 3.8 | 8003 | 0.032 | 1.385 | 178 | 4.1 | 4.1 | 0.846 | 0.846 | 89.3 | 37.3 | 58.8 | 20.2 |
| 5gag | G | 3.8 | 8003 | 0.032 | 1.385 | 176 | 16.3 | 15.4 | 0.516 | 0.516 | 85.2 | 16.7 | 60.2 | 17 |
| 5gag | K | 3.8 | 8003 | 0.032 | 1.385 | 142 | 2.2 | 2.2 | 0.881 | 0.881 | 92.3 | 73.3 | 85.9 | 6.6 |
| 5gag | P | 3.8 | 8003 | 0.032 | 1.385 | 117 | 3.1 | 3.1 | 0.816 | 0.816 | 90.6 | 51.9 | 81.2 | 8.4 |
| 5gag | V | 3.8 | 8003 | 0.032 | 1.385 | 103 | 4.3 | 4.3 | 0.743 | 0.758 | 81.4 | 56.6 | 58.8 | 6.7 |
| 5gag | W | 3.8 | 8003 | 0.032 | 1.385 | 94 | 4.3 | 4.3 | 0.848 | 0.848 | 90.4 | 49.4 | 76.6 | 6.9 |
| 5gag | Z | 3.8 | 8003 | 0.032 | 1.385 | 62 | 3.2 | 3.2 | 0.798 | 0.822 | 88.7 | 49.1 | 88.7 | 92.7 |
| 5gah | G | 3.8 | 8004 | 0.05 | 1.385 | 176 | 5.9 | 5.9 | 0.790 | 0.790 | 78.4 | 30.4 | 69.3 | 9 |
| 5gah | K | 3.8 | 8004 | 0.05 | 1.385 | 142 | 2.7 | 2.7 | 0.863 | 0.863 | 90.8 | 62.8 | 80.3 | 3.5 |
| 5gah | P | 3.8 | 8004 | 0.05 | 1.385 | 117 | 2.8 | 2.8 | 0.864 | 0.864 | 91.5 | 57.9 | 79.5 | 14 |
| 5gah | V | 3.8 | 8004 | 0.05 | 1.385 | 102 | 5.3 | 3.1 | 0.809 | 0.841 | 88.2 | 50 | 68.6 | 2.9 |
| 5gah | W | 3.8 | 8004 | 0.05 | 1.385 | 94 | 4.3 | 4.3 | 0.861 | 0.861 | 95.7 | 57.8 | 62.8 | 5.1 |
| 5gah | Z | 3.8 | 8004 | 0.05 | 1.385 | 62 | 2.0 | 2.0 | 0.825 | 0.825 | 91.9 | 71.9 | 91.9 | 57.9 |
| 5gam | d | 3.7 | 8011 | 0.012 | 1.43 | 82 | 4.4 | 4.4 | 0.838 | 0.838 | 92.7 | 15.8 | 69.5 | 1.8 |
| 5gan | D | 3.7 | 8012 | 0.036 | 1.43 | 140 | 3.7 | 3.7 | 0.855 | 0.855 | 89.3 | 60.8 | 72.1 | 10.9 |
| 5gan | K | 3.7 | 8012 | 0.036 | 1.43 | 124 | 2.5 | 2.5 | 0.917 | 0.917 | 95.2 | 70.3 | 88.7 | 26.4 |
| 5gao | A | 4.2 | 8013 | 0.02 | 1.43 | 255 | 18.5 | 18.3 | 0.374 | 0.427 | 75.7 | 25.4 | 50.6 | 5.4 |
| 5gap | D | 3.6 | 8014 | 0.02 | 1.43 | 140 | 2.7 | 2.7 | 0.876 | 0.876 | 92.9 | 66.2 | 80 | 5.4 |
| 5gap | K | 3.6 | 8014 | 0.02 | 1.43 | 124 | 2.7 | 2.7 | 0.910 | 0.910 | 92.7 | 67.8 | 86.3 | 63.6 |
| 5gaq | A | 3.1 | 8015 | 0.07 | 1.43 | 288 | 2.8 | 2.8 | 0.910 | 0.918 | 94.4 | 74.6 | 89.6 | 20.9 |
| 5gaq | B | 3.1 | 8015 | 0.07 | 1.43 | 288 | 7.4 | 7.4 | 0.839 | 0.890 | 90.6 | 57.5 | 35.8 | 4.9 |
| 5gaq | C | 3.1 | 8015 | 0.07 | 1.43 | 288 | 5.2 | 5.0 | 0.919 | 0.920 | 93.8 | 63.3 | 87.8 | 13 |
| 5gaq | D | 3.1 | 8015 | 0.07 | 1.43 | 288 | 4.0 | 4.0 | 0.927 | 0.927 | 94.4 | 69.1 | 86.8 | 10.8 |
| 5gaq | E | 3.1 | 8015 | 0.07 | 1.43 | 288 | 2.1 | 2.1 | 0.945 | 0.945 | 96.5 | 82 | 90.6 | 66.3 |
| 5gaq | F | 3.1 | 8015 | 0.07 | 1.43 | 288 | 2.3 | 2.3 | 0.927 | 0.927 | 94.1 | 85.6 | 91 | 34.7 |
| 5gaq | G | 3.1 | 8015 | 0.07 | 1.43 | 288 | 2.2 | 2.2 | 0.943 | 0.943 | 95.5 | 82.9 | 89.9 | 55.2 |
| 5gaq | H | 3.1 | 8015 | 0.07 | 1.43 | 288 | 34.0 | 11.6 | 0.638 | 0.858 | 87.2 | 30.7 | 88.2 | 18.5 |
| 5gaq | I | 3.1 | 8015 | 0.07 | 1.43 | 288 | 2.5 | 2.5 | 0.936 | 0.936 | 94.8 | 63.7 | 89.6 | 5.8 |
| 5gjq | B | 4.4 | 9511 | 0.033 | 1.07 | 244 | 19.3 | 17.3 | 0.363 | 0.449 | 67.2 | 9.1 | 61.5 | 6.7 |
| 5gjq | C | 4.4 | 9511 | 0.033 | 1.07 | 233 | 14.4 | 14.4 | 0.511 | 0.511 | 73 | 17.1 | 53.6 | 7.2 |
| 5gjq | d | 4.4 | 9511 | 0.033 | 1.07 | 199 | 18.1 | 15.4 | 0.274 | 0.402 | 73.4 | 6.2 | 70.9 | 7.8 |
| 5gjq | D | 4.4 | 9511 | 0.033 | 1.07 | 250 | 18.3 | 17.1 | 0.295 | 0.410 | 73.6 | 11.4 | 71.2 | 17.4 |

| | | | | | | | | | | | | | | |
|---|---|---|---|---|---|---|---|---|---|---|---|---|---|---|
| 5gjq | E | 4.4 | 9511 | 0.033 | 1.07 | 243 | 17.2 | 15.8 | 0.558 | 0.558 | 72.8 | 14.1 | 58 | 8.5 |
| 5gjq | F | 4.4 | 9511 | 0.033 | 1.07 | 234 | 17.2 | 16.3 | 0.387 | 0.520 | 76.1 | 4.5 | 60.3 | 9.2 |
| 5gjq | G | 4.4 | 9511 | 0.033 | 1.07 | 238 | 13.6 | 13.0 | 0.560 | 0.568 | 69.3 | 17.6 | 58 | 9.4 |
| 5gjq | h | 4.4 | 9511 | 0.033 | 1.07 | 244 | 18.4 | 16.7 | 0.427 | 0.560 | 74.2 | 16.6 | 69.3 | 8.9 |
| 5gjq | i | 4.4 | 9511 | 0.033 | 1.07 | 231 | 18.5 | 15.9 | 0.324 | 0.491 | 77.9 | 10.6 | 72.7 | 5.4 |
| 5gjq | j | 4.4 | 9511 | 0.033 | 1.07 | 250 | 11.1 | 9.9 | 0.615 | 0.680 | 76 | 20.5 | 68.4 | 8.2 |
| 5gjq | k | 4.4 | 9511 | 0.033 | 1.07 | 243 | 18.2 | 17.8 | 0.331 | 0.347 | 71.2 | 8.7 | 53.9 | 6.1 |
| 5gjq | l | 4.4 | 9511 | 0.033 | 1.07 | 234 | 20.2 | 13.2 | 0.364 | 0.580 | 73.1 | 5.3 | 71.4 | 9 |
| 5gjq | m | 4.4 | 9511 | 0.033 | 1.07 | 238 | 20.9 | 18.7 | 0.400 | 0.453 | 72.3 | 22.1 | 60.9 | 3.4 |
| 5gjq | r | 4.4 | 9511 | 0.033 | 1.07 | 199 | 12.0 | 10.3 | 0.431 | 0.611 | 77.9 | 16.8 | 67.8 | 9.6 |
| 5gmk | Q | 3.4 | 9525 | 0.0413 | 1.306 | 185 | 20.2 | 20.2 | 0.392 | 0.635 | 63.2 | 12.8 | 63.8 | 6.8 |
| 5gmk | R | 3.4 | 9525 | 0.0413 | 1.306 | 261 | 18.1 | 15.1 | 0.632 | 0.770 | 85.1 | 31.1 | 72 | 14.4 |
| 5gmk | T | 3.4 | 9525 | 0.0413 | 1.306 | 157 | 1.8 | 1.8 | 0.916 | 0.916 | 94.3 | 83.8 | 77.1 | 45.5 |
| 5h1s | I | 3.5 | 9572 | 0.132 | 1.28 | 173 | 8.3 | 5.8 | 0.846 | 0.846 | 91.9 | 53.5 | 75.7 | 19.1 |
| 5irx | E | 3 | 8117 | 3.5 | 1.2156 | 75 | 1.8 | 1.8 | 0.835 | 0.835 | 93.3 | 80 | 40 | 6.7 |
| 5irx | F | 3 | 8117 | 3.5 | 1.2156 | 75 | 8.0 | 4.3 | 0.774 | 0.781 | 86.7 | 38.5 | 53.3 | 12.5 |
| 5iyb | C | 3.9 | 8136 | 0.01 | 1.31 | 275 | 18.7 | 18.0 | 0.678 | 0.797 | 78.2 | 17.7 | 71.6 | 8.6 |
| 5iyb | D | 3.9 | 8136 | 0.01 | 1.31 | 129 | 19.3 | 7.8 | 0.268 | 0.568 | 60.5 | 7.7 | 45 | 3.4 |
| 5iyb | E | 3.9 | 8136 | 0.01 | 1.31 | 210 | 12.0 | 9.1 | 0.739 | 0.750 | 72.4 | 29.6 | 71.9 | 15.9 |
| 5iyb | H | 3.9 | 8136 | 0.01 | 1.31 | 150 | 16.4 | 15.0 | 0.293 | 0.503 | 67.3 | 7.9 | 60.7 | 6.6 |
| 5iyc | C | 3.9 | 8137 | 0.01 | 1.31 | 275 | 13.9 | 10.3 | 0.788 | 0.791 | 78.2 | 23.3 | 62.5 | 6.4 |
| 5iyc | D | 3.9 | 8137 | 0.01 | 1.31 | 129 | 16.4 | 12.8 | 0.318 | 0.406 | 58.1 | 9.3 | 40.3 | 3.8 |
| 5iyc | E | 3.9 | 8137 | 0.01 | 1.31 | 210 | 13.7 | 9.5 | 0.576 | 0.684 | 77.1 | 19.1 | 71 | 7.4 |
| 5iyc | H | 3.9 | 8137 | 0.01 | 1.31 | 150 | 19.9 | 14.4 | 0.295 | 0.371 | 66 | 10.1 | 40 | 6.7 |
| 5iyd | C | 3.9 | 8138 | 0.01 | 1.31 | 275 | 4.5 | 4.5 | 0.860 | 0.860 | 86.2 | 51.1 | 63.3 | 4 |
| 5iyd | D | 3.9 | 8138 | 0.01 | 1.31 | 129 | 15.6 | 12.7 | 0.339 | 0.518 | 58.9 | 3.9 | 43.4 | 3.6 |
| 5iyd | E | 3.9 | 8138 | 0.01 | 1.31 | 210 | 11.0 | 6.4 | 0.715 | 0.808 | 79.5 | 36.5 | 68.6 | 5.6 |
| 5iyd | H | 3.9 | 8138 | 0.01 | 1.31 | 150 | 16.6 | 15.7 | 0.300 | 0.429 | 72 | 6.5 | 33.3 | 12 |
| 5juy | I | 4.1 | 8178 | 0.03 | 1.368 | 104 | 10.1 | 10.1 | 0.342 | 0.353 | 59.6 | 3.2 | 22.1 | 17.4 |
| 5juy | L | 4.1 | 8178 | 0.03 | 1.368 | 104 | 10.0 | 9.4 | 0.385 | 0.385 | 54.8 | 7 | 24 | 12 |
| 5kel | A | 4.3 | 8240 | 0.025 | 1.31 | 235 | 19.5 | 18.3 | 0.439 | 0.520 | 73.2 | 14 | 51.1 | 9.2 |
| 5kel | C | 4.3 | 8240 | 0.025 | 1.31 | 121 | 15.8 | 15.4 | 0.272 | 0.302 | 57.9 | 7.1 | 32.2 | 2.6 |
| 5kel | D | 4.3 | 8240 | 0.025 | 1.31 | 107 | 15.6 | 12.7 | 0.288 | 0.363 | 67.3 | 12.5 | 49.5 | 11.3 |
| 5kel | E | 4.3 | 8240 | 0.025 | 1.31 | 235 | 22.6 | 17.4 | 0.321 | 0.480 | 73.6 | 3.5 | 61.7 | 4.8 |
| 5kel | F | 4.3 | 8240 | 0.025 | 1.31 | 235 | 20.2 | 14.8 | 0.259 | 0.661 | 75.7 | 6.7 | 71.5 | 6 |
| 5kel | H | 4.3 | 8240 | 0.025 | 1.31 | 120 | 15.1 | 14.7 | 0.279 | 0.284 | 64.2 | 2.6 | 51.7 | 6.5 |
| 5kel | J | 4.3 | 8240 | 0.025 | 1.31 | 121 | 15.5 | 14.7 | 0.264 | 0.417 | 70.2 | 11.8 | 33.1 | 0 |

| | | | | | | | | | | | | | | |
|---|---|---|---|---|---|---|---|---|---|---|---|---|---|---|
| 5kel | L | 4.3 | 8240 | 0.025 | 1.31 | 107 | 14.1 | 13.0 | 0.288 | 0.513 | 76.6 | 12.2 | 44.9 | 6.2 |
| 5kel | M | 4.3 | 8240 | 0.025 | 1.31 | 121 | 17.0 | 14.9 | 0.323 | 0.345 | 66.1 | 11.2 | 41.3 | 8 |
| 5kel | N | 4.3 | 8240 | 0.025 | 1.31 | 107 | 13.8 | 13.3 | 0.262 | 0.364 | 70.1 | 10.7 | 43 | 6.5 |
| 5kel | O | 4.3 | 8240 | 0.025 | 1.31 | 107 | 13.9 | 13.7 | 0.300 | 0.337 | 67.3 | 9.7 | 43.9 | 8.5 |
| 5kel | P | 4.3 | 8240 | 0.025 | 1.31 | 120 | 15.3 | 14.2 | 0.275 | 0.338 | 69.2 | 7.2 | 34.2 | 4.9 |
| 5kel | Q | 4.3 | 8240 | 0.025 | 1.31 | 120 | 14.2 | 14.2 | 0.258 | 0.344 | 60.8 | 2.7 | 46.7 | 8.9 |
| 5kel | T | 4.3 | 8240 | 0.025 | 1.31 | 107 | 13.7 | 13.6 | 0.266 | 0.389 | 67.3 | 9.7 | 62.6 | 4.5 |
| 5kel | U | 4.3 | 8240 | 0.025 | 1.31 | 107 | 14.7 | 13.6 | 0.247 | 0.313 | 71 | 11.8 | 48.6 | 3.8 |
| 5ken | C | 4.3 | 8242 | 0.011 | 1.31 | 118 | 16.1 | 14.9 | 0.243 | 0.276 | 54.2 | 10.9 | 29.7 | 5.7 |
| 5ken | D | 4.3 | 8242 | 0.011 | 1.31 | 107 | 17.1 | 15.2 | 0.295 | 0.300 | 69.2 | 9.5 | 43.9 | 4.3 |
| 5ken | E | 4.3 | 8242 | 0.011 | 1.31 | 235 | 21.5 | 19.2 | 0.240 | 0.279 | 58.3 | 5.8 | 34.5 | 3.7 |
| 5ken | G | 4.3 | 8242 | 0.011 | 1.31 | 118 | 15.8 | 14.7 | 0.289 | 0.333 | 59.3 | 5.7 | 33.1 | 7.7 |
| 5ken | H | 4.3 | 8242 | 0.011 | 1.31 | 107 | 13.7 | 13.7 | 0.292 | 0.292 | 69.2 | 5.4 | 41.1 | 2.3 |
| 5ken | I | 4.3 | 8242 | 0.011 | 1.31 | 107 | 14.4 | 14.4 | 0.220 | 0.264 | 51.4 | 3.6 | 34.6 | 2.7 |
| 5ken | J | 4.3 | 8242 | 0.011 | 1.31 | 121 | 17.0 | 16.8 | 0.255 | 0.306 | 57.9 | 5.7 | 35.5 | 7 |
| 5ken | K | 4.3 | 8242 | 0.011 | 1.31 | 235 | 23.6 | 17.7 | 0.248 | 0.342 | 63.8 | 10 | 25.5 | 3.3 |
| 5ken | N | 4.3 | 8242 | 0.011 | 1.31 | 118 | 16.4 | 14.3 | 0.279 | 0.289 | 53.4 | 7.9 | 29.7 | 2.9 |
| 5ken | O | 4.3 | 8242 | 0.011 | 1.31 | 107 | 14.8 | 14.0 | 0.268 | 0.289 | 56.1 | 3.3 | 40.2 | 7 |
| 5ken | P | 4.3 | 8242 | 0.011 | 1.31 | 107 | 14.1 | 14.1 | 0.313 | 0.313 | 57.9 | 3.2 | 35.5 | 7.9 |
| 5ken | Q | 4.3 | 8242 | 0.011 | 1.31 | 121 | 15.9 | 15.5 | 0.285 | 0.358 | 56.2 | 5.9 | 34.7 | 11.9 |
| 5kgf | M | 4.5 | 8246 | 0.04 | 1.45 | 76 | 12.5 | 10.5 | 0.303 | 0.303 | 61.8 | 19.1 | 56.6 | 7 |
| 5kgf | O | 4.5 | 8246 | 0.04 | 1.45 | 76 | 10.5 | 8.5 | 0.282 | 0.420 | 73.7 | 8.9 | 51.3 | 7.7 |
| 5lc5 | E | 4.4 | 4032 | 0.165 | 1.33 | 186 | 9.9 | 9.9 | 0.643 | 0.643 | 83.3 | 38.7 | 72 | 6 |
| 5lc5 | S | 4.4 | 4032 | 0.165 | 1.33 | 80 | 13.5 | 12.3 | 0.242 | 0.282 | 66.2 | 5.7 | 43.8 | 0 |
| 5lc5 | T | 4.4 | 4032 | 0.165 | 1.33 | 75 | 11.5 | 5.8 | 0.312 | 0.587 | 70.7 | 7.5 | 78.7 | 5.1 |
| 5lc5 | U | 4.4 | 4032 | 0.165 | 1.33 | 85 | 3.9 | 3.9 | 0.731 | 0.731 | 84.7 | 29.2 | 71.8 | 4.9 |
| 5lc5 | V | 4.4 | 4032 | 0.165 | 1.33 | 106 | 8.2 | 7.1 | 0.678 | 0.678 | 75.5 | 45 | 87.7 | 8.6 |
| 5ldw | E | 4.3 | 4040 | 0.165 | 1.33 | 186 | 7.9 | 4.4 | 0.800 | 0.800 | 82.8 | 29.9 | 68.8 | 3.9 |
| 5ldw | S | 4.3 | 4040 | 0.165 | 1.33 | 80 | 13.3 | 7.4 | 0.339 | 0.616 | 78.8 | 22.2 | 52.5 | 9.5 |
| 5ldw | T | 4.3 | 4040 | 0.165 | 1.33 | 75 | 9.1 | 9.1 | 0.358 | 0.358 | 78.7 | 6.8 | 76 | 0 |
| 5ldw | U | 4.3 | 4040 | 0.165 | 1.33 | 85 | 3.7 | 3.7 | 0.748 | 0.748 | 87.1 | 40.5 | 81.2 | 24.6 |
| 5ldw | V | 4.3 | 4040 | 0.165 | 1.33 | 106 | 7.1 | 3.3 | 0.729 | 0.771 | 83 | 51.1 | 84.9 | 10 |
| 5ldw | W | 4.3 | 4040 | 0.165 | 1.33 | 111 | 5.0 | 5.0 | 0.789 | 0.794 | 85.6 | 22.1 | 83.8 | 34.4 |
| 5lj3 | d | 3.8 | 4055 | 0.027 | 1.43 | 82 | 12.7 | 8.6 | 0.345 | 0.644 | 78 | 20.3 | 51.2 | 9.5 |
| 5lj3 | L | 3.8 | 4055 | 0.027 | 1.43 | 155 | 4.4 | 4.4 | 0.826 | 0.826 | 87.1 | 42.2 | 78.1 | 11.6 |
| 5lj3 | M | 3.8 | 4055 | 0.027 | 1.43 | 252 | 8.3 | 8.3 | 0.782 | 0.796 | 79 | 32.7 | 72.2 | 15.9 |
| 5lj3 | N | 3.8 | 4055 | 0.027 | 1.43 | 209 | 22.5 | 20.1 | 0.436 | 0.601 | 59.3 | 24.2 | 70.8 | 7.4 |

| | | | | | | | | | | | | | | |
|---|---|---|---|---|---|---|---|---|---|---|---|---|---|---|
| 5ll6 | P | 3.9 | 4071 | 0.038 | 1.084 | 206 | 8.9 | 5.1 | 0.785 | 0.786 | 79.1 | 16.6 | 68.9 | 6.3 |
| 5lmn | C | 3.6 | 4073 | 0.09 | 1.34 | 206 | 7.9 | 6.2 | 0.769 | 0.824 | 87.9 | 61.9 | 80.1 | 3.6 |
| 5lmn | D | 3.6 | 4073 | 0.09 | 1.34 | 208 | 5.3 | 4.2 | 0.804 | 0.836 | 82.7 | 49.4 | 56.2 | 8.5 |
| 5lmn | E | 3.6 | 4073 | 0.09 | 1.34 | 150 | 4.2 | 4.2 | 0.813 | 0.813 | 88.7 | 82.7 | 86 | 18.6 |
| 5lmn | F | 3.6 | 4073 | 0.09 | 1.34 | 101 | 11.3 | 11.3 | 0.399 | 0.524 | 78.2 | 11.4 | 67.3 | 10.3 |
| 5lmn | G | 3.6 | 4073 | 0.09 | 1.34 | 155 | 2.3 | 2.3 | 0.903 | 0.903 | 94.2 | 71.2 | 83.9 | 6.9 |
| 5lmn | H | 3.6 | 4073 | 0.09 | 1.34 | 138 | 10.0 | 8.7 | 0.731 | 0.747 | 79.7 | 41.8 | 79.7 | 32.7 |
| 5lmo | C | 4.3 | 4074 | 0.05 | 1.34 | 206 | 20.2 | 15.4 | 0.285 | 0.428 | 62.6 | 6.2 | 44.7 | 6.5 |
| 5lmo | D | 4.3 | 4074 | 0.05 | 1.34 | 208 | 7.3 | 7.3 | 0.715 | 0.715 | 71.6 | 26.2 | 45.7 | 7.4 |
| 5lmo | F | 4.3 | 4074 | 0.05 | 1.34 | 101 | 14.4 | 13.0 | 0.399 | 0.471 | 68.3 | 11.6 | 40.6 | 2.4 |
| 5lmo | X | 4.3 | 4074 | 0.05 | 1.34 | 168 | 15.3 | 12.9 | 0.286 | 0.431 | 64.3 | 11.1 | 31.5 | 5.7 |
| 5lmq | C | 4.2 | 4076 | 0.075 | 1.34 | 206 | 12.6 | 12.6 | 0.542 | 0.544 | 73.8 | 22.4 | 59.7 | 8.9 |
| 5lmq | F | 4.2 | 4076 | 0.075 | 1.34 | 101 | 13.1 | 9.7 | 0.418 | 0.479 | 72.3 | 15.1 | 80.2 | 23.5 |
| 5lmq | X | 4.2 | 4076 | 0.075 | 1.34 | 168 | 12.9 | 12.0 | 0.407 | 0.479 | 75 | 16.7 | 53.6 | 7.8 |
| 5lmr | C | 4.5 | 4077 | 0.075 | 1.34 | 206 | 14.2 | 12.4 | 0.580 | 0.583 | 71.8 | 18.2 | 32.5 | 10.4 |
| 5lmr | F | 4.5 | 4077 | 0.075 | 1.34 | 101 | 13.9 | 12.7 | 0.433 | 0.435 | 72.3 | 13.7 | 76.2 | 10.4 |
| 5lmr | X | 4.5 | 4077 | 0.075 | 1.34 | 168 | 14.0 | 12.6 | 0.414 | 0.429 | 68.5 | 8.7 | 54.8 | 6.5 |
| 5lmt | C | 4.2 | 4079 | 0.08 | 1.34 | 206 | 17.7 | 11.5 | 0.430 | 0.565 | 76.7 | 10.1 | 26.2 | 1.9 |
| 5lmt | F | 4.2 | 4079 | 0.08 | 1.34 | 101 | 16.2 | 13.6 | 0.234 | 0.454 | 76.2 | 3.9 | 55.4 | 3.6 |
| 5lmu | C | 4 | 4080 | 0.1 | 1.34 | 206 | 9.2 | 9.2 | 0.760 | 0.760 | 84 | 41 | 69.9 | 3.5 |
| 5lmu | F | 4 | 4080 | 0.1 | 1.34 | 101 | 15.6 | 11.1 | 0.337 | 0.479 | 81.2 | 11 | 86.1 | 4.6 |
| 5lmu | G | 4 | 4080 | 0.1 | 1.34 | 155 | 5.0 | 5.0 | 0.772 | 0.781 | 81.3 | 65.9 | 40.6 | 1.6 |
| 5lmx | E | 4.9 | 4088 | 0.14 | 1.77 | 215 | 21.5 | 13.1 | 0.452 | 0.576 | 65.1 | 8.6 | 33 | 5.6 |
| 5lmx | F | 4.9 | 4088 | 0.14 | 1.77 | 100 | 20.2 | 16.1 | 0.433 | 0.447 | 65 | 4.6 | 60 | 5 |
| 5lmx | H | 4.9 | 4088 | 0.14 | 1.77 | 134 | 17.6 | 13.7 | 0.267 | 0.292 | 66.4 | 3.4 | 41.8 | 3.6 |
| 5m0q | B | 4.9 | 4138 | 0.03 | 1.35 | 256 | 17.3 | 17.3 | 0.482 | 0.482 | 58.2 | 7.4 | 33.2 | 7.1 |
| 5m0q | C | 4.9 | 4138 | 0.03 | 1.35 | 261 | 23.4 | 20.7 | 0.310 | 0.380 | 58.6 | 2.6 | 15.7 | 12.2 |
| 5m0q | E | 4.9 | 4138 | 0.03 | 1.35 | 257 | 16.3 | 12.2 | 0.392 | 0.534 | 64.6 | 9.6 | 36.6 | 4.3 |
| 5m0q | F | 4.9 | 4138 | 0.03 | 1.35 | 266 | 23.9 | 22.0 | 0.323 | 0.400 | 60.5 | 7.5 | 37.6 | 5 |
| 5m0q | G | 4.9 | 4138 | 0.03 | 1.35 | 205 | 22.6 | 18.9 | 0.252 | 0.266 | 61.5 | 7.9 | 29.8 | 3.3 |
| 5m0q | H | 4.9 | 4138 | 0.03 | 1.35 | 217 | 18.8 | 15.0 | 0.319 | 0.330 | 55.3 | 11.7 | 27.2 | 6.8 |
| 5m0q | J | 4.9 | 4138 | 0.03 | 1.35 | 256 | 25.0 | 15.1 | 0.255 | 0.519 | 62.5 | 6.2 | 39.1 | 3 |
| 5m0q | K | 4.9 | 4138 | 0.03 | 1.35 | 261 | 22.5 | 16.3 | 0.482 | 0.482 | 61.7 | 4.3 | 24.5 | 7.8 |
| 5m0q | M | 4.9 | 4138 | 0.03 | 1.35 | 256 | 13.0 | 12.9 | 0.534 | 0.563 | 64.5 | 13.3 | 41.4 | 2.8 |
| 5m0q | N | 4.9 | 4138 | 0.03 | 1.35 | 264 | 22.9 | 14.3 | 0.423 | 0.516 | 59.5 | 8.3 | 20.8 | 5.5 |
| 5m0q | O | 4.9 | 4138 | 0.03 | 1.35 | 205 | 21.1 | 16.5 | 0.268 | 0.431 | 66.8 | 5.1 | 24.4 | 6 |
| 5m0q | P | 4.9 | 4138 | 0.03 | 1.35 | 217 | 16.5 | 13.4 | 0.385 | 0.420 | 60.4 | 3.8 | 40.6 | 2.3 |

| | | | | | | | | | | | | | | |
|---|---|---|---|---|---|---|---|---|---|---|---|---|---|---|
| 5m32 | A | 3.8 | 4146 | 0.05 | 1.27 | 230 | 2.0 | 2.0 | 0.939 | 0.939 | 97.4 | 84.4 | 83.9 | 45.1 |
| 5m32 | B | 3.8 | 4146 | 0.05 | 1.27 | 234 | 2.6 | 2.6 | 0.892 | 0.892 | 91.9 | 67.9 | 89.7 | 36.2 |
| 5m32 | C | 3.8 | 4146 | 0.05 | 1.27 | 234 | 1.9 | 1.9 | 0.932 | 0.932 | 95.3 | 82.1 | 79.5 | 45.2 |
| 5m32 | D | 3.8 | 4146 | 0.05 | 1.27 | 233 | 3.3 | 3.3 | 0.896 | 0.896 | 93.1 | 66.8 | 81.1 | 27.5 |
| 5m32 | E | 3.8 | 4146 | 0.05 | 1.27 | 234 | 1.9 | 1.9 | 0.941 | 0.941 | 97 | 83.3 | 84.6 | 50 |
| 5m32 | G | 3.8 | 4146 | 0.05 | 1.27 | 233 | 9.0 | 2.9 | 0.775 | 0.918 | 94.8 | 69.7 | 91.4 | 34.3 |
| 5m32 | i | 3.8 | 4146 | 0.05 | 1.27 | 230 | 13.9 | 11.1 | 0.536 | 0.574 | 75.2 | 27.2 | 75.7 | 5.2 |
| 5m32 | J | 3.8 | 4146 | 0.05 | 1.27 | 196 | 2.0 | 2.0 | 0.938 | 0.938 | 96.9 | 84.2 | 88.3 | 15.6 |
| 5m32 | O | 3.8 | 4146 | 0.05 | 1.27 | 230 | 2.9 | 2.9 | 0.887 | 0.887 | 92.2 | 66.5 | 76.1 | 31.4 |
| 5m32 | P | 3.8 | 4146 | 0.05 | 1.27 | 244 | 17.8 | 17.3 | 0.387 | 0.558 | 80.3 | 8.7 | 71.3 | 33.3 |
| 5m32 | q | 3.8 | 4146 | 0.05 | 1.27 | 259 | 25.7 | 25.7 | 0.280 | 0.408 | 59.5 | 2.6 | 56 | 6.2 |
| 5m32 | Q | 3.8 | 4146 | 0.05 | 1.27 | 235 | 3.6 | 3.6 | 0.876 | 0.876 | 89.4 | 60.5 | 80.4 | 29.1 |
| 5m32 | R | 3.8 | 4146 | 0.05 | 1.27 | 233 | 5.9 | 5.4 | 0.854 | 0.858 | 85.4 | 44.7 | 74.2 | 7.5 |
| 5m32 | S | 3.8 | 4146 | 0.05 | 1.27 | 238 | 3.1 | 3.1 | 0.887 | 0.887 | 90.8 | 68.5 | 83.2 | 37.9 |
| 5m32 | U | 3.8 | 4146 | 0.05 | 1.27 | 241 | 3.4 | 3.4 | 0.888 | 0.888 | 90.5 | 64.7 | 79.3 | 4.7 |
| 5m32 | X | 3.8 | 4146 | 0.05 | 1.27 | 196 | 2.4 | 2.4 | 0.922 | 0.922 | 94.4 | 77.8 | 86.2 | 5.3 |
| 5m3f | E | 3.8 | 4147 | 0.045 | 1.05 | 212 | 8.0 | 8.0 | 0.739 | 0.744 | 82.1 | 46 | 68.9 | 6.8 |
| 5m3f | H | 3.8 | 4147 | 0.045 | 1.05 | 131 | 11.2 | 11.2 | 0.734 | 0.734 | 78.6 | 38.8 | 71.8 | 3.2 |
| 5m3m | H | 4 | 4148 | 0.04 | 1.05 | 131 | 18.0 | 15.8 | 0.272 | 0.405 | 73.3 | 9.4 | 71.8 | 4.3 |
| 5m5w | E | 3.8 | 3446 | 0.03 | 1.35 | 214 | 3.4 | 3.4 | 0.860 | 0.860 | 88.3 | 44.4 | 77.6 | 8.4 |
| 5m5w | H | 3.8 | 3446 | 0.03 | 1.35 | 134 | 16.6 | 14.1 | 0.466 | 0.605 | 76.1 | 16.7 | 62.7 | 8.3 |
| 5m5x | E | 4 | 3447 | 0.03 | 1.35 | 214 | 19.2 | 12.1 | 0.381 | 0.581 | 63.6 | 8.1 | 60.3 | 7.8 |
| 5m5x | H | 4 | 3447 | 0.03 | 1.35 | 134 | 14.9 | 13.6 | 0.272 | 0.431 | 70.1 | 3.2 | 25.4 | 8.8 |
| 5m5y | E | 4 | 3448 | 0.03 | 1.35 | 214 | 7.6 | 7.6 | 0.725 | 0.735 | 74.8 | 12.5 | 69.2 | 9.5 |
| 5m5y | H | 4 | 3448 | 0.03 | 1.35 | 134 | 15.2 | 14.0 | 0.240 | 0.267 | 62.7 | 6 | 35.8 | 8.3 |
| 5m64 | E | 4.6 | 3449 | 0.045 | 1.35 | 214 | 15.8 | 14.2 | 0.439 | 0.479 | 64.5 | 10.1 | 45.3 | 3.1 |
| 5m64 | H | 4.6 | 3449 | 0.045 | 1.35 | 134 | 16.9 | 15.2 | 0.262 | 0.309 | 63.4 | 11.8 | 34.3 | 2.2 |
| 5mmi | D | 3.2 | 3531 | 0.08 | 1.39 | 221 | 10.1 | 7.9 | 0.736 | 0.840 | 91 | 36.3 | 77.4 | 20.5 |
| 5mmi | K | 3.2 | 3531 | 0.08 | 1.39 | 203 | 2.4 | 2.4 | 0.911 | 0.911 | 92.1 | 80.7 | 79.8 | 30.9 |
| 5mmi | L | 3.2 | 3531 | 0.08 | 1.39 | 121 | 5.2 | 4.5 | 0.889 | 0.889 | 93.4 | 54.9 | 84.3 | 5.9 |
| 5mmi | O | 3.2 | 3531 | 0.08 | 1.39 | 116 | 2.1 | 2.1 | 0.914 | 0.914 | 95.7 | 85.6 | 86.2 | 71 |
| 5mmi | P | 3.2 | 3531 | 0.08 | 1.39 | 122 | 3.9 | 3.9 | 0.808 | 0.808 | 86.1 | 28.6 | 78.7 | 8.3 |
| 5mmi | Q | 3.2 | 3531 | 0.08 | 1.39 | 118 | 2.0 | 2.0 | 0.904 | 0.904 | 94.9 | 72.3 | 78 | 8.7 |
| 5mmi | S | 3.2 | 3531 | 0.08 | 1.39 | 170 | 5.6 | 5.2 | 0.869 | 0.869 | 90.6 | 66.2 | 34.7 | 5.1 |
| 5mmi | T | 3.2 | 3531 | 0.08 | 1.39 | 172 | 4.0 | 3.9 | 0.894 | 0.899 | 91.9 | 46.2 | 72.7 | 24.8 |
| 5mmi | Z | 3.2 | 3531 | 0.08 | 1.39 | 101 | 2.7 | 2.7 | 0.871 | 0.871 | 92.1 | 69.9 | 87.1 | 13.6 |
| 5mmj | c | 3.6 | 3532 | 0.06 | 1.39 | 216 | 15.2 | 14.0 | 0.511 | 0.546 | 83.8 | 22.1 | 70.8 | 7.8 |

| | | | | | | | | | | | | | | |
|---|---|---|---|---|---|---|---|---|---|---|---|---|---|---|
| 5mmj | d | 3.6 | 3532 | 0.06 | 1.39 | 199 | 3.2 | 3.2 | 0.865 | 0.865 | 90.5 | 45.6 | 77.4 | 40.3 |
| 5mmj | e | 3.6 | 3532 | 0.06 | 1.39 | 187 | 18.4 | 16.9 | 0.736 | 0.736 | 81.3 | 28.3 | 68.4 | 15.6 |
| 5mmj | f | 3.6 | 3532 | 0.06 | 1.39 | 113 | 17.9 | 11.4 | 0.478 | 0.643 | 78.8 | 5.6 | 69.9 | 5.1 |
| 5mps | a | 3.9 | 3539 | 0.036 | 1.43 | 137 | 3.7 | 3.7 | 0.762 | 0.762 | 81.8 | 42.9 | 87.6 | 30 |
| 5mps | d | 3.9 | 3539 | 0.036 | 1.43 | 82 | 12.6 | 10.8 | 0.248 | 0.544 | 81.7 | 7.5 | 40.2 | 6.1 |
| 5mps | L | 3.9 | 3539 | 0.036 | 1.43 | 155 | 7.1 | 4.9 | 0.802 | 0.822 | 83.9 | 55.4 | 74.2 | 1.7 |
| 5mps | M | 3.9 | 3539 | 0.036 | 1.43 | 252 | 10.5 | 8.2 | 0.737 | 0.737 | 81 | 27 | 48.4 | 4.1 |
| 5mps | N | 3.9 | 3539 | 0.036 | 1.43 | 227 | 21.9 | 21.0 | 0.581 | 0.583 | 58.1 | 6.8 | 66.5 | 6 |
| 5mq0 | a | 4.2 | 3541 | 0.03 | 1.43 | 137 | 4.2 | 4.2 | 0.727 | 0.727 | 75.2 | 37.9 | 43.8 | 10 |
| 5mq0 | L | 4.2 | 3541 | 0.03 | 1.43 | 155 | 19.2 | 17.2 | 0.309 | 0.572 | 73.5 | 7 | 33.5 | 3.8 |
| 5mq0 | M | 4.2 | 3541 | 0.03 | 1.43 | 252 | 20.9 | 13.7 | 0.384 | 0.550 | 74.6 | 7.4 | 52.8 | 3 |
| 5mq0 | N | 4.2 | 3541 | 0.03 | 1.43 | 227 | 21.7 | 20.5 | 0.526 | 0.528 | 53.7 | 8.2 | 40.5 | 4.3 |
| 5n61 | E | 3.4 | 3593 | 0.065 | 1.13 | 212 | 4.5 | 4.5 | 0.823 | 0.858 | 94.3 | 49.5 | 72.2 | 19 |
| 5n61 | F | 3.4 | 3593 | 0.065 | 1.13 | 100 | 1.9 | 1.9 | 0.926 | 0.926 | 96 | 87.5 | 91 | 37.4 |
| 5n61 | G | 3.4 | 3593 | 0.065 | 1.13 | 193 | 9.1 | 8.6 | 0.700 | 0.705 | 80.3 | 46.5 | 38.9 | 4 |
| 5n61 | H | 3.4 | 3593 | 0.065 | 1.13 | 131 | 15.8 | 13.4 | 0.525 | 0.546 | 88.5 | 33.6 | 87.8 | 11.3 |
| 5n61 | J | 3.4 | 3593 | 0.065 | 1.13 | 69 | 1.9 | 1.9 | 0.856 | 0.856 | 94.2 | 72.3 | 82.6 | 33.3 |
| 5n61 | K | 3.4 | 3593 | 0.065 | 1.13 | 101 | 1.2 | 1.2 | 0.927 | 0.927 | 97 | 95.9 | 91.1 | 35.9 |
| 5nj3 | C | 3.8 | 3654 | 0.038 | 1.039 | 208 | 16.0 | 15.7 | 0.504 | 0.572 | 70.2 | 11 | 33.7 | 20 |
| 5nj3 | D | 3.8 | 3654 | 0.038 | 1.039 | 210 | 14.2 | 13.9 | 0.638 | 0.682 | 81.4 | 45 | 41.4 | 9.2 |
| 5nj3 | E | 3.8 | 3654 | 0.038 | 1.039 | 208 | 15.7 | 15.4 | 0.386 | 0.479 | 73.6 | 15.7 | 29.3 | 9.8 |
| 5nj3 | F | 3.8 | 3654 | 0.038 | 1.039 | 210 | 19.6 | 16.8 | 0.609 | 0.609 | 76.7 | 16.1 | 49.5 | 8.7 |
| 5nsr | A | 3.8 | 3695 | 0.008 | 1.06 | 233 | 26.4 | 24.1 | 0.437 | 0.437 | 67.4 | 14 | 45.9 | 4.7 |
| 5nsr | B | 3.8 | 3695 | 0.008 | 1.06 | 223 | 25.0 | 24.1 | 0.437 | 0.440 | 69.5 | 18.1 | 45.7 | 12.7 |
| 5o31 | E | 4.1 | 3731 | 0.14 | 1.38 | 186 | 5.1 | 5.1 | 0.805 | 0.805 | 80.6 | 18.7 | 53.2 | 5.1 |
| 5o31 | q | 4.1 | 3731 | 0.14 | 1.38 | 138 | 12.5 | 4.6 | 0.740 | 0.803 | 74.6 | 34 | 74.6 | 6.8 |
| 5o31 | S | 4.1 | 3731 | 0.14 | 1.38 | 80 | 12.2 | 9.7 | 0.303 | 0.578 | 72.5 | 6.9 | 45 | 2.8 |
| 5o31 | T | 4.1 | 3731 | 0.14 | 1.38 | 75 | 7.4 | 7.4 | 0.526 | 0.526 | 66.7 | 22 | 74.7 | 5.4 |
| 5o31 | U | 4.1 | 3731 | 0.14 | 1.38 | 85 | 2.7 | 2.7 | 0.774 | 0.774 | 84.7 | 58.3 | 83.5 | 70.4 |
| 5o31 | V | 4.1 | 3731 | 0.14 | 1.38 | 106 | 4.9 | 4.9 | 0.751 | 0.755 | 82.1 | 20.7 | 78.3 | 9.6 |
| 5o31 | W | 4.1 | 3731 | 0.14 | 1.38 | 111 | 3.7 | 3.7 | 0.817 | 0.817 | 86.5 | 45.8 | 88.3 | 32.7 |
| 5o5j | C | 3.5 | 3748 | 0.1 | 1.39 | 208 | 3.0 | 3.0 | 0.898 | 0.898 | 91.3 | 64.2 | 74.5 | 6.5 |
| 5o5j | D | 3.5 | 3748 | 0.1 | 1.39 | 200 | 4.3 | 4.3 | 0.853 | 0.853 | 87.5 | 40.6 | 65.5 | 9.2 |
| 5o5j | F | 3.5 | 3748 | 0.1 | 1.39 | 96 | 3.1 | 3.1 | 0.870 | 0.870 | 91.7 | 54.5 | 84.4 | 8.6 |
| 5o5j | G | 3.5 | 3748 | 0.1 | 1.39 | 155 | 5.2 | 5.1 | 0.834 | 0.841 | 87.7 | 43.4 | 80.6 | 20.8 |
| 5o5j | H | 3.5 | 3748 | 0.1 | 1.39 | 131 | 2.8 | 2.8 | 0.881 | 0.881 | 92.4 | 77.7 | 86.3 | 46 |
| 5o5j | O | 3.5 | 3748 | 0.1 | 1.39 | 88 | 3.6 | 3.6 | 0.844 | 0.844 | 88.6 | 60.3 | 75 | 78.8 |

| | | | | | | | | | | | | | | |
|---|---|---|---|---|---|---|---|---|---|---|---|---|---|---|
| 5o5j | P | 3.5 | 3748 | 0.1 | 1.39 | 113 | 1.8 | 1.8 | 0.889 | 0.889 | 94.7 | 77.6 | 58.4 | 18.2 |
| 5o5j | R | 3.5 | 3748 | 0.1 | 1.39 | 65 | 2.7 | 2.7 | 0.843 | 0.843 | 93.8 | 47.5 | 78.5 | 52.9 |
| 5o60 | D | 3.2 | 3750 | 0.1 | 1.39 | 214 | 4.1 | 4.1 | 0.893 | 0.893 | 92.1 | 66.5 | 83.6 | 12.8 |
| 5o60 | E | 3.2 | 3750 | 0.1 | 1.39 | 209 | 1.7 | 1.7 | 0.938 | 0.938 | 96.7 | 84.2 | 76.1 | 46.5 |
| 5o60 | F | 3.2 | 3750 | 0.1 | 1.39 | 182 | 7.2 | 7.2 | 0.772 | 0.772 | 78.6 | 34.3 | 69.8 | 4.7 |
| 5o60 | G | 3.2 | 3750 | 0.1 | 1.39 | 176 | 9.0 | 8.5 | 0.756 | 0.756 | 77.3 | 36 | 68.8 | 10.7 |
| 5o60 | K | 3.2 | 3750 | 0.1 | 1.39 | 146 | 2.4 | 2.4 | 0.911 | 0.911 | 93.8 | 64.2 | 89.7 | 35.1 |
| 5o60 | N | 3.2 | 3750 | 0.1 | 1.39 | 136 | 1.4 | 1.4 | 0.944 | 0.944 | 97.8 | 94.7 | 95.6 | 44.6 |
| 5o60 | O | 3.2 | 3750 | 0.1 | 1.39 | 118 | 2.1 | 2.1 | 0.914 | 0.914 | 95.8 | 81.4 | 88.1 | 50 |
| 5o60 | P | 3.2 | 3750 | 0.1 | 1.39 | 126 | 3.9 | 3.9 | 0.815 | 0.845 | 84.1 | 55.7 | 80.2 | 25.7 |
| 5o60 | Q | 3.2 | 3750 | 0.1 | 1.39 | 113 | 2.6 | 2.6 | 0.873 | 0.873 | 92 | 65.4 | 87.6 | 5.1 |
| 5o60 | S | 3.2 | 3750 | 0.1 | 1.39 | 100 | 2.2 | 2.2 | 0.904 | 0.904 | 95 | 64.2 | 76 | 7.9 |
| 5o60 | T | 3.2 | 3750 | 0.1 | 1.39 | 114 | 3.1 | 3.1 | 0.899 | 0.899 | 93.9 | 81.3 | 80.7 | 33.7 |
| 5o60 | U | 3.2 | 3750 | 0.1 | 1.39 | 97 | 2.7 | 2.7 | 0.860 | 0.860 | 90.7 | 77.3 | 75.3 | 2.7 |
| 5o60 | W | 3.2 | 3750 | 0.1 | 1.39 | 192 | 6.9 | 6.9 | 0.765 | 0.765 | 88.5 | 49.4 | 55.2 | 11.3 |
| 5o60 | Z | 3.2 | 3750 | 0.1 | 1.39 | 64 | 1.4 | 1.4 | 0.831 | 0.831 | 98.4 | 100 | 85.9 | 9.1 |
| 5o9z | I | 4.5 | 3766 | 0.04 | 1.16 | 176 | 17.0 | 16.2 | 0.522 | 0.542 | 62.5 | 6.4 | 42.6 | 6.7 |
| 5o9z | J | 4.5 | 3766 | 0.04 | 1.16 | 135 | 15.1 | 11.9 | 0.336 | 0.465 | 65.9 | 4.5 | 59.3 | 5 |
| 5o9z | O | 4.5 | 3766 | 0.04 | 1.16 | 126 | 14.7 | 12.0 | 0.393 | 0.409 | 69.8 | 9.1 | 49.2 | 3.2 |
| 5oa1 | E | 4.4 | 3727 | 0.01 | 1.35 | 215 | 19.4 | 17.2 | 0.365 | 0.522 | 61.9 | 6 | 47 | 4 |
| 5oa1 | H | 4.4 | 3727 | 0.01 | 1.35 | 134 | 15.3 | 14.4 | 0.298 | 0.321 | 59 | 3.8 | 32.1 | 11.6 |
| 5of4 | E | 4.4 | 3802 | 0.03 | 1.32 | 184 | 14.8 | 14.2 | 0.361 | 0.435 | 67.4 | 8.9 | 70.7 | 6.2 |
| 5of4 | F | 4.4 | 3802 | 0.03 | 1.32 | 205 | 15.8 | 15.8 | 0.463 | 0.463 | 70.7 | 11.7 | 63.4 | 5.4 |
| 5oik | C | 3.7 | 3817 | 0.022 | 1.35 | 259 | 3.0 | 3.0 | 0.927 | 0.927 | 93.8 | 66.3 | 84.6 | 11 |
| 5oik | E | 3.7 | 3817 | 0.022 | 1.35 | 209 | 6.5 | 4.1 | 0.741 | 0.861 | 88.5 | 45.4 | 80.9 | 4.7 |
| 5oik | F | 3.7 | 3817 | 0.022 | 1.35 | 82 | 3.5 | 3.5 | 0.809 | 0.815 | 89 | 61.6 | 74.4 | 47.5 |
| 5oik | G | 3.7 | 3817 | 0.022 | 1.35 | 171 | 20.6 | 13.6 | 0.263 | 0.324 | 62 | 4.7 | 20.5 | 5.7 |
| 5oik | H | 3.7 | 3817 | 0.022 | 1.35 | 148 | 3.6 | 3.6 | 0.882 | 0.882 | 92.6 | 52.6 | 70.3 | 9.6 |
| 5oik | I | 3.7 | 3817 | 0.022 | 1.35 | 117 | 11.8 | 7.8 | 0.613 | 0.701 | 76.9 | 24.4 | 39.3 | 4.3 |
| 5oik | J | 3.7 | 3817 | 0.022 | 1.35 | 67 | 2.0 | 2.0 | 0.793 | 0.793 | 92.5 | 85.5 | 70.1 | 17 |
| 5oik | K | 3.7 | 3817 | 0.022 | 1.35 | 115 | 2.6 | 2.6 | 0.873 | 0.873 | 90.4 | 73.1 | 81.7 | 60.6 |
| 5ool | 7 | 3.1 | 3842 | 0.0853 | 1.06 | 287 | 22.2 | 19.2 | 0.521 | 0.627 | 74.2 | 6.6 | 55.7 | 7.5 |
| 5ool | c | 3.1 | 3842 | 0.0853 | 1.06 | 275 | 10.1 | 8.4 | 0.795 | 0.795 | 83.3 | 29.7 | 71.6 | 18.8 |
| 5oom | 7 | 3 | 3843 | 0.075 | 1.06 | 287 | 18.9 | 15.5 | 0.620 | 0.629 | 75.3 | 14.8 | 57.1 | 8.5 |
| 5oqj | 3 | 4.7 | 3846 | 0.066 | 1.37 | 138 | 9.2 | 8.4 | 0.505 | 0.514 | 66.7 | 10.9 | 32.6 | 4.4 |
| 5oqj | C | 4.7 | 3846 | 0.066 | 1.37 | 262 | 3.2 | 3.2 | 0.898 | 0.898 | 89.7 | 55.7 | 85.1 | 12.1 |
| 5oqj | D | 4.7 | 3846 | 0.066 | 1.37 | 157 | 18.1 | 16.2 | 0.303 | 0.331 | 55.4 | 10.3 | 52.2 | 4.9 |

| | | | | | | | | | | | | | | |
|---|---|---|---|---|---|---|---|---|---|---|---|---|---|---|
| 5oqj | E | 4.7 | 3846 | 0.066 | 1.37 | 213 | 17.3 | 14.0 | 0.529 | 0.652 | 75.1 | 35.6 | 66.7 | 2.8 |
| 5oqj | F | 4.7 | 3846 | 0.066 | 1.37 | 83 | 6.8 | 3.7 | 0.742 | 0.805 | 83.1 | 30.4 | 85.5 | 2.8 |
| 5oqj | G | 4.7 | 3846 | 0.066 | 1.37 | 171 | 19.9 | 16.8 | 0.228 | 0.311 | 63.2 | 6.5 | 31 | 3.8 |
| 5oqj | H | 4.7 | 3846 | 0.066 | 1.37 | 136 | 14.0 | 10.6 | 0.587 | 0.587 | 75 | 21.6 | 52.2 | 8.5 |
| 5oqj | I | 4.7 | 3846 | 0.066 | 1.37 | 116 | 18.4 | 13.7 | 0.283 | 0.559 | 76.7 | 9 | 67.2 | 9 |
| 5oqj | O | 4.7 | 3846 | 0.066 | 1.37 | 180 | 18.0 | 14.8 | 0.307 | 0.323 | 58.3 | 4.8 | 42.2 | 9.2 |
| 5sy1 | C | 3.9 | 8315 | 0.035 | 1.255 | 147 | 12.9 | 12.3 | 0.567 | 0.688 | 74.8 | 46.4 | 59.2 | 4.6 |
| 5sy1 | D | 3.9 | 8315 | 0.035 | 1.255 | 147 | 12.9 | 9.4 | 0.682 | 0.697 | 74.1 | 36.7 | 61.2 | 10 |
| 5t0g | b | 4.4 | 8334 | 0.0033 | 0.86 | 191 | 16.0 | 14.3 | 0.306 | 0.368 | 66.5 | 4.7 | 48.2 | 8.7 |
| 5t0g | G | 4.4 | 8334 | 0.0033 | 0.86 | 240 | 17.6 | 15.3 | 0.256 | 0.430 | 68.8 | 6.7 | 53.3 | 3.1 |
| 5t0g | H | 4.4 | 8334 | 0.0033 | 0.86 | 233 | 16.6 | 16.6 | 0.301 | 0.396 | 67.4 | 5.7 | 63.5 | 8.1 |
| 5t0g | I | 4.4 | 8334 | 0.0033 | 0.86 | 250 | 16.1 | 16.1 | 0.481 | 0.540 | 66.8 | 15 | 47.2 | 4.2 |
| 5t0g | J | 4.4 | 8334 | 0.0033 | 0.86 | 239 | 20.9 | 17.4 | 0.272 | 0.340 | 71.1 | 9.4 | 44.8 | 7.5 |
| 5t0g | L | 4.4 | 8334 | 0.0033 | 0.86 | 238 | 18.3 | 16.8 | 0.349 | 0.499 | 65.5 | 8.3 | 39.9 | 10.5 |
| 5tqq | H | 3.8 | 8435 | 0.019 | 1.3 | 111 | 14.8 | 12.2 | 0.439 | 0.499 | 78.4 | 13.8 | 78.4 | 9.2 |
| 5tqq | I | 3.8 | 8435 | 0.019 | 1.3 | 111 | 13.4 | 13.2 | 0.293 | 0.366 | 67.6 | 8 | 46.8 | 3.8 |
| 5tqq | L | 3.8 | 8435 | 0.019 | 1.3 | 107 | 11.5 | 10.2 | 0.442 | 0.634 | 74.8 | 3.8 | 75.7 | 4.9 |
| 5tqq | M | 3.8 | 8435 | 0.019 | 1.3 | 107 | 10.1 | 10.1 | 0.673 | 0.673 | 75.7 | 23.5 | 68.2 | 12.3 |
| 5tr1 | H | 4 | 8454 | 0.0176 | 1.3 | 111 | 15.6 | 13.1 | 0.285 | 0.309 | 68.5 | 6.6 | 34.2 | 5.3 |
| 5tr1 | I | 4 | 8454 | 0.0176 | 1.3 | 111 | 15.8 | 13.2 | 0.245 | 0.284 | 64 | 7 | 32.4 | 8.3 |
| 5tr1 | L | 4 | 8454 | 0.0176 | 1.3 | 107 | 13.1 | 13.1 | 0.321 | 0.422 | 66.4 | 18.3 | 50.5 | 3.7 |
| 5tr1 | M | 4 | 8454 | 0.0176 | 1.3 | 107 | 14.0 | 13.3 | 0.294 | 0.434 | 67.3 | 8.3 | 48.6 | 13.5 |
| 5u07 | D | 3.8 | 8477 | 0.07 | 1.23 | 260 | 20.5 | 14.8 | 0.291 | 0.755 | 77.7 | 7.9 | 84.6 | 5.5 |
| 5u07 | J | 3.8 | 8477 | 0.07 | 1.23 | 170 | 13.6 | 12.1 | 0.298 | 0.298 | 44.1 | 16 | 78.8 | 20.1 |
| 5u07 | N | 3.8 | 8477 | 0.07 | 1.23 | 241 | 16.1 | 16.1 | 0.715 | 0.715 | 81.3 | 44.4 | 58.5 | 8.5 |
| 5u0a | J | 3.3 | 8478 | 0.07 | 1.23 | 173 | 15.3 | 11.5 | 0.301 | 0.540 | 83.2 | 29.2 | 89 | 63.6 |
| 5u0a | L | 3.3 | 8478 | 0.07 | 1.23 | 165 | 11.6 | 11.6 | 0.683 | 0.683 | 87.3 | 62.5 | 83.6 | 25.4 |
| 5u0a | N | 3.3 | 8478 | 0.07 | 1.23 | 241 | 16.4 | 5.8 | 0.811 | 0.880 | 88.8 | 66.4 | 78.4 | 9 |
| 5u0p | 2 | 4.4 | 8479 | 0.016 | 1.31 | 78 | 12.0 | 9.4 | 0.339 | 0.339 | 67.9 | 13.2 | 51.3 | 5 |
| 5u0p | R | 4.4 | 8479 | 0.016 | 1.31 | 207 | 20.5 | 19.1 | 0.254 | 0.334 | 67.1 | 7.2 | 37.7 | 3.8 |
| 5u0p | T | 4.4 | 8479 | 0.016 | 1.31 | 179 | 18.3 | 15.9 | 0.292 | 0.326 | 62.6 | 6.2 | 52 | 7.5 |
| 5u1c | A | 3.9 | 8481 | 0.08 | 1.31 | 251 | 15.4 | 14.5 | 0.434 | 0.636 | 76.9 | 15.5 | 51.8 | 15.4 |
| 5u1c | B | 3.9 | 8481 | 0.08 | 1.31 | 172 | 17.3 | 13.3 | 0.463 | 0.493 | 70.9 | 7.4 | 66.9 | 5.2 |
| 5u1c | C | 3.9 | 8481 | 0.08 | 1.31 | 251 | 22.1 | 14.2 | 0.388 | 0.446 | 74.1 | 15.6 | 68.1 | 8.2 |
| 5u1c | D | 3.9 | 8481 | 0.08 | 1.31 | 172 | 12.7 | 11.9 | 0.391 | 0.419 | 70.9 | 11.5 | 41.9 | 12.5 |
| 5u8t | A | 4.9 | 8519 | 0.03 | 1.3 | 208 | 9.0 | 9.0 | 0.619 | 0.666 | 70.7 | 6.8 | 55.8 | 5.2 |
| 5ujz | G | 4.8 | 8561 | 2.3 | 1.64 | 236 | 19.0 | 18.1 | 0.242 | 0.302 | 57.2 | 6.7 | 29.7 | 7.1 |

| | | | | | | | | | | | | | | |
|---|---|---|---|---|---|---|---|---|---|---|---|---|---|---|
| 5ujz | H | 4.8 | 8561 | 2.3 | 1.64 | 236 | 17.6 | 17.6 | 0.267 | 0.299 | 61.9 | 8.2 | 25.4 | 8.3 |
| 5ujz | I | 4.8 | 8561 | 2.3 | 1.64 | 236 | 20.2 | 18.6 | 0.265 | 0.292 | 59.3 | 8.6 | 31.4 | 6.8 |
| 5uk0 | G | 4.8 | 8562 | 2.08 | 1.64 | 236 | 21.1 | 18.8 | 0.240 | 0.260 | 58.9 | 6.5 | 44.9 | 2.8 |
| 5uk0 | H | 4.8 | 8562 | 2.08 | 1.64 | 236 | 17.5 | 17.5 | 0.261 | 0.284 | 62.3 | 8.2 | 36 | 5.9 |
| 5uk0 | I | 4.8 | 8562 | 2.08 | 1.64 | 236 | 21.5 | 17.7 | 0.265 | 0.313 | 64.8 | 7.2 | 31.4 | 8.1 |
| 5uk1 | B | 4.8 | 8563 | 1.6 | 1.64 | 173 | 17.1 | 15.3 | 0.379 | 0.379 | 67.1 | 9.5 | 64.2 | 8.1 |
| 5uk1 | G | 4.8 | 8563 | 1.6 | 1.64 | 236 | 18.3 | 16.8 | 0.263 | 0.279 | 49.2 | 7.8 | 22 | 1.9 |
| 5uk1 | H | 4.8 | 8563 | 1.6 | 1.64 | 236 | 22.0 | 16.3 | 0.263 | 0.296 | 55.1 | 8.5 | 25.4 | 10 |
| 5uk1 | I | 4.8 | 8563 | 1.6 | 1.64 | 236 | 18.8 | 18.8 | 0.287 | 0.294 | 50 | 5.9 | 31.4 | 8.1 |
| 5uk2 | G | 4.8 | 8564 | 2.52 | 1.64 | 236 | 19.4 | 17.3 | 0.276 | 0.276 | 55.1 | 3.1 | 53.8 | 4.7 |
| 5uk2 | H | 4.8 | 8564 | 2.52 | 1.64 | 236 | 17.5 | 16.9 | 0.279 | 0.305 | 57.2 | 7.4 | 26.7 | 12.7 |
| 5uk2 | I | 4.8 | 8564 | 2.52 | 1.64 | 236 | 21.3 | 17.2 | 0.234 | 0.325 | 57.2 | 5.2 | 41.1 | 3.1 |
| 5up6 | G | 3.7 | 8584 | 0.02 | 1.3 | 230 | 4.8 | 4.6 | 0.920 | 0.920 | 94.8 | 54.6 | 81.3 | 40.1 |
| 5up6 | H | 3.7 | 8584 | 0.02 | 1.3 | 217 | 14.6 | 11.7 | 0.586 | 0.704 | 74.2 | 17.4 | 61.8 | 14.9 |
| 5upa | G | 4.1 | 8585 | 0.015 | 1.3 | 228 | 10.0 | 9.1 | 0.766 | 0.769 | 82.9 | 37.6 | 76.3 | 20.7 |
| 5upa | H | 4.1 | 8585 | 0.015 | 1.3 | 217 | 14.5 | 14.5 | 0.610 | 0.610 | 80.6 | 22.3 | 50.2 | 13.8 |
| 5upa | K | 4.1 | 8585 | 0.015 | 1.3 | 79 | 5.1 | 3.2 | 0.763 | 0.763 | 84.8 | 49.3 | 82.3 | 7.7 |
| 5upc | G | 4.4 | 8586 | 0.018 | 1.3 | 230 | 19.8 | 18.9 | 0.427 | 0.513 | 76.5 | 11.9 | 66.5 | 4.6 |
| 5upc | H | 4.4 | 8586 | 0.018 | 1.3 | 217 | 15.4 | 14.2 | 0.577 | 0.577 | 75.1 | 22.1 | 63.6 | 6.5 |
| 5uz5 | C | 3.7 | 8622 | 0.06 | 1.36 | 131 | 15.9 | 9.1 | 0.277 | 0.681 | 74 | 8.2 | 59.5 | 10.3 |
| 5uz5 | F | 3.7 | 8622 | 0.06 | 1.36 | 187 | 17.0 | 16.4 | 0.425 | 0.466 | 80.7 | 19.9 | 73.8 | 29.7 |
| 5uz5 | G | 3.7 | 8622 | 0.06 | 1.36 | 229 | 4.8 | 4.8 | 0.839 | 0.839 | 88.2 | 52 | 75.5 | 37 |
| 5uz5 | L | 3.7 | 8622 | 0.06 | 1.36 | 119 | 1.6 | 1.6 | 0.905 | 0.905 | 95.8 | 83.3 | 77.3 | 27.2 |
| 5uz5 | M | 3.7 | 8622 | 0.06 | 1.36 | 96 | 1.8 | 1.8 | 0.883 | 0.883 | 94.8 | 85.7 | 82.3 | 12.7 |
| 5uz5 | N | 3.7 | 8622 | 0.06 | 1.36 | 92 | 1.6 | 1.6 | 0.910 | 0.910 | 96.7 | 89.9 | 83.7 | 6.5 |
| 5uz5 | O | 3.7 | 8622 | 0.06 | 1.36 | 74 | 13.0 | 8.9 | 0.312 | 0.717 | 78.4 | 8.6 | 58.1 | 4.7 |
| 5uz5 | P | 3.7 | 8622 | 0.06 | 1.36 | 68 | 13.0 | 9.6 | 0.351 | 0.659 | 83.8 | 33.3 | 76.5 | 3.8 |
| 5uz5 | Q | 3.7 | 8622 | 0.06 | 1.36 | 75 | 7.7 | 4.9 | 0.770 | 0.770 | 88 | 31.8 | 64 | 10.4 |
| 5uz7 | N | 4.1 | 8623 | 0.05 | 1.06 | 128 | 15.4 | 11.2 | 0.327 | 0.615 | 81.2 | 6.7 | 81.2 | 4.8 |
| 5uz9 | C | 3.4 | 8624 | 0.0274 | 1.03 | 293 | 24.0 | 21.6 | 0.407 | 0.416 | 75.1 | 20.5 | 71.3 | 6.2 |
| 5uz9 | I | 3.4 | 8624 | 0.0274 | 1.03 | 76 | 6.9 | 4.5 | 0.637 | 0.762 | 84.2 | 59.4 | 77.6 | 8.5 |
| 5uz9 | J | 3.4 | 8624 | 0.0274 | 1.03 | 76 | 2.9 | 2.9 | 0.775 | 0.775 | 90.8 | 47.8 | 82.9 | 41.3 |
| 5v8l | G | 4.3 | 8643 | 0.03 | 1.31 | 121 | 16.8 | 14.7 | 0.404 | 0.467 | 75.2 | 8.8 | 35.5 | 11.6 |
| 5v8l | H | 4.3 | 8643 | 0.03 | 1.31 | 121 | 13.2 | 11.1 | 0.613 | 0.622 | 74.4 | 31.1 | 47.9 | 3.4 |
| 5v8l | I | 4.3 | 8643 | 0.03 | 1.31 | 121 | 9.9 | 6.8 | 0.756 | 0.769 | 85.1 | 21.4 | 52.9 | 4.7 |
| 5v8l | J | 4.3 | 8643 | 0.03 | 1.31 | 140 | 21.7 | 16.7 | 0.388 | 0.390 | 67.1 | 8.5 | 23.6 | 9.1 |
| 5v8l | K | 4.3 | 8643 | 0.03 | 1.31 | 98 | 13.8 | 11.8 | 0.303 | 0.406 | 75.5 | 10.8 | 59.2 | 8.6 |

| | | | | | | | | | | | | | | |
|---|---|---|---|---|---|---|---|---|---|---|---|---|---|---|
| 5v8l | L | 4.3 | 8643 | 0.03 | 1.31 | 98 | 13.4 | 11.4 | 0.289 | 0.394 | 69.4 | 7.4 | 54.1 | 7.5 |
| 5v8l | N | 4.3 | 8643 | 0.03 | 1.31 | 112 | 15.6 | 14.6 | 0.268 | 0.270 | 65.2 | 11 | 22.3 | 12 |
| 5v8m | H | 4.4 | 8644 | 0.035 | 1.31 | 121 | 13.7 | 12.8 | 0.342 | 0.489 | 75.2 | 3.3 | 49.6 | 3.3 |
| 5v8m | I | 4.4 | 8644 | 0.035 | 1.31 | 129 | 13.9 | 8.3 | 0.328 | 0.715 | 75.2 | 11.3 | 70.5 | 4.4 |
| 5v8m | J | 4.4 | 8644 | 0.035 | 1.31 | 129 | 5.4 | 5.1 | 0.776 | 0.776 | 82.9 | 49.5 | 58.1 | 2.7 |
| 5v8m | L | 4.4 | 8644 | 0.035 | 1.31 | 98 | 13.5 | 13.5 | 0.312 | 0.322 | 73.5 | 6.9 | 53.1 | 3.8 |
| 5v8m | R | 4.4 | 8644 | 0.035 | 1.31 | 121 | 5.9 | 4.9 | 0.575 | 0.803 | 84.3 | 11.8 | 77.7 | 7.4 |
| 5v8m | S | 4.4 | 8644 | 0.035 | 1.31 | 121 | 15.1 | 14.4 | 0.339 | 0.493 | 76 | 9.8 | 43.8 | 7.5 |
| 5v8m | T | 4.4 | 8644 | 0.035 | 1.31 | 98 | 14.0 | 12.7 | 0.340 | 0.355 | 69.4 | 11.8 | 44.9 | 15.9 |
| 5v8m | U | 4.4 | 8644 | 0.035 | 1.31 | 98 | 14.7 | 13.1 | 0.351 | 0.434 | 68.4 | 14.9 | 41.8 | 2.4 |
| 5vai | N | 4.1 | 8653 | 0.055 | 1 | 128 | 13.7 | 12.3 | 0.329 | 0.436 | 64.1 | 7.3 | 39.1 | 6 |
| 5vfo | g | 3.5 | 8662 | 0.01 | 0.75 | 240 | 16.9 | 16.4 | 0.363 | 0.676 | 77.5 | 7 | 77.5 | 28 |
| 5vfo | G | 3.5 | 8662 | 0.01 | 0.75 | 239 | 3.0 | 3.0 | 0.892 | 0.892 | 90.8 | 54.8 | 87.4 | 17.7 |
| 5vfo | h | 3.5 | 8662 | 0.01 | 0.75 | 232 | 12.3 | 12.3 | 0.631 | 0.631 | 81 | 25.5 | 69.4 | 44.1 |
| 5vfo | H | 3.5 | 8662 | 0.01 | 0.75 | 230 | 20.2 | 18.2 | 0.554 | 0.554 | 77 | 32.2 | 78.7 | 31.5 |
| 5vfo | i | 3.5 | 8662 | 0.01 | 0.75 | 250 | 18.8 | 16.4 | 0.556 | 0.556 | 76.4 | 22 | 79.2 | 5.6 |
| 5vfo | I | 3.5 | 8662 | 0.01 | 0.75 | 248 | 3.8 | 3.8 | 0.894 | 0.894 | 91.5 | 62.6 | 84.7 | 20.5 |
| 5vfo | j | 3.5 | 8662 | 0.01 | 0.75 | 239 | 8.9 | 6.8 | 0.794 | 0.799 | 79.9 | 36.6 | 77.4 | 35.1 |
| 5vfo | J | 3.5 | 8662 | 0.01 | 0.75 | 239 | 7.0 | 4.5 | 0.828 | 0.835 | 83.7 | 31.5 | 85.4 | 22.5 |
| 5vfo | k | 3.5 | 8662 | 0.01 | 0.75 | 228 | 5.1 | 5.1 | 0.870 | 0.870 | 88.6 | 33.7 | 81.6 | 22.6 |
| 5vfo | K | 3.5 | 8662 | 0.01 | 0.75 | 228 | 4.4 | 4.4 | 0.881 | 0.881 | 89.9 | 55.6 | 83.8 | 31.4 |
| 5vfo | l | 3.5 | 8662 | 0.01 | 0.75 | 238 | 19.5 | 16.7 | 0.282 | 0.610 | 73.5 | 6.3 | 85.3 | 8.4 |
| 5vfo | L | 3.5 | 8662 | 0.01 | 0.75 | 238 | 3.5 | 3.5 | 0.899 | 0.899 | 93.7 | 61.9 | 86.1 | 4.9 |
| 5vfo | q | 3.5 | 8662 | 0.01 | 0.75 | 199 | 6.0 | 4.5 | 0.886 | 0.886 | 92 | 48.1 | 90.5 | 31.1 |
| 5vfo | Q | 3.5 | 8662 | 0.01 | 0.75 | 199 | 4.5 | 3.7 | 0.899 | 0.899 | 93 | 64.3 | 89.4 | 4.5 |
| 5vfr | b | 4.9 | 8665 | 0.008 | 0.75 | 191 | 17.4 | 15.2 | 0.280 | 0.320 | 51.3 | 5.1 | 24.1 | 8.7 |
| 5vfr | g | 4.9 | 8665 | 0.008 | 0.75 | 240 | 18.2 | 18.2 | 0.316 | 0.316 | 62.1 | 6.7 | 56.7 | 5.1 |
| 5vfr | G | 4.9 | 8665 | 0.008 | 0.75 | 239 | 18.8 | 16.2 | 0.274 | 0.480 | 59.8 | 7 | 55.6 | 7.5 |
| 5vfr | h | 4.9 | 8665 | 0.008 | 0.75 | 232 | 20.4 | 19.3 | 0.282 | 0.320 | 65.9 | 5.2 | 65.5 | 9.2 |
| 5vfr | H | 4.9 | 8665 | 0.008 | 0.75 | 230 | 19.0 | 17.6 | 0.266 | 0.432 | 65.7 | 5.3 | 37 | 8.2 |
| 5vfr | i | 4.9 | 8665 | 0.008 | 0.75 | 250 | 20.1 | 19.9 | 0.252 | 0.389 | 61.2 | 11.1 | 41.6 | 4.8 |
| 5vfr | I | 4.9 | 8665 | 0.008 | 0.75 | 248 | 21.0 | 17.6 | 0.286 | 0.359 | 59.7 | 7.4 | 55.2 | 5.1 |
| 5vfr | j | 4.9 | 8665 | 0.008 | 0.75 | 239 | 19.0 | 15.7 | 0.269 | 0.288 | 67.8 | 4.3 | 56.5 | 8.1 |
| 5vfr | J | 4.9 | 8665 | 0.008 | 0.75 | 239 | 19.7 | 16.7 | 0.350 | 0.415 | 68.2 | 12.9 | 57.7 | 8 |
| 5vfr | k | 4.9 | 8665 | 0.008 | 0.75 | 228 | 18.7 | 17.0 | 0.298 | 0.339 | 66.7 | 11.8 | 28.9 | 15.2 |
| 5vfr | K | 4.9 | 8665 | 0.008 | 0.75 | 228 | 18.2 | 16.7 | 0.354 | 0.412 | 61.4 | 2.1 | 67.1 | 9.2 |
| 5vfr | l | 4.9 | 8665 | 0.008 | 0.75 | 238 | 18.4 | 17.8 | 0.270 | 0.359 | 66 | 8.9 | 58 | 9.4 |

| | | | | | | | | | | | | | | |
|---|---|---|---|---|---|---|---|---|---|---|---|---|---|---|
| 5vfr | L | 4.9 | 8665 | 0.008 | 0.75 | 238 | 15.4 | 15.4 | 0.458 | 0.463 | 66 | 15.9 | 70.2 | 6 |
| 5vfr | q | 4.9 | 8665 | 0.008 | 0.75 | 199 | 14.4 | 14.4 | 0.475 | 0.486 | 62.3 | 15.3 | 45.2 | 6.7 |
| 5vfr | Q | 4.9 | 8665 | 0.008 | 0.75 | 199 | 18.1 | 16.8 | 0.340 | 0.351 | 66.3 | 8.3 | 77.9 | 7.1 |
| 5vgz | b | 4.5 | 8672 | 0.004 | 0.98 | 191 | 16.7 | 16.1 | 0.325 | 0.365 | 64.9 | 8.1 | 34.6 | 7.6 |
| 5vgz | d | 4.5 | 8672 | 0.004 | 0.98 | 257 | 21.6 | 12.5 | 0.297 | 0.544 | 58.8 | 4.6 | 44.7 | 7.8 |
| 5vhy | E | 4.6 | 8687 | 0.028 | 1.04 | 179 | 10.2 | 9.9 | 0.555 | 0.555 | 70.9 | 6.3 | 65.4 | 10.3 |
| 5vhy | F | 4.6 | 8687 | 0.028 | 1.04 | 179 | 22.7 | 14.9 | 0.442 | 0.517 | 70.4 | 11.1 | 57.5 | 7.8 |
| 5vn3 | A | 3.7 | 8713 | 0.07 | 1.31 | 136 | 9.8 | 9.8 | 0.617 | 0.683 | 79.4 | 52.8 | 72.8 | 28.3 |
| 5vn3 | B | 3.7 | 8713 | 0.07 | 1.31 | 136 | 11.7 | 10.0 | 0.599 | 0.653 | 79.4 | 43.5 | 78.7 | 52.3 |
| 5vn3 | C | 3.7 | 8713 | 0.07 | 1.31 | 176 | 20.3 | 16.1 | 0.490 | 0.630 | 74.4 | 11.5 | 40.9 | 13.9 |
| 5vn3 | D | 3.7 | 8713 | 0.07 | 1.31 | 136 | 11.6 | 10.2 | 0.622 | 0.687 | 82.4 | 43.8 | 72.8 | 32.3 |
| 5vn3 | E | 3.7 | 8713 | 0.07 | 1.31 | 176 | 20.0 | 18.2 | 0.489 | 0.599 | 74.4 | 8.4 | 29 | 2 |
| 5vn3 | F | 3.7 | 8713 | 0.07 | 1.31 | 176 | 19.2 | 16.8 | 0.335 | 0.457 | 69.9 | 6.5 | 42 | 14.9 |
| 5vn3 | H | 3.7 | 8713 | 0.07 | 1.31 | 127 | 18.4 | 15.4 | 0.351 | 0.382 | 72.4 | 13 | 43.3 | 10.9 |
| 5vn3 | K | 3.7 | 8713 | 0.07 | 1.31 | 127 | 14.1 | 13.2 | 0.464 | 0.488 | 72.4 | 13 | 53.5 | 4.4 |
| 5vn3 | L | 3.7 | 8713 | 0.07 | 1.31 | 110 | 15.3 | 14.8 | 0.275 | 0.373 | 67.3 | 6.8 | 42.7 | 17 |
| 5vn3 | M | 3.7 | 8713 | 0.07 | 1.31 | 127 | 15.1 | 13.9 | 0.415 | 0.551 | 78 | 6.1 | 34.6 | 11.4 |
| 5vn3 | N | 3.7 | 8713 | 0.07 | 1.31 | 110 | 12.6 | 12.6 | 0.363 | 0.421 | 69.1 | 3.9 | 47.3 | 7.7 |
| 5vn3 | O | 3.7 | 8713 | 0.07 | 1.31 | 110 | 16.4 | 14.1 | 0.299 | 0.299 | 67.3 | 10.8 | 32.7 | 13.9 |
| 5vn8 | A | 3.6 | 8717 | 0.08 | 1.31 | 139 | 10.4 | 10.4 | 0.609 | 0.684 | 79.1 | 59.1 | 82 | 58.8 |
| 5vn8 | B | 3.6 | 8717 | 0.08 | 1.31 | 139 | 10.7 | 9.6 | 0.623 | 0.623 | 85.6 | 73.1 | 78.4 | 62.4 |
| 5vn8 | F | 3.6 | 8717 | 0.08 | 1.31 | 127 | 17.6 | 12.7 | 0.383 | 0.529 | 73.2 | 7.5 | 46.5 | 5.1 |
| 5vn8 | H | 3.6 | 8717 | 0.08 | 1.31 | 127 | 16.0 | 15.0 | 0.413 | 0.422 | 72.4 | 7.6 | 63.8 | 4.9 |
| 5vn8 | I | 3.6 | 8717 | 0.08 | 1.31 | 127 | 17.9 | 14.0 | 0.301 | 0.493 | 72.4 | 6.5 | 41.7 | 5.7 |
| 5vn8 | J | 3.6 | 8717 | 0.08 | 1.31 | 108 | 13.8 | 12.2 | 0.442 | 0.442 | 67.6 | 6.8 | 32.4 | 5.7 |
| 5vn8 | K | 3.6 | 8717 | 0.08 | 1.31 | 108 | 12.2 | 12.2 | 0.333 | 0.333 | 73.1 | 5.1 | 43.5 | 2.1 |
| 5vt0 | G | 3.8 | 8732 | 0.014 | 1.3 | 228 | 2.9 | 2.9 | 0.918 | 0.918 | 94.3 | 66.5 | 82.5 | 8.5 |
| 5vt0 | H | 3.8 | 8732 | 0.014 | 1.3 | 217 | 12.2 | 11.9 | 0.559 | 0.615 | 71.9 | 44.9 | 63.6 | 29.7 |
| 5vt0 | K | 3.8 | 8732 | 0.014 | 1.3 | 79 | 1.3 | 1.3 | 0.889 | 0.889 | 97.5 | 97.4 | 83.5 | 9.1 |
| 5vy3 | 0 | 3.1 | 8741 | 0.06 | 0.91 | 221 | 3.2 | 3.2 | 0.907 | 0.907 | 91 | 67.7 | 86 | 62.1 |
| 5vy3 | A | 3.1 | 8741 | 0.06 | 0.91 | 221 | 8.3 | 8.0 | 0.790 | 0.808 | 86.4 | 72.3 | 88.2 | 62.6 |
| 5vy3 | C | 3.1 | 8741 | 0.06 | 0.91 | 221 | 4.3 | 4.3 | 0.922 | 0.922 | 93.2 | 66.5 | 85.5 | 70.4 |
| 5vy3 | E | 3.1 | 8741 | 0.06 | 0.91 | 221 | 5.3 | 5.3 | 0.824 | 0.828 | 91 | 68.7 | 89.1 | 65 |
| 5vy3 | G | 3.1 | 8741 | 0.06 | 0.91 | 221 | 1.9 | 1.9 | 0.938 | 0.938 | 95 | 81.4 | 85.1 | 64.9 |
| 5vy3 | I | 3.1 | 8741 | 0.06 | 0.91 | 221 | 2.2 | 2.2 | 0.941 | 0.941 | 95.9 | 84.4 | 86 | 62.1 |
| 5vy3 | K | 3.1 | 8741 | 0.06 | 0.91 | 221 | 8.4 | 7.1 | 0.786 | 0.831 | 87.8 | 63.9 | 87.3 | 61.7 |
| 5vy3 | M | 3.1 | 8741 | 0.06 | 0.91 | 221 | 2.5 | 2.5 | 0.935 | 0.935 | 94.1 | 85.1 | 86.9 | 83.3 |

| | | | | | | | | | | | | | | |
|---|---|---|---|---|---|---|---|---|---|---|---|---|---|---|
| 5vy3 | O | 3.1 | 8741 | 0.06 | 0.91 | 221 | 6.1 | 6.1 | 0.829 | 0.830 | 92.3 | 79.9 | 89.6 | 72.2 |
| 5vy3 | Q | 3.1 | 8741 | 0.06 | 0.91 | 221 | 5.3 | 5.3 | 0.827 | 0.827 | 91.9 | 71.4 | 88.2 | 62.6 |
| 5vy3 | S | 3.1 | 8741 | 0.06 | 0.91 | 221 | 2.4 | 2.4 | 0.928 | 0.928 | 93.7 | 77.3 | 84.6 | 62.6 |
| 5vy3 | U | 3.1 | 8741 | 0.06 | 0.91 | 221 | 6.6 | 4.3 | 0.799 | 0.897 | 88.2 | 68.7 | 83.7 | 55.7 |
| 5vy3 | W | 3.1 | 8741 | 0.06 | 0.91 | 221 | 3.3 | 3.3 | 0.911 | 0.911 | 91 | 75.6 | 83.3 | 54.9 |
| 5vy3 | Y | 3.1 | 8741 | 0.06 | 0.91 | 221 | 9.1 | 8.1 | 0.829 | 0.829 | 83.3 | 73.4 | 90 | 77.9 |
| 5vy4 | 0 | 3.3 | 8742 | 0.06 | 0.91 | 221 | 6.1 | 6.1 | 0.794 | 0.811 | 87.3 | 47.7 | 86.4 | 70.7 |
| 5vy4 | A | 3.3 | 8742 | 0.06 | 0.91 | 221 | 1.8 | 1.8 | 0.943 | 0.943 | 96.4 | 81.2 | 89.6 | 71.2 |
| 5vy4 | C | 3.3 | 8742 | 0.06 | 0.91 | 221 | 5.0 | 5.0 | 0.828 | 0.832 | 90 | 73.9 | 88.7 | 74.5 |
| 5vy4 | E | 3.3 | 8742 | 0.06 | 0.91 | 221 | 3.6 | 3.6 | 0.918 | 0.918 | 93.2 | 62.6 | 86.9 | 55.2 |
| 5vy4 | G | 3.3 | 8742 | 0.06 | 0.91 | 221 | 4.1 | 4.1 | 0.932 | 0.932 | 94.6 | 86.1 | 86.4 | 64.9 |
| 5vy4 | I | 3.3 | 8742 | 0.06 | 0.91 | 221 | 2.1 | 2.1 | 0.950 | 0.950 | 96.4 | 80.3 | 88.7 | 74 |
| 5vy4 | K | 3.3 | 8742 | 0.06 | 0.91 | 221 | 2.6 | 2.6 | 0.940 | 0.940 | 94.6 | 86.1 | 84.2 | 37.1 |
| 5vy4 | M | 3.3 | 8742 | 0.06 | 0.91 | 221 | 1.3 | 1.3 | 0.956 | 0.956 | 97.7 | 94.4 | 86 | 33.2 |
| 5vy4 | O | 3.3 | 8742 | 0.06 | 0.91 | 221 | 3.2 | 3.2 | 0.934 | 0.934 | 95.5 | 92.9 | 86.9 | 69.8 |
| 5vy4 | Q | 3.3 | 8742 | 0.06 | 0.91 | 221 | 2.8 | 2.8 | 0.903 | 0.903 | 91.9 | 71.9 | 87.8 | 52.1 |
| 5vy4 | S | 3.3 | 8742 | 0.06 | 0.91 | 221 | 2.2 | 2.2 | 0.933 | 0.933 | 94.6 | 82.3 | 87.3 | 51.8 |
| 5vy4 | U | 3.3 | 8742 | 0.06 | 0.91 | 221 | 8.7 | 7.5 | 0.797 | 0.829 | 85.5 | 66.7 | 86 | 75.8 |
| 5vy4 | W | 3.3 | 8742 | 0.06 | 0.91 | 221 | 2.7 | 2.7 | 0.909 | 0.909 | 92.3 | 73.5 | 87.8 | 45.4 |
| 5vy4 | Y | 3.3 | 8742 | 0.06 | 0.91 | 221 | 10.5 | 7.8 | 0.844 | 0.844 | 85.1 | 61.2 | 88.7 | 75 |
| 5w5y | E | 3.8 | 8771 | 0.04 | 1.3 | 215 | 11.4 | 8.4 | 0.724 | 0.736 | 80.9 | 35.1 | 68.8 | 7.4 |
| 5w5y | F | 3.8 | 8771 | 0.04 | 1.3 | 83 | 3.5 | 3.5 | 0.815 | 0.815 | 89.2 | 64.9 | 84.3 | 38.6 |
| 5w5y | G | 3.8 | 8771 | 0.04 | 1.3 | 201 | 20.9 | 17.8 | 0.326 | 0.425 | 70.1 | 5 | 46.8 | 6.4 |
| 5w5y | H | 3.8 | 8771 | 0.04 | 1.3 | 133 | 14.5 | 11.7 | 0.344 | 0.503 | 74.4 | 5.1 | 61.7 | 7.3 |
| 5w5y | K | 3.8 | 8771 | 0.04 | 1.3 | 103 | 3.3 | 3.3 | 0.855 | 0.855 | 91.3 | 45.7 | 90.3 | 17.2 |
| 5w64 | E | 4.2 | 8774 | 0.04 | 1.3 | 215 | 13.5 | 13.5 | 0.557 | 0.557 | 71.2 | 25.5 | 35.3 | 3.9 |
| 5w64 | F | 4.2 | 8774 | 0.04 | 1.3 | 83 | 4.6 | 4.6 | 0.715 | 0.725 | 83.1 | 39.1 | 73.5 | 9.8 |
| 5w64 | G | 4.2 | 8774 | 0.04 | 1.3 | 201 | 17.8 | 16.5 | 0.370 | 0.370 | 63.7 | 13.3 | 31.8 | 1.6 |
| 5w64 | H | 4.2 | 8774 | 0.04 | 1.3 | 133 | 14.5 | 13.9 | 0.236 | 0.296 | 69.2 | 9.8 | 40.6 | 5.6 |
| 5w65 | E | 4.3 | 8775 | 0.04 | 1.3 | 215 | 14.7 | 11.5 | 0.541 | 0.689 | 77.2 | 24.7 | 68.8 | 4.7 |
| 5w65 | F | 4.3 | 8775 | 0.04 | 1.3 | 83 | 2.3 | 2.3 | 0.821 | 0.821 | 89.2 | 55.4 | 84.3 | 5.7 |
| 5w65 | G | 4.3 | 8775 | 0.04 | 1.3 | 201 | 17.4 | 17.0 | 0.424 | 0.441 | 69.2 | 7.9 | 41.8 | 6 |
| 5w65 | H | 4.3 | 8775 | 0.04 | 1.3 | 133 | 16.5 | 14.9 | 0.277 | 0.405 | 73.7 | 11.2 | 56.4 | 12 |
| 5w65 | K | 4.3 | 8775 | 0.04 | 1.3 | 103 | 2.9 | 2.9 | 0.849 | 0.849 | 91.3 | 50 | 86.4 | 31.5 |
| 5w66 | E | 3.9 | 8776 | 0.04 | 1.3 | 215 | 15.6 | 14.4 | 0.505 | 0.541 | 71.2 | 15.7 | 60 | 4.7 |
| 5w66 | F | 3.9 | 8776 | 0.04 | 1.3 | 83 | 3.6 | 3.6 | 0.797 | 0.797 | 89.2 | 56.8 | 78.3 | 49.2 |
| 5w66 | G | 3.9 | 8776 | 0.04 | 1.3 | 201 | 25.8 | 18.4 | 0.215 | 0.449 | 64.7 | 6.2 | 22.9 | 4.3 |

| | | | | | | | | | | | | | | |
|---|---|---|---|---|---|---|---|---|---|---|---|---|---|---|
| 5w66 | H | 3.9 | 8776 | 0.04 | 1.3 | 133 | 15.6 | 14.1 | 0.263 | 0.394 | 66.9 | 6.7 | 24.1 | 9.4 |
| 5w66 | K | 3.9 | 8776 | 0.04 | 1.3 | 103 | 8.8 | 8.8 | 0.548 | 0.589 | 72.8 | 6.7 | 75.7 | 7.7 |
| 5w9h | B | 4 | 8783 | 0.015 | 1.02 | 119 | 14.3 | 13.4 | 0.299 | 0.339 | 63.9 | 3.9 | 41.2 | 18.4 |
| 5w9h | C | 4 | 8783 | 0.015 | 1.02 | 111 | 12.3 | 12.3 | 0.388 | 0.417 | 55.9 | 3.2 | 56.8 | 6.3 |
| 5w9h | E | 4 | 8783 | 0.015 | 1.02 | 119 | 15.5 | 14.5 | 0.259 | 0.324 | 67.2 | 13.8 | 38.7 | 6.5 |
| 5w9h | F | 4 | 8783 | 0.015 | 1.02 | 111 | 15.1 | 13.9 | 0.285 | 0.286 | 64.9 | 11.1 | 36.9 | 9.8 |
| 5w9h | H | 4 | 8783 | 0.015 | 1.02 | 119 | 16.5 | 13.7 | 0.368 | 0.368 | 71.4 | 5.9 | 35.3 | 9.5 |
| 5w9h | I | 4 | 8783 | 0.015 | 1.02 | 111 | 15.8 | 12.5 | 0.248 | 0.292 | 57.7 | 7.8 | 37.8 | 9.5 |
| 5w9i | C | 3.6 | 8784 | 0.015 | 1.02 | 119 | 14.8 | 12.4 | 0.531 | 0.531 | 80.7 | 6.2 | 60.5 | 8.3 |
| 5w9i | D | 3.6 | 8784 | 0.015 | 1.02 | 111 | 14.1 | 12.3 | 0.313 | 0.615 | 76.6 | 11.8 | 41.4 | 4.3 |
| 5w9i | G | 3.6 | 8784 | 0.015 | 1.02 | 119 | 9.9 | 7.9 | 0.668 | 0.722 | 83.2 | 43.4 | 68.9 | 6.1 |
| 5w9i | H | 3.6 | 8784 | 0.015 | 1.02 | 111 | 15.8 | 11.7 | 0.443 | 0.443 | 68.5 | 10.5 | 64 | 11.3 |
| 5w9i | K | 3.6 | 8784 | 0.015 | 1.02 | 119 | 6.9 | 6.9 | 0.767 | 0.767 | 85.7 | 45.1 | 64.7 | 7.8 |
| 5w9i | L | 3.6 | 8784 | 0.015 | 1.02 | 111 | 15.0 | 11.2 | 0.275 | 0.530 | 70.3 | 12.8 | 66.7 | 5.4 |
| 5w9j | B | 4.8 | 8785 | 0.015 | 1.02 | 119 | 16.9 | 15.3 | 0.225 | 0.332 | 52.1 | 6.5 | 34.5 | 7.3 |
| 5w9j | C | 4.8 | 8785 | 0.015 | 1.02 | 111 | 15.8 | 14.4 | 0.245 | 0.291 | 65.8 | 8.2 | 29.7 | 12.1 |
| 5w9j | E | 4.8 | 8785 | 0.015 | 1.02 | 119 | 15.7 | 15.0 | 0.257 | 0.292 | 61.3 | 2.7 | 44.5 | 3.8 |
| 5w9j | F | 4.8 | 8785 | 0.015 | 1.02 | 111 | 15.4 | 13.8 | 0.238 | 0.281 | 62.2 | 0 | 39.6 | 11.4 |
| 5w9j | H | 4.8 | 8785 | 0.015 | 1.02 | 119 | 16.8 | 15.1 | 0.246 | 0.282 | 58.8 | 5.7 | 35.3 | 9.5 |
| 5w9j | I | 4.8 | 8785 | 0.015 | 1.02 | 111 | 14.3 | 13.3 | 0.285 | 0.296 | 57.7 | 6.2 | 44.1 | 8.2 |
| 5w9k | B | 4.6 | 8786 | 0.015 | 1.02 | 119 | 15.3 | 14.1 | 0.267 | 0.300 | 58.8 | 12.9 | 41.2 | 4.1 |
| 5w9k | C | 4.6 | 8786 | 0.015 | 1.02 | 111 | 15.8 | 15.2 | 0.283 | 0.310 | 62.2 | 4.3 | 26.1 | 6.9 |
| 5w9k | E | 4.6 | 8786 | 0.015 | 1.02 | 119 | 15.8 | 14.2 | 0.270 | 0.302 | 62.2 | 2.7 | 26.9 | 12.5 |
| 5w9k | F | 4.6 | 8786 | 0.015 | 1.02 | 111 | 16.7 | 14.3 | 0.265 | 0.322 | 64 | 5.6 | 45 | 10 |
| 5w9k | H | 4.6 | 8786 | 0.015 | 1.02 | 119 | 16.7 | 14.7 | 0.272 | 0.299 | 65.5 | 5.1 | 32.8 | 5.1 |
| 5w9l | E | 4.8 | 8787 | 0.015 | 1.02 | 119 | 16.4 | 13.9 | 0.293 | 0.357 | 68.1 | 8.6 | 34.5 | 12.2 |
| 5w9l | F | 4.8 | 8787 | 0.015 | 1.02 | 111 | 14.9 | 14.3 | 0.264 | 0.292 | 64.9 | 4.2 | 38.7 | 14 |
| 5w9l | H | 4.8 | 8787 | 0.015 | 1.02 | 119 | 14.7 | 14.4 | 0.260 | 0.305 | 58.8 | 4.3 | 36.1 | 9.3 |
| 5w9l | I | 4.8 | 8787 | 0.015 | 1.02 | 111 | 16.6 | 14.7 | 0.260 | 0.293 | 62.2 | 2.9 | 31.5 | 14.3 |
| 5w9m | B | 4.7 | 8788 | 0.015 | 1.02 | 119 | 13.4 | 13.4 | 0.276 | 0.276 | 67.2 | 10 | 38.7 | 0 |
| 5w9m | C | 4.7 | 8788 | 0.015 | 1.02 | 111 | 15.9 | 13.3 | 0.292 | 0.326 | 57.7 | 6.2 | 27.9 | 3.2 |
| 5w9m | H | 4.7 | 8788 | 0.015 | 1.02 | 119 | 16.9 | 14.8 | 0.298 | 0.358 | 68.1 | 2.5 | 45.4 | 5.6 |
| 5w9m | I | 4.7 | 8788 | 0.015 | 1.02 | 111 | 15.1 | 14.7 | 0.240 | 0.283 | 65.8 | 4.1 | 41.4 | 2.2 |
| 5w9o | B | 4.5 | 8790 | 0.015 | 1.02 | 119 | 13.6 | 13.2 | 0.278 | 0.362 | 67.2 | 15 | 30.3 | 5.6 |
| 5w9o | C | 4.5 | 8790 | 0.015 | 1.02 | 111 | 14.7 | 13.8 | 0.234 | 0.293 | 59.5 | 4.5 | 22.5 | 12 |
| 5w9o | E | 4.5 | 8790 | 0.015 | 1.02 | 119 | 17.5 | 14.5 | 0.299 | 0.304 | 67.2 | 5 | 34.5 | 7.3 |
| 5w9o | F | 4.5 | 8790 | 0.015 | 1.02 | 111 | 14.2 | 14.2 | 0.251 | 0.315 | 63.1 | 5.7 | 41.4 | 19.6 |

| | | | | | | | | | | | | | | |
|---|---|---|---|---|---|---|---|---|---|---|---|---|---|---|
| 5w9o | H | 4.5 | 8790 | 0.015 | 1.02 | 119 | 15.6 | 14.5 | 0.341 | 0.391 | 73.1 | 5.7 | 42.9 | 11.8 |
| 5w9o | I | 4.5 | 8790 | 0.015 | 1.02 | 111 | 14.3 | 13.5 | 0.288 | 0.302 | 61.3 | 4.4 | 32.4 | 8.3 |
| 5w9p | D | 4 | 8791 | 0.015 | 1.02 | 119 | 14.4 | 12.8 | 0.497 | 0.669 | 79 | 14.9 | 58 | 5.8 |
| 5w9p | E | 4 | 8791 | 0.015 | 1.02 | 111 | 14.6 | 13.2 | 0.311 | 0.369 | 68.5 | 13.2 | 45 | 8 |
| 5w9p | F | 4 | 8791 | 0.015 | 1.02 | 119 | 16.2 | 13.2 | 0.457 | 0.515 | 82.4 | 5.1 | 67.2 | 3.8 |
| 5w9p | G | 4 | 8791 | 0.015 | 1.02 | 111 | 16.6 | 12.7 | 0.229 | 0.293 | 63.1 | 4.3 | 38.7 | 4.7 |
| 5w9p | K | 4 | 8791 | 0.015 | 1.02 | 119 | 16.3 | 13.4 | 0.334 | 0.527 | 84.9 | 12.9 | 68.9 | 1.2 |
| 5w9p | L | 4 | 8791 | 0.015 | 1.02 | 111 | 13.3 | 13.0 | 0.301 | 0.388 | 63.1 | 8.6 | 47.7 | 5.7 |
| 5wc0 | A | 4.4 | 8794 | 0.043 | 1.31 | 275 | 28.1 | 19.3 | 0.334 | 0.407 | 71.3 | 4.6 | 57.1 | 5.7 |
| 5wc0 | B | 4.4 | 8794 | 0.043 | 1.31 | 277 | 23.8 | 18.5 | 0.367 | 0.573 | 70.4 | 11.3 | 61 | 5.9 |
| 5wc0 | C | 4.4 | 8794 | 0.043 | 1.31 | 277 | 17.8 | 17.8 | 0.480 | 0.497 | 70 | 17 | 59.2 | 7.3 |
| 5wc0 | D | 4.4 | 8794 | 0.043 | 1.31 | 277 | 22.6 | 20.7 | 0.363 | 0.479 | 78 | 7.9 | 65.7 | 6.6 |
| 5wc0 | E | 4.4 | 8794 | 0.043 | 1.31 | 277 | 25.7 | 21.2 | 0.481 | 0.487 | 69.3 | 26.6 | 66.8 | 1.6 |
| 5wc0 | F | 4.4 | 8794 | 0.043 | 1.31 | 277 | 21.8 | 17.3 | 0.285 | 0.390 | 65.7 | 5.5 | 43.7 | 10.7 |
| 5wfe | A | 3.6 | 8827 | 0.8 | 1.07 | 262 | 18.7 | 7.9 | 0.481 | 0.757 | 72.1 | 7.4 | 64.5 | 24.3 |
| 5wfe | B | 3.6 | 8827 | 0.8 | 1.07 | 279 | 20.1 | 8.5 | 0.303 | 0.704 | 73.5 | 11.2 | 69.5 | 20.1 |
| 5wfe | C | 3.6 | 8827 | 0.8 | 1.07 | 256 | 18.5 | 15.2 | 0.389 | 0.417 | 60.5 | 6.5 | 48.8 | 4.8 |
| 5wfe | F | 3.6 | 8827 | 0.8 | 1.07 | 94 | 12.0 | 11.5 | 0.311 | 0.364 | 85.1 | 18.8 | 60.6 | 29.8 |
| 5x8r | f | 3.7 | 6710 | 0.05 | 1.05 | 111 | 15.7 | 9.7 | 0.320 | 0.697 | 80.2 | 10.1 | 53.2 | 5.1 |
| 5xlp | C | 4.2 | 6731 | 0.036 | 1.31 | 287 | 21.0 | 21.0 | 0.269 | 0.426 | 64.1 | 8.7 | 44.9 | 8.5 |
| 5xnm | o | 3.2 | 6742 | 0.026 | 1.04 | 214 | 18.7 | 18.0 | 0.435 | 0.481 | 74.8 | 6.2 | 56.5 | 7.4 |
| 5xnm | O | 3.2 | 6742 | 0.026 | 1.04 | 214 | 20.6 | 17.0 | 0.436 | 0.493 | 71 | 12.5 | 55.1 | 5.9 |
| 5xnm | y | 3.2 | 6742 | 0.026 | 1.04 | 219 | 1.8 | 1.8 | 0.931 | 0.931 | 95 | 77.9 | 80.8 | 48.6 |
| 5xon | C | 3.8 | 6747 | 0.0666 | 1.533 | 263 | 2.5 | 2.5 | 0.923 | 0.923 | 94.7 | 75.9 | 87.5 | 27.8 |
| 5xon | E | 3.8 | 6747 | 0.0666 | 1.533 | 213 | 3.0 | 3.0 | 0.894 | 0.894 | 90.6 | 63.2 | 72.8 | 21.3 |
| 5xon | F | 3.8 | 6747 | 0.0666 | 1.533 | 84 | 3.3 | 3.3 | 0.855 | 0.855 | 92.9 | 52.6 | 85.7 | 26.4 |
| 5xon | H | 3.8 | 6747 | 0.0666 | 1.533 | 133 | 18.3 | 9.5 | 0.393 | 0.614 | 85.7 | 6.1 | 78.9 | 13.3 |
| 5xon | K | 3.8 | 6747 | 0.0666 | 1.533 | 113 | 2.1 | 2.1 | 0.907 | 0.907 | 95.6 | 78.7 | 88.5 | 38 |
| 5xsy | B | 4 | 6770 | 0.1 | 1.338 | 172 | 15.7 | 15.7 | 0.493 | 0.653 | 69.8 | 6.7 | 30.8 | 9.4 |
| 5xtb | F | 3.4 | 6771 | 0.0516 | 1.083 | 83 | 2.9 | 2.9 | 0.798 | 0.798 | 90.4 | 70.7 | 63.9 | 7.5 |
| 5xtb | O | 3.4 | 6771 | 0.0516 | 1.083 | 212 | 2.7 | 2.7 | 0.905 | 0.905 | 92.9 | 64 | 71.7 | 5.9 |
| 5xtc | c | 3.7 | 6772 | 0.0677 | 1.083 | 153 | 2.0 | 2.0 | 0.907 | 0.907 | 94.1 | 80.6 | 83 | 58.3 |
| 5xtc | d | 3.7 | 6772 | 0.0677 | 1.083 | 171 | 2.5 | 2.5 | 0.914 | 0.914 | 93 | 74.8 | 88.9 | 53.9 |
| 5xtc | p | 3.7 | 6772 | 0.0677 | 1.083 | 172 | 2.8 | 2.8 | 0.927 | 0.927 | 94.8 | 81 | 84.3 | 9.7 |
| 5xtc | u | 3.7 | 6772 | 0.0677 | 1.083 | 169 | 3.3 | 3.3 | 0.866 | 0.866 | 88.8 | 48.7 | 23.7 | 2.5 |
| 5xtc | V | 3.7 | 6772 | 0.0677 | 1.083 | 140 | 3.2 | 3.2 | 0.859 | 0.859 | 91.4 | 48.4 | 74.3 | 6.7 |
| 5xtd | c | 3.7 | 6773 | 0.0525 | 1.083 | 153 | 2.1 | 2.1 | 0.888 | 0.888 | 93.5 | 83.9 | 79.1 | 10.7 |

| | | | | | | | | | | | | | | |
|---|---|---|---|---|---|---|---|---|---|---|---|---|---|---|
| 5xtd | C | 3.7 | 6773 | 0.0525 | 1.083 | 156 | 19.5 | 16.3 | 0.332 | 0.389 | 78.2 | 4.1 | 84 | 21.4 |
| 5xtd | d | 3.7 | 6773 | 0.0525 | 1.083 | 171 | 2.2 | 2.2 | 0.913 | 0.913 | 94.2 | 76.4 | 87.7 | 70.7 |
| 5xtd | g | 3.7 | 6773 | 0.0525 | 1.083 | 119 | 1.8 | 1.8 | 0.896 | 0.896 | 95.8 | 80.7 | 93.3 | 33.3 |
| 5xtd | p | 3.7 | 6773 | 0.0525 | 1.083 | 172 | 2.9 | 2.9 | 0.888 | 0.888 | 90.7 | 62.2 | 82.6 | 38 |
| 5xtd | u | 3.7 | 6773 | 0.0525 | 1.083 | 169 | 2.6 | 2.6 | 0.890 | 0.890 | 93.5 | 48.1 | 85.2 | 2.8 |
| 5xtd | V | 3.7 | 6773 | 0.0525 | 1.083 | 140 | 3.2 | 3.2 | 0.853 | 0.853 | 90.7 | 45.7 | 75 | 6.7 |
| 5xte | E | 3.4 | 6774 | 0.0783 | 1.083 | 74 | 5.6 | 4.6 | 0.790 | 0.790 | 90.5 | 23.9 | 78.4 | 75.9 |
| 5xte | F | 3.4 | 6774 | 0.0783 | 1.083 | 106 | 1.7 | 1.7 | 0.913 | 0.913 | 95.3 | 95 | 86.8 | 58.7 |
| 5xte | G | 3.4 | 6774 | 0.0783 | 1.083 | 51 | 1.7 | 1.7 | 0.866 | 0.866 | 96.1 | 77.6 | 82.4 | 78.6 |
| 5xte | H | 3.4 | 6774 | 0.0783 | 1.083 | 241 | 2.9 | 2.9 | 0.911 | 0.911 | 92.1 | 74.3 | 73 | 26.7 |
| 5xte | R | 3.4 | 6774 | 0.0783 | 1.083 | 74 | 3.2 | 3.2 | 0.807 | 0.807 | 89.2 | 57.6 | 78.4 | 82.8 |
| 5xte | S | 3.4 | 6774 | 0.0783 | 1.083 | 106 | 2.3 | 2.3 | 0.879 | 0.879 | 92.5 | 72.4 | 82.1 | 52.9 |
| 5xte | U | 3.4 | 6774 | 0.0783 | 1.083 | 241 | 2.8 | 2.8 | 0.932 | 0.932 | 95.4 | 80 | 77.6 | 39.6 |
| 5xxu | A | 3.4 | 6780 | 0.03 | 1.32 | 198 | 4.4 | 4.4 | 0.870 | 0.870 | 89.9 | 64 | 85.9 | 36.5 |
| 5xxu | B | 3.4 | 6780 | 0.03 | 1.32 | 212 | 7.5 | 7.5 | 0.791 | 0.791 | 87.3 | 52.4 | 76.4 | 30.9 |
| 5xxu | E | 3.4 | 6780 | 0.03 | 1.32 | 259 | 2.1 | 2.1 | 0.943 | 0.943 | 96.5 | 80.4 | 91.1 | 6.4 |
| 5xxu | H | 3.4 | 6780 | 0.03 | 1.32 | 177 | 5.3 | 4.3 | 0.860 | 0.860 | 88.7 | 28.7 | 80.8 | 25.9 |
| 5xyi | A | 3.4 | 6788 | 0.036 | 1.32 | 204 | 2.1 | 2.1 | 0.924 | 0.924 | 93.1 | 80 | 85.3 | 30.5 |
| 5xyi | B | 3.4 | 6788 | 0.036 | 1.32 | 219 | 4.9 | 4.9 | 0.888 | 0.888 | 90.9 | 63.3 | 79.9 | 6.3 |
| 5xyi | H | 3.4 | 6788 | 0.036 | 1.32 | 163 | 4.1 | 4.1 | 0.831 | 0.831 | 85.3 | 61.9 | 78.5 | 18.8 |
| 5xym | O | 3.1 | 6789 | 0.044 | 1.32 | 107 | 2.8 | 2.8 | 0.868 | 0.868 | 94.4 | 72.3 | 90.7 | 27.8 |
| 5y81 | C | 4.7 | 6816 | 0.0225 | 1.3 | 140 | 18.5 | 13.8 | 0.258 | 0.420 | 60.7 | 4.7 | 83.6 | 2.6 |
| 5y81 | H | 4.7 | 6816 | 0.0225 | 1.3 | 154 | 16.9 | 16.9 | 0.219 | 0.277 | 54.5 | 9.5 | 29.9 | 10.9 |
| 5y88 | e | 3.5 | 6817 | 0.05 | 1.306 | 82 | 10.8 | 10.8 | 0.682 | 0.682 | 78 | 17.2 | 63.4 | 3.8 |
| 5y88 | L | 3.5 | 6817 | 0.05 | 1.306 | 157 | 8.7 | 6.9 | 0.703 | 0.741 | 84.7 | 65.4 | 68.8 | 23.1 |
| 5y88 | M | 3.5 | 6817 | 0.05 | 1.306 | 182 | 22.8 | 19.7 | 0.632 | 0.632 | 64.3 | 29.1 | 37.4 | 4.4 |
| 5y88 | N | 3.5 | 6817 | 0.05 | 1.306 | 261 | 4.6 | 4.6 | 0.831 | 0.837 | 85.1 | 43.7 | 54.8 | 7 |
| 5ylz | L | 3.6 | 6839 | 0.034 | 1.06 | 157 | 3.5 | 3.5 | 0.859 | 0.859 | 88.5 | 72.7 | 76.4 | 25 |
| 5ylz | N | 3.6 | 6839 | 0.034 | 1.06 | 261 | 15.0 | 14.0 | 0.409 | 0.522 | 69.3 | 8.3 | 56.3 | 4.8 |
| 5ylz | S | 3.6 | 6839 | 0.034 | 1.06 | 207 | 3.6 | 3.6 | 0.882 | 0.882 | 91.8 | 71.6 | 78.3 | 66 |
| 5yq7 | C | 4.1 | 6828 | 0.03 | 1.12 | 293 | 24.6 | 22.9 | 0.309 | 0.763 | 72.7 | 6.1 | 31.7 | 4.3 |
| 5z3g | H | 3.7 | 6878 | 0.09 | 1.38 | 255 | 24.3 | 16.8 | 0.259 | 0.311 | 60.9 | 7.1 | 30.9 | 5.1 |
| 5z3g | J | 3.7 | 6878 | 0.09 | 1.38 | 239 | 2.7 | 2.7 | 0.896 | 0.896 | 90.4 | 69.9 | 78.2 | 20.3 |
| 5z3g | K | 3.7 | 6878 | 0.09 | 1.38 | 162 | 19.0 | 17.0 | 0.256 | 0.360 | 72.2 | 9.4 | 66.7 | 8.3 |
| 5z3g | N | 3.7 | 6878 | 0.09 | 1.38 | 226 | 15.5 | 13.3 | 0.306 | 0.345 | 53.5 | 6.6 | 11.5 | 3.8 |
| 5z3g | O | 3.7 | 6878 | 0.09 | 1.38 | 190 | 16.2 | 13.9 | 0.563 | 0.605 | 78.4 | 38.9 | 78.9 | 6.7 |
| 5z3g | Q | 3.7 | 6878 | 0.09 | 1.38 | 137 | 11.5 | 7.8 | 0.661 | 0.808 | 81 | 35.1 | 71.5 | 45.9 |

| | | | | | | | | | | | | | | |
|---|---|---|---|---|---|---|---|---|---|---|---|---|---|---|
| 5z3g | S | 3.7 | 6878 | 0.09 | 1.38 | 197 | 16.8 | 6.5 | 0.436 | 0.795 | 84.8 | 35.3 | 77.2 | 21.1 |
| 5z3g | W | 3.7 | 6878 | 0.09 | 1.38 | 170 | 16.4 | 10.4 | 0.663 | 0.689 | 77.6 | 14.4 | 71.2 | 3.3 |
| 5z3g | X | 3.7 | 6878 | 0.09 | 1.38 | 202 | 7.3 | 7.3 | 0.692 | 0.736 | 77.7 | 38.9 | 87.6 | 44.1 |
| 5z3o | D | 3.6 | 6880 | 0.04 | 1.32 | 93 | 0.9 | 0.9 | 0.944 | 0.944 | 100 | 100 | 98.9 | 91.3 |
| 5z3o | H | 3.6 | 6880 | 0.04 | 1.32 | 93 | 2.8 | 2.8 | 0.887 | 0.887 | 93.5 | 90.8 | 93.5 | 81.6 |
| 5z58 | z | 4.9 | 6891 | 0.0346 | 1.338 | 177 | 16.7 | 15.9 | 0.422 | 0.430 | 61 | 8.3 | 26 | 6.5 |
| 5z62 | B | 3.3 | 6896 | 0.032 | 1.091 | 227 | 6.7 | 6.7 | 0.832 | 0.845 | 86.3 | 52 | 81.1 | 5.4 |
| 5z62 | C | 3.3 | 6896 | 0.032 | 1.091 | 260 | 2.3 | 2.3 | 0.935 | 0.935 | 95.8 | 76.3 | 91.5 | 73.1 |
| 5z62 | E | 3.3 | 6896 | 0.032 | 1.091 | 109 | 2.8 | 2.8 | 0.872 | 0.872 | 95.4 | 50 | 79.8 | 27.6 |
| 5z62 | F | 3.3 | 6896 | 0.032 | 1.091 | 98 | 4.7 | 4.7 | 0.773 | 0.773 | 83.7 | 51.2 | 73.5 | 8.3 |
| 5z62 | H | 3.3 | 6896 | 0.032 | 1.091 | 82 | 2.4 | 2.4 | 0.838 | 0.838 | 92.7 | 64.5 | 87.8 | 36.1 |
| 5z62 | I | 3.3 | 6896 | 0.032 | 1.091 | 73 | 3.0 | 3.0 | 0.835 | 0.835 | 91.8 | 73.1 | 83.6 | 6.6 |
| 5zdh | b | 3.2 | 6917 | 0.025 | 1.32 | 84 | 10.7 | 9.4 | 0.496 | 0.496 | 76.2 | 26.6 | 71.4 | 3.3 |
| 5zdh | X | 3.2 | 6917 | 0.025 | 1.32 | 84 | 12.4 | 9.5 | 0.423 | 0.448 | 70.2 | 5.1 | 61.9 | 7.7 |
| 5zgb | D | 3.6 | 6929 | 0.0562 | 1.05 | 119 | 4.5 | 3.9 | 0.855 | 0.855 | 92.4 | 58.2 | 52.9 | 6.3 |
| 5zgb | E | 3.6 | 6929 | 0.0562 | 1.05 | 61 | 3.3 | 3.3 | 0.783 | 0.783 | 88.5 | 42.6 | 37.7 | 0 |
| 5zji | 2 | 3.3 | 6932 | 0.05 | 1.06 | 207 | 2.4 | 2.4 | 0.920 | 0.920 | 93.2 | 73.1 | 81.6 | 47.9 |
| 5zji | 3 | 3.3 | 6932 | 0.05 | 1.06 | 221 | 2.5 | 2.5 | 0.908 | 0.908 | 93.2 | 67.5 | 80.5 | 44.4 |
| 5zji | 4 | 3.3 | 6932 | 0.05 | 1.06 | 199 | 2.2 | 2.2 | 0.928 | 0.928 | 95.5 | 74.7 | 80.4 | 56.9 |
| 5zji | C | 3.3 | 6932 | 0.05 | 1.06 | 81 | 2.8 | 2.8 | 0.856 | 0.856 | 91.4 | 73 | 56.8 | 0 |
| 5zji | D | 3.3 | 6932 | 0.05 | 1.06 | 142 | 1.5 | 1.5 | 0.933 | 0.933 | 96.5 | 91.2 | 89.4 | 15.7 |
| 5zji | E | 3.3 | 6932 | 0.05 | 1.06 | 68 | 2.0 | 2.0 | 0.872 | 0.872 | 95.6 | 80 | 73.5 | 18 |
| 5zji | F | 3.3 | 6932 | 0.05 | 1.06 | 158 | 2.3 | 2.3 | 0.905 | 0.905 | 94.3 | 77.2 | 88.6 | 54.3 |
| 5zwn | c | 3.4 | 6973 | 0.045 | 1.338 | 101 | 2.6 | 2.6 | 0.877 | 0.877 | 93.1 | 85.1 | 78.2 | 8.9 |
| 5zwn | e | 3.4 | 6973 | 0.045 | 1.338 | 74 | 3.6 | 3.6 | 0.816 | 0.816 | 89.2 | 68.2 | 90.5 | 10.4 |
| 5zwn | f | 3.4 | 6973 | 0.045 | 1.338 | 68 | 2.3 | 2.3 | 0.846 | 0.846 | 91.2 | 75.8 | 86.8 | 35.6 |
| 5zwn | V | 3.4 | 6973 | 0.045 | 1.338 | 187 | 6.5 | 3.5 | 0.851 | 0.885 | 90.9 | 60.6 | 85 | 34.6 |
| 5zwn | W | 3.4 | 6973 | 0.045 | 1.338 | 229 | 5.2 | 4.2 | 0.847 | 0.895 | 90 | 67.5 | 83.4 | 46.1 |
| 5zwo | E | 3.9 | 6974 | 0.022 | 1.338 | 138 | 3.7 | 3.7 | 0.842 | 0.842 | 89.9 | 52.4 | 62.3 | 39.5 |
| 5zwo | M | 3.9 | 6974 | 0.022 | 1.338 | 126 | 2.6 | 2.6 | 0.878 | 0.878 | 92.1 | 75.9 | 86.5 | 42.2 |
| 6a91 | B | 3.2 | 6996 | 0.08 | 1.091 | 57 | 3.0 | 3.0 | 0.831 | 0.831 | 94.7 | 79.6 | 78.9 | 31.1 |
| 6a96 | K | 3.5 | 6998 | 7 | 1.07 | 125 | 14.9 | 13.1 | 0.340 | 0.723 | 75.2 | 6.4 | 70.4 | 11.4 |
| 6a96 | L | 3.5 | 6998 | 7 | 1.07 | 124 | 15.2 | 15.2 | 0.285 | 0.304 | 66.1 | 12.2 | 52.4 | 3.1 |
| 6a96 | O | 3.5 | 6998 | 7 | 1.07 | 125 | 15.6 | 14.4 | 0.319 | 0.319 | 65.6 | 4.9 | 37.6 | 6.4 |
| 6agb | J | 3.5 | 9616 | 0.03 | 1.32 | 293 | 6.3 | 3.9 | 0.913 | 0.913 | 93.2 | 81 | 79.9 | 32.5 |
| 6agf | B | 3.2 | 9617 | 0.05 | 1.091 | 173 | 1.7 | 1.7 | 0.938 | 0.938 | 96.5 | 88.6 | 86.1 | 42.3 |
| 6ahu | I | 3.7 | 9627 | 0.0256 | 1.32 | 236 | 12.2 | 9.4 | 0.668 | 0.670 | 80.1 | 36.5 | 59.3 | 4.3 |

| | | | | | | | | | | | | | | |
|---|---|---|---|---|---|---|---|---|---|---|---|---|---|---|
| 6ahu | J | 3.7 | 9627 | 0.0256 | 1.32 | 247 | 4.9 | 4.9 | 0.840 | 0.840 | 84.6 | 47.8 | 78.9 | 13.8 |
| 6asx | G | 3.8 | 7002 | 0.01 | 1.03 | 220 | 9.8 | 4.3 | 0.882 | 0.894 | 90.5 | 64.3 | 88.6 | 16.4 |
| 6asx | H | 3.8 | 7002 | 0.01 | 1.03 | 220 | 11.5 | 10.0 | 0.611 | 0.643 | 85 | 45.5 | 76.4 | 22 |
| 6avo | b | 3.8 | 7010 | 0.0383 | 1.2 | 230 | 8.4 | 6.0 | 0.773 | 0.813 | 83 | 48.7 | 62.2 | 5.6 |
| 6avo | G | 3.8 | 7010 | 0.0383 | 1.2 | 234 | 4.2 | 4.2 | 0.855 | 0.855 | 87.2 | 58.3 | 80.3 | 4.8 |
| 6avo | I | 3.8 | 7010 | 0.0383 | 1.2 | 232 | 16.6 | 13.3 | 0.312 | 0.651 | 76.3 | 8.5 | 68.1 | 25.9 |
| 6avo | K | 3.8 | 7010 | 0.0383 | 1.2 | 240 | 19.6 | 15.8 | 0.413 | 0.425 | 83.3 | 24.5 | 73.3 | 5.1 |
| 6avo | L | 3.8 | 7010 | 0.0383 | 1.2 | 234 | 4.6 | 4.6 | 0.848 | 0.848 | 84.6 | 52 | 70.9 | 8.4 |
| 6avo | N | 3.8 | 7010 | 0.0383 | 1.2 | 232 | 11.9 | 8.9 | 0.639 | 0.743 | 74.6 | 32.4 | 55.2 | 35.2 |
| 6avo | P | 3.8 | 7010 | 0.0383 | 1.2 | 230 | 6.6 | 5.5 | 0.817 | 0.823 | 83.5 | 47.4 | 80.9 | 34.9 |
| 6avo | R | 3.8 | 7010 | 0.0383 | 1.2 | 240 | 10.1 | 8.7 | 0.752 | 0.761 | 79.6 | 55 | 75.4 | 30.9 |
| 6avo | T | 3.8 | 7010 | 0.0383 | 1.2 | 196 | 12.7 | 12.6 | 0.509 | 0.516 | 83.2 | 34.4 | 82.7 | 29.6 |
| 6avo | V | 3.8 | 7010 | 0.0383 | 1.2 | 196 | 6.4 | 6.0 | 0.826 | 0.826 | 84.2 | 46.7 | 82.7 | 35.8 |
| 6az1 | a | 2.7 | 7024 | 0.05 | 1.02 | 72 | 5.1 | 5.1 | 0.752 | 0.752 | 90.3 | 61.5 | 83.3 | 86.7 |
| 6az1 | B | 2.7 | 7024 | 0.05 | 1.02 | 211 | 1.3 | 1.3 | 0.961 | 0.961 | 97.6 | 96.1 | 90 | 68.4 |
| 6az1 | E | 2.7 | 7024 | 0.05 | 1.02 | 260 | 3.8 | 3.8 | 0.934 | 0.934 | 95 | 66.4 | 95.8 | 8.8 |
| 6b3j | N | 3.3 | 7039 | 0.04 | 1.06 | 126 | 11.8 | 8.4 | 0.458 | 0.780 | 81 | 14.7 | 76.2 | 7.3 |
| 6b46 | C | 3.1 | 7050 | 3.71 | 0.8254 | 293 | 8.7 | 7.5 | 0.809 | 0.809 | 81.9 | 33.8 | 64.2 | 35.6 |
| 6b46 | I | 3.1 | 7050 | 3.71 | 0.8254 | 79 | 2.3 | 2.3 | 0.855 | 0.855 | 92.4 | 72.6 | 74.7 | 35.6 |
| 6b47 | C | 3.2 | 7051 | 0.07 | 1.0724 | 293 | 22.1 | 22.1 | 0.484 | 0.502 | 78.5 | 23.5 | 65.5 | 24.5 |
| 6b6h | A | 3.9 | 7059 | 0.13 | 1.35 | 230 | 25.8 | 24.5 | 0.398 | 0.398 | 73.5 | 5.3 | 30.4 | 8.6 |
| 6b6h | B | 3.9 | 7059 | 0.13 | 1.35 | 228 | 26.9 | 25.9 | 0.265 | 0.317 | 63.2 | 4.9 | 36.8 | 7.1 |
| 6b70 | C | 3.7 | 7062 | 0.024 | 1.073 | 218 | 18.0 | 17.1 | 0.391 | 0.444 | 70.6 | 3.9 | 54.6 | 10.1 |
| 6b70 | D | 3.7 | 7062 | 0.024 | 1.073 | 211 | 19.1 | 17.7 | 0.341 | 0.426 | 72 | 12.5 | 28 | 8.5 |
| 6bk8 | F | 3.3 | 7109 | 0.06 | 1.36 | 199 | 20.7 | 19.3 | 0.690 | 0.690 | 67.8 | 36.3 | 63.8 | 4.7 |
| 6bk8 | G | 3.3 | 7109 | 0.06 | 1.36 | 255 | 6.5 | 6.5 | 0.789 | 0.848 | 91.8 | 71.4 | 80 | 21.6 |
| 6bk8 | I | 3.3 | 7109 | 0.06 | 1.36 | 156 | 1.8 | 1.8 | 0.946 | 0.946 | 98.1 | 89.5 | 84.6 | 40.9 |
| 6bk8 | N | 3.3 | 7109 | 0.06 | 1.36 | 167 | 2.3 | 2.3 | 0.915 | 0.915 | 94 | 71.3 | 92.8 | 38.1 |
| 6bk8 | S | 3.3 | 7109 | 0.06 | 1.36 | 238 | 5.8 | 5.8 | 0.850 | 0.850 | 90.8 | 81 | 89.9 | 28.5 |
| 6btm | A | 3.4 | 7286 | 0.39 | 1.1 | 221 | 2.5 | 2.5 | 0.912 | 0.912 | 94.1 | 68.3 | 71.9 | 11.9 |
| 6btm | D | 3.4 | 7286 | 0.39 | 1.1 | 172 | 2.7 | 2.7 | 0.888 | 0.888 | 91.9 | 59.5 | 81.4 | 25 |
| 6btm | E | 3.4 | 7286 | 0.39 | 1.1 | 161 | 2.6 | 2.6 | 0.913 | 0.913 | 95 | 73.9 | 86.3 | 44.6 |
| 6buz | N | 3.9 | 7293 | 0.0014 | 0.42 | 134 | 14.5 | 12.0 | 0.287 | 0.317 | 63.4 | 4.7 | 78.4 | 6.7 |
| 6bzo | A | 3.4 | 7319 | 0.347 | 1.1 | 226 | 2.8 | 2.8 | 0.928 | 0.928 | 93.4 | 83.4 | 88.5 | 58.5 |
| 6bzo | B | 3.4 | 7319 | 0.347 | 1.1 | 237 | 4.3 | 4.3 | 0.901 | 0.901 | 92 | 48.6 | 75.9 | 7.2 |
| 6bzo | E | 3.4 | 7319 | 0.347 | 1.1 | 83 | 2.0 | 2.0 | 0.857 | 0.857 | 91.6 | 77.6 | 88 | 72.6 |
| 6bzo | J | 3.4 | 7319 | 0.347 | 1.1 | 107 | 2.6 | 2.6 | 0.861 | 0.861 | 93.5 | 66 | 69.2 | 10.8 |

| | | | | | | | | | | | | | | |
|---|---|---|---|---|---|---|---|---|---|---|---|---|---|---|
| 6c04 | A | 3.3 | 7320 | 0.372 | 1.1 | 226 | 1.8 | 1.8 | 0.950 | 0.950 | 96 | 88.9 | 88.5 | 60.5 |
| 6c04 | B | 3.3 | 7320 | 0.372 | 1.1 | 237 | 2.4 | 2.4 | 0.922 | 0.922 | 93.7 | 68 | 87.3 | 40.6 |
| 6c04 | E | 3.3 | 7320 | 0.372 | 1.1 | 83 | 3.4 | 3.4 | 0.871 | 0.871 | 94 | 73.1 | 68.7 | 50.9 |
| 6c04 | J | 3.3 | 7320 | 0.372 | 1.1 | 108 | 5.9 | 2.8 | 0.826 | 0.854 | 88.9 | 59.4 | 94.4 | 19.6 |
| 6c0f | D | 3.7 | 7324 | 0.02 | 1.3 | 190 | 3.9 | 3.9 | 0.877 | 0.877 | 92.6 | 67 | 77.9 | 37.8 |
| 6c0f | F | 3.7 | 7324 | 0.02 | 1.3 | 242 | 13.0 | 10.6 | 0.573 | 0.826 | 83.5 | 28.2 | 73.1 | 5.6 |
| 6c0f | I | 3.7 | 7324 | 0.02 | 1.3 | 288 | 3.9 | 3.9 | 0.913 | 0.913 | 92.7 | 39 | 80.9 | 9 |
| 6c0f | K | 3.7 | 7324 | 0.02 | 1.3 | 260 | 18.4 | 15.4 | 0.324 | 0.542 | 79.2 | 13.1 | 66.2 | 8.7 |
| 6c0f | O | 3.7 | 7324 | 0.02 | 1.3 | 184 | 14.1 | 13.3 | 0.539 | 0.835 | 85.9 | 29.1 | 75.5 | 10.1 |
| 6c0f | Q | 3.7 | 7324 | 0.02 | 1.3 | 132 | 2.2 | 2.2 | 0.896 | 0.896 | 96.2 | 79.5 | 78 | 27.2 |
| 6c0f | t | 3.7 | 7324 | 0.02 | 1.3 | 248 | 24.2 | 18.2 | 0.372 | 0.408 | 78.6 | 6.2 | 62.1 | 9.7 |
| 6c0f | y | 3.7 | 7324 | 0.02 | 1.3 | 226 | 14.8 | 14.2 | 0.320 | 0.351 | 56.2 | 7.1 | 23 | 9.6 |
| 6c0f | z | 3.7 | 7324 | 0.02 | 1.3 | 243 | 10.4 | 4.4 | 0.722 | 0.864 | 90.5 | 52.3 | 81.1 | 25.4 |
| 6c0w | K | 4 | 7326 | 0.02 | 1.02 | 205 | 7.7 | 7.7 | 0.733 | 0.733 | 78.5 | 29.2 | 74.6 | 6.5 |
| 6c26 | C | 3.5 | 7336 | 0.04 | 1.088 | 248 | 5.9 | 5.9 | 0.817 | 0.832 | 90.7 | 60 | 81.9 | 50.2 |
| 6c5v | C | 4.8 | 7344 | 0.08 | 1.36 | 177 | 14.8 | 14.0 | 0.411 | 0.433 | 66.7 | 5.9 | 28.8 | 2 |
| 6c5v | H | 4.8 | 7344 | 0.08 | 1.36 | 214 | 16.2 | 16.2 | 0.287 | 0.313 | 65 | 6.5 | 41.1 | 8 |
| 6c5v | L | 4.8 | 7344 | 0.08 | 1.36 | 212 | 19.7 | 17.6 | 0.418 | 0.418 | 64.2 | 12.5 | 30.2 | 12.5 |
| 6c66 | I | 3.7 | 7347 | 0.03 | 1.24 | 177 | 10.0 | 10.0 | 0.665 | 0.665 | 89.8 | 65.4 | 88.1 | 50 |
| 6c66 | K | 3.7 | 7347 | 0.03 | 1.24 | 165 | 11.7 | 11.7 | 0.447 | 0.550 | 86.7 | 36.4 | 87.9 | 69.7 |
| 6c66 | M | 3.7 | 7347 | 0.03 | 1.24 | 244 | 3.2 | 3.2 | 0.886 | 0.886 | 91 | 60.4 | 69.3 | 5.3 |
| 6c6s | D | 3.7 | 7349 | 0.02 | 1.3 | 161 | 20.8 | 8.1 | 0.244 | 0.668 | 72.7 | 18.8 | 46 | 6.8 |
| 6c6s | G | 3.7 | 7349 | 0.02 | 1.3 | 221 | 4.7 | 4.5 | 0.908 | 0.908 | 93.2 | 65.5 | 85.1 | 6.4 |
| 6c6s | H | 3.7 | 7349 | 0.02 | 1.3 | 219 | 17.7 | 13.1 | 0.573 | 0.580 | 77.6 | 24.7 | 73.5 | 7.5 |
| 6c6s | K | 3.7 | 7349 | 0.02 | 1.3 | 83 | 2.2 | 2.2 | 0.859 | 0.859 | 94 | 71.8 | 80.7 | 7.5 |
| 6c6t | D | 3.5 | 7350 | 0.016 | 1.3 | 85 | 3.9 | 3.9 | 0.767 | 0.783 | 82.4 | 65.7 | 70.6 | 5 |
| 6c6t | G | 3.5 | 7350 | 0.016 | 1.3 | 221 | 2.9 | 2.9 | 0.929 | 0.929 | 94.1 | 80.8 | 90 | 32.7 |
| 6c6t | H | 3.5 | 7350 | 0.016 | 1.3 | 219 | 15.0 | 11.0 | 0.580 | 0.693 | 81.3 | 44.4 | 79.5 | 15.5 |
| 6c6u | G | 3.7 | 7351 | 0.013 | 1.07 | 221 | 3.8 | 3.8 | 0.878 | 0.890 | 92.3 | 61.3 | 89.6 | 52.5 |
| 6c6u | H | 3.7 | 7351 | 0.013 | 1.07 | 218 | 12.6 | 11.6 | 0.702 | 0.702 | 81.7 | 32 | 73.4 | 9.4 |
| 6c6u | N | 3.7 | 7351 | 0.013 | 1.07 | 98 | 3.4 | 3.4 | 0.822 | 0.822 | 93.9 | 62 | 46.9 | 2.2 |
| 6cb1 | F | 4.6 | 7445 | 0.018 | 1.3 | 242 | 22.0 | 21.0 | 0.393 | 0.549 | 67.4 | 6.7 | 51.7 | 5.6 |
| 6cb1 | z | 4.6 | 7445 | 0.018 | 1.3 | 243 | 11.3 | 7.3 | 0.554 | 0.757 | 78.6 | 7.3 | 54.3 | 8.3 |
| 6cde | 1 | 3.8 | 7459 | 0.544 | 1.06 | 132 | 4.8 | 4.8 | 0.807 | 0.807 | 83.3 | 60 | 78 | 4.9 |
| 6cde | 3 | 3.8 | 7459 | 0.544 | 1.06 | 113 | 14.4 | 14.0 | 0.333 | 0.355 | 71.7 | 9.9 | 23 | 3.8 |
| 6cde | 5 | 3.8 | 7459 | 0.544 | 1.06 | 128 | 1.6 | 1.6 | 0.923 | 0.923 | 96.9 | 86.3 | 89.8 | 6.1 |
| 6cde | 6 | 3.8 | 7459 | 0.544 | 1.06 | 103 | 5.9 | 2.8 | 0.870 | 0.883 | 92.2 | 63.2 | 55.3 | 10.5 |

| | | | | | | | | | | | | | | |
|---|---|---|---|---|---|---|---|---|---|---|---|---|---|---|
| 6cde | 7 | 3.8 | 7459 | 0.544 | 1.06 | 132 | 17.3 | 16.8 | 0.320 | 0.406 | 74.2 | 4.1 | 31.8 | 9.5 |
| 6cde | 8 | 3.8 | 7459 | 0.544 | 1.06 | 109 | 14.5 | 13.7 | 0.255 | 0.419 | 71.6 | 10.3 | 43.1 | 2.1 |
| 6cde | d | 3.8 | 7459 | 0.544 | 1.06 | 132 | 6.0 | 5.1 | 0.781 | 0.800 | 82.6 | 33.9 | 76.5 | 51.5 |
| 6cde | D | 3.8 | 7459 | 0.544 | 1.06 | 132 | 4.2 | 4.2 | 0.815 | 0.818 | 85.6 | 44.2 | 72 | 50.5 |
| 6cde | h | 3.8 | 7459 | 0.544 | 1.06 | 113 | 15.5 | 13.8 | 0.303 | 0.378 | 61.1 | 4.3 | 27.4 | 6.5 |
| 6cde | H | 3.8 | 7459 | 0.544 | 1.06 | 113 | 14.5 | 14.1 | 0.307 | 0.351 | 63.7 | 12.5 | 26.5 | 3.3 |
| 6cde | m | 3.8 | 7459 | 0.544 | 1.06 | 132 | 18.0 | 16.4 | 0.347 | 0.416 | 71.2 | 6.4 | 23.5 | 0 |
| 6cde | M | 3.8 | 7459 | 0.544 | 1.06 | 132 | 15.0 | 15.0 | 0.355 | 0.452 | 78 | 23.3 | 33.3 | 6.8 |
| 6cde | n | 3.8 | 7459 | 0.544 | 1.06 | 109 | 15.7 | 14.4 | 0.490 | 0.490 | 72.5 | 7.6 | 41.3 | 6.7 |
| 6cde | N | 3.8 | 7459 | 0.544 | 1.06 | 109 | 14.1 | 13.1 | 0.445 | 0.468 | 73.4 | 25 | 62.4 | 4.4 |
| 6cde | q | 3.8 | 7459 | 0.544 | 1.06 | 128 | 1.8 | 1.8 | 0.933 | 0.933 | 96.9 | 83.1 | 54.7 | 1.4 |
| 6cde | Q | 3.8 | 7459 | 0.544 | 1.06 | 128 | 2.5 | 2.5 | 0.916 | 0.916 | 96.1 | 82.1 | 89.1 | 7 |
| 6cde | r | 3.8 | 7459 | 0.544 | 1.06 | 103 | 2.7 | 2.7 | 0.916 | 0.916 | 97.1 | 80 | 71.8 | 24.3 |
| 6cde | R | 3.8 | 7459 | 0.544 | 1.06 | 103 | 2.1 | 2.1 | 0.877 | 0.877 | 93.2 | 67.7 | 81.6 | 7.1 |
| 6cdi | 3 | 3.6 | 7460 | 1.01 | 1.07 | 116 | 14.0 | 13.2 | 0.292 | 0.342 | 70.7 | 12.2 | 40.5 | 6.4 |
| 6cdi | 4 | 3.6 | 7460 | 1.01 | 1.07 | 113 | 14.8 | 14.8 | 0.270 | 0.343 | 68.1 | 10.4 | 29.2 | 9.1 |
| 6cdi | 5 | 3.6 | 7460 | 1.01 | 1.07 | 132 | 19.6 | 16.5 | 0.483 | 0.488 | 75.8 | 6 | 35.6 | 4.3 |
| 6cdi | 6 | 3.6 | 7460 | 1.01 | 1.07 | 105 | 15.7 | 12.0 | 0.300 | 0.432 | 73.3 | 10.4 | 61.9 | 4.6 |
| 6cdi | 7 | 3.6 | 7460 | 1.01 | 1.07 | 102 | 10.8 | 8.7 | 0.694 | 0.737 | 79.4 | 61.7 | 82.4 | 9.5 |
| 6cdi | 8 | 3.6 | 7460 | 1.01 | 1.07 | 128 | 8.4 | 2.3 | 0.858 | 0.895 | 89.1 | 63.2 | 84.4 | 5.6 |
| 6cdi | A | 3.6 | 7460 | 1.01 | 1.07 | 132 | 6.9 | 6.5 | 0.801 | 0.812 | 84.1 | 48.6 | 78 | 41.7 |
| 6cdi | c | 3.6 | 7460 | 1.01 | 1.07 | 132 | 6.6 | 6.6 | 0.804 | 0.804 | 84.8 | 59.8 | 82.6 | 31.2 |
| 6cdi | D | 3.6 | 7460 | 1.01 | 1.07 | 132 | 7.0 | 5.8 | 0.756 | 0.768 | 78.8 | 29.8 | 85.6 | 38.1 |
| 6cdi | h | 3.6 | 7460 | 1.01 | 1.07 | 116 | 12.5 | 12.5 | 0.371 | 0.427 | 68.1 | 10.1 | 53.4 | 4.8 |
| 6cdi | H | 3.6 | 7460 | 1.01 | 1.07 | 116 | 15.1 | 13.5 | 0.363 | 0.369 | 74.1 | 10.5 | 55.2 | 6.2 |
| 6cdi | l | 3.6 | 7460 | 1.01 | 1.07 | 113 | 15.4 | 14.8 | 0.304 | 0.304 | 64.6 | 4.1 | 23.9 | 11.1 |
| 6cdi | L | 3.6 | 7460 | 1.01 | 1.07 | 113 | 16.1 | 14.2 | 0.279 | 0.309 | 65.5 | 12.2 | 21.2 | 4.2 |
| 6cdi | m | 3.6 | 7460 | 1.01 | 1.07 | 132 | 16.4 | 15.6 | 0.445 | 0.465 | 68.9 | 15.4 | 39.4 | 5.8 |
| 6cdi | M | 3.6 | 7460 | 1.01 | 1.07 | 132 | 15.2 | 14.2 | 0.519 | 0.543 | 79.5 | 17.1 | 37.9 | 10 |
| 6cdi | n | 3.6 | 7460 | 1.01 | 1.07 | 105 | 14.4 | 13.3 | 0.314 | 0.402 | 77.1 | 6.2 | 51.4 | 5.6 |
| 6cdi | N | 3.6 | 7460 | 1.01 | 1.07 | 105 | 14.7 | 12.6 | 0.393 | 0.448 | 70.5 | 18.9 | 67.6 | 21.1 |
| 6cdi | q | 3.6 | 7460 | 1.01 | 1.07 | 128 | 1.7 | 1.7 | 0.924 | 0.924 | 96.9 | 80.6 | 69.5 | 5.6 |
| 6cdi | Q | 3.6 | 7460 | 1.01 | 1.07 | 128 | 1.8 | 1.8 | 0.909 | 0.909 | 95.3 | 82 | 86.7 | 7.2 |
| 6cdi | r | 3.6 | 7460 | 1.01 | 1.07 | 102 | 2.3 | 2.3 | 0.870 | 0.870 | 95.1 | 70.1 | 52 | 11.3 |
| 6cdi | R | 3.6 | 7460 | 1.01 | 1.07 | 102 | 2.2 | 2.2 | 0.887 | 0.887 | 94.1 | 84.4 | 88.2 | 6.7 |
| 6cfw | A | 3.7 | 7468 | 0.048 | 1.088 | 165 | 4.5 | 3.6 | 0.873 | 0.887 | 88.5 | 51.4 | 85.5 | 48.2 |
| 6cfw | C | 3.7 | 7468 | 0.048 | 1.088 | 114 | 1.7 | 1.7 | 0.901 | 0.901 | 95.6 | 86.2 | 93 | 56.6 |

| | | | | | | | | | | | | | | |
|---|---|---|---|---|---|---|---|---|---|---|---|---|---|---|
| 6cfw | F | 3.7 | 7468 | 0.048 | 1.088 | 145 | 2.4 | 2.4 | 0.910 | 0.910 | 94.5 | 65.7 | 84.1 | 27.9 |
| 6cfw | J | 3.7 | 7468 | 0.048 | 1.088 | 143 | 2.3 | 2.3 | 0.909 | 0.909 | 95.1 | 83.1 | 90.9 | 18.5 |
| 6cfw | K | 3.7 | 7468 | 0.048 | 1.088 | 166 | 7.3 | 4.4 | 0.842 | 0.858 | 85.5 | 42.3 | 72.9 | 9.9 |
| 6cfw | N | 3.7 | 7468 | 0.048 | 1.088 | 121 | 2.1 | 2.1 | 0.901 | 0.901 | 95 | 75.7 | 71.1 | 11.6 |
| 6cm3 | B | 3.5 | 7516 | 0.0386 | 1.31 | 118 | 10.3 | 10.0 | 0.594 | 0.740 | 82.2 | 46.4 | 81.4 | 46.9 |
| 6cm3 | C | 3.5 | 7516 | 0.0386 | 1.31 | 118 | 19.8 | 8.9 | 0.657 | 0.719 | 83.1 | 33.7 | 89 | 37.1 |
| 6cm3 | G | 3.5 | 7516 | 0.0386 | 1.31 | 97 | 2.4 | 2.4 | 0.890 | 0.890 | 95.9 | 75.3 | 52.6 | 9.8 |
| 6cm3 | H | 3.5 | 7516 | 0.0386 | 1.31 | 97 | 7.2 | 7.2 | 0.782 | 0.782 | 83.5 | 69.1 | 67 | 9.2 |
| 6cm3 | I | 3.5 | 7516 | 0.0386 | 1.31 | 97 | 1.8 | 1.8 | 0.883 | 0.883 | 94.8 | 84.8 | 77.3 | 6.7 |
| 6cm3 | J | 3.5 | 7516 | 0.0386 | 1.31 | 107 | 15.5 | 13.8 | 0.256 | 0.326 | 59.8 | 12.5 | 28 | 10 |
| 6cm3 | K | 3.5 | 7516 | 0.0386 | 1.31 | 127 | 17.1 | 15.0 | 0.305 | 0.442 | 77.2 | 7.1 | 48 | 3.3 |
| 6cm3 | L | 3.5 | 7516 | 0.0386 | 1.31 | 107 | 13.9 | 12.6 | 0.300 | 0.410 | 73.8 | 15.2 | 24.3 | 7.7 |
| 6cm3 | M | 3.5 | 7516 | 0.0386 | 1.31 | 127 | 16.0 | 15.6 | 0.359 | 0.457 | 74 | 8.5 | 54.3 | 8.7 |
| 6cm3 | N | 3.5 | 7516 | 0.0386 | 1.31 | 107 | 16.1 | 14.9 | 0.387 | 0.387 | 72 | 7.8 | 33.6 | 13.9 |
| 6cm3 | O | 3.5 | 7516 | 0.0386 | 1.31 | 127 | 16.6 | 13.6 | 0.386 | 0.479 | 78.7 | 5 | 63.8 | 8.6 |
| 6cm3 | P | 3.5 | 7516 | 0.0386 | 1.31 | 131 | 10.6 | 8.4 | 0.823 | 0.823 | 87 | 41.2 | 74 | 8.2 |
| 6cm3 | Q | 3.5 | 7516 | 0.0386 | 1.31 | 107 | 8.5 | 6.8 | 0.794 | 0.794 | 86.9 | 51.6 | 85 | 6.6 |
| 6cm3 | R | 3.5 | 7516 | 0.0386 | 1.31 | 131 | 12.2 | 8.1 | 0.720 | 0.819 | 84 | 59.1 | 71.8 | 8.5 |
| 6cm3 | S | 3.5 | 7516 | 0.0386 | 1.31 | 107 | 5.7 | 5.7 | 0.778 | 0.778 | 85 | 44 | 69.2 | 8.1 |
| 6cm3 | T | 3.5 | 7516 | 0.0386 | 1.31 | 131 | 2.6 | 2.6 | 0.902 | 0.902 | 95.4 | 66.4 | 71 | 8.6 |
| 6cm3 | U | 3.5 | 7516 | 0.0386 | 1.31 | 107 | 5.0 | 5.0 | 0.823 | 0.823 | 86.9 | 53.8 | 73.8 | 6.3 |
| 6cm9 | H | 3.7 | 7457 | 0.0726 | 1.067 | 163 | 4.3 | 4.3 | 0.798 | 0.798 | 80.4 | 61.1 | 79.8 | 10 |
| 6cnb | E | 4.1 | 7530 | 0.0334 | 1.18 | 215 | 6.8 | 6.8 | 0.776 | 0.781 | 84.7 | 59.9 | 57.2 | 7.3 |
| 6cnb | F | 4.1 | 7530 | 0.0334 | 1.18 | 83 | 2.0 | 2.0 | 0.873 | 0.873 | 97.6 | 74.1 | 86.7 | 5.6 |
| 6cnb | G | 4.1 | 7530 | 0.0334 | 1.18 | 184 | 21.3 | 17.7 | 0.237 | 0.288 | 64.7 | 6.7 | 22.8 | 11.9 |
| 6cnb | H | 4.1 | 7530 | 0.0334 | 1.18 | 140 | 17.2 | 14.4 | 0.485 | 0.563 | 85 | 26.1 | 45.7 | 6.2 |
| 6cnb | K | 4.1 | 7530 | 0.0334 | 1.18 | 101 | 2.5 | 2.5 | 0.882 | 0.882 | 94.1 | 83.2 | 89.1 | 35.6 |
| 6cnb | P | 4.1 | 7530 | 0.0334 | 1.18 | 277 | 11.1 | 9.2 | 0.559 | 0.736 | 74.4 | 10.2 | 19.5 | 7.4 |
| 6cnc | E | 4.1 | 7531 | 0.04 | 1.18 | 215 | 7.6 | 6.4 | 0.750 | 0.753 | 80.9 | 48.3 | 65.6 | 3.5 |
| 6cnc | F | 4.1 | 7531 | 0.04 | 1.18 | 83 | 4.0 | 4.0 | 0.796 | 0.798 | 88 | 61.6 | 92.8 | 37.7 |
| 6cnc | G | 4.1 | 7531 | 0.04 | 1.18 | 184 | 20.5 | 17.2 | 0.321 | 0.340 | 69 | 5.5 | 15.2 | 17.9 |
| 6cnc | H | 4.1 | 7531 | 0.04 | 1.18 | 140 | 17.3 | 14.8 | 0.349 | 0.538 | 77.9 | 9.2 | 57.1 | 13.8 |
| 6cnc | K | 4.1 | 7531 | 0.04 | 1.18 | 101 | 2.8 | 2.8 | 0.886 | 0.886 | 95 | 68.8 | 88.1 | 6.7 |
| 6cnc | M | 4.1 | 7531 | 0.04 | 1.18 | 164 | 20.7 | 17.5 | 0.352 | 0.352 | 65.9 | 12 | 30.5 | 8 |
| 6cnc | P | 4.1 | 7531 | 0.04 | 1.18 | 277 | 11.5 | 11.1 | 0.644 | 0.656 | 71.1 | 12.2 | 29.2 | 11.1 |
| 6cnd | E | 4.8 | 7532 | 0.035 | 1.18 | 215 | 15.9 | 14.4 | 0.627 | 0.647 | 75.8 | 18.4 | 47 | 3 |
| 6cnd | F | 4.8 | 7532 | 0.035 | 1.18 | 83 | 2.4 | 2.4 | 0.799 | 0.799 | 90.4 | 73.3 | 44.6 | 0 |

| | | | | | | | | | | | | | | |
|---|---|---|---|---|---|---|---|---|---|---|---|---|---|---|
| 6cnd | G | 4.8 | 7532 | 0.035 | 1.18 | 184 | 21.4 | 17.6 | 0.285 | 0.286 | 59.2 | 6.4 | 18.5 | 8.8 |
| 6cnd | H | 4.8 | 7532 | 0.035 | 1.18 | 140 | 17.1 | 15.9 | 0.231 | 0.329 | 64.3 | 7.8 | 55.7 | 3.8 |
| 6cnf | E | 4.5 | 7533 | 0.035 | 1.18 | 215 | 11.3 | 9.7 | 0.586 | 0.618 | 72.1 | 32.3 | 58.6 | 3.2 |
| 6cnf | F | 4.5 | 7533 | 0.035 | 1.18 | 83 | 3.0 | 3.0 | 0.778 | 0.778 | 86.7 | 62.5 | 43.4 | 8.3 |
| 6cnf | G | 4.5 | 7533 | 0.035 | 1.18 | 184 | 16.3 | 15.2 | 0.288 | 0.290 | 61.4 | 8.8 | 20.1 | 13.5 |
| 6cnf | H | 4.5 | 7533 | 0.035 | 1.18 | 140 | 14.5 | 14.5 | 0.252 | 0.312 | 62.1 | 6.9 | 28.6 | 2.5 |
| 6cnf | M | 4.5 | 7533 | 0.035 | 1.18 | 164 | 19.7 | 17.5 | 0.190 | 0.258 | 61 | 5 | 34.8 | 3.5 |
| 6cnf | P | 4.5 | 7533 | 0.035 | 1.18 | 277 | 20.5 | 10.9 | 0.283 | 0.538 | 64.3 | 5.1 | 46.9 | 3.8 |
| 6cnj | F | 3.7 | 7535 | 0.0327 | 1.07 | 218 | 18.3 | 18.0 | 0.324 | 0.519 | 70.2 | 4.6 | 37.2 | 6.2 |
| 6cnj | G | 3.7 | 7535 | 0.0327 | 1.07 | 218 | 17.7 | 16.7 | 0.445 | 0.454 | 71.6 | 17.3 | 43.1 | 5.3 |
| 6cnj | H | 3.7 | 7535 | 0.0327 | 1.07 | 218 | 23.4 | 20.2 | 0.314 | 0.501 | 71.1 | 6.5 | 40.4 | 8 |
| 6cnj | I | 3.7 | 7535 | 0.0327 | 1.07 | 218 | 14.8 | 12.2 | 0.613 | 0.661 | 77.5 | 47.3 | 25.7 | 16.1 |
| 6cnj | J | 3.7 | 7535 | 0.0327 | 1.07 | 218 | 19.5 | 18.9 | 0.266 | 0.452 | 70.2 | 9.2 | 37.6 | 2.4 |
| 6cnj | K | 3.7 | 7535 | 0.0327 | 1.07 | 218 | 20.3 | 15.2 | 0.280 | 0.636 | 73.9 | 14.9 | 51.8 | 38.1 |
| 6cnk | F | 3.9 | 7536 | 0.0325 | 1.07 | 218 | 17.0 | 17.0 | 0.403 | 0.403 | 70.2 | 11.8 | 26.1 | 5.3 |
| 6cnk | G | 3.9 | 7536 | 0.0325 | 1.07 | 218 | 17.2 | 17.1 | 0.576 | 0.576 | 72.5 | 24.7 | 39 | 11.8 |
| 6cnk | J | 3.9 | 7536 | 0.0325 | 1.07 | 218 | 18.6 | 17.5 | 0.403 | 0.403 | 68.3 | 5.4 | 44.5 | 4.1 |
| 6cnk | K | 3.9 | 7536 | 0.0325 | 1.07 | 218 | 20.2 | 20.2 | 0.445 | 0.446 | 67.9 | 7.4 | 37.6 | 6.1 |
| 6cnm | E | 3.4 | 7537 | 0.04 | 1.03 | 66 | 11.4 | 8.6 | 0.247 | 0.465 | 63.6 | 9.5 | 54.5 | 5.6 |
| 6cnm | F | 3.4 | 7537 | 0.04 | 1.03 | 66 | 5.2 | 5.2 | 0.602 | 0.602 | 74.2 | 38.8 | 50 | 0 |
| 6cnm | G | 3.4 | 7537 | 0.04 | 1.03 | 66 | 10.7 | 6.9 | 0.245 | 0.475 | 75.8 | 12 | 60.6 | 2.5 |
| 6cnm | H | 3.4 | 7537 | 0.04 | 1.03 | 66 | 6.6 | 6.6 | 0.346 | 0.492 | 62.1 | 4.9 | 54.5 | 2.8 |
| 6cnn | E | 3.5 | 7538 | 0.019 | 1.03 | 146 | 6.7 | 4.3 | 0.735 | 0.874 | 91.8 | 65.7 | 77.4 | 13.3 |
| 6cnn | F | 3.5 | 7538 | 0.019 | 1.03 | 146 | 3.3 | 3.3 | 0.885 | 0.885 | 92.5 | 60 | 77.4 | 20.4 |
| 6cnn | G | 3.5 | 7538 | 0.019 | 1.03 | 146 | 4.3 | 4.3 | 0.842 | 0.842 | 88.4 | 35.7 | 71.9 | 5.7 |
| 6cnn | H | 3.5 | 7538 | 0.019 | 1.03 | 146 | 2.9 | 2.9 | 0.861 | 0.861 | 89 | 46.2 | 58.9 | 17.4 |
| 6cno | E | 4.7 | 7539 | 0.16 | 1.03 | 146 | 6.2 | 5.6 | 0.741 | 0.741 | 76.7 | 18.8 | 54.1 | 6.3 |
| 6cno | F | 4.7 | 7539 | 0.16 | 1.03 | 146 | 5.0 | 5.0 | 0.752 | 0.752 | 80.8 | 11.9 | 51.4 | 5.3 |
| 6cno | G | 4.7 | 7539 | 0.16 | 1.03 | 146 | 5.4 | 4.2 | 0.746 | 0.776 | 77.4 | 20.4 | 30.8 | 2.2 |
| 6cno | H | 4.7 | 7539 | 0.16 | 1.03 | 146 | 4.0 | 4.0 | 0.739 | 0.739 | 80.1 | 33.3 | 61 | 6.7 |
| 6cp3 | G | 3.8 | 7546 | 0.0324 | 1.23 | 269 | 17.0 | 12.1 | 0.675 | 0.675 | 80.7 | 23.5 | 75.8 | 22.5 |
| 6cp3 | H | 3.8 | 7546 | 0.0324 | 1.23 | 132 | 12.2 | 10.6 | 0.437 | 0.522 | 78.8 | 9.6 | 59.8 | 3.8 |
| 6cp3 | Y | 3.8 | 7546 | 0.0324 | 1.23 | 166 | 5.1 | 5.1 | 0.814 | 0.814 | 84.3 | 32.1 | 80.7 | 5.2 |
| 6cp6 | G | 3.6 | 7548 | 0.0218 | 1.23 | 269 | 13.6 | 12.9 | 0.657 | 0.675 | 78.1 | 34.8 | 64.3 | 14.5 |
| 6cp6 | Y | 3.6 | 7548 | 0.0218 | 1.23 | 166 | 6.4 | 6.4 | 0.784 | 0.784 | 83.1 | 36.2 | 75.3 | 4 |
| 6crq | D | 4.2 | 7568 | 0.015 | 1.02 | 125 | 16.6 | 8.8 | 0.569 | 0.769 | 80 | 10 | 80.8 | 7.9 |
| 6crq | E | 4.2 | 7568 | 0.015 | 1.02 | 100 | 12.1 | 11.0 | 0.546 | 0.546 | 75 | 33.3 | 59 | 5.1 |

| | | | | | | | | | | | | | | |
|---|---|---|---|---|---|---|---|---|---|---|---|---|---|---|
| 6crq | I | 4.2 | 7568 | 0.015 | 1.02 | 125 | 7.9 | 7.9 | 0.755 | 0.755 | 85.6 | 49.5 | 68.8 | 3.5 |
| 6crq | J | 4.2 | 7568 | 0.015 | 1.02 | 125 | 10.1 | 8.0 | 0.626 | 0.626 | 84.8 | 45.3 | 73.6 | 7.6 |
| 6crq | K | 4.2 | 7568 | 0.015 | 1.02 | 100 | 13.2 | 9.3 | 0.473 | 0.565 | 73 | 30.1 | 42 | 4.8 |
| 6crq | L | 4.2 | 7568 | 0.015 | 1.02 | 100 | 14.2 | 12.4 | 0.299 | 0.485 | 72 | 9.7 | 43 | 4.7 |
| 6cue | 1 | 4 | 7621 | 1.04 | 1.1 | 132 | 4.5 | 4.5 | 0.788 | 0.788 | 83.3 | 34.5 | 71.2 | 6.4 |
| 6cue | 5 | 4 | 7621 | 1.04 | 1.1 | 132 | 18.8 | 16.7 | 0.266 | 0.305 | 71.2 | 6.4 | 24.2 | 3.1 |
| 6cue | 6 | 4 | 7621 | 1.04 | 1.1 | 107 | 16.1 | 13.0 | 0.356 | 0.356 | 65.4 | 12.9 | 31.8 | 0 |
| 6cue | 8 | 4 | 7621 | 1.04 | 1.1 | 102 | 13.7 | 13.4 | 0.281 | 0.495 | 77.5 | 7.6 | 55.9 | 5.3 |
| 6cue | d | 4 | 7621 | 1.04 | 1.1 | 132 | 3.7 | 3.7 | 0.814 | 0.828 | 85.6 | 56.6 | 72.7 | 31.2 |
| 6cue | D | 4 | 7621 | 1.04 | 1.1 | 132 | 6.8 | 6.1 | 0.760 | 0.766 | 80.3 | 44.3 | 72.7 | 32.3 |
| 6cue | m | 4 | 7621 | 1.04 | 1.1 | 132 | 17.9 | 16.3 | 0.202 | 0.421 | 64.4 | 8.2 | 25.8 | 2.9 |
| 6cue | M | 4 | 7621 | 1.04 | 1.1 | 132 | 15.2 | 15.2 | 0.243 | 0.376 | 72 | 7.4 | 35.6 | 8.5 |
| 6cue | n | 4 | 7621 | 1.04 | 1.1 | 107 | 13.7 | 12.6 | 0.331 | 0.362 | 73.8 | 7.6 | 35.5 | 5.3 |
| 6cue | N | 4 | 7621 | 1.04 | 1.1 | 107 | 11.9 | 11.9 | 0.406 | 0.418 | 76.6 | 6.1 | 38.3 | 12.2 |
| 6cue | r | 4 | 7621 | 1.04 | 1.1 | 102 | 14.7 | 12.6 | 0.305 | 0.627 | 82.4 | 11.9 | 49 | 4 |
| 6cue | R | 4 | 7621 | 1.04 | 1.1 | 102 | 7.9 | 7.9 | 0.722 | 0.722 | 87.3 | 40.4 | 68.6 | 15.7 |
| 6cuf | 3 | 4 | 7622 | 1 | 1.06 | 116 | 15.2 | 13.5 | 0.256 | 0.294 | 60.3 | 5.7 | 20.7 | 8.3 |
| 6cuf | 5 | 4 | 7622 | 1 | 1.06 | 132 | 17.0 | 15.1 | 0.248 | 0.302 | 67.4 | 7.9 | 29.5 | 7.7 |
| 6cuf | 6 | 4 | 7622 | 1 | 1.06 | 105 | 16.0 | 13.2 | 0.262 | 0.410 | 64.8 | 11.8 | 38.1 | 0 |
| 6cuf | 7 | 4 | 7622 | 1 | 1.06 | 102 | 14.3 | 10.6 | 0.309 | 0.486 | 79.4 | 8.6 | 51 | 5.8 |
| 6cuf | 8 | 4 | 7622 | 1 | 1.06 | 128 | 15.6 | 12.9 | 0.290 | 0.548 | 76.6 | 8.2 | 51.6 | 3 |
| 6cuf | A | 4 | 7622 | 1 | 1.06 | 132 | 6.2 | 6.2 | 0.772 | 0.772 | 83.3 | 32.7 | 72 | 8.4 |
| 6cuf | c | 4 | 7622 | 1 | 1.06 | 132 | 4.6 | 4.6 | 0.785 | 0.785 | 84.1 | 52.3 | 72.7 | 6.2 |
| 6cuf | D | 4 | 7622 | 1 | 1.06 | 132 | 4.0 | 4.0 | 0.812 | 0.812 | 85.6 | 48.7 | 65.9 | 4.6 |
| 6cuf | h | 4 | 7622 | 1 | 1.06 | 116 | 16.8 | 14.0 | 0.227 | 0.289 | 59.5 | 4.3 | 31.9 | 10.8 |
| 6cuf | H | 4 | 7622 | 1 | 1.06 | 116 | 17.0 | 15.8 | 0.257 | 0.296 | 66.4 | 5.2 | 30.2 | 0 |
| 6cuf | m | 4 | 7622 | 1 | 1.06 | 132 | 18.1 | 16.0 | 0.272 | 0.274 | 68.9 | 5.5 | 34.8 | 2.2 |
| 6cuf | M | 4 | 7622 | 1 | 1.06 | 132 | 19.4 | 16.0 | 0.306 | 0.337 | 62.9 | 4.8 | 34.1 | 4.4 |
| 6cuf | n | 4 | 7622 | 1 | 1.06 | 105 | 16.1 | 13.8 | 0.261 | 0.337 | 67.6 | 5.6 | 31.4 | 0 |
| 6cuf | N | 4 | 7622 | 1 | 1.06 | 105 | 14.4 | 13.6 | 0.255 | 0.328 | 65.7 | 8.7 | 30.5 | 3.1 |
| 6cuf | q | 4 | 7622 | 1 | 1.06 | 128 | 11.1 | 9.4 | 0.627 | 0.653 | 80.5 | 16.5 | 40.6 | 5.8 |
| 6cuf | Q | 4 | 7622 | 1 | 1.06 | 128 | 16.1 | 11.6 | 0.349 | 0.480 | 78.1 | 8 | 43.8 | 5.4 |
| 6cuf | r | 4 | 7622 | 1 | 1.06 | 102 | 13.7 | 13.0 | 0.369 | 0.480 | 82.4 | 7.1 | 53.9 | 3.6 |
| 6cuf | R | 4 | 7622 | 1 | 1.06 | 102 | 15.4 | 13.0 | 0.290 | 0.470 | 73.5 | 14.7 | 67.6 | 4.3 |
| 6cxc | G | 3.9 | 7774 | 0.0232 | 1.2546 | 114 | 12.9 | 12.9 | 0.289 | 0.313 | 58.8 | 7.5 | 43.9 | 4 |
| 6cxc | I | 3.9 | 7774 | 0.0232 | 1.2546 | 114 | 14.3 | 13.6 | 0.295 | 0.330 | 57.9 | 1.5 | 46.5 | 5.7 |
| 6cxc | J | 3.9 | 7774 | 0.0232 | 1.2546 | 108 | 16.6 | 14.8 | 0.288 | 0.329 | 58.3 | 9.5 | 36.1 | 10.3 |

| | | | | | | | | | | | | | | |
|---|---|---|---|---|---|---|---|---|---|---|---|---|---|---|
| 6cxc | K | 3.9 | 7774 | 0.0232 | 1.2546 | 108 | 14.5 | 14.0 | 0.262 | 0.312 | 63 | 7.4 | 44.4 | 4.2 |
| 6cxc | X | 3.9 | 7774 | 0.0232 | 1.2546 | 114 | 16.2 | 14.7 | 0.241 | 0.275 | 59.6 | 5.9 | 47.4 | 3.7 |
| 6cxc | Y | 3.9 | 7774 | 0.0232 | 1.2546 | 108 | 14.7 | 13.4 | 0.252 | 0.271 | 68.5 | 6.8 | 34.3 | 5.4 |
| 6d6q | A | 3.5 | 7808 | 0.025 | 1.07 | 287 | 2.1 | 2.1 | 0.942 | 0.942 | 94.8 | 80.9 | 90.6 | 25 |
| 6d6q | C | 3.5 | 7808 | 0.025 | 1.07 | 265 | 2.9 | 2.9 | 0.920 | 0.920 | 93.2 | 63.2 | 84.2 | 9 |
| 6d6q | D | 3.5 | 7808 | 0.025 | 1.07 | 208 | 1.9 | 1.9 | 0.931 | 0.931 | 96.2 | 84.5 | 94.2 | 57.7 |
| 6d6q | E | 3.5 | 7808 | 0.025 | 1.07 | 286 | 13.8 | 12.0 | 0.713 | 0.725 | 85.7 | 45.7 | 87.4 | 32 |
| 6d6q | F | 3.5 | 7808 | 0.025 | 1.07 | 252 | 12.5 | 12.0 | 0.767 | 0.768 | 89.7 | 65 | 81.3 | 32.2 |
| 6d6q | I | 3.5 | 7808 | 0.025 | 1.07 | 183 | 11.6 | 11.0 | 0.517 | 0.576 | 74.9 | 24.1 | 56.3 | 7.8 |
| 6d6r | A | 3.5 | 7809 | 0.037 | 1.07 | 287 | 11.6 | 11.6 | 0.698 | 0.698 | 83.6 | 42.5 | 91.6 | 46 |
| 6d6r | C | 3.5 | 7809 | 0.037 | 1.07 | 265 | 8.2 | 7.2 | 0.843 | 0.843 | 87.5 | 51.3 | 89.8 | 47.1 |
| 6d6r | D | 3.5 | 7809 | 0.037 | 1.07 | 208 | 6.7 | 3.9 | 0.853 | 0.883 | 91.8 | 64.4 | 94.2 | 50.5 |
| 6d6r | E | 3.5 | 7809 | 0.037 | 1.07 | 286 | 13.9 | 12.9 | 0.689 | 0.695 | 83.9 | 34.2 | 78.3 | 54 |
| 6d6r | F | 3.5 | 7809 | 0.037 | 1.07 | 252 | 17.1 | 15.5 | 0.625 | 0.634 | 77.4 | 32.8 | 82.1 | 35.3 |
| 6d6r | I | 3.5 | 7809 | 0.037 | 1.07 | 183 | 13.7 | 12.4 | 0.438 | 0.457 | 67.2 | 13.8 | 62.3 | 26.3 |
| 6d6t | I | 3.9 | 7816 | 0.024 | 1.07 | 106 | 14.5 | 12.5 | 0.545 | 0.755 | 87.7 | 33.3 | 50 | 5.7 |
| 6d6t | J | 3.9 | 7816 | 0.024 | 1.07 | 116 | 1.7 | 1.7 | 0.935 | 0.935 | 97.4 | 77.9 | 86.2 | 32 |
| 6d6t | K | 3.9 | 7816 | 0.024 | 1.07 | 116 | 3.2 | 3.2 | 0.895 | 0.895 | 95.7 | 70.3 | 87.1 | 5.9 |
| 6d6t | L | 3.9 | 7816 | 0.024 | 1.07 | 106 | 2.2 | 2.2 | 0.868 | 0.868 | 92.5 | 65.3 | 85.8 | 11 |
| 6d6u | I | 3.9 | 7817 | 0.024 | 1.07 | 106 | 1.9 | 1.9 | 0.896 | 0.896 | 97.2 | 84.5 | 92.5 | 5.1 |
| 6d6u | J | 3.9 | 7817 | 0.024 | 1.07 | 116 | 1.9 | 1.9 | 0.900 | 0.900 | 95.7 | 73.9 | 75.9 | 5.7 |
| 6d6u | K | 3.9 | 7817 | 0.024 | 1.07 | 116 | 3.1 | 3.1 | 0.902 | 0.902 | 95.7 | 64 | 81 | 7.4 |
| 6d6u | L | 3.9 | 7817 | 0.024 | 1.07 | 106 | 7.3 | 2.2 | 0.847 | 0.859 | 92.5 | 64.3 | 76.4 | 4.9 |
| 6d6v | D | 4.8 | 7821 | 0.055 | 1.36 | 185 | 21.9 | 19.4 | 0.269 | 0.332 | 63.8 | 6.8 | 37.3 | 2.9 |
| 6d6v | E | 4.8 | 7821 | 0.055 | 1.36 | 149 | 18.7 | 14.6 | 0.297 | 0.411 | 66.4 | 6.1 | 44.3 | 3 |
| 6d6v | F | 4.8 | 7821 | 0.055 | 1.36 | 113 | 16.5 | 14.0 | 0.271 | 0.448 | 61.9 | 10 | 45.1 | 2 |
| 6d7w | A | 3.8 | 7826 | 0.014 | 0.84 | 275 | 5.2 | 5.2 | 0.827 | 0.845 | 83.6 | 30 | 62.2 | 19.9 |
| 6d7w | B | 3.8 | 7826 | 0.014 | 0.84 | 274 | 6.8 | 6.0 | 0.827 | 0.830 | 80.7 | 24.9 | 55.1 | 31.1 |
| 6d7w | C | 3.8 | 7826 | 0.014 | 0.84 | 269 | 6.1 | 4.8 | 0.812 | 0.832 | 82.2 | 38.9 | 72.1 | 7.2 |
| 6d7w | D | 3.8 | 7826 | 0.014 | 0.84 | 269 | 7.6 | 4.9 | 0.838 | 0.838 | 84 | 32.7 | 58 | 7.1 |
| 6d83 | C | 4.3 | 7453 | 0.0303 | 1.067 | 165 | 14.1 | 13.9 | 0.283 | 0.367 | 63 | 8.7 | 55.2 | 2.2 |
| 6d83 | H | 4.3 | 7453 | 0.0303 | 1.067 | 163 | 15.7 | 13.6 | 0.256 | 0.381 | 74.8 | 9 | 35.6 | 1.7 |
| 6d9h | R | 3.6 | 7835 | 0.06 | 1.06 | 288 | 3.3 | 3.3 | 0.914 | 0.914 | 91.7 | 63.3 | 83.3 | 56.7 |
| 6dcq | B | 3.1 | 7858 | 0.0583 | 1.03 | 122 | 7.2 | 6.0 | 0.837 | 0.837 | 86.1 | 55.2 | 88.5 | 55.6 |
| 6dcq | D | 3.1 | 7858 | 0.0583 | 1.03 | 146 | 2.5 | 2.5 | 0.927 | 0.927 | 95.9 | 66.4 | 84.9 | 33.1 |
| 6dcq | H | 3.1 | 7858 | 0.0583 | 1.03 | 132 | 18.2 | 14.6 | 0.282 | 0.578 | 80.3 | 7.5 | 62.9 | 26.5 |
| 6dcq | M | 3.1 | 7858 | 0.0583 | 1.03 | 135 | 7.5 | 5.6 | 0.784 | 0.845 | 88.1 | 75.6 | 79.3 | 35.5 |

| | | | | | | | | | | | | | | |
|---|---|---|---|---|---|---|---|---|---|---|---|---|---|---|
| 6dcq | N | 3.1 | 7858 | 0.0583 | 1.03 | 112 | 14.5 | 12.0 | 0.440 | 0.497 | 75.9 | 18.8 | 43.8 | 4.1 |
| 6dde | E | 3.5 | 7868 | 3.91 | 1.04 | 232 | 16.7 | 14.8 | 0.500 | 0.590 | 72.4 | 5.4 | 29.7 | 5.8 |
| 6dde | R | 3.5 | 7868 | 3.91 | 1.04 | 281 | 10.6 | 9.1 | 0.449 | 0.781 | 79 | 8.6 | 49.5 | 9.4 |
| 6ddf | R | 3.5 | 7869 | 3.81 | 1.04 | 281 | 10.2 | 6.2 | 0.683 | 0.812 | 81.9 | 33.5 | 79.4 | 13.9 |
| 6ddg | V | 3.1 | 7870 | 0.03 | 1.06 | 143 | 2.3 | 2.3 | 0.927 | 0.927 | 95.8 | 78.1 | 88.8 | 7.9 |
| 6dfg | B | 4.4 | 7875 | 0.04 | 1.03 | 115 | 15.1 | 8.0 | 0.295 | 0.736 | 74.8 | 9.3 | 71.3 | 6.1 |
| 6dfg | E | 4.4 | 7875 | 0.04 | 1.03 | 115 | 15.7 | 13.1 | 0.263 | 0.618 | 71.3 | 9.8 | 80 | 6.5 |
| 6dfg | F | 4.4 | 7875 | 0.04 | 1.03 | 115 | 14.0 | 13.6 | 0.578 | 0.605 | 79.1 | 7.7 | 78.3 | 10 |
| 6dfg | G | 4.4 | 7875 | 0.04 | 1.03 | 129 | 16.6 | 16.6 | 0.266 | 0.324 | 69 | 7.9 | 36.4 | 4.3 |
| 6dfg | H | 4.4 | 7875 | 0.04 | 1.03 | 129 | 16.9 | 16.5 | 0.271 | 0.359 | 63.6 | 8.5 | 29.5 | 2.6 |
| 6dfg | I | 4.4 | 7875 | 0.04 | 1.03 | 129 | 16.1 | 15.9 | 0.285 | 0.355 | 66.7 | 9.3 | 34.1 | 9.1 |
| 6dfg | J | 4.4 | 7875 | 0.04 | 1.03 | 106 | 15.2 | 13.2 | 0.315 | 0.315 | 68.9 | 6.8 | 28.3 | 3.3 |
| 6dfg | K | 4.4 | 7875 | 0.04 | 1.03 | 106 | 14.2 | 14.2 | 0.255 | 0.368 | 62.3 | 9.1 | 26.4 | 0 |
| 6dfg | L | 4.4 | 7875 | 0.04 | 1.03 | 106 | 12.2 | 12.2 | 0.288 | 0.302 | 63.2 | 11.9 | 30.2 | 6.2 |
| 6dfh | B | 3.9 | 7876 | 0.03 | 1.03 | 128 | 19.3 | 16.6 | 0.285 | 0.658 | 78.1 | 9 | 63.3 | 35.8 |
| 6dfh | E | 3.9 | 7876 | 0.03 | 1.03 | 128 | 18.0 | 11.3 | 0.671 | 0.741 | 75 | 12.5 | 69.5 | 3.4 |
| 6dfh | F | 3.9 | 7876 | 0.03 | 1.03 | 128 | 21.1 | 7.2 | 0.351 | 0.732 | 76.6 | 16.3 | 39.1 | 2 |
| 6dfh | G | 3.9 | 7876 | 0.03 | 1.03 | 128 | 16.4 | 11.7 | 0.344 | 0.533 | 71.9 | 6.5 | 35.9 | 8.7 |
| 6dfh | H | 3.9 | 7876 | 0.03 | 1.03 | 128 | 15.4 | 14.3 | 0.301 | 0.436 | 66.4 | 7.1 | 29.7 | 7.9 |
| 6dfh | I | 3.9 | 7876 | 0.03 | 1.03 | 128 | 17.4 | 14.6 | 0.313 | 0.454 | 75 | 8.3 | 51.6 | 7.6 |
| 6dfh | J | 3.9 | 7876 | 0.03 | 1.03 | 106 | 14.2 | 13.8 | 0.582 | 0.582 | 80.2 | 23.5 | 54.7 | 8.6 |
| 6dfh | K | 3.9 | 7876 | 0.03 | 1.03 | 106 | 13.2 | 6.9 | 0.505 | 0.767 | 87.7 | 6.5 | 64.2 | 10.3 |
| 6dfh | L | 3.9 | 7876 | 0.03 | 1.03 | 106 | 15.0 | 11.3 | 0.314 | 0.475 | 68.9 | 6.8 | 57.5 | 4.9 |
| 6did | C | 4.7 | 7896 | 0.35 | 1.15 | 217 | 19.1 | 17.9 | 0.281 | 0.299 | 54.8 | 10.1 | 30 | 9.2 |
| 6did | D | 4.7 | 7896 | 0.35 | 1.15 | 213 | 19.9 | 14.6 | 0.250 | 0.335 | 60.1 | 3.9 | 22.5 | 8.3 |
| 6did | E | 4.7 | 7896 | 0.35 | 1.15 | 217 | 22.1 | 18.7 | 0.263 | 0.310 | 56.7 | 7.3 | 26.3 | 7 |
| 6did | H | 4.7 | 7896 | 0.35 | 1.15 | 217 | 19.5 | 16.9 | 0.261 | 0.303 | 58.5 | 11 | 23 | 6 |
| 6did | K | 4.7 | 7896 | 0.35 | 1.15 | 213 | 16.7 | 16.7 | 0.352 | 0.352 | 60.1 | 12.5 | 21.1 | 6.7 |
| 6did | L | 4.7 | 7896 | 0.35 | 1.15 | 213 | 19.6 | 17.2 | 0.282 | 0.313 | 54.9 | 10.3 | 23.9 | 9.8 |
| 6djp | D | 4.8 | 7939 | 0.25 | 1.67 | 107 | 15.7 | 15.1 | 0.236 | 0.303 | 58.9 | 3.2 | 31.8 | 11.8 |
| 6djp | E | 4.8 | 7939 | 0.25 | 1.67 | 115 | 13.9 | 13.4 | 0.278 | 0.291 | 58.3 | 7.5 | 41.7 | 4.2 |
| 6djp | F | 4.8 | 7939 | 0.25 | 1.67 | 108 | 16.2 | 13.9 | 0.253 | 0.283 | 62 | 1.5 | 50 | 20.4 |
| 6dmy | B | 3.6 | 7968 | 0.05 | 1.091 | 151 | 9.8 | 9.8 | 0.637 | 0.637 | 82.1 | 45.2 | 70.9 | 3.7 |
| 6dnf | A | 3.2 | 7971 | 2.8 | 0.861 | 225 | 33.1 | 26.2 | 0.650 | 0.694 | 78.2 | 43.8 | 77.8 | 19.4 |
| 6dnf | B | 3.2 | 7971 | 2.8 | 0.861 | 225 | 43.6 | 31.3 | 0.405 | 0.487 | 81.8 | 3.8 | 73.8 | 16.9 |
| 6dnf | C | 3.2 | 7971 | 2.8 | 0.861 | 225 | 34.0 | 24.6 | 0.581 | 0.581 | 69.3 | 32.7 | 77.3 | 19 |
| 6dnf | D | 3.2 | 7971 | 2.8 | 0.861 | 225 | 41.0 | 29.4 | 0.409 | 0.589 | 82.2 | 10.8 | 69.3 | 20.5 |

| | | | | | | | | | | | | | | |
|---|---|---|---|---|---|---|---|---|---|---|---|---|---|---|
| 6drd | C | 3.9 | 7997 | 0.035 | 1.31 | 262 | 14.0 | 13.9 | 0.615 | 0.680 | 88.2 | 33.3 | 81.7 | 17.3 |
| 6drd | E | 3.9 | 7997 | 0.035 | 1.31 | 185 | 9.9 | 7.6 | 0.614 | 0.724 | 81.1 | 36.7 | 77.8 | 18.8 |
| 6drd | F | 3.9 | 7997 | 0.035 | 1.31 | 68 | 14.0 | 9.6 | 0.358 | 0.403 | 79.4 | 13 | 76.5 | 40.4 |
| 6drd | H | 3.9 | 7997 | 0.035 | 1.31 | 146 | 11.5 | 10.0 | 0.589 | 0.735 | 82.2 | 19.2 | 61 | 6.7 |
| 6drd | I | 3.9 | 7997 | 0.035 | 1.31 | 110 | 10.4 | 10.4 | 0.666 | 0.674 | 79.1 | 39.1 | 47.3 | 5.8 |
| 6drd | K | 3.9 | 7997 | 0.035 | 1.31 | 111 | 2.4 | 2.4 | 0.864 | 0.864 | 92.8 | 60.2 | 86.5 | 8.3 |
| 6dt0 | A | 3.7 | 8911 | 0.07 | 1.15 | 236 | 32.3 | 30.2 | 0.715 | 0.715 | 83.5 | 41.6 | 77.1 | 29.1 |
| 6dt0 | B | 3.7 | 8911 | 0.07 | 1.15 | 236 | 29.4 | 20.1 | 0.686 | 0.730 | 81.8 | 39.9 | 82.2 | 36.1 |
| 6dt0 | C | 3.7 | 8911 | 0.07 | 1.15 | 236 | 34.5 | 25.4 | 0.677 | 0.738 | 82.6 | 29.7 | 81.4 | 14.6 |
| 6dt0 | D | 3.7 | 8911 | 0.07 | 1.15 | 236 | 27.1 | 23.3 | 0.664 | 0.748 | 77.1 | 25.3 | 84.3 | 28.1 |
| 6dw1 | A | 3.1 | 8923 | 7.68 | 0.649 | 210 | 1.6 | 1.6 | 0.945 | 0.945 | 97.1 | 86.8 | 86.7 | 9.9 |
| 6dw1 | B | 3.1 | 8923 | 7.68 | 0.649 | 210 | 2.6 | 2.6 | 0.925 | 0.925 | 95.2 | 68.5 | 86.2 | 35.9 |
| 6dw1 | C | 3.1 | 8923 | 7.68 | 0.649 | 210 | 2.1 | 2.1 | 0.937 | 0.937 | 95.7 | 73.6 | 84.8 | 5.1 |
| 6dw1 | D | 3.1 | 8923 | 7.68 | 0.649 | 208 | 3.6 | 3.6 | 0.881 | 0.881 | 88.9 | 65.9 | 76.4 | 10.1 |
| 6dw1 | E | 3.1 | 8923 | 7.68 | 0.649 | 208 | 3.0 | 3.0 | 0.924 | 0.924 | 94.2 | 73.5 | 94.7 | 7.6 |
| 6dwb | A | 3.3 | 8924 | 0.15 | 1.35 | 78 | 7.8 | 7.6 | 0.481 | 0.493 | 76.9 | 5 | 82.1 | 95.3 |
| 6dwb | C | 3.3 | 8924 | 0.15 | 1.35 | 78 | 1.4 | 1.4 | 0.912 | 0.912 | 97.4 | 90.8 | 92.3 | 54.2 |
| 6dwb | D | 3.3 | 8924 | 0.15 | 1.35 | 78 | 1.3 | 1.3 | 0.912 | 0.912 | 96.2 | 96 | 88.5 | 55.1 |
| 6dwb | E | 3.3 | 8924 | 0.15 | 1.35 | 78 | 1.2 | 1.2 | 0.936 | 0.936 | 97.4 | 98.7 | 92.3 | 56.9 |
| 6dwb | F | 3.3 | 8924 | 0.15 | 1.35 | 78 | 1.4 | 1.4 | 0.927 | 0.927 | 96.2 | 90.7 | 87.2 | 98.5 |
| 6dwb | G | 3.3 | 8924 | 0.15 | 1.35 | 78 | 0.8 | 0.8 | 0.947 | 0.947 | 100 | 100 | 93.6 | 100 |
| 6dwb | I | 3.3 | 8924 | 0.15 | 1.35 | 78 | 0.7 | 0.7 | 0.955 | 0.955 | 100 | 100 | 87.2 | 57.4 |
| 6dzk | C | 3.6 | 8934 | 0.02 | 1.07 | 208 | 9.3 | 4.8 | 0.796 | 0.807 | 81.2 | 37.3 | 71.2 | 7.4 |
| 6dzk | D | 3.6 | 8934 | 0.02 | 1.07 | 200 | 7.1 | 7.1 | 0.781 | 0.835 | 81.5 | 39.9 | 58.5 | 4.3 |
| 6dzk | E | 3.6 | 8934 | 0.02 | 1.07 | 180 | 3.8 | 3.8 | 0.874 | 0.874 | 90.6 | 63.2 | 79.4 | 22.4 |
| 6dzk | F | 3.6 | 8934 | 0.02 | 1.07 | 96 | 3.4 | 3.4 | 0.799 | 0.799 | 84.4 | 54.3 | 71.9 | 4.3 |
| 6dzk | H | 3.6 | 8934 | 0.02 | 1.07 | 131 | 3.6 | 3.6 | 0.834 | 0.842 | 88.5 | 64.7 | 80.2 | 28.6 |
| 6dzl | A | 4.1 | 8935 | 0.0455 | 1.15 | 213 | 16.3 | 15.4 | 0.606 | 0.607 | 75.6 | 6.2 | 48.8 | 4.8 |
| 6dzl | B | 4.1 | 8935 | 0.0455 | 1.15 | 213 | 17.9 | 16.7 | 0.544 | 0.603 | 76.5 | 14.7 | 46.5 | 11.1 |
| 6dzl | C | 4.1 | 8935 | 0.0455 | 1.15 | 213 | 19.4 | 17.7 | 0.404 | 0.430 | 75.1 | 11.9 | 74.6 | 8.8 |
| 6dzl | G | 4.1 | 8935 | 0.0455 | 1.15 | 121 | 11.1 | 11.1 | 0.391 | 0.601 | 68.6 | 8.4 | 53.7 | 15.4 |
| 6dzl | H | 4.1 | 8935 | 0.0455 | 1.15 | 121 | 15.6 | 14.1 | 0.300 | 0.675 | 73.6 | 6.7 | 50.4 | 13.1 |
| 6dzl | I | 4.1 | 8935 | 0.0455 | 1.15 | 121 | 11.0 | 11.0 | 0.702 | 0.702 | 76.9 | 30.1 | 52.1 | 1.6 |
| 6dzl | J | 4.1 | 8935 | 0.0455 | 1.15 | 106 | 15.5 | 13.4 | 0.356 | 0.359 | 74.5 | 6.3 | 42.5 | 0 |
| 6dzl | K | 4.1 | 8935 | 0.0455 | 1.15 | 106 | 14.9 | 13.3 | 0.268 | 0.498 | 65.1 | 13 | 43.4 | 10.9 |
| 6dzl | L | 4.1 | 8935 | 0.0455 | 1.15 | 106 | 16.4 | 11.8 | 0.388 | 0.490 | 73.6 | 12.8 | 39.6 | 0 |
| 6dzm | A | 4.3 | 8936 | 0.0393 | 1.31 | 224 | 19.7 | 18.6 | 0.435 | 0.530 | 75.9 | 8.8 | 38.8 | 4.6 |

| | | | | | | | | | | | | | | |
|---|---|---|---|---|---|---|---|---|---|---|---|---|---|---|
| 6dzm | B | 4.3 | 8936 | 0.0393 | 1.31 | 224 | 25.8 | 16.8 | 0.265 | 0.480 | 69.6 | 7.1 | 26.3 | 0 |
| 6dzm | C | 4.3 | 8936 | 0.0393 | 1.31 | 224 | 22.9 | 16.5 | 0.482 | 0.508 | 73.7 | 9.7 | 35.7 | 8.8 |
| 6dzm | G | 4.3 | 8936 | 0.0393 | 1.31 | 120 | 15.5 | 14.8 | 0.304 | 0.419 | 61.7 | 10.8 | 54.2 | 1.5 |
| 6dzm | H | 4.3 | 8936 | 0.0393 | 1.31 | 106 | 13.5 | 12.9 | 0.267 | 0.339 | 68.9 | 5.5 | 38.7 | 7.3 |
| 6dzm | I | 4.3 | 8936 | 0.0393 | 1.31 | 120 | 14.7 | 14.6 | 0.319 | 0.450 | 73.3 | 9.1 | 28.3 | 8.8 |
| 6dzm | J | 4.3 | 8936 | 0.0393 | 1.31 | 120 | 16.5 | 14.3 | 0.318 | 0.413 | 70 | 8.3 | 36.7 | 6.8 |
| 6dzm | K | 4.3 | 8936 | 0.0393 | 1.31 | 106 | 14.6 | 12.6 | 0.344 | 0.412 | 67.9 | 4.2 | 56.6 | 8.3 |
| 6dzm | L | 4.3 | 8936 | 0.0393 | 1.31 | 106 | 15.9 | 11.3 | 0.296 | 0.296 | 74.5 | 7.6 | 32.1 | 8.8 |
| 6dzp | G | 3.4 | 8937 | 0.033 | 1.07 | 176 | 14.1 | 12.2 | 0.573 | 0.629 | 76.7 | 10.4 | 65.3 | 6.1 |
| 6dzp | P | 3.4 | 8937 | 0.033 | 1.07 | 126 | 4.2 | 4.2 | 0.793 | 0.793 | 86.5 | 71.6 | 78.6 | 45.5 |
| 6dzp | W | 3.4 | 8937 | 0.033 | 1.07 | 192 | 10.6 | 8.2 | 0.662 | 0.876 | 89.1 | 30.4 | 73.4 | 28.4 |
| 6dzp | Z | 3.4 | 8937 | 0.033 | 1.07 | 64 | 2.1 | 2.1 | 0.832 | 0.832 | 93.8 | 83.3 | 92.2 | 45.8 |
| 6dzt | M | 3 | 8938 | 0.025 | 0.9926 | 226 | 11.9 | 11.9 | 0.559 | 0.597 | 79.2 | 45.8 | 75.2 | 44.1 |
| 6dzt | N | 3 | 8938 | 0.025 | 0.9926 | 226 | 12.2 | 11.0 | 0.709 | 0.709 | 85.8 | 42.8 | 77 | 25.3 |
| 6e0c | A | 2.6 | 8945 | 0.014 | 1.005 | 99 | 1.5 | 1.5 | 0.920 | 0.920 | 96 | 91.6 | 83.8 | 88 |
| 6e0c | B | 2.6 | 8945 | 0.014 | 1.005 | 80 | 1.5 | 1.5 | 0.926 | 0.926 | 96.2 | 94.8 | 95 | 75 |
| 6e0c | C | 2.6 | 8945 | 0.014 | 1.005 | 107 | 1.8 | 1.8 | 0.932 | 0.932 | 97.2 | 82.7 | 87.9 | 45.7 |
| 6e0c | D | 2.6 | 8945 | 0.014 | 1.005 | 95 | 1.3 | 1.3 | 0.927 | 0.927 | 97.9 | 93.5 | 91.6 | 87.4 |
| 6e0c | E | 2.6 | 8945 | 0.014 | 1.005 | 99 | 2.1 | 2.1 | 0.900 | 0.900 | 93.9 | 83.9 | 88.9 | 98.9 |
| 6e0c | F | 2.6 | 8945 | 0.014 | 1.005 | 80 | 1.1 | 1.1 | 0.926 | 0.926 | 96.2 | 96.1 | 90 | 79.2 |
| 6e0c | G | 2.6 | 8945 | 0.014 | 1.005 | 108 | 1.3 | 1.3 | 0.948 | 0.948 | 97.2 | 100 | 88.9 | 82.3 |
| 6e0c | H | 2.6 | 8945 | 0.014 | 1.005 | 94 | 1.4 | 1.4 | 0.925 | 0.925 | 96.8 | 87.9 | 97.9 | 100 |
| 6e0c | M | 2.6 | 8945 | 0.014 | 1.005 | 229 | 14.7 | 13.9 | 0.554 | 0.628 | 84.7 | 41.2 | 88.6 | 26.1 |
| 6e0c | N | 2.6 | 8945 | 0.014 | 1.005 | 229 | 14.7 | 11.6 | 0.588 | 0.742 | 87.3 | 47 | 80.8 | 47 |
| 6e0p | A | 2.6 | 8949 | 0.014 | 1.005 | 99 | 1.8 | 1.8 | 0.903 | 0.903 | 96 | 84.2 | 82.8 | 100 |
| 6e0p | B | 2.6 | 8949 | 0.014 | 1.005 | 80 | 1.6 | 1.6 | 0.928 | 0.928 | 97.5 | 96.2 | 92.5 | 100 |
| 6e0p | C | 2.6 | 8949 | 0.014 | 1.005 | 107 | 2.5 | 2.5 | 0.911 | 0.911 | 94.4 | 78.2 | 88.8 | 88.4 |
| 6e0p | D | 2.6 | 8949 | 0.014 | 1.005 | 95 | 1.4 | 1.4 | 0.933 | 0.933 | 97.9 | 87.1 | 97.9 | 97.8 |
| 6e0p | E | 2.6 | 8949 | 0.014 | 1.005 | 99 | 1.1 | 1.1 | 0.947 | 0.947 | 98 | 99 | 83.8 | 89.2 |
| 6e0p | F | 2.6 | 8949 | 0.014 | 1.005 | 80 | 1.4 | 1.4 | 0.921 | 0.921 | 96.2 | 97.4 | 96.2 | 100 |
| 6e0p | G | 2.6 | 8949 | 0.014 | 1.005 | 108 | 1.3 | 1.3 | 0.946 | 0.946 | 96.3 | 92.3 | 92.6 | 91 |
| 6e0p | H | 2.6 | 8949 | 0.014 | 1.005 | 94 | 1.0 | 1.0 | 0.951 | 0.951 | 98.9 | 96.8 | 96.8 | 100 |
| 6e0p | M | 2.6 | 8949 | 0.014 | 1.005 | 229 | 14.3 | 11.6 | 0.726 | 0.729 | 90 | 48.1 | 77.7 | 48.9 |
| 6e0p | N | 2.6 | 8949 | 0.014 | 1.005 | 229 | 13.9 | 13.3 | 0.605 | 0.694 | 90 | 45.6 | 80.3 | 40.2 |
| 6e11 | G | 4.2 | 8952 | 0.0405 | 1.04 | 191 | 4.2 | 4.2 | 0.852 | 0.852 | 88.5 | 42 | 84.8 | 34 |
| 6e1h | C | 3.5 | 8955 | 0.436 | 1 | 164 | 3.2 | 3.2 | 0.849 | 0.849 | 87.2 | 46.9 | 57.3 | 10.6 |
| 6e2f | E | 3.9 | 8961 | 0.34 | 1.08 | 148 | 5.0 | 5.0 | 0.762 | 0.762 | 80.4 | 30.3 | 39.2 | 5.2 |

| | | | | | | | | | | | | | | |
|---|---|---|---|---|---|---|---|---|---|---|---|---|---|---|
| 6e2g | E | 3.6 | 8962 | 0.4 | 1.06 | 148 | 5.1 | 5.1 | 0.754 | 0.754 | 77.7 | 22.6 | 33.8 | 0 |
| 6e3y | E | 3.3 | 8978 | 0.04 | 1.06 | 115 | 7.9 | 3.0 | 0.514 | 0.803 | 85.2 | 39.8 | 83.5 | 6.2 |
| 6e3y | N | 3.3 | 8978 | 0.04 | 1.06 | 126 | 14.7 | 8.6 | 0.357 | 0.745 | 84.9 | 6.5 | 61.9 | 5.1 |
| 6edu | G | 4.1 | 9038 | 0.041 | 1.31 | 97 | 13.4 | 6.0 | 0.430 | 0.751 | 76.3 | 5.4 | 29.9 | 13.8 |
| 6edu | H | 4.1 | 9038 | 0.041 | 1.31 | 97 | 11.4 | 8.5 | 0.633 | 0.633 | 74.2 | 16.7 | 59.8 | 3.4 |
| 6edu | I | 4.1 | 9038 | 0.041 | 1.31 | 97 | 13.6 | 11.7 | 0.519 | 0.581 | 77.3 | 8 | 58.8 | 3.5 |
| 6edu | J | 4.1 | 9038 | 0.041 | 1.31 | 124 | 17.2 | 14.7 | 0.339 | 0.385 | 71.8 | 11.2 | 42.7 | 3.8 |
| 6edu | K | 4.1 | 9038 | 0.041 | 1.31 | 111 | 14.0 | 12.5 | 0.282 | 0.308 | 64.9 | 13.9 | 47.7 | 3.8 |
| 6edu | M | 4.1 | 9038 | 0.041 | 1.31 | 111 | 15.5 | 14.3 | 0.281 | 0.317 | 63.1 | 10 | 38.7 | 9.3 |
| 6edu | N | 4.1 | 9038 | 0.041 | 1.31 | 124 | 15.4 | 14.1 | 0.271 | 0.328 | 66.9 | 6 | 31.5 | 7.7 |
| 6edu | O | 4.1 | 9038 | 0.041 | 1.31 | 111 | 16.4 | 14.0 | 0.254 | 0.287 | 60.4 | 4.5 | 32.4 | 8.3 |
| 6edu | P | 4.1 | 9038 | 0.041 | 1.31 | 131 | 18.4 | 13.9 | 0.272 | 0.479 | 74 | 6.2 | 38.2 | 6 |
| 6edu | Q | 4.1 | 9038 | 0.041 | 1.31 | 107 | 13.2 | 11.9 | 0.405 | 0.602 | 74.8 | 6.2 | 58.9 | 14.3 |
| 6edu | R | 4.1 | 9038 | 0.041 | 1.31 | 131 | 16.2 | 15.4 | 0.335 | 0.417 | 71.8 | 13.8 | 27.5 | 8.3 |
| 6edu | S | 4.1 | 9038 | 0.041 | 1.31 | 107 | 15.5 | 13.9 | 0.225 | 0.308 | 64.5 | 7.2 | 57 | 9.8 |
| 6edu | T | 4.1 | 9038 | 0.041 | 1.31 | 131 | 16.6 | 15.3 | 0.294 | 0.411 | 74.8 | 6.1 | 19.8 | 0 |
| 6edu | U | 4.1 | 9038 | 0.041 | 1.31 | 107 | 15.5 | 12.4 | 0.312 | 0.448 | 64.5 | 11.6 | 57 | 6.6 |
| 6ee8 | A | 3.9 | 9039 | 0.3 | 1.3 | 225 | 15.6 | 14.4 | 0.586 | 0.607 | 80.4 | 17.7 | 48.9 | 5.5 |
| 6ee8 | B | 3.9 | 9039 | 0.3 | 1.3 | 237 | 20.8 | 18.3 | 0.319 | 0.338 | 78.5 | 9.7 | 66.2 | 8.9 |
| 6ee8 | M | 3.9 | 9039 | 0.3 | 1.3 | 159 | 9.9 | 9.3 | 0.558 | 0.684 | 75.5 | 22.5 | 69.2 | 3.6 |
| 6eec | A | 3.6 | 9041 | 0.3 | 1.3 | 225 | 3.2 | 3.2 | 0.917 | 0.924 | 93.8 | 67.3 | 85.8 | 45.1 |
| 6eec | B | 3.6 | 9041 | 0.3 | 1.3 | 237 | 3.2 | 3.2 | 0.902 | 0.902 | 92.8 | 64.5 | 76.4 | 24.3 |
| 6eec | E | 3.6 | 9041 | 0.3 | 1.3 | 83 | 2.9 | 2.9 | 0.819 | 0.819 | 88 | 63 | 72.3 | 48.3 |
| 6eec | M | 3.6 | 9041 | 0.3 | 1.3 | 159 | 9.3 | 6.5 | 0.737 | 0.791 | 80.5 | 30.5 | 80.5 | 40.6 |
| 6ef0 | A | 4.4 | 9042 | 0.03 | 1.03 | 238 | 7.0 | 5.2 | 0.796 | 0.818 | 78.2 | 26.9 | 78.6 | 3.7 |
| 6ef0 | B | 4.4 | 9042 | 0.03 | 1.03 | 250 | 10.0 | 10.0 | 0.705 | 0.705 | 79.2 | 46 | 74.8 | 11.2 |
| 6ef0 | C | 4.4 | 9042 | 0.03 | 1.03 | 244 | 5.7 | 5.7 | 0.805 | 0.815 | 79.5 | 42.3 | 75.8 | 21.6 |
| 6ef0 | D | 4.4 | 9042 | 0.03 | 1.03 | 242 | 20.9 | 19.2 | 0.569 | 0.569 | 81 | 34.7 | 74.8 | 36.5 |
| 6ef0 | E | 4.4 | 9042 | 0.03 | 1.03 | 249 | 6.6 | 6.6 | 0.780 | 0.780 | 77.9 | 45.9 | 69.1 | 22.7 |
| 6ef0 | F | 4.4 | 9042 | 0.03 | 1.03 | 234 | 16.0 | 16.0 | 0.575 | 0.580 | 75.2 | 30.1 | 80.3 | 7.4 |
| 6ef0 | H | 4.4 | 9042 | 0.03 | 1.03 | 257 | 21.4 | 11.8 | 0.256 | 0.700 | 71.2 | 4.9 | 52.1 | 6 |
| 6ef0 | I | 4.4 | 9042 | 0.03 | 1.03 | 271 | 21.7 | 15.8 | 0.353 | 0.451 | 66.8 | 7.2 | 51.3 | 6.5 |
| 6ef0 | J | 4.4 | 9042 | 0.03 | 1.03 | 272 | 21.7 | 13.1 | 0.581 | 0.647 | 76.1 | 11.1 | 80.1 | 4.6 |
| 6ef0 | K | 4.4 | 9042 | 0.03 | 1.03 | 272 | 19.7 | 17.8 | 0.353 | 0.430 | 75.7 | 5.8 | 76.1 | 6.3 |
| 6ef0 | L | 4.4 | 9042 | 0.03 | 1.03 | 273 | 22.0 | 16.6 | 0.267 | 0.528 | 75.8 | 4.8 | 68.5 | 6.4 |
| 6ef0 | M | 4.4 | 9042 | 0.03 | 1.03 | 258 | 25.6 | 17.9 | 0.279 | 0.360 | 60.5 | 9 | 55.8 | 5.6 |
| 6ef1 | A | 4.7 | 9043 | 0.035 | 1.03 | 239 | 9.2 | 9.2 | 0.784 | 0.784 | 82.8 | 39.4 | 56.5 | 5.2 |

| | | | | | | | | | | | | | | |
|---|---|---|---|---|---|---|---|---|---|---|---|---|---|---|
| 6ef1 | B | 4.7 | 9043 | 0.035 | 1.03 | 250 | 21.6 | 17.2 | 0.275 | 0.325 | 72.8 | 6.6 | 71.6 | 7.8 |
| 6ef1 | C | 4.7 | 9043 | 0.035 | 1.03 | 238 | 5.4 | 5.4 | 0.817 | 0.817 | 83.2 | 37.9 | 76.9 | 13.1 |
| 6ef1 | D | 4.7 | 9043 | 0.035 | 1.03 | 234 | 9.7 | 6.7 | 0.753 | 0.792 | 76.5 | 35.2 | 71.8 | 26.8 |
| 6ef1 | E | 4.7 | 9043 | 0.035 | 1.03 | 242 | 17.5 | 13.9 | 0.653 | 0.689 | 71.5 | 24.9 | 53.7 | 6.9 |
| 6ef1 | F | 4.7 | 9043 | 0.035 | 1.03 | 233 | 20.1 | 19.3 | 0.362 | 0.428 | 69.1 | 16.1 | 70.4 | 7.9 |
| 6ef1 | H | 4.7 | 9043 | 0.035 | 1.03 | 254 | 23.7 | 17.4 | 0.249 | 0.479 | 62.2 | 8.9 | 50 | 7.1 |
| 6ef1 | I | 4.7 | 9043 | 0.035 | 1.03 | 271 | 23.8 | 22.0 | 0.220 | 0.280 | 53.5 | 9.7 | 64.2 | 6.3 |
| 6ef1 | J | 4.7 | 9043 | 0.035 | 1.03 | 273 | 20.1 | 19.0 | 0.401 | 0.505 | 62.3 | 7.1 | 62.3 | 10 |
| 6ef1 | K | 4.7 | 9043 | 0.035 | 1.03 | 276 | 17.7 | 16.4 | 0.448 | 0.498 | 62.7 | 12.1 | 63.8 | 5.7 |
| 6ef1 | L | 4.7 | 9043 | 0.035 | 1.03 | 264 | 21.9 | 19.7 | 0.231 | 0.263 | 63.6 | 7.1 | 32.2 | 11.8 |
| 6ef1 | M | 4.7 | 9043 | 0.035 | 1.03 | 262 | 20.5 | 18.4 | 0.220 | 0.399 | 66.8 | 8 | 39.3 | 10.7 |
| 6ef2 | A | 4.3 | 9044 | 0.04 | 1.03 | 238 | 15.2 | 14.4 | 0.462 | 0.666 | 78.6 | 23.5 | 85.3 | 20.7 |
| 6ef2 | B | 4.3 | 9044 | 0.04 | 1.03 | 249 | 12.5 | 12.5 | 0.702 | 0.702 | 78.3 | 33.3 | 78.3 | 42.6 |
| 6ef2 | C | 4.3 | 9044 | 0.04 | 1.03 | 241 | 6.0 | 6.0 | 0.830 | 0.833 | 82.2 | 61.1 | 83 | 26.5 |
| 6ef2 | D | 4.3 | 9044 | 0.04 | 1.03 | 241 | 17.9 | 17.0 | 0.483 | 0.560 | 79.7 | 28.1 | 85.9 | 32.9 |
| 6ef2 | E | 4.3 | 9044 | 0.04 | 1.03 | 247 | 15.2 | 6.5 | 0.627 | 0.806 | 81 | 31 | 91.5 | 22.6 |
| 6ef2 | F | 4.3 | 9044 | 0.04 | 1.03 | 232 | 16.1 | 16.1 | 0.301 | 0.484 | 81.9 | 10.5 | 88.8 | 5.8 |
| 6ef2 | H | 4.3 | 9044 | 0.04 | 1.03 | 265 | 19.8 | 15.2 | 0.455 | 0.485 | 66.8 | 16.4 | 27.5 | 4.1 |
| 6ef2 | I | 4.3 | 9044 | 0.04 | 1.03 | 260 | 10.7 | 9.4 | 0.704 | 0.704 | 81.2 | 43.6 | 35.8 | 9.7 |
| 6ef2 | J | 4.3 | 9044 | 0.04 | 1.03 | 262 | 9.6 | 8.2 | 0.743 | 0.790 | 84.4 | 50.2 | 82.1 | 17.2 |
| 6ef2 | K | 4.3 | 9044 | 0.04 | 1.03 | 259 | 16.8 | 15.4 | 0.434 | 0.471 | 81.1 | 11 | 79.2 | 12.7 |
| 6ef2 | L | 4.3 | 9044 | 0.04 | 1.03 | 264 | 21.9 | 20.1 | 0.286 | 0.379 | 69.7 | 8.2 | 74.6 | 9.1 |
| 6ef2 | M | 4.3 | 9044 | 0.04 | 1.03 | 270 | 15.2 | 14.6 | 0.577 | 0.577 | 76.3 | 35.4 | 45.9 | 6.5 |
| 6ef3 | 4 | 4.2 | 9045 | 0.045 | 1.03 | 195 | 3.6 | 3.6 | 0.867 | 0.867 | 88.7 | 68.2 | 83.1 | 3.7 |
| 6ef3 | A | 4.2 | 9045 | 0.045 | 1.03 | 247 | 6.9 | 6.4 | 0.817 | 0.820 | 82.2 | 49.8 | 85.4 | 42.7 |
| 6ef3 | B | 4.2 | 9045 | 0.045 | 1.03 | 250 | 12.3 | 6.0 | 0.697 | 0.823 | 82.8 | 48.3 | 86.4 | 17.6 |
| 6ef3 | C | 4.2 | 9045 | 0.045 | 1.03 | 242 | 20.6 | 16.8 | 0.587 | 0.640 | 78.5 | 36.3 | 82.6 | 6.5 |
| 6ef3 | D | 4.2 | 9045 | 0.045 | 1.03 | 241 | 14.0 | 13.1 | 0.694 | 0.694 | 80.1 | 47.7 | 91.7 | 32.1 |
| 6ef3 | E | 4.2 | 9045 | 0.045 | 1.03 | 249 | 10.5 | 7.6 | 0.619 | 0.763 | 78.3 | 41.5 | 84.3 | 13.3 |
| 6ef3 | F | 4.2 | 9045 | 0.045 | 1.03 | 234 | 16.4 | 16.4 | 0.622 | 0.626 | 82.5 | 30.1 | 82.1 | 6.2 |
| 6ef3 | r | 4.2 | 9045 | 0.045 | 1.03 | 280 | 15.5 | 13.7 | 0.505 | 0.654 | 80.4 | 28.9 | 82.9 | 10.8 |
| 6ef3 | u | 4.2 | 9045 | 0.045 | 1.03 | 75 | 12.6 | 9.7 | 0.333 | 0.482 | 72 | 1.9 | 54.7 | 2.4 |
| 6eml | P | 3.6 | 3886 | 0.0722 | 1.084 | 206 | 6.2 | 6.2 | 0.833 | 0.833 | 85 | 43.4 | 85.9 | 16.9 |
| 6et5 | H | 2.9 | 3951 | 0.0156 | 1.06 | 257 | 3.8 | 3.8 | 0.888 | 0.888 | 89.5 | 73 | 80.9 | 20.2 |
| 6et5 | L | 2.9 | 3951 | 0.0156 | 1.06 | 273 | 2.3 | 2.3 | 0.940 | 0.940 | 95.6 | 75.1 | 91.9 | 62.9 |
| 6eti | C | 3.1 | 3953 | 0.038 | 0.84 | 107 | 2.1 | 2.1 | 0.912 | 0.912 | 96.3 | 78.6 | 79.4 | 40 |
| 6eti | D | 3.1 | 3953 | 0.038 | 0.84 | 118 | 1.9 | 1.9 | 0.936 | 0.936 | 97.5 | 86.1 | 93.2 | 19.1 |

| | | | | | | | | | | | | | | |
|---|---|---|---|---|---|---|---|---|---|---|---|---|---|---|
| 6eti | E | 3.1 | 3953 | 0.038 | 0.84 | 107 | 1.5 | 1.5 | 0.930 | 0.930 | 97.2 | 87.5 | 69.2 | 6.8 |
| 6eti | F | 3.1 | 3953 | 0.038 | 0.84 | 118 | 1.9 | 1.9 | 0.920 | 0.920 | 94.9 | 83 | 94.9 | 6.2 |
| 6eu0 | E | 4 | 3955 | 0.02 | 1.06 | 215 | 14.7 | 11.8 | 0.571 | 0.571 | 73 | 27.4 | 63.7 | 18.2 |
| 6exn | a | 3.7 | 3979 | 0.04 | 1.12 | 171 | 9.2 | 4.7 | 0.795 | 0.818 | 84.2 | 41 | 82.5 | 22 |
| 6exn | L | 3.7 | 3979 | 0.04 | 1.12 | 156 | 2.9 | 2.9 | 0.892 | 0.892 | 91.7 | 74.8 | 74.4 | 50 |
| 6exn | M | 3.7 | 3979 | 0.04 | 1.12 | 255 | 13.8 | 6.9 | 0.647 | 0.741 | 80 | 38.2 | 69 | 3.4 |
| 6exv | E | 3.6 | 3981 | 0.0577 | 1.07 | 209 | 3.6 | 3.6 | 0.894 | 0.894 | 91.4 | 73.3 | 82.8 | 49.1 |
| 6eyd | A | 4.2 | 3983 | 0.1 | 1.34 | 220 | 28.5 | 26.0 | 0.311 | 0.384 | 62.3 | 8 | 42.3 | 8.6 |
| 6eyd | B | 4.2 | 3983 | 0.1 | 1.34 | 233 | 27.8 | 26.3 | 0.247 | 0.287 | 68.7 | 7.5 | 51.9 | 7.4 |
| 6eyd | F | 4.2 | 3983 | 0.1 | 1.34 | 217 | 27.6 | 13.4 | 0.268 | 0.587 | 61.8 | 10.4 | 43.8 | 10.5 |
| 6ezm | A | 3.2 | 4160 | 0.116 | 1.065 | 212 | 9.9 | 8.2 | 0.808 | 0.825 | 81.6 | 45.1 | 76.4 | 57.4 |
| 6ezm | B | 3.2 | 4160 | 0.116 | 1.065 | 212 | 6.8 | 6.0 | 0.876 | 0.877 | 88.2 | 79.1 | 81.1 | 63.4 |
| 6ezm | C | 3.2 | 4160 | 0.116 | 1.065 | 212 | 5.8 | 5.8 | 0.892 | 0.892 | 91.5 | 84 | 84.9 | 47.8 |
| 6ezm | D | 3.2 | 4160 | 0.116 | 1.065 | 212 | 7.9 | 5.8 | 0.805 | 0.872 | 89.6 | 62.6 | 84.4 | 82.1 |
| 6ezm | E | 3.2 | 4160 | 0.116 | 1.065 | 212 | 5.1 | 5.1 | 0.892 | 0.892 | 91 | 58.5 | 82.1 | 43.7 |
| 6ezm | F | 3.2 | 4160 | 0.116 | 1.065 | 212 | 11.7 | 7.7 | 0.781 | 0.781 | 91.5 | 61.3 | 82.1 | 64.4 |
| 6ezm | G | 3.2 | 4160 | 0.116 | 1.065 | 212 | 6.7 | 6.7 | 0.860 | 0.860 | 87.3 | 64.9 | 83.5 | 76.8 |
| 6ezm | H | 3.2 | 4160 | 0.116 | 1.065 | 212 | 5.2 | 4.8 | 0.816 | 0.891 | 90.6 | 68.2 | 85.4 | 69.1 |
| 6ezm | I | 3.2 | 4160 | 0.116 | 1.065 | 212 | 6.9 | 5.7 | 0.798 | 0.815 | 89.6 | 54.7 | 81.1 | 51.2 |
| 6ezm | J | 3.2 | 4160 | 0.116 | 1.065 | 212 | 7.0 | 7.0 | 0.874 | 0.884 | 89.2 | 74.1 | 75.9 | 72 |
| 6ezm | K | 3.2 | 4160 | 0.116 | 1.065 | 212 | 9.7 | 6.1 | 0.823 | 0.842 | 86.3 | 38.8 | 82.1 | 66.7 |
| 6ezm | L | 3.2 | 4160 | 0.116 | 1.065 | 212 | 6.9 | 5.6 | 0.803 | 0.819 | 89.6 | 42.6 | 69.8 | 79.7 |
| 6ezm | M | 3.2 | 4160 | 0.116 | 1.065 | 212 | 9.3 | 7.9 | 0.771 | 0.785 | 85.4 | 60.8 | 78.8 | 66.5 |
| 6ezm | N | 3.2 | 4160 | 0.116 | 1.065 | 212 | 5.9 | 5.9 | 0.889 | 0.889 | 91.5 | 70.6 | 74.1 | 59.9 |
| 6ezm | O | 3.2 | 4160 | 0.116 | 1.065 | 212 | 8.0 | 5.8 | 0.810 | 0.810 | 88.7 | 43.1 | 80.7 | 64.9 |
| 6ezm | P | 3.2 | 4160 | 0.116 | 1.065 | 212 | 7.3 | 5.9 | 0.873 | 0.873 | 89.6 | 64.7 | 80.2 | 57.6 |
| 6ezm | Q | 3.2 | 4160 | 0.116 | 1.065 | 212 | 7.2 | 5.3 | 0.851 | 0.851 | 90.6 | 67.7 | 80.2 | 84.7 |
| 6ezm | R | 3.2 | 4160 | 0.116 | 1.065 | 212 | 7.5 | 7.2 | 0.849 | 0.854 | 90.6 | 70.8 | 76.4 | 60.5 |
| 6ezm | S | 3.2 | 4160 | 0.116 | 1.065 | 212 | 6.8 | 6.8 | 0.868 | 0.868 | 90.1 | 63.9 | 78.8 | 85 |
| 6ezm | T | 3.2 | 4160 | 0.116 | 1.065 | 212 | 6.7 | 6.7 | 0.860 | 0.860 | 86.3 | 66.7 | 79.7 | 59.2 |
| 6ezm | U | 3.2 | 4160 | 0.116 | 1.065 | 212 | 4.9 | 4.9 | 0.832 | 0.839 | 90.6 | 63 | 78.8 | 47.9 |
| 6ezm | V | 3.2 | 4160 | 0.116 | 1.065 | 212 | 8.2 | 5.8 | 0.807 | 0.866 | 88.7 | 53.7 | 81.1 | 62.8 |
| 6ezm | W | 3.2 | 4160 | 0.116 | 1.065 | 212 | 7.6 | 5.3 | 0.837 | 0.869 | 90.1 | 77 | 87.3 | 69.7 |
| 6ezm | X | 3.2 | 4160 | 0.116 | 1.065 | 212 | 7.5 | 6.5 | 0.865 | 0.871 | 88.7 | 55.3 | 73.6 | 41 |
| 6f0x | F | 4.6 | 4166 | 0.0765 | 1.43 | 282 | 19.7 | 17.5 | 0.379 | 0.423 | 67.4 | 4.7 | 33.7 | 12.6 |
| 6f0x | P | 4.6 | 4166 | 0.0765 | 1.43 | 194 | 17.3 | 15.6 | 0.378 | 0.378 | 68.6 | 7.5 | 70.6 | 5.8 |
| 6f1t | K | 3.5 | 4168 | 0.046 | 1.34 | 278 | 14.3 | 14.3 | 0.645 | 0.645 | 74.8 | 14.9 | 50 | 3.6 |

| | | | | | | | | | | | | | | |
|---|---|---|---|---|---|---|---|---|---|---|---|---|---|---|
| 6f1t | L | 3.5 | 4168 | 0.046 | 1.34 | 269 | 16.2 | 10.6 | 0.718 | 0.770 | 79.9 | 22.8 | 68.8 | 8.1 |
| 6f1u | K | 3.4 | 4169 | 0.09 | 1.34 | 278 | 15.8 | 9.7 | 0.623 | 0.715 | 75.9 | 20.9 | 58.6 | 6.7 |
| 6f1u | L | 3.4 | 4169 | 0.09 | 1.34 | 269 | 11.4 | 6.3 | 0.766 | 0.806 | 75.5 | 26.1 | 70.3 | 9.5 |
| 6f2d | H | 4.2 | 4173 | 0.07 | 0.86 | 89 | 10.0 | 5.0 | 0.370 | 0.768 | 87.6 | 6.4 | 92.1 | 46.3 |
| 6f40 | E | 3.7 | 4180 | 0.05 | 1.35 | 214 | 12.3 | 8.9 | 0.783 | 0.808 | 84.6 | 40.3 | 70.6 | 7.9 |
| 6f40 | F | 3.7 | 4180 | 0.05 | 1.35 | 83 | 1.3 | 1.3 | 0.895 | 0.895 | 96.4 | 97.5 | 80.7 | 49.3 |
| 6f40 | H | 3.7 | 4180 | 0.05 | 1.35 | 140 | 17.8 | 15.7 | 0.311 | 0.572 | 84.3 | 10.2 | 80.7 | 22.1 |
| 6f40 | K | 3.7 | 4180 | 0.05 | 1.35 | 101 | 1.3 | 1.3 | 0.930 | 0.930 | 98 | 94.9 | 92.1 | 48.4 |
| 6f40 | P | 3.7 | 4180 | 0.05 | 1.35 | 246 | 11.6 | 10.5 | 0.660 | 0.726 | 78.9 | 20.1 | 62.2 | 25.5 |
| 6f41 | E | 4.3 | 4181 | 0.045 | 1.35 | 214 | 8.5 | 8.5 | 0.705 | 0.705 | 76.6 | 45.7 | 65.4 | 4.3 |
| 6f41 | F | 4.3 | 4181 | 0.045 | 1.35 | 83 | 3.0 | 3.0 | 0.802 | 0.802 | 88 | 53.4 | 72.3 | 6.7 |
| 6f41 | H | 4.3 | 4181 | 0.045 | 1.35 | 140 | 14.8 | 14.8 | 0.237 | 0.399 | 72.1 | 5 | 53.6 | 4 |
| 6f41 | K | 4.3 | 4181 | 0.045 | 1.35 | 101 | 7.0 | 7.0 | 0.712 | 0.712 | 84.2 | 51.8 | 86.1 | 35.6 |
| 6f41 | P | 4.3 | 4181 | 0.045 | 1.35 | 246 | 9.8 | 8.3 | 0.726 | 0.759 | 76.4 | 26.1 | 25.2 | 12.9 |
| 6f44 | E | 4.2 | 4183 | 0.045 | 1.35 | 214 | 17.1 | 12.0 | 0.565 | 0.646 | 69.6 | 30.2 | 33.2 | 2.8 |
| 6f44 | F | 4.2 | 4183 | 0.045 | 1.35 | 83 | 3.1 | 3.1 | 0.800 | 0.800 | 89.2 | 71.6 | 43.4 | 11.1 |
| 6f44 | G | 4.2 | 4183 | 0.045 | 1.35 | 180 | 21.0 | 17.4 | 0.240 | 0.250 | 60 | 9.3 | 16.1 | 3.4 |
| 6f44 | H | 4.2 | 4183 | 0.045 | 1.35 | 140 | 16.4 | 14.9 | 0.223 | 0.379 | 70.7 | 6.1 | 60.7 | 2.4 |
| 6f44 | K | 4.2 | 4183 | 0.045 | 1.35 | 101 | 12.7 | 8.9 | 0.452 | 0.677 | 86.1 | 6.9 | 79.2 | 2.5 |
| 6f6w | A | 3.8 | 4192 | 0.1 | 1.34 | 221 | 25.0 | 18.6 | 0.387 | 0.399 | 67.9 | 8 | 46.6 | 4.9 |
| 6f6w | B | 3.8 | 4192 | 0.1 | 1.34 | 233 | 19.5 | 17.2 | 0.392 | 0.392 | 71.2 | 7.8 | 51.1 | 11.8 |
| 6fai | A | 3.4 | 4214 | 0.01 | 1.39 | 206 | 5.9 | 4.8 | 0.852 | 0.852 | 87.4 | 53.3 | 77.2 | 35.2 |
| 6fai | B | 3.4 | 4214 | 0.01 | 1.39 | 214 | 10.2 | 10.2 | 0.632 | 0.632 | 79.9 | 29.8 | 62.1 | 24.8 |
| 6fai | C | 3.4 | 4214 | 0.01 | 1.39 | 217 | 3.3 | 3.3 | 0.898 | 0.898 | 92.2 | 57 | 76.5 | 14.5 |
| 6fai | E | 3.4 | 4214 | 0.01 | 1.39 | 260 | 9.1 | 4.7 | 0.840 | 0.896 | 88.5 | 33.9 | 72.3 | 6.4 |
| 6fai | h | 3.4 | 4214 | 0.01 | 1.39 | 181 | 6.5 | 6.5 | 0.753 | 0.753 | 82.3 | 52.3 | 67.4 | 18.9 |
| 6fai | H | 3.4 | 4214 | 0.01 | 1.39 | 184 | 11.1 | 10.2 | 0.563 | 0.636 | 84.2 | 16.1 | 64.1 | 21.2 |
| 6fai | J | 3.4 | 4214 | 0.01 | 1.39 | 185 | 3.2 | 3.2 | 0.901 | 0.901 | 92.4 | 47.4 | 79.5 | 46.3 |
| 6fai | L | 3.4 | 4214 | 0.01 | 1.39 | 140 | 3.5 | 3.5 | 0.841 | 0.853 | 87.1 | 59.8 | 77.9 | 5.5 |
| 6fai | N | 3.4 | 4214 | 0.01 | 1.39 | 150 | 2.5 | 2.5 | 0.894 | 0.894 | 92.7 | 67.6 | 81.3 | 55.7 |
| 6fai | V | 3.4 | 4214 | 0.01 | 1.39 | 87 | 3.5 | 3.5 | 0.794 | 0.794 | 87.4 | 61.8 | 40.2 | 8.6 |
| 6fai | W | 3.4 | 4214 | 0.01 | 1.39 | 129 | 6.2 | 6.2 | 0.869 | 0.869 | 92.2 | 63 | 91.5 | 8.5 |
| 6fai | Y | 3.4 | 4214 | 0.01 | 1.39 | 134 | 3.3 | 3.3 | 0.870 | 0.873 | 91 | 62.3 | 70.1 | 9.6 |
| 6fbv | A | 3.5 | 4230 | 0.03 | 1.061 | 223 | 2.1 | 2.1 | 0.925 | 0.925 | 93.7 | 81.3 | 85.2 | 34.7 |
| 6fbv | E | 3.5 | 4230 | 0.03 | 1.061 | 83 | 1.9 | 1.9 | 0.881 | 0.881 | 96.4 | 86.2 | 84.3 | 40 |
| 6feq | C | 3.6 | 4246 | 0.51 | 1.065 | 107 | 2.1 | 2.1 | 0.866 | 0.866 | 94.4 | 78.2 | 64.5 | 10.1 |
| 6feq | D | 3.6 | 4246 | 0.51 | 1.065 | 118 | 13.7 | 11.3 | 0.436 | 0.535 | 82.2 | 15.5 | 63.6 | 6.7 |

| | | | | | | | | | | | | | | |
|---|---|---|---|---|---|---|---|---|---|---|---|---|---|---|
| 6feq | E | 3.6 | 4246 | 0.51 | 1.065 | 107 | 11.7 | 9.2 | 0.674 | 0.690 | 84.1 | 18.9 | 79.4 | 4.7 |
| 6feq | F | 3.6 | 4246 | 0.51 | 1.065 | 118 | 13.9 | 13.6 | 0.426 | 0.448 | 72.9 | 23.3 | 44.1 | 7.7 |
| 6ff4 | 1 | 3.4 | 4255 | 0.04 | 1.16 | 122 | 2.8 | 2.8 | 0.899 | 0.899 | 95.1 | 76.7 | 82 | 35 |
| 6ff4 | L | 3.4 | 4255 | 0.04 | 1.16 | 103 | 1.9 | 1.9 | 0.903 | 0.903 | 96.1 | 87.9 | 89.3 | 40.2 |
| 6ff4 | O | 3.4 | 4255 | 0.04 | 1.16 | 253 | 7.1 | 4.3 | 0.761 | 0.837 | 79.8 | 61.4 | 71.5 | 13.3 |
| 6ff4 | Q | 3.4 | 4255 | 0.04 | 1.16 | 138 | 2.1 | 2.1 | 0.913 | 0.913 | 96.4 | 85 | 71 | 45.9 |
| 6ff4 | s | 3.4 | 4255 | 0.04 | 1.16 | 175 | 3.4 | 3.4 | 0.887 | 0.887 | 90.3 | 58.2 | 73.7 | 14 |
| 6ff4 | y | 3.4 | 4255 | 0.04 | 1.16 | 100 | 2.3 | 2.3 | 0.894 | 0.894 | 93 | 57 | 69 | 7.2 |
| 6ff4 | Y | 3.4 | 4255 | 0.04 | 1.16 | 95 | 11.1 | 3.3 | 0.416 | 0.788 | 89.5 | 7.1 | 72.6 | 5.8 |
| 6flp | A | 4.1 | 4274 | 0.27 | 1.1 | 229 | 15.8 | 12.5 | 0.665 | 0.684 | 77.7 | 12.9 | 74.2 | 6.5 |
| 6flp | B | 4.1 | 4274 | 0.27 | 1.1 | 217 | 26.3 | 17.8 | 0.346 | 0.527 | 72.8 | 10.1 | 59 | 8.6 |
| 6flp | E | 4.1 | 4274 | 0.27 | 1.1 | 90 | 4.4 | 4.4 | 0.744 | 0.744 | 84.4 | 50 | 35.6 | 3.1 |
| 6fn1 | B | 3.6 | 4281 | 0.025 | 0.84 | 220 | 14.9 | 14.5 | 0.636 | 0.636 | 81.4 | 23.5 | 45.9 | 11.9 |
| 6fn1 | C | 3.6 | 4281 | 0.025 | 0.84 | 225 | 6.0 | 6.0 | 0.855 | 0.855 | 86.7 | 62.1 | 70.7 | 11.3 |
| 6fn4 | B | 4.1 | 4282 | 0.07 | 1.387 | 220 | 19.8 | 17.7 | 0.291 | 0.291 | 74.5 | 5.5 | 40.5 | 12.4 |
| 6fn4 | C | 4.1 | 4282 | 0.07 | 1.387 | 225 | 11.3 | 10.6 | 0.730 | 0.730 | 78.7 | 51.4 | 69.3 | 7.7 |
| 6fo0 | D | 4.1 | 4286 | 0.114 | 1.065 | 239 | 3.8 | 3.8 | 0.865 | 0.865 | 89.5 | 51.9 | 54.8 | 6.9 |
| 6fo0 | F | 4.1 | 4286 | 0.114 | 1.065 | 98 | 2.6 | 2.6 | 0.846 | 0.846 | 91.8 | 74.4 | 88.8 | 8 |
| 6fo0 | Q | 4.1 | 4286 | 0.114 | 1.065 | 239 | 4.8 | 4.8 | 0.866 | 0.866 | 88.7 | 33.5 | 35.6 | 7.1 |
| 6fo0 | S | 4.1 | 4286 | 0.114 | 1.065 | 98 | 2.4 | 2.4 | 0.884 | 0.884 | 94.9 | 60.2 | 89.8 | 60.2 |
| 6fo0 | U | 4.1 | 4286 | 0.114 | 1.065 | 65 | 3.8 | 3.8 | 0.719 | 0.719 | 86.2 | 30.4 | 90.8 | 84.7 |
| 6fo2 | D | 4.4 | 4288 | 0.0548 | 1.047 | 237 | 6.6 | 3.2 | 0.827 | 0.865 | 85.7 | 58.1 | 56.1 | 3 |
| 6fo2 | F | 4.4 | 4288 | 0.0548 | 1.047 | 98 | 2.7 | 2.7 | 0.841 | 0.841 | 92.9 | 64.8 | 87.8 | 39.5 |
| 6fo2 | H | 4.4 | 4288 | 0.0548 | 1.047 | 65 | 2.8 | 2.8 | 0.697 | 0.702 | 86.2 | 53.6 | 46.2 | 16.7 |
| 6fo2 | Q | 4.4 | 4288 | 0.0548 | 1.047 | 237 | 6.7 | 6.7 | 0.843 | 0.843 | 86.5 | 41 | 57.4 | 7.4 |
| 6fo2 | S | 4.4 | 4288 | 0.0548 | 1.047 | 98 | 2.2 | 2.2 | 0.855 | 0.855 | 92.9 | 83.5 | 46.9 | 8.7 |
| 6fo2 | U | 4.4 | 4288 | 0.0548 | 1.047 | 65 | 3.3 | 3.3 | 0.686 | 0.686 | 87.7 | 52.6 | 87.7 | 1.8 |
| 6fo6 | D | 4.1 | 4292 | 0.173 | 1.067 | 237 | 3.4 | 3.4 | 0.887 | 0.887 | 90.3 | 52.8 | 66.2 | 26.8 |
| 6fo6 | F | 4.1 | 4292 | 0.173 | 1.067 | 98 | 2.6 | 2.6 | 0.858 | 0.858 | 92.9 | 75.8 | 93.9 | 39.1 |
| 6fo6 | H | 4.1 | 4292 | 0.173 | 1.067 | 65 | 2.9 | 2.9 | 0.779 | 0.779 | 89.2 | 41.4 | 96.9 | 98.4 |
| 6fo6 | Q | 4.1 | 4292 | 0.173 | 1.067 | 237 | 6.1 | 3.3 | 0.873 | 0.885 | 88.6 | 47.1 | 63.7 | 28.5 |
| 6fo6 | S | 4.1 | 4292 | 0.173 | 1.067 | 98 | 2.1 | 2.1 | 0.900 | 0.900 | 94.9 | 81.7 | 93.9 | 7.6 |
| 6fo6 | U | 4.1 | 4292 | 0.173 | 1.067 | 65 | 4.3 | 4.3 | 0.762 | 0.762 | 89.2 | 32.8 | 92.3 | 6.7 |
| 6g18 | A | 3.6 | 4337 | 0.0583 | 1.084 | 216 | 4.5 | 4.5 | 0.857 | 0.889 | 92.1 | 66.3 | 90.7 | 58.2 |
| 6g18 | B | 3.6 | 4337 | 0.0583 | 1.084 | 213 | 2.2 | 2.2 | 0.925 | 0.925 | 94.4 | 69.2 | 83.1 | 33.9 |
| 6g18 | C | 3.6 | 4337 | 0.0583 | 1.084 | 218 | 2.4 | 2.4 | 0.930 | 0.934 | 94.5 | 71.4 | 61 | 42.9 |
| 6g18 | E | 3.6 | 4337 | 0.0583 | 1.084 | 262 | 2.2 | 2.2 | 0.939 | 0.939 | 95 | 82.7 | 85.1 | 11.7 |

| | | | | | | | | | | | | | | |
|---|---|---|---|---|---|---|---|---|---|---|---|---|---|---|
| 6g18 | H | 3.6 | 4337 | 0.0583 | 1.084 | 186 | 4.3 | 4.3 | 0.875 | 0.875 | 90.3 | 47.6 | 77.4 | 6.9 |
| 6g18 | w | 3.6 | 4337 | 0.0583 | 1.084 | 249 | 3.4 | 3.4 | 0.868 | 0.868 | 88.4 | 47.7 | 80.7 | 27.4 |
| 6g18 | x | 3.6 | 4337 | 0.0583 | 1.084 | 178 | 1.7 | 1.7 | 0.936 | 0.936 | 95.5 | 85.9 | 88.2 | 68.8 |
| 6g18 | Y | 3.6 | 4337 | 0.0583 | 1.084 | 124 | 1.5 | 1.5 | 0.925 | 0.925 | 95.2 | 87.3 | 87.9 | 4.6 |
| 6g2j | E | 3.3 | 4345 | 0.075 | 1.05 | 212 | 2.3 | 2.3 | 0.913 | 0.913 | 94.3 | 76.5 | 84.9 | 37.2 |
| 6g2j | S | 3.3 | 4345 | 0.075 | 1.05 | 83 | 12.4 | 8.5 | 0.505 | 0.722 | 91.6 | 17.1 | 91.6 | 65.8 |
| 6g4s | A | 4 | 4348 | 0.05 | 1.084 | 216 | 3.0 | 3.0 | 0.900 | 0.900 | 91.2 | 56.3 | 47.2 | 6.9 |
| 6g4s | B | 4 | 4348 | 0.05 | 1.084 | 213 | 11.0 | 11.0 | 0.779 | 0.779 | 80.3 | 25.7 | 72.3 | 5.2 |
| 6g4s | C | 4 | 4348 | 0.05 | 1.084 | 218 | 3.7 | 3.7 | 0.867 | 0.875 | 89.9 | 43.9 | 72 | 6.4 |
| 6g4s | E | 4 | 4348 | 0.05 | 1.084 | 262 | 4.6 | 4.6 | 0.884 | 0.884 | 89.7 | 36.2 | 59.2 | 5.8 |
| 6g4s | F | 4 | 4348 | 0.05 | 1.084 | 189 | 11.3 | 6.0 | 0.673 | 0.760 | 83.1 | 21 | 55.6 | 1.9 |
| 6g4s | G | 4 | 4348 | 0.05 | 1.084 | 230 | 17.6 | 17.4 | 0.563 | 0.571 | 75.7 | 20.1 | 64.8 | 9.4 |
| 6g4s | H | 4 | 4348 | 0.05 | 1.084 | 186 | 6.8 | 6.3 | 0.764 | 0.769 | 78.5 | 51.4 | 70.4 | 6.1 |
| 6g4s | I | 4 | 4348 | 0.05 | 1.084 | 205 | 7.4 | 7.4 | 0.763 | 0.763 | 80.5 | 33.3 | 53.7 | 8.2 |
| 6g4s | J | 4 | 4348 | 0.05 | 1.084 | 180 | 3.2 | 3.2 | 0.875 | 0.875 | 90.6 | 52.8 | 53.3 | 8.3 |
| 6g4s | L | 4 | 4348 | 0.05 | 1.084 | 151 | 6.7 | 5.1 | 0.827 | 0.837 | 86.1 | 30.8 | 45 | 7.4 |
| 6g4s | N | 4 | 4348 | 0.05 | 1.084 | 149 | 2.4 | 2.4 | 0.852 | 0.852 | 90.6 | 69.6 | 26.2 | 5.1 |
| 6g4s | R | 4 | 4348 | 0.05 | 1.084 | 120 | 12.8 | 5.8 | 0.509 | 0.704 | 69.2 | 10.8 | 59.2 | 5.6 |
| 6g4s | V | 4 | 4348 | 0.05 | 1.084 | 82 | 7.3 | 3.9 | 0.725 | 0.738 | 85.4 | 27.1 | 74.4 | 4.9 |
| 6g4s | W | 4 | 4348 | 0.05 | 1.084 | 129 | 10.2 | 5.7 | 0.658 | 0.796 | 86 | 36 | 82.9 | 19.6 |
| 6g4s | x | 4 | 4348 | 0.05 | 1.084 | 175 | 8.9 | 4.8 | 0.620 | 0.828 | 89.7 | 42 | 86.9 | 6.6 |
| 6g4s | X | 4 | 4348 | 0.05 | 1.084 | 141 | 13.6 | 11.4 | 0.772 | 0.772 | 78 | 13.6 | 85.1 | 7.5 |
| 6g4s | Y | 4 | 4348 | 0.05 | 1.084 | 124 | 6.0 | 4.1 | 0.725 | 0.820 | 87.9 | 59.6 | 57.3 | 4.2 |
| 6g4w | B | 4.5 | 4349 | 0.04 | 1.084 | 213 | 21.7 | 16.3 | 0.501 | 0.501 | 68.5 | 9.6 | 31.5 | 7.5 |
| 6g4w | F | 4.5 | 4349 | 0.04 | 1.084 | 189 | 13.4 | 8.9 | 0.677 | 0.705 | 75.1 | 6.3 | 56.1 | 1.9 |
| 6g4w | H | 4.5 | 4349 | 0.04 | 1.084 | 186 | 17.8 | 14.2 | 0.278 | 0.520 | 64.5 | 10.8 | 66.1 | 4.9 |
| 6g4w | x | 4.5 | 4349 | 0.04 | 1.084 | 175 | 8.6 | 8.6 | 0.673 | 0.691 | 74.9 | 19.1 | 72.6 | 8.7 |
| 6g51 | A | 4.1 | 4350 | 0.04 | 1.084 | 216 | 11.3 | 10.7 | 0.770 | 0.770 | 86.6 | 32.1 | 72.7 | 7 |
| 6g51 | B | 4.1 | 4350 | 0.04 | 1.084 | 213 | 21.5 | 18.1 | 0.275 | 0.502 | 71.8 | 6.5 | 46.9 | 5 |
| 6g51 | C | 4.1 | 4350 | 0.04 | 1.084 | 218 | 12.9 | 12.9 | 0.769 | 0.769 | 78.4 | 39.8 | 71.6 | 5.1 |
| 6g51 | E | 4.1 | 4350 | 0.04 | 1.084 | 262 | 14.8 | 14.3 | 0.563 | 0.563 | 67.6 | 11.9 | 56.1 | 3.4 |
| 6g51 | H | 4.1 | 4350 | 0.04 | 1.084 | 186 | 14.6 | 14.6 | 0.380 | 0.390 | 66.7 | 10.5 | 61.8 | 7 |
| 6g51 | J | 4.1 | 4350 | 0.04 | 1.084 | 180 | 6.4 | 5.0 | 0.793 | 0.809 | 81.7 | 36.7 | 68.3 | 9.8 |
| 6g51 | x | 4.1 | 4350 | 0.04 | 1.084 | 178 | 5.3 | 5.3 | 0.793 | 0.808 | 82 | 36.3 | 76.4 | 29.4 |
| 6g51 | Y | 4.1 | 4350 | 0.04 | 1.084 | 124 | 10.3 | 10.2 | 0.657 | 0.657 | 76.6 | 23.2 | 64.5 | 6.2 |
| 6g53 | A | 4.5 | 4351 | 0.04 | 1.084 | 216 | 9.5 | 9.5 | 0.671 | 0.671 | 75 | 9.9 | 66.2 | 7.7 |
| 6g53 | B | 4.5 | 4351 | 0.04 | 1.084 | 213 | 20.3 | 18.1 | 0.253 | 0.299 | 62.9 | 10.4 | 34.3 | 5.5 |

| | | | | | | | | | | | | | | |
|---|---|---|---|---|---|---|---|---|---|---|---|---|---|---|
| 6g53 | C | 4.5 | 4351 | 0.04 | 1.084 | 218 | 13.5 | 13.5 | 0.449 | 0.459 | 67.4 | 4.1 | 48.6 | 13.2 |
| 6g53 | H | 4.5 | 4351 | 0.04 | 1.084 | 186 | 17.1 | 15.9 | 0.274 | 0.353 | 65.1 | 8.3 | 53.8 | 8 |
| 6g53 | x | 4.5 | 4351 | 0.04 | 1.084 | 178 | 13.9 | 12.7 | 0.330 | 0.444 | 68 | 10.7 | 55.6 | 6.1 |
| 6g5h | A | 3.6 | 4352 | 0.06 | 1.084 | 206 | 9.5 | 6.2 | 0.745 | 0.802 | 84.5 | 52.3 | 72.3 | 32.2 |
| 6g5h | B | 3.6 | 4352 | 0.06 | 1.084 | 213 | 18.0 | 11.7 | 0.655 | 0.724 | 75.1 | 18.8 | 54.9 | 7.7 |
| 6g5h | C | 3.6 | 4352 | 0.06 | 1.084 | 218 | 3.4 | 3.4 | 0.902 | 0.902 | 92.7 | 63.4 | 85.3 | 13.4 |
| 6g5h | E | 3.6 | 4352 | 0.06 | 1.084 | 262 | 5.8 | 4.4 | 0.815 | 0.883 | 89.7 | 44.3 | 69.8 | 7.1 |
| 6g5h | Y | 3.6 | 4352 | 0.06 | 1.084 | 124 | 3.1 | 3.1 | 0.860 | 0.860 | 88.7 | 68.2 | 84.7 | 21.9 |
| 6g5i | A | 3.5 | 4353 | 0.03 | 1.084 | 216 | 2.5 | 2.5 | 0.915 | 0.915 | 93.5 | 69.3 | 83.3 | 6.7 |
| 6g5i | B | 3.5 | 4353 | 0.03 | 1.084 | 213 | 14.4 | 10.1 | 0.651 | 0.680 | 83.1 | 44.1 | 69 | 32 |
| 6g5i | C | 3.5 | 4353 | 0.03 | 1.084 | 218 | 7.4 | 7.4 | 0.829 | 0.829 | 86.7 | 46.6 | 77.1 | 10.1 |
| 6g5i | E | 3.5 | 4353 | 0.03 | 1.084 | 262 | 3.5 | 3.5 | 0.895 | 0.895 | 90.1 | 61.4 | 31.3 | 4.9 |
| 6g5i | x | 3.5 | 4353 | 0.03 | 1.084 | 175 | 9.3 | 7.2 | 0.698 | 0.806 | 87.4 | 30.1 | 54.3 | 7.4 |
| 6g5i | Y | 3.5 | 4353 | 0.03 | 1.084 | 124 | 8.5 | 8.5 | 0.679 | 0.743 | 84.7 | 36.2 | 65.3 | 11.1 |
| 6g72 | E | 3.9 | 4356 | 0.075 | 1.33 | 212 | 13.0 | 13.0 | 0.655 | 0.669 | 80.7 | 16.4 | 78.3 | 4.8 |
| 6g72 | P | 3.9 | 4356 | 0.075 | 1.33 | 291 | 15.4 | 11.2 | 0.655 | 0.711 | 81.1 | 38.1 | 80.4 | 10.7 |
| 6g72 | R | 3.9 | 4356 | 0.075 | 1.33 | 95 | 6.7 | 4.0 | 0.734 | 0.781 | 82.1 | 48.7 | 55.8 | 5.7 |
| 6g72 | S | 3.9 | 4356 | 0.075 | 1.33 | 83 | 13.8 | 12.0 | 0.280 | 0.445 | 74.7 | 3.2 | 67.5 | 12.5 |
| 6g72 | T | 3.9 | 4356 | 0.075 | 1.33 | 75 | 11.6 | 8.1 | 0.351 | 0.421 | 66.7 | 18 | 65.3 | 14.3 |
| 6g72 | V | 3.9 | 4356 | 0.075 | 1.33 | 112 | 4.1 | 4.1 | 0.797 | 0.797 | 85.7 | 53.1 | 86.6 | 6.2 |
| 6g72 | W | 3.9 | 4356 | 0.075 | 1.33 | 114 | 3.7 | 3.7 | 0.831 | 0.831 | 89.5 | 58.8 | 89.5 | 49 |
| 6g72 | Y | 3.9 | 4356 | 0.075 | 1.33 | 140 | 5.0 | 4.0 | 0.832 | 0.832 | 87.1 | 39.3 | 80 | 34.8 |
| 6g79 | S | 3.8 | 4358 | 0.07 | 1.06 | 262 | 3.8 | 3.8 | 0.866 | 0.866 | 87.8 | 58.7 | 88.5 | 28.4 |
| 6g90 | F | 4 | 4364 | 0.0369 | 1.13 | 175 | 8.2 | 6.6 | 0.738 | 0.761 | 80.6 | 44.7 | 81.1 | 6.3 |
| 6g90 | G | 4 | 4364 | 0.0369 | 1.13 | 216 | 10.8 | 10.8 | 0.748 | 0.748 | 77.3 | 50.9 | 88.4 | 6.8 |
| 6gcs | 8 | 4.3 | 4384 | 0.045 | 1.09 | 82 | 5.2 | 3.6 | 0.736 | 0.736 | 89 | 27.4 | 82.9 | 7.4 |
| 6gcs | f | 4.3 | 4384 | 0.045 | 1.09 | 80 | 4.5 | 4.5 | 0.695 | 0.695 | 78.8 | 41.3 | 70 | 8.9 |
| 6gcs | F | 4.3 | 4384 | 0.045 | 1.09 | 119 | 3.8 | 3.8 | 0.777 | 0.777 | 82.4 | 49 | 50.4 | 8.3 |
| 6gcs | H | 4.3 | 4384 | 0.045 | 1.09 | 185 | 8.6 | 6.8 | 0.715 | 0.725 | 81.1 | 18 | 72.4 | 6 |
| 6gcs | J | 4.3 | 4384 | 0.045 | 1.09 | 140 | 4.1 | 4.1 | 0.808 | 0.808 | 88.6 | 36.3 | 72.9 | 26.5 |
| 6gcs | O | 4.3 | 4384 | 0.045 | 1.09 | 77 | 11.2 | 9.6 | 0.389 | 0.490 | 81.8 | 17.5 | 81.8 | 7.9 |
| 6gcs | P | 4.3 | 4384 | 0.045 | 1.09 | 116 | 4.4 | 4.4 | 0.759 | 0.795 | 81 | 42.6 | 77.6 | 5.6 |
| 6gcs | Q | 4.3 | 4384 | 0.045 | 1.09 | 85 | 4.2 | 4.2 | 0.733 | 0.733 | 83.5 | 22.5 | 78.8 | 6 |
| 6gcs | U | 4.3 | 4384 | 0.045 | 1.09 | 158 | 5.7 | 5.7 | 0.824 | 0.827 | 87.3 | 30.4 | 81 | 18.8 |
| 6gcs | X | 4.3 | 4384 | 0.045 | 1.09 | 117 | 5.7 | 5.6 | 0.803 | 0.803 | 85.5 | 24 | 80.3 | 4.3 |
| 6gdg | A | 4.1 | 4390 | 0.0631 | 1.07 | 264 | 10.2 | 7.4 | 0.762 | 0.762 | 76.1 | 20.4 | 45.8 | 7.4 |
| 6gdg | E | 4.1 | 4390 | 0.0631 | 1.07 | 128 | 16.0 | 14.6 | 0.357 | 0.357 | 68.8 | 5.7 | 58.6 | 5.3 |

| | | | | | | | | | | | | | | |
|---|---|---|---|---|---|---|---|---|---|---|---|---|---|---|
| 6gfw | A | 3.7 | 4397 | 0.035 | 1.08 | 233 | 1.7 | 1.7 | 0.939 | 0.939 | 96.1 | 90.6 | 81.1 | 20.1 |
| 6gfw | B | 3.7 | 4397 | 0.035 | 1.08 | 235 | 13.0 | 12.4 | 0.666 | 0.666 | 86 | 35.6 | 51.1 | 10.8 |
| 6gh5 | B | 3.4 | 0001 | 0.018 | 1.06 | 235 | 24.4 | 13.3 | 0.261 | 0.704 | 81.3 | 3.1 | 65.5 | 17.5 |
| 6gh5 | E | 3.4 | 0001 | 0.018 | 1.06 | 75 | 2.3 | 2.3 | 0.831 | 0.831 | 93.3 | 62.9 | 82.7 | 37.1 |
| 6gh6 | A | 4.1 | 0002 | 0.016 | 1.08 | 233 | 12.3 | 11.9 | 0.565 | 0.675 | 78.5 | 38.8 | 74.7 | 6.3 |
| 6gh6 | B | 4.1 | 0002 | 0.016 | 1.08 | 235 | 12.5 | 12.5 | 0.553 | 0.564 | 78.3 | 32.1 | 45.5 | 37.4 |
| 6giq | D | 3.2 | 0004 | 0.33 | 1.06 | 248 | 1.6 | 1.6 | 0.960 | 0.960 | 97.6 | 92.1 | 84.7 | 31 |
| 6giq | F | 3.2 | 0004 | 0.33 | 1.06 | 74 | 2.3 | 2.3 | 0.854 | 0.854 | 90.5 | 82.1 | 85.1 | 31.7 |
| 6giq | G | 3.2 | 0004 | 0.33 | 1.06 | 126 | 1.2 | 1.2 | 0.937 | 0.937 | 97.6 | 94.3 | 91.3 | 54.8 |
| 6giq | O | 3.2 | 0004 | 0.33 | 1.06 | 248 | 1.3 | 1.3 | 0.963 | 0.963 | 98.8 | 96.3 | 87.1 | 60.2 |
| 6giq | Q | 3.2 | 0004 | 0.33 | 1.06 | 75 | 3.2 | 3.2 | 0.874 | 0.874 | 94.7 | 43.7 | 84 | 38.1 |
| 6giq | R | 3.2 | 0004 | 0.33 | 1.06 | 126 | 1.5 | 1.5 | 0.937 | 0.937 | 97.6 | 90.2 | 89.7 | 46 |
| 6giq | T | 3.2 | 0004 | 0.33 | 1.06 | 54 | 1.8 | 1.8 | 0.845 | 0.845 | 96.3 | 92.3 | 87 | 63.8 |
| 6gmh | C | 3.1 | 0031 | 0.0157 | 1.049 | 263 | 2.6 | 2.6 | 0.945 | 0.945 | 95.1 | 75.2 | 87.1 | 47.6 |
| 6gmh | E | 3.1 | 0031 | 0.0157 | 1.049 | 209 | 2.4 | 2.4 | 0.923 | 0.923 | 95.7 | 80 | 84.2 | 11.4 |
| 6gmh | F | 3.1 | 0031 | 0.0157 | 1.049 | 82 | 3.8 | 3.8 | 0.880 | 0.880 | 93.9 | 66.2 | 82.9 | 55.9 |
| 6gmh | H | 3.1 | 0031 | 0.0157 | 1.049 | 148 | 3.4 | 3.4 | 0.887 | 0.887 | 91.9 | 69.9 | 92.6 | 26.3 |
| 6gmh | I | 3.1 | 0031 | 0.0157 | 1.049 | 117 | 2.7 | 2.7 | 0.854 | 0.854 | 89.7 | 56.2 | 56.4 | 4.5 |
| 6gmh | J | 3.1 | 0031 | 0.0157 | 1.049 | 67 | 2.3 | 2.3 | 0.837 | 0.837 | 91 | 85.2 | 91 | 54.1 |
| 6gmh | K | 3.1 | 0031 | 0.0157 | 1.049 | 115 | 1.6 | 1.6 | 0.935 | 0.935 | 97.4 | 90.2 | 92.2 | 3.8 |
| 6gml | C | 3.2 | 0038 | 0.00596 | 1.2277 | 258 | 2.6 | 2.6 | 0.947 | 0.947 | 95.3 | 78.5 | 86.4 | 48.9 |
| 6gml | E | 3.2 | 0038 | 0.00596 | 1.2277 | 209 | 3.6 | 3.6 | 0.884 | 0.884 | 90.4 | 65.1 | 76.1 | 32.7 |
| 6gml | F | 3.2 | 0038 | 0.00596 | 1.2277 | 82 | 3.6 | 3.6 | 0.843 | 0.843 | 91.5 | 45.3 | 89 | 94.5 |
| 6gml | H | 3.2 | 0038 | 0.00596 | 1.2277 | 148 | 3.0 | 3.0 | 0.882 | 0.882 | 91.9 | 67.6 | 87.2 | 7.8 |
| 6gml | I | 3.2 | 0038 | 0.00596 | 1.2277 | 117 | 5.1 | 5.0 | 0.815 | 0.815 | 85.5 | 41 | 74.4 | 24.1 |
| 6gml | J | 3.2 | 0038 | 0.00596 | 1.2277 | 67 | 1.8 | 1.8 | 0.876 | 0.876 | 97 | 89.2 | 79.1 | 1.9 |
| 6gml | K | 3.2 | 0038 | 0.00596 | 1.2277 | 115 | 1.8 | 1.8 | 0.912 | 0.912 | 96.5 | 85.6 | 93 | 80.4 |
| 6gov | U | 3.7 | 0043 | 0.0138 | 1.35 | 230 | 1.7 | 1.7 | 0.944 | 0.944 | 96.1 | 85.5 | 85.7 | 30.5 |
| 6gov | V | 3.7 | 0043 | 0.0138 | 1.35 | 220 | 12.4 | 12.2 | 0.602 | 0.602 | 84.5 | 37.6 | 60.9 | 45.5 |
| 6h3c | B | 3.9 | 0132 | 0.00816 | 0.86 | 258 | 13.1 | 6.1 | 0.849 | 0.852 | 86.8 | 25.4 | 77.1 | 7 |
| 6h3c | G | 3.9 | 0132 | 0.00816 | 0.86 | 258 | 16.9 | 8.4 | 0.511 | 0.722 | 79.8 | 33 | 75.6 | 31.3 |
| 6h67 | E | 3.6 | 0146 | 0.0162 | 1.06 | 212 | 13.9 | 12.1 | 0.638 | 0.648 | 69.8 | 18.9 | 53.3 | 3.5 |
| 6h67 | H | 3.6 | 0146 | 0.0162 | 1.06 | 131 | 16.2 | 14.6 | 0.288 | 0.451 | 67.9 | 7.9 | 31.3 | 0 |
| 6h68 | E | 4.6 | 0147 | 0.0239 | 1.06 | 215 | 19.8 | 16.3 | 0.231 | 0.351 | 64.2 | 8 | 30.2 | 6.2 |
| 6h68 | H | 4.6 | 0147 | 0.0239 | 1.06 | 132 | 16.0 | 14.5 | 0.317 | 0.317 | 56.1 | 4.1 | 16.7 | 13.6 |
| 6hco | C | 3.6 | 0196 | 0.32 | 0.812 | 107 | 15.7 | 13.6 | 0.391 | 0.657 | 82.2 | 4.5 | 70.1 | 6.7 |
| 6hco | D | 3.6 | 0196 | 0.32 | 0.812 | 118 | 15.7 | 13.3 | 0.365 | 0.520 | 67.8 | 6.2 | 61.9 | 6.8 |

| | | | | | | | | | | | | | | |
|---|---|---|---|---|---|---|---|---|---|---|---|---|---|---|
| 6hco | E | 3.6 | 0196 | 0.32 | 0.812 | 107 | 15.3 | 8.4 | 0.386 | 0.706 | 86 | 6.5 | 63.6 | 5.9 |
| 6hco | F | 3.6 | 0196 | 0.32 | 0.812 | 118 | 16.5 | 13.3 | 0.250 | 0.434 | 68.6 | 8.6 | 30.5 | 5.6 |
| 6hjn | B | 3.3 | 0234 | 1 | 1.078 | 173 | 1.4 | 1.4 | 0.944 | 0.944 | 98.8 | 92.4 | 89 | 42.9 |
| 6hjn | D | 3.3 | 0234 | 1 | 1.078 | 173 | 2.2 | 2.2 | 0.926 | 0.926 | 95.4 | 81.8 | 91.9 | 64.2 |
| 6hjn | F | 3.3 | 0234 | 1 | 1.078 | 173 | 1.7 | 1.7 | 0.942 | 0.942 | 97.7 | 88.2 | 87.3 | 31.1 |
| 6hjp | B | 3.3 | 0235 | 1 | 1.078 | 174 | 1.7 | 1.7 | 0.944 | 0.944 | 97.7 | 90 | 96 | 70.1 |
| 6hjp | D | 3.3 | 0235 | 1 | 1.078 | 174 | 1.3 | 1.3 | 0.954 | 0.954 | 98.3 | 95.9 | 96.6 | 13.7 |
| 6hjp | F | 3.3 | 0235 | 1 | 1.078 | 174 | 1.7 | 1.7 | 0.941 | 0.941 | 97.1 | 94.7 | 96 | 55.1 |
| 6hjp | G | 3.3 | 0235 | 1 | 1.078 | 216 | 12.5 | 9.9 | 0.752 | 0.763 | 87.5 | 56.1 | 64.8 | 43.6 |
| 6hjp | H | 3.3 | 0235 | 1 | 1.078 | 214 | 16.7 | 14.0 | 0.379 | 0.493 | 75.7 | 9.9 | 44.4 | 4.2 |
| 6hjp | I | 3.3 | 0235 | 1 | 1.078 | 216 | 10.2 | 10.2 | 0.760 | 0.762 | 87 | 54.3 | 58.8 | 27.6 |
| 6hjp | J | 3.3 | 0235 | 1 | 1.078 | 214 | 22.2 | 17.6 | 0.539 | 0.582 | 66.4 | 7.7 | 42.5 | 5.5 |
| 6hjp | K | 3.3 | 0235 | 1 | 1.078 | 216 | 10.1 | 9.7 | 0.712 | 0.762 | 87.5 | 61.9 | 57.9 | 6.4 |
| 6hjp | L | 3.3 | 0235 | 1 | 1.078 | 214 | 19.6 | 17.3 | 0.522 | 0.561 | 75.7 | 13.6 | 46.7 | 9 |
| 6hjq | B | 4.1 | 0236 | 1 | 1.078 | 203 | 3.4 | 3.4 | 0.848 | 0.848 | 86.7 | 61.9 | 68.5 | 12.9 |
| 6hjq | D | 4.1 | 0236 | 1 | 1.078 | 203 | 20.4 | 5.4 | 0.362 | 0.800 | 77.3 | 17.8 | 68.5 | 10.8 |
| 6hjq | F | 4.1 | 0236 | 1 | 1.078 | 203 | 13.4 | 4.9 | 0.805 | 0.848 | 81.3 | 19.4 | 76.4 | 9.7 |
| 6hjq | G | 4.1 | 0236 | 1 | 1.078 | 216 | 20.9 | 19.1 | 0.280 | 0.477 | 68.5 | 6.8 | 34.7 | 8 |
| 6hjq | H | 4.1 | 0236 | 1 | 1.078 | 214 | 19.6 | 17.5 | 0.391 | 0.421 | 61.7 | 8.3 | 43.5 | 7.5 |
| 6hjq | I | 4.1 | 0236 | 1 | 1.078 | 216 | 18.7 | 18.4 | 0.470 | 0.508 | 72.2 | 7.7 | 45.4 | 12.2 |
| 6hjq | J | 4.1 | 0236 | 1 | 1.078 | 214 | 18.3 | 17.3 | 0.272 | 0.309 | 68.7 | 4.1 | 29 | 6.5 |
| 6hjq | K | 4.1 | 0236 | 1 | 1.078 | 216 | 22.3 | 20.0 | 0.260 | 0.482 | 73.1 | 3.2 | 45.4 | 15.3 |
| 6hjq | L | 4.1 | 0236 | 1 | 1.078 | 214 | 25.6 | 17.7 | 0.237 | 0.399 | 65.9 | 5.7 | 23.4 | 14 |
| 6hjr | B | 4.2 | 0237 | 1 | 1.078 | 203 | 22.5 | 6.5 | 0.361 | 0.784 | 82.3 | 15 | 71.9 | 5.5 |
| 6hjr | D | 4.2 | 0237 | 1 | 1.078 | 203 | 22.4 | 4.9 | 0.360 | 0.817 | 75.4 | 5.9 | 62.6 | 3.9 |
| 6hjr | F | 4.2 | 0237 | 1 | 1.078 | 203 | 12.7 | 5.8 | 0.730 | 0.796 | 75.9 | 32.5 | 69 | 18.6 |
| 6hko | E | 3.4 | 0238 | 0.0388 | 1.04 | 214 | 7.8 | 7.8 | 0.790 | 0.790 | 86 | 59.2 | 79.4 | 22.9 |
| 6hko | F | 3.4 | 0238 | 0.0388 | 1.04 | 100 | 1.2 | 1.2 | 0.933 | 0.933 | 98 | 92.9 | 92 | 44.6 |
| 6hko | H | 3.4 | 0238 | 0.0388 | 1.04 | 131 | 3.7 | 3.7 | 0.868 | 0.878 | 90.1 | 66.9 | 93.9 | 8.9 |
| 6hko | K | 3.4 | 0238 | 0.0388 | 1.04 | 98 | 3.1 | 3.1 | 0.893 | 0.893 | 93.9 | 64.1 | 96.9 | 43.2 |
| 6hlq | E | 3.2 | 0239 | 0.066 | 1.04 | 213 | 2.2 | 2.2 | 0.936 | 0.936 | 95.8 | 91.7 | 86.9 | 34.1 |
| 6hlq | H | 3.2 | 0239 | 0.066 | 1.04 | 131 | 6.3 | 6.3 | 0.806 | 0.806 | 86.3 | 78.8 | 87.8 | 31.3 |
| 6hlr | E | 3.2 | 0240 | 0.0493 | 1.04 | 212 | 2.1 | 2.1 | 0.941 | 0.941 | 96.2 | 83.8 | 84.9 | 59.4 |
| 6hlr | F | 3.2 | 0240 | 0.0493 | 1.04 | 100 | 2.0 | 2.0 | 0.928 | 0.928 | 98 | 89.8 | 95 | 64.2 |
| 6hlr | H | 3.2 | 0240 | 0.0493 | 1.04 | 131 | 4.6 | 4.6 | 0.874 | 0.874 | 89.3 | 82.9 | 90.8 | 29.4 |
| 6hlr | K | 3.2 | 0240 | 0.0493 | 1.04 | 98 | 1.5 | 1.5 | 0.927 | 0.927 | 98 | 89.6 | 93.9 | 78.3 |
| 6hls | E | 3.2 | 0241 | 0.0777 | 1.04 | 214 | 3.0 | 3.0 | 0.904 | 0.904 | 93 | 74.4 | 82.2 | 21 |

| | | | | | | | | | | | | | | |
|---|---|---|---|---|---|---|---|---|---|---|---|---|---|---|
| 6hls | F | 3.2 | 0241 | 0.0777 | 1.04 | 100 | 1.5 | 1.5 | 0.928 | 0.928 | 98 | 86.7 | 82 | 96.3 |
| 6hls | H | 3.2 | 0241 | 0.0777 | 1.04 | 131 | 14.0 | 11.6 | 0.518 | 0.635 | 87.8 | 31.3 | 86.3 | 3.5 |
| 6hls | K | 3.2 | 0241 | 0.0777 | 1.04 | 98 | 2.7 | 2.7 | 0.897 | 0.897 | 94.9 | 49.5 | 95.9 | 90.4 |
| 6hn4 | E | 4.2 | 0246 | 0.03 | 1.04 | 202 | 17.2 | 17.1 | 0.334 | 0.403 | 64.4 | 5.4 | 32.2 | 1.5 |
| 6hra | C | 3.7 | 0257 | 0.04 | 1.012 | 187 | 2.4 | 2.4 | 0.914 | 0.914 | 95.2 | 69.1 | 87.2 | 33.1 |
| 6hrb | C | 4 | 0258 | 0.0595 | 1.012 | 189 | 9.4 | 8.3 | 0.752 | 0.759 | 76.7 | 35.9 | 73.5 | 7.9 |
| 6hu9 | b | 3.3 | 0262 | 0.0263 | 1.3861 | 236 | 12.3 | 7.2 | 0.718 | 0.802 | 85.6 | 40.1 | 63.6 | 34 |
| 6hu9 | D | 3.3 | 0262 | 0.0263 | 1.3861 | 247 | 1.7 | 1.7 | 0.959 | 0.959 | 98 | 81.4 | 92.3 | 28.1 |
| 6hu9 | e | 3.3 | 0262 | 0.0263 | 1.3861 | 133 | 4.2 | 3.8 | 0.881 | 0.881 | 89.5 | 49.6 | 87.2 | 73.3 |
| 6hu9 | f | 3.3 | 0262 | 0.0263 | 1.3861 | 102 | 3.3 | 3.3 | 0.808 | 0.808 | 89.2 | 75.8 | 78.4 | 76.2 |
| 6hu9 | F | 3.3 | 0262 | 0.0263 | 1.3861 | 75 | 1.8 | 1.8 | 0.880 | 0.880 | 96 | 87.5 | 85.3 | 40.6 |
| 6hu9 | G | 3.3 | 0262 | 0.0263 | 1.3861 | 126 | 1.7 | 1.7 | 0.945 | 0.945 | 96 | 93.4 | 78.6 | 33.3 |
| 6hu9 | O | 3.3 | 0262 | 0.0263 | 1.3861 | 247 | 1.2 | 1.2 | 0.962 | 0.962 | 96.8 | 91.6 | 84.6 | 49.3 |
| 6hu9 | Q | 3.3 | 0262 | 0.0263 | 1.3861 | 75 | 1.9 | 1.9 | 0.888 | 0.888 | 93.3 | 80 | 84 | 36.5 |
| 6hu9 | R | 3.3 | 0262 | 0.0263 | 1.3861 | 126 | 1.4 | 1.4 | 0.949 | 0.949 | 97.6 | 93.5 | 85.7 | 32.4 |
| 6hu9 | T | 3.3 | 0262 | 0.0263 | 1.3861 | 57 | 1.7 | 1.7 | 0.890 | 0.890 | 94.7 | 88.9 | 89.5 | 58.8 |
| 6hug | G | 3.1 | 0275 | 0.062 | 1.055 | 121 | 2.7 | 2.7 | 0.907 | 0.907 | 95 | 81.7 | 61.2 | 5.4 |
| 6huj | G | 3 | 0279 | 0.042 | 1.07 | 121 | 4.5 | 4.5 | 0.831 | 0.831 | 86 | 55.8 | 79.3 | 21.9 |
| 6huk | G | 3.7 | 0280 | 0.07 | 1.07 | 121 | 15.0 | 9.5 | 0.340 | 0.734 | 77.7 | 11.7 | 56.2 | 5.9 |
| 6huo | G | 3.3 | 0282 | 0.06 | 1.07 | 121 | 3.9 | 3.9 | 0.871 | 0.871 | 90.9 | 67.3 | 59.5 | 6.9 |
| 6hup | G | 3.6 | 0283 | 0.025 | 0.895 | 121 | 12.8 | 11.0 | 0.619 | 0.619 | 73.6 | 24.7 | 61.2 | 9.5 |
| 6hwh | L | 3.3 | 0289 | 0.475 | 1.06 | 281 | 2.6 | 2.6 | 0.931 | 0.931 | 94.3 | 79.6 | 83.3 | 53 |
| 6hwh | O | 3.3 | 0289 | 0.475 | 1.06 | 142 | 2.1 | 2.1 | 0.900 | 0.900 | 93 | 75.8 | 89.4 | 26.8 |
| 6hwh | P | 3.3 | 0289 | 0.475 | 1.06 | 281 | 11.2 | 5.0 | 0.740 | 0.868 | 87.2 | 53.9 | 85.1 | 37.2 |
| 6hwh | T | 3.3 | 0289 | 0.475 | 1.06 | 142 | 4.0 | 4.0 | 0.855 | 0.863 | 90.1 | 62.5 | 82.4 | 54.7 |
| 6hwh | Z | 3.3 | 0289 | 0.475 | 1.06 | 184 | 1.6 | 1.6 | 0.943 | 0.943 | 96.7 | 89.9 | 90.2 | 40.4 |
| 6hz4 | A | 3.6 | 0310 | 0.06 | 1.05 | 290 | 7.8 | 7.8 | 0.793 | 0.793 | 86.2 | 49.6 | 65.5 | 17.9 |
| 6hz4 | B | 3.6 | 0310 | 0.06 | 1.05 | 290 | 12.3 | 11.8 | 0.713 | 0.757 | 80.7 | 29.9 | 76.2 | 9.5 |
| 6hz4 | C | 3.6 | 0310 | 0.06 | 1.05 | 290 | 7.8 | 5.5 | 0.867 | 0.867 | 87.6 | 66.1 | 77.2 | 4.5 |
| 6hz4 | D | 3.6 | 0310 | 0.06 | 1.05 | 284 | 2.6 | 2.6 | 0.909 | 0.909 | 91.2 | 68 | 76.8 | 10.1 |
| 6hz4 | E | 3.6 | 0310 | 0.06 | 1.05 | 285 | 15.9 | 11.0 | 0.590 | 0.720 | 78.6 | 52.2 | 69.1 | 14.7 |
| 6hz4 | F | 3.6 | 0310 | 0.06 | 1.05 | 285 | 3.5 | 3.5 | 0.871 | 0.871 | 87.4 | 54.2 | 67.4 | 7.8 |
| 6hz4 | N | 3.6 | 0310 | 0.06 | 1.05 | 152 | 7.7 | 7.7 | 0.645 | 0.658 | 77.6 | 24.6 | 73.7 | 8 |
| 6hz5 | A | 4.2 | 0311 | 0.024 | 1.05 | 290 | 22.2 | 17.0 | 0.404 | 0.404 | 71 | 10.7 | 55.9 | 7.4 |
| 6hz5 | B | 4.2 | 0311 | 0.024 | 1.05 | 290 | 6.0 | 6.0 | 0.796 | 0.796 | 78.6 | 28.5 | 59.7 | 10.4 |
| 6hz5 | C | 4.2 | 0311 | 0.024 | 1.05 | 290 | 10.5 | 10.5 | 0.670 | 0.670 | 73.8 | 38.8 | 62.8 | 5.5 |
| 6hz5 | D | 4.2 | 0311 | 0.024 | 1.05 | 284 | 18.8 | 15.8 | 0.454 | 0.679 | 69.7 | 12.1 | 65.1 | 8.6 |

| | | | | | | | | | | | | | | |
|---|---|---|---|---|---|---|---|---|---|---|---|---|---|---|
| 6hz5 | E | 4.2 | 0311 | 0.024 | 1.05 | 285 | 22.0 | 17.6 | 0.279 | 0.395 | 70.2 | 5.5 | 60.7 | 5.2 |
| 6hz5 | F | 4.2 | 0311 | 0.024 | 1.05 | 285 | 17.9 | 17.6 | 0.416 | 0.416 | 68.1 | 11.3 | 42.8 | 4.1 |
| 6hz5 | G | 4.2 | 0311 | 0.024 | 1.05 | 290 | 24.1 | 18.8 | 0.369 | 0.491 | 71 | 11.7 | 27.2 | 8.9 |
| 6hz5 | H | 4.2 | 0311 | 0.024 | 1.05 | 290 | 24.7 | 21.2 | 0.365 | 0.476 | 76.9 | 16.1 | 64.1 | 9.1 |
| 6hz5 | I | 4.2 | 0311 | 0.024 | 1.05 | 290 | 20.9 | 15.7 | 0.362 | 0.525 | 67.6 | 11.7 | 62.4 | 7.7 |
| 6hz5 | J | 4.2 | 0311 | 0.024 | 1.05 | 284 | 21.0 | 12.2 | 0.284 | 0.674 | 72.5 | 5.3 | 24.3 | 10.1 |
| 6hz5 | K | 4.2 | 0311 | 0.024 | 1.05 | 285 | 18.9 | 18.9 | 0.286 | 0.435 | 66 | 7.4 | 51.6 | 4.8 |
| 6hz5 | L | 4.2 | 0311 | 0.024 | 1.05 | 285 | 22.4 | 22.4 | 0.241 | 0.269 | 71.2 | 4.9 | 45.6 | 7.7 |
| 6hz6 | A | 4.3 | 0312 | 0.026 | 1.05 | 290 | 23.2 | 17.2 | 0.271 | 0.401 | 71 | 8.7 | 52.8 | 7.8 |
| 6hz6 | B | 4.3 | 0312 | 0.026 | 1.05 | 290 | 18.8 | 16.5 | 0.419 | 0.492 | 72.4 | 10 | 62.1 | 3.9 |
| 6hz6 | C | 4.3 | 0312 | 0.026 | 1.05 | 290 | 18.6 | 15.4 | 0.380 | 0.540 | 66.6 | 10.4 | 51 | 8.1 |
| 6hz6 | D | 4.3 | 0312 | 0.026 | 1.05 | 284 | 12.6 | 10.8 | 0.706 | 0.706 | 76.8 | 38.1 | 45.1 | 3.9 |
| 6hz6 | E | 4.3 | 0312 | 0.026 | 1.05 | 285 | 20.2 | 18.2 | 0.436 | 0.436 | 64.9 | 4.9 | 53 | 7.3 |
| 6hz6 | F | 4.3 | 0312 | 0.026 | 1.05 | 285 | 24.5 | 20.7 | 0.291 | 0.388 | 64.9 | 1.6 | 34 | 9.3 |
| 6hz6 | G | 4.3 | 0312 | 0.026 | 1.05 | 290 | 22.1 | 15.5 | 0.404 | 0.494 | 72.4 | 11.9 | 52.8 | 5.9 |
| 6hz6 | H | 4.3 | 0312 | 0.026 | 1.05 | 290 | 23.2 | 21.0 | 0.257 | 0.312 | 69.3 | 5.5 | 23.8 | 10.1 |
| 6hz6 | I | 4.3 | 0312 | 0.026 | 1.05 | 290 | 14.2 | 10.7 | 0.680 | 0.694 | 73.1 | 23.1 | 56.2 | 8.6 |
| 6hz6 | J | 4.3 | 0312 | 0.026 | 1.05 | 284 | 21.9 | 20.3 | 0.343 | 0.358 | 74.3 | 7.1 | 67.3 | 4.2 |
| 6hz6 | K | 4.3 | 0312 | 0.026 | 1.05 | 285 | 18.8 | 18.2 | 0.315 | 0.315 | 64.6 | 4.3 | 57.5 | 7.3 |
| 6hz6 | L | 4.3 | 0312 | 0.026 | 1.05 | 285 | 24.5 | 20.5 | 0.287 | 0.287 | 58.9 | 6 | 39.6 | 2.7 |
| 6hz7 | A | 4.3 | 0313 | 0.026 | 1.05 | 290 | 21.3 | 20.0 | 0.342 | 0.499 | 72.4 | 12.4 | 48.3 | 6.4 |
| 6hz7 | B | 4.3 | 0313 | 0.026 | 1.05 | 290 | 18.7 | 16.1 | 0.380 | 0.490 | 69.3 | 16.4 | 68.3 | 17.7 |
| 6hz7 | C | 4.3 | 0313 | 0.026 | 1.05 | 290 | 20.5 | 15.2 | 0.432 | 0.526 | 70.3 | 10.8 | 62.4 | 8.8 |
| 6hz7 | D | 4.3 | 0313 | 0.026 | 1.05 | 284 | 22.1 | 18.6 | 0.319 | 0.326 | 63.7 | 5.5 | 57.7 | 4.9 |
| 6hz7 | E | 4.3 | 0313 | 0.026 | 1.05 | 285 | 15.0 | 15.0 | 0.534 | 0.587 | 68.4 | 17.4 | 50.5 | 9 |
| 6hz7 | F | 4.3 | 0313 | 0.026 | 1.05 | 285 | 22.4 | 18.4 | 0.242 | 0.327 | 64.9 | 8.1 | 49.5 | 5 |
| 6hz7 | G | 4.3 | 0313 | 0.026 | 1.05 | 290 | 21.8 | 16.4 | 0.298 | 0.558 | 71.4 | 6.8 | 54.8 | 5 |
| 6hz7 | H | 4.3 | 0313 | 0.026 | 1.05 | 290 | 24.0 | 16.6 | 0.241 | 0.400 | 63.1 | 7.7 | 41.7 | 5.8 |
| 6hz7 | I | 4.3 | 0313 | 0.026 | 1.05 | 290 | 17.0 | 17.0 | 0.390 | 0.475 | 64.5 | 10.2 | 67.6 | 6.6 |
| 6hz7 | J | 4.3 | 0313 | 0.026 | 1.05 | 284 | 20.3 | 16.8 | 0.337 | 0.419 | 72.2 | 7.8 | 32.7 | 11.8 |
| 6hz7 | K | 4.3 | 0313 | 0.026 | 1.05 | 285 | 25.6 | 19.3 | 0.377 | 0.383 | 70.9 | 6.4 | 63.5 | 4.4 |
| 6hz7 | L | 4.3 | 0313 | 0.026 | 1.05 | 285 | 21.4 | 20.3 | 0.248 | 0.284 | 66 | 6.4 | 55.4 | 3.2 |
| 6hz8 | A | 4.3 | 0314 | 0.026 | 1.05 | 290 | 23.3 | 12.4 | 0.383 | 0.556 | 68.3 | 10.1 | 41 | 7.6 |
| 6hz8 | B | 4.3 | 0314 | 0.026 | 1.05 | 290 | 17.4 | 17.4 | 0.329 | 0.425 | 70.7 | 11.2 | 38.6 | 5.4 |
| 6hz8 | C | 4.3 | 0314 | 0.026 | 1.05 | 290 | 12.0 | 12.0 | 0.643 | 0.669 | 67.9 | 24.4 | 67.9 | 4.6 |
| 6hz8 | D | 4.3 | 0314 | 0.026 | 1.05 | 284 | 22.8 | 21.2 | 0.329 | 0.338 | 69.4 | 8.1 | 52.5 | 5.4 |
| 6hz8 | E | 4.3 | 0314 | 0.026 | 1.05 | 285 | 21.8 | 19.9 | 0.272 | 0.426 | 60.7 | 9.2 | 54.7 | 6.4 |

| | | | | | | | | | | | | | | |
|---|---|---|---|---|---|---|---|---|---|---|---|---|---|---|
| 6hz8 | F | 4.3 | 0314 | 0.026 | 1.05 | 285 | 25.9 | 21.8 | 0.251 | 0.311 | 62.1 | 6.8 | 36.8 | 6.7 |
| 6hz8 | G | 4.3 | 0314 | 0.026 | 1.05 | 290 | 21.7 | 20.0 | 0.283 | 0.363 | 65.5 | 8.9 | 47.9 | 7.2 |
| 6hz8 | H | 4.3 | 0314 | 0.026 | 1.05 | 290 | 16.5 | 15.1 | 0.567 | 0.567 | 71.4 | 18.4 | 57.6 | 5.4 |
| 6hz8 | I | 4.3 | 0314 | 0.026 | 1.05 | 290 | 19.6 | 15.8 | 0.370 | 0.461 | 66.9 | 12.9 | 49.3 | 3.5 |
| 6hz8 | J | 4.3 | 0314 | 0.026 | 1.05 | 284 | 20.8 | 17.1 | 0.257 | 0.417 | 65.1 | 7 | 59.2 | 4.8 |
| 6hz8 | K | 4.3 | 0314 | 0.026 | 1.05 | 285 | 17.4 | 15.7 | 0.279 | 0.405 | 69.5 | 8.1 | 44.6 | 3.1 |
| 6hz8 | L | 4.3 | 0314 | 0.026 | 1.05 | 285 | 23.9 | 20.2 | 0.227 | 0.347 | 67.4 | 4.7 | 45.6 | 7.7 |
| 6hz9 | A | 4.8 | 0315 | 0.023 | 1.05 | 290 | 20.4 | 17.6 | 0.306 | 0.373 | 62.4 | 6.1 | 39.3 | 3.5 |
| 6hz9 | B | 4.8 | 0315 | 0.023 | 1.05 | 290 | 24.2 | 19.0 | 0.282 | 0.482 | 64.5 | 5.3 | 43.1 | 7.2 |
| 6hz9 | C | 4.8 | 0315 | 0.023 | 1.05 | 290 | 24.9 | 22.2 | 0.356 | 0.369 | 59.3 | 7 | 24.1 | 5.7 |
| 6hz9 | D | 4.8 | 0315 | 0.023 | 1.05 | 284 | 16.7 | 16.7 | 0.373 | 0.380 | 62.7 | 9.6 | 48.2 | 4.4 |
| 6hz9 | E | 4.8 | 0315 | 0.023 | 1.05 | 285 | 20.4 | 17.0 | 0.273 | 0.302 | 62.5 | 9.6 | 48.1 | 5.8 |
| 6hz9 | F | 4.8 | 0315 | 0.023 | 1.05 | 285 | 24.3 | 19.4 | 0.269 | 0.341 | 57.9 | 10.9 | 27.4 | 5.1 |
| 6hz9 | G | 4.8 | 0315 | 0.023 | 1.05 | 290 | 20.3 | 19.6 | 0.319 | 0.342 | 59.7 | 7.5 | 31.7 | 4.3 |
| 6hz9 | H | 4.8 | 0315 | 0.023 | 1.05 | 290 | 19.7 | 18.0 | 0.296 | 0.312 | 61 | 7.3 | 61 | 6.2 |
| 6hz9 | I | 4.8 | 0315 | 0.023 | 1.05 | 290 | 23.2 | 16.8 | 0.282 | 0.528 | 63.4 | 6.5 | 59.3 | 6.4 |
| 6hz9 | J | 4.8 | 0315 | 0.023 | 1.05 | 284 | 21.7 | 17.9 | 0.295 | 0.386 | 61.3 | 5.2 | 50 | 3.5 |
| 6hz9 | K | 4.8 | 0315 | 0.023 | 1.05 | 285 | 19.3 | 18.1 | 0.349 | 0.358 | 58.6 | 6 | 49.8 | 5.6 |
| 6hz9 | L | 4.8 | 0315 | 0.023 | 1.05 | 285 | 21.6 | 21.3 | 0.263 | 0.312 | 61.1 | 4 | 45.3 | 7 |
| 6i53 | G | 3.2 | 4411 | 0.04 | 1.07 | 123 | 8.3 | 4.6 | 0.831 | 0.831 | 86.2 | 85.8 | 78 | 7.3 |
| 6i84 | C | 4.4 | 4429 | 0.0173 | 1.05 | 266 | 26.1 | 24.2 | 0.287 | 0.296 | 63.9 | 5.3 | 29.7 | 2.5 |
| 6i84 | E | 4.4 | 4429 | 0.0173 | 1.05 | 214 | 18.6 | 17.9 | 0.250 | 0.348 | 60.3 | 7 | 18.2 | 10.3 |
| 6i84 | H | 4.4 | 4429 | 0.0173 | 1.05 | 134 | 18.8 | 16.6 | 0.255 | 0.301 | 53.7 | 8.3 | 36.6 | 2 |
| 6icz | N | 3 | 9645 | 0.03 | 1.338 | 143 | 2.8 | 2.8 | 0.915 | 0.915 | 95.1 | 80.1 | 76.2 | 22.9 |
| 6icz | S | 3 | 9645 | 0.03 | 1.338 | 159 | 14.6 | 13.5 | 0.416 | 0.477 | 73 | 22.4 | 63.5 | 6.9 |
| 6id0 | N | 2.9 | 9646 | 0.025 | 1.338 | 143 | 1.6 | 1.6 | 0.925 | 0.925 | 95.1 | 91.9 | 72 | 45.6 |
| 6id0 | S | 2.9 | 9646 | 0.025 | 1.338 | 159 | 17.0 | 15.3 | 0.306 | 0.391 | 80.5 | 2.3 | 42.8 | 5.9 |
| 6id1 | N | 2.9 | 9647 | 0.02 | 1.338 | 143 | 2.4 | 2.4 | 0.906 | 0.906 | 93 | 77.4 | 77.6 | 45 |
| 6ifk | B | 3.2 | 9653 | 0.065 | 1.04 | 295 | 3.2 | 3.2 | 0.917 | 0.917 | 92.2 | 71.3 | 86.1 | 10.2 |
| 6ifk | C | 3.2 | 9653 | 0.065 | 1.04 | 121 | 1.6 | 1.6 | 0.931 | 0.931 | 95.9 | 92.2 | 81.8 | 71.7 |
| 6ifk | D | 3.2 | 9653 | 0.065 | 1.04 | 121 | 3.4 | 3.4 | 0.882 | 0.882 | 90.9 | 50 | 82.6 | 25 |
| 6ifk | E | 3.2 | 9653 | 0.065 | 1.04 | 207 | 2.2 | 2.2 | 0.930 | 0.930 | 95.7 | 79.8 | 85.5 | 10.7 |
| 6ifk | F | 3.2 | 9653 | 0.065 | 1.04 | 216 | 2.0 | 2.0 | 0.941 | 0.941 | 95.8 | 77.3 | 83.3 | 30 |
| 6ifl | B | 3.2 | 9654 | 0.07 | 1.04 | 121 | 2.8 | 2.8 | 0.875 | 0.875 | 92.6 | 77.7 | 89.3 | 81.5 |
| 6ifl | C | 3.2 | 9654 | 0.07 | 1.04 | 121 | 3.1 | 3.1 | 0.914 | 0.914 | 95 | 68.7 | 85.1 | 68.9 |
| 6ifl | D | 3.2 | 9654 | 0.07 | 1.04 | 217 | 1.7 | 1.7 | 0.963 | 0.963 | 98.6 | 91.1 | 92.2 | 38 |
| 6ifl | E | 3.2 | 9654 | 0.07 | 1.04 | 217 | 2.1 | 2.1 | 0.941 | 0.941 | 95.9 | 78.4 | 90.3 | 17.3 |

| | | | | | | | | | | | | | | |
|---|---|---|---|---|---|---|---|---|---|---|---|---|---|---|
| 6ifl | F | 3.2 | 9654 | 0.07 | 1.04 | 205 | 1.7 | 1.7 | 0.941 | 0.941 | 96.1 | 87.8 | 90.7 | 50.5 |
| 6ifl | G | 3.2 | 9654 | 0.07 | 1.04 | 273 | 2.8 | 2.8 | 0.929 | 0.929 | 93.8 | 75.4 | 90.1 | 30.9 |
| 6ifr | B | 3.4 | 9656 | 0.062 | 1.04 | 288 | 16.4 | 9.7 | 0.789 | 0.821 | 78.8 | 22.5 | 85.4 | 14.6 |
| 6ifr | C | 3.4 | 9656 | 0.062 | 1.04 | 121 | 1.6 | 1.6 | 0.923 | 0.923 | 98.3 | 91.6 | 77.7 | 75.5 |
| 6ifr | D | 3.4 | 9656 | 0.062 | 1.04 | 121 | 3.2 | 3.2 | 0.847 | 0.847 | 92.6 | 67.9 | 83.5 | 37.6 |
| 6ifr | E | 3.4 | 9656 | 0.062 | 1.04 | 205 | 2.5 | 2.5 | 0.903 | 0.908 | 92.7 | 81.6 | 78 | 23.1 |
| 6ifr | F | 3.4 | 9656 | 0.062 | 1.04 | 217 | 2.4 | 2.4 | 0.919 | 0.919 | 93.5 | 71.4 | 88 | 17.3 |
| 6ifr | G | 3.4 | 9656 | 0.062 | 1.04 | 217 | 2.4 | 2.4 | 0.925 | 0.925 | 94 | 74 | 88 | 20.9 |
| 6ifu | B | 3.1 | 9657 | 0.076 | 1.04 | 121 | 1.8 | 1.8 | 0.926 | 0.926 | 95 | 87 | 83.5 | 29.7 |
| 6ifu | C | 3.1 | 9657 | 0.076 | 1.04 | 121 | 1.5 | 1.5 | 0.930 | 0.930 | 95.9 | 82.8 | 89.3 | 56.5 |
| 6ifu | G | 3.1 | 9657 | 0.076 | 1.04 | 272 | 2.9 | 2.9 | 0.909 | 0.909 | 92.6 | 73.4 | 84.6 | 26.1 |
| 6ify | B | 3.8 | 9658 | 0.048 | 1.04 | 294 | 22.6 | 19.2 | 0.272 | 0.415 | 63.6 | 8.6 | 39.8 | 6.8 |
| 6ify | C | 3.8 | 9658 | 0.048 | 1.04 | 121 | 4.9 | 4.9 | 0.761 | 0.761 | 82.6 | 35 | 51.2 | 8.1 |
| 6ify | D | 3.8 | 9658 | 0.048 | 1.04 | 121 | 4.5 | 4.5 | 0.743 | 0.743 | 81 | 25.5 | 79.3 | 8.3 |
| 6ifz | B | 3.6 | 9659 | 0.033 | 1.04 | 293 | 23.2 | 19.6 | 0.442 | 0.496 | 79.5 | 5.6 | 67.2 | 21.3 |
| 6ifz | C | 3.6 | 9659 | 0.033 | 1.04 | 121 | 3.7 | 3.7 | 0.897 | 0.897 | 93.4 | 49.6 | 81.8 | 26.3 |
| 6ifz | D | 3.6 | 9659 | 0.033 | 1.04 | 121 | 3.2 | 3.2 | 0.866 | 0.866 | 93.4 | 68.1 | 82.6 | 34 |
| 6ig0 | B | 3.4 | 9660 | 0.056 | 1.04 | 295 | 9.5 | 9.5 | 0.831 | 0.831 | 88.1 | 37.7 | 80.7 | 22.3 |
| 6ig0 | C | 3.4 | 9660 | 0.056 | 1.04 | 121 | 2.4 | 2.4 | 0.892 | 0.892 | 92.6 | 81.2 | 86.8 | 75.2 |
| 6ig0 | D | 3.4 | 9660 | 0.056 | 1.04 | 121 | 2.2 | 2.2 | 0.891 | 0.891 | 93.4 | 80.5 | 77.7 | 23.4 |
| 6ig0 | E | 3.4 | 9660 | 0.056 | 1.04 | 207 | 6.6 | 4.8 | 0.871 | 0.892 | 88.4 | 44.3 | 86 | 22.5 |
| 6ig0 | F | 3.4 | 9660 | 0.056 | 1.04 | 216 | 2.6 | 2.6 | 0.925 | 0.925 | 94.9 | 68.8 | 88 | 12.6 |
| 6ig0 | G | 3.4 | 9660 | 0.056 | 1.04 | 217 | 2.2 | 2.2 | 0.939 | 0.939 | 96.3 | 78.9 | 90.8 | 29.9 |
| 6igz | 2 | 3.5 | 9670 | 0.034 | 0.8727 | 212 | 2.2 | 2.2 | 0.919 | 0.919 | 94.3 | 84 | 85.4 | 51.4 |
| 6igz | 3 | 3.5 | 9670 | 0.034 | 0.8727 | 226 | 3.4 | 3.4 | 0.894 | 0.894 | 92.5 | 58.4 | 80.1 | 58.6 |
| 6igz | 6 | 3.5 | 9670 | 0.034 | 0.8727 | 229 | 4.4 | 4.4 | 0.877 | 0.877 | 90.4 | 32.4 | 83.4 | 29.3 |
| 6igz | 7 | 3.5 | 9670 | 0.034 | 0.8727 | 228 | 2.8 | 2.8 | 0.893 | 0.893 | 90.8 | 56.5 | 72.4 | 32.1 |
| 6igz | D | 3.5 | 9670 | 0.034 | 0.8727 | 142 | 6.4 | 4.7 | 0.848 | 0.859 | 89.4 | 57.5 | 83.8 | 10.9 |
| 6igz | E | 3.5 | 9670 | 0.034 | 0.8727 | 61 | 4.5 | 4.5 | 0.731 | 0.731 | 83.6 | 68.6 | 90.2 | 20 |
| 6igz | F | 3.5 | 9670 | 0.034 | 0.8727 | 163 | 3.4 | 3.4 | 0.878 | 0.878 | 90.8 | 48 | 76.7 | 51.2 |
| 6ijj | 3 | 2.9 | 9678 | 0.07 | 1.04 | 220 | 2.3 | 2.3 | 0.931 | 0.931 | 95.5 | 87.2 | 73.3 | 55.6 |
| 6ijj | 4 | 2.9 | 9678 | 0.07 | 1.04 | 210 | 3.2 | 3.2 | 0.882 | 0.882 | 89.5 | 60.6 | 79 | 41 |
| 6ijj | 5 | 2.9 | 9678 | 0.07 | 1.04 | 226 | 1.5 | 1.5 | 0.945 | 0.945 | 97.3 | 91.4 | 85.4 | 59.1 |
| 6ijj | 6 | 2.9 | 9678 | 0.07 | 1.04 | 230 | 2.1 | 2.1 | 0.939 | 0.939 | 96.5 | 82.4 | 78.3 | 31.7 |
| 6ijj | 7 | 2.9 | 9678 | 0.07 | 1.04 | 213 | 2.5 | 2.5 | 0.942 | 0.942 | 95.8 | 71.6 | 88.3 | 42.6 |
| 6ijj | 8 | 2.9 | 9678 | 0.07 | 1.04 | 215 | 1.7 | 1.7 | 0.949 | 0.949 | 96.7 | 83.7 | 83.3 | 19 |
| 6ijj | C | 2.9 | 9678 | 0.07 | 1.04 | 80 | 5.6 | 2.2 | 0.850 | 0.850 | 90 | 50 | 85 | 5.9 |

| | | | | | | | | | | | | | | |
|---|---|---|---|---|---|---|---|---|---|---|---|---|---|---|
| 6ijj | D | 2.9 | 9678 | 0.07 | 1.04 | 144 | 2.0 | 2.0 | 0.930 | 0.930 | 96.5 | 84.9 | 84.7 | 19.7 |
| 6ijj | E | 2.9 | 9678 | 0.07 | 1.04 | 64 | 3.6 | 3.6 | 0.854 | 0.854 | 92.2 | 47.5 | 87.5 | 23.2 |
| 6ijj | F | 2.9 | 9678 | 0.07 | 1.04 | 164 | 2.7 | 2.7 | 0.904 | 0.904 | 95.1 | 71.2 | 83.5 | 49.6 |
| 6ijo | D | 3.3 | 9680 | 0.06 | 1.04 | 144 | 2.1 | 2.1 | 0.905 | 0.905 | 94.4 | 75 | 80.6 | 34.5 |
| 6ijo | E | 3.3 | 9680 | 0.06 | 1.04 | 64 | 4.3 | 3.7 | 0.796 | 0.811 | 87.5 | 48.2 | 68.8 | 11.4 |
| 6ir9 | C | 3.8 | 9713 | 0.0165 | 1.49 | 263 | 4.8 | 4.8 | 0.862 | 0.862 | 87.1 | 52.4 | 74.9 | 9.1 |
| 6ir9 | E | 3.8 | 9713 | 0.0165 | 1.49 | 213 | 5.2 | 5.2 | 0.814 | 0.814 | 84.5 | 32.8 | 64.8 | 21.7 |
| 6ir9 | F | 3.8 | 9713 | 0.0165 | 1.49 | 84 | 2.0 | 2.0 | 0.861 | 0.861 | 95.2 | 80 | 86.9 | 72.6 |
| 6ir9 | H | 3.8 | 9713 | 0.0165 | 1.49 | 133 | 16.6 | 13.4 | 0.486 | 0.487 | 78.9 | 6.7 | 34.6 | 6.5 |
| 6ir9 | K | 3.8 | 9713 | 0.0165 | 1.49 | 113 | 13.1 | 7.4 | 0.590 | 0.802 | 77.9 | 51.1 | 85 | 8.3 |
| 6irm | A | 3.4 | 9718 | 0.0216 | 1.07 | 98 | 1.5 | 1.5 | 0.940 | 0.940 | 98 | 87.5 | 92.9 | 98.9 |
| 6irm | B | 3.4 | 9718 | 0.0216 | 1.07 | 88 | 3.6 | 3.6 | 0.865 | 0.865 | 92 | 60.5 | 89.8 | 64.6 |
| 6irm | C | 3.4 | 9718 | 0.0216 | 1.07 | 107 | 1.6 | 1.6 | 0.899 | 0.899 | 95.3 | 83.3 | 88.8 | 55.8 |
| 6irm | D | 3.4 | 9718 | 0.0216 | 1.07 | 93 | 2.6 | 2.6 | 0.920 | 0.920 | 95.7 | 87.6 | 93.5 | 86.2 |
| 6irm | E | 3.4 | 9718 | 0.0216 | 1.07 | 95 | 1.2 | 1.2 | 0.941 | 0.941 | 98.9 | 93.6 | 92.6 | 61.4 |
| 6irm | F | 3.4 | 9718 | 0.0216 | 1.07 | 80 | 2.0 | 2.0 | 0.877 | 0.877 | 93.8 | 88 | 88.8 | 59.2 |
| 6irm | G | 3.4 | 9718 | 0.0216 | 1.07 | 107 | 1.6 | 1.6 | 0.917 | 0.917 | 97.2 | 95.2 | 91.6 | 45.9 |
| 6irm | H | 3.4 | 9718 | 0.0216 | 1.07 | 93 | 1.3 | 1.3 | 0.946 | 0.946 | 98.9 | 92.4 | 95.7 | 76.4 |
| 6iro | A | 3.4 | 9720 | 0.0253 | 1.07 | 98 | 1.3 | 1.3 | 0.927 | 0.927 | 99 | 96.9 | 92.9 | 52.7 |
| 6iro | B | 3.4 | 9720 | 0.0253 | 1.07 | 87 | 3.0 | 3.0 | 0.870 | 0.870 | 90.8 | 75.9 | 87.4 | 77.6 |
| 6iro | C | 3.4 | 9720 | 0.0253 | 1.07 | 107 | 1.6 | 1.6 | 0.937 | 0.937 | 98.1 | 87.6 | 86 | 75 |
| 6iro | D | 3.4 | 9720 | 0.0253 | 1.07 | 93 | 1.2 | 1.2 | 0.928 | 0.928 | 97.8 | 92.3 | 89.2 | 72.3 |
| 6iro | F | 3.4 | 9720 | 0.0253 | 1.07 | 86 | 3.2 | 3.2 | 0.843 | 0.843 | 86 | 81.1 | 84.9 | 71.2 |
| 6iro | G | 3.4 | 9720 | 0.0253 | 1.07 | 107 | 1.5 | 1.5 | 0.934 | 0.934 | 97.2 | 89.4 | 84.1 | 83.3 |
| 6iro | H | 3.4 | 9720 | 0.0253 | 1.07 | 93 | 1.3 | 1.3 | 0.920 | 0.920 | 96.8 | 94.4 | 92.5 | 96.5 |
| 6itc | C | 3.5 | 9731 | 0.0242 | 1.05 | 112 | 14.6 | 13.3 | 0.248 | 0.489 | 71.4 | 15 | 48.2 | 5.6 |
| 6itc | V | 3.5 | 9731 | 0.0242 | 1.05 | 116 | 14.4 | 13.6 | 0.453 | 0.537 | 74.1 | 12.8 | 55.2 | 6.2 |
| 6iyc | D | 2.6 | 9751 | 0.024 | 1.091 | 100 | 2.1 | 2.1 | 0.876 | 0.876 | 93 | 74.2 | 84 | 39.3 |
| 6j4x | C | 4.3 | 0672 | 0.016 | 1.49 | 263 | 28.0 | 20.8 | 0.344 | 0.385 | 66.9 | 9.7 | 35.7 | 4.3 |
| 6j4x | E | 4.3 | 0672 | 0.016 | 1.49 | 213 | 15.6 | 15.6 | 0.571 | 0.571 | 73.2 | 13.5 | 61.5 | 5.3 |
| 6j4x | F | 4.3 | 0672 | 0.016 | 1.49 | 84 | 8.0 | 8.0 | 0.629 | 0.641 | 84.5 | 69 | 72.6 | 8.2 |
| 6j4x | H | 4.3 | 0672 | 0.016 | 1.49 | 133 | 16.8 | 14.7 | 0.272 | 0.302 | 52.6 | 11.4 | 33.8 | 6.7 |
| 6j4y | C | 4.3 | 0673 | 0.016 | 1.49 | 263 | 21.9 | 21.9 | 0.431 | 0.435 | 73.8 | 20.1 | 48.7 | 7 |
| 6j4y | E | 4.3 | 0673 | 0.016 | 1.49 | 213 | 14.7 | 11.0 | 0.707 | 0.757 | 80.8 | 22.7 | 59.2 | 7.9 |
| 6j4y | F | 4.3 | 0673 | 0.016 | 1.49 | 84 | 4.3 | 4.3 | 0.758 | 0.758 | 85.7 | 23.6 | 77.4 | 4.6 |
| 6j4y | H | 4.3 | 0673 | 0.016 | 1.49 | 133 | 14.7 | 14.7 | 0.325 | 0.331 | 72.9 | 6.2 | 22.6 | 3.3 |
| 6j4z | C | 4.1 | 0674 | 0.016 | 1.49 | 263 | 23.1 | 20.4 | 0.341 | 0.403 | 77.6 | 6.9 | 66.5 | 5.7 |

| | | | | | | | | | | | | | | |
|---|---|---|---|---|---|---|---|---|---|---|---|---|---|---|
| 6j4z | E | 4.1 | 0674 | 0.016 | 1.49 | 213 | 12.9 | 10.5 | 0.668 | 0.668 | 76.5 | 18.4 | 60.1 | 7.8 |
| 6j4z | F | 4.1 | 0674 | 0.016 | 1.49 | 84 | 4.1 | 4.1 | 0.752 | 0.752 | 82.1 | 53.6 | 52.4 | 2.3 |
| 6j4z | H | 4.1 | 0674 | 0.016 | 1.49 | 133 | 17.6 | 15.6 | 0.267 | 0.316 | 67.7 | 11.1 | 38.3 | 3.9 |
| 6j50 | C | 4.7 | 0675 | 0.016 | 1.49 | 263 | 25.6 | 21.9 | 0.305 | 0.523 | 62 | 7.4 | 49.8 | 3.8 |
| 6j50 | E | 4.7 | 0675 | 0.016 | 1.49 | 213 | 20.9 | 19.8 | 0.266 | 0.398 | 65.3 | 7.2 | 62.4 | 4.5 |
| 6j50 | F | 4.7 | 0675 | 0.016 | 1.49 | 84 | 15.5 | 11.3 | 0.291 | 0.472 | 72.6 | 3.3 | 64.3 | 38.9 |
| 6j50 | H | 4.7 | 0675 | 0.016 | 1.49 | 133 | 16.1 | 13.3 | 0.286 | 0.339 | 66.9 | 7.9 | 25.6 | 5.9 |
| 6j51 | C | 4.2 | 0676 | 0.016 | 1.49 | 263 | 19.0 | 16.3 | 0.589 | 0.597 | 70.7 | 19.4 | 68.1 | 5 |
| 6j51 | E | 4.2 | 0676 | 0.016 | 1.49 | 213 | 18.8 | 14.8 | 0.467 | 0.573 | 81.7 | 13.2 | 58.7 | 6.4 |
| 6j51 | F | 4.2 | 0676 | 0.016 | 1.49 | 84 | 3.8 | 3.8 | 0.773 | 0.800 | 86.9 | 72.6 | 81 | 14.7 |
| 6j51 | H | 4.2 | 0676 | 0.016 | 1.49 | 133 | 16.3 | 12.3 | 0.488 | 0.613 | 68.4 | 9.9 | 29.3 | 0 |
| 6j5i | G | 3.3 | 0677 | 0.0188 | 1.38 | 272 | 13.7 | 13.7 | 0.614 | 0.614 | 74.6 | 36.5 | 54 | 38.8 |
| 6j5i | S | 3.3 | 0677 | 0.0188 | 1.38 | 187 | 9.7 | 9.0 | 0.694 | 0.694 | 78.1 | 32.9 | 72.2 | 5.9 |
| 6j5j | G | 3.5 | 0669 | 0.018 | 1.38 | 272 | 13.9 | 6.1 | 0.651 | 0.877 | 90.4 | 41.5 | 76.1 | 26.6 |
| 6j5j | H | 3.5 | 0669 | 0.018 | 1.38 | 132 | 12.0 | 9.8 | 0.642 | 0.680 | 77.3 | 23.5 | 57.6 | 10.5 |
| 6j5j | S | 3.5 | 0669 | 0.018 | 1.38 | 187 | 10.5 | 8.6 | 0.715 | 0.800 | 87.7 | 13.4 | 73.3 | 27.7 |
| 6j5u | C | 3.9 | 0681 | 0.03 | 1.091 | 172 | 14.9 | 13.7 | 0.341 | 0.351 | 61.6 | 5.7 | 28.5 | 6.1 |
| 6j5v | C | 4.3 | 0682 | 0.022 | 1.091 | 172 | 12.9 | 12.9 | 0.284 | 0.330 | 64 | 5.5 | 50 | 9.3 |
| 6j6g | l | 3.2 | 0686 | 0.019 | 1.33 | 83 | 2.5 | 2.5 | 0.834 | 0.834 | 92.8 | 61 | 50.6 | 7.1 |
| 6j6g | Q | 3.2 | 0686 | 0.019 | 1.33 | 292 | 23.4 | 21.7 | 0.480 | 0.482 | 43.5 | 9.4 | 58.6 | 13.5 |
| 6j6g | R | 3.2 | 0686 | 0.019 | 1.33 | 261 | 4.2 | 4.2 | 0.932 | 0.932 | 93.5 | 68.9 | 78.9 | 7.8 |
| 6j6g | T | 3.2 | 0686 | 0.019 | 1.33 | 157 | 1.8 | 1.8 | 0.917 | 0.917 | 95.5 | 82 | 84.1 | 24.2 |
| 6j6h | l | 3.6 | 0687 | 0.024 | 1.33 | 83 | 12.2 | 10.1 | 0.664 | 0.664 | 88 | 21.9 | 57.8 | 4.2 |
| 6j6h | Q | 3.6 | 0687 | 0.024 | 1.33 | 292 | 23.6 | 22.4 | 0.474 | 0.474 | 47.6 | 8.6 | 44.2 | 6.2 |
| 6j6h | R | 3.6 | 0687 | 0.024 | 1.33 | 261 | 4.2 | 4.2 | 0.859 | 0.859 | 88.1 | 44.3 | 69.3 | 15.5 |
| 6j6h | T | 3.6 | 0687 | 0.024 | 1.33 | 157 | 2.0 | 2.0 | 0.911 | 0.911 | 95.5 | 76.7 | 80.3 | 29.4 |
| 6j6i | A | 3.7 | 0688 | 0.013 | 1.091 | 193 | 17.1 | 16.5 | 0.280 | 0.369 | 68.4 | 12.9 | 58 | 3.6 |
| 6j6n | Q | 3.9 | 0691 | 0.028 | 1.33 | 292 | 23.1 | 21.1 | 0.393 | 0.393 | 44.5 | 9.2 | 53.1 | 4.5 |
| 6j6n | R | 3.9 | 0691 | 0.028 | 1.33 | 261 | 21.8 | 16.2 | 0.251 | 0.448 | 70.9 | 8.6 | 33.7 | 3.4 |
| 6j6n | T | 3.9 | 0691 | 0.028 | 1.33 | 157 | 7.8 | 5.0 | 0.718 | 0.736 | 80.3 | 24.6 | 68.2 | 49.5 |
| 6j6q | Q | 3.7 | 0692 | 0.0116 | 1.33 | 292 | 23.0 | 22.1 | 0.278 | 0.475 | 44.9 | 9.2 | 61.6 | 12.2 |
| 6j6q | R | 3.7 | 0692 | 0.0116 | 1.33 | 261 | 20.5 | 8.8 | 0.601 | 0.647 | 85.8 | 24.6 | 64.4 | 5.4 |
| 6j6q | T | 3.7 | 0692 | 0.0116 | 1.33 | 157 | 3.0 | 3.0 | 0.877 | 0.877 | 90.4 | 61.3 | 38.9 | 4.9 |
| 6j8g | B | 3.2 | 9781 | 0.07 | 1.091 | 173 | 2.5 | 2.5 | 0.905 | 0.915 | 94.8 | 69.5 | 86.1 | 7.4 |
| 6j8i | B | 3.2 | 9782 | 0.06 | 1.091 | 173 | 12.6 | 10.1 | 0.610 | 0.630 | 87.9 | 37.5 | 80.9 | 4.3 |
| 6j9e | A | 3.4 | 9785 | 0.013 | 1.014 | 211 | 4.0 | 4.0 | 0.889 | 0.889 | 91.5 | 76.7 | 89.6 | 21.7 |
| 6j9e | B | 3.4 | 9785 | 0.013 | 1.014 | 208 | 2.7 | 2.7 | 0.893 | 0.893 | 89.9 | 81.3 | 74 | 25.3 |

| | | | | | | | | | | | | | | |
|---|---|---|---|---|---|---|---|---|---|---|---|---|---|---|
| 6j9e | E | 3.4 | 9785 | 0.013 | 1.014 | 65 | 2.7 | 2.7 | 0.783 | 0.783 | 87.7 | 71.9 | 89.2 | 84.5 |
| 6j9e | J | 3.4 | 9785 | 0.013 | 1.014 | 66 | 1.6 | 1.6 | 0.856 | 0.856 | 97 | 92.2 | 83.3 | 36.4 |
| 6j9f | A | 4 | 9786 | 0.013 | 1.014 | 215 | 6.1 | 6.1 | 0.842 | 0.842 | 87 | 53.5 | 82.3 | 18.6 |
| 6j9f | B | 4 | 9786 | 0.013 | 1.014 | 203 | 19.2 | 13.9 | 0.600 | 0.618 | 74.9 | 13.8 | 67 | 19.9 |
| 6j9f | E | 4 | 9786 | 0.013 | 1.014 | 64 | 3.9 | 3.9 | 0.680 | 0.680 | 81.2 | 48.1 | 73.4 | 6.4 |
| 6j9f | J | 4 | 9786 | 0.013 | 1.014 | 66 | 6.1 | 6.1 | 0.761 | 0.761 | 89.4 | 47.5 | 69.7 | 37 |
| 6jbq | A | 4 | 9792 | 0.03 | 1.014 | 216 | 19.4 | 11.7 | 0.575 | 0.594 | 74.5 | 26.1 | 55.6 | 5 |
| 6jbq | B | 4 | 9792 | 0.03 | 1.014 | 219 | 11.2 | 11.0 | 0.599 | 0.825 | 84.9 | 42.5 | 81.3 | 29.2 |
| 6jmr | E | 4.1 | 9850 | 0.035 | 1.49 | 215 | 18.9 | 17.9 | 0.317 | 0.394 | 69.8 | 10 | 40.9 | 9.1 |
| 6jmr | F | 4.1 | 9850 | 0.035 | 1.49 | 218 | 16.3 | 16.3 | 0.405 | 0.423 | 67.4 | 10.9 | 31.2 | 4.4 |
| 6jnx | A | 4.1 | 9852 | 0.026 | 1.307 | 219 | 11.3 | 11.3 | 0.552 | 0.552 | 77.6 | 36.5 | 73.1 | 17.5 |
| 6jnx | B | 4.1 | 9852 | 0.026 | 1.307 | 218 | 14.6 | 14.3 | 0.578 | 0.589 | 73.4 | 21.9 | 65.1 | 7.7 |
| 6jo5 | 3 | 2.9 | 9853 | 0.05 | 1.12 | 202 | 2.9 | 2.9 | 0.921 | 0.921 | 93.6 | 70.4 | 81.7 | 27.9 |
| 6jo5 | 4 | 2.9 | 9853 | 0.05 | 1.12 | 203 | 2.2 | 2.2 | 0.919 | 0.919 | 93.6 | 79.5 | 76.8 | 60.9 |
| 6jo5 | 5 | 2.9 | 9853 | 0.05 | 1.12 | 223 | 2.0 | 2.0 | 0.933 | 0.933 | 94.2 | 79.5 | 80.7 | 44.4 |
| 6jo5 | 6 | 2.9 | 9853 | 0.05 | 1.12 | 229 | 1.8 | 1.8 | 0.952 | 0.952 | 97.4 | 81.6 | 83.4 | 38.2 |
| 6jo5 | 8 | 2.9 | 9853 | 0.05 | 1.12 | 217 | 1.8 | 1.8 | 0.944 | 0.944 | 96.3 | 85.2 | 90.3 | 64.8 |
| 6jo5 | C | 2.9 | 9853 | 0.05 | 1.12 | 80 | 1.7 | 1.7 | 0.877 | 0.877 | 95 | 89.5 | 60 | 33.3 |
| 6jo5 | D | 2.9 | 9853 | 0.05 | 1.12 | 144 | 1.5 | 1.5 | 0.941 | 0.941 | 97.9 | 87.9 | 88.9 | 62.5 |
| 6jo5 | E | 2.9 | 9853 | 0.05 | 1.12 | 61 | 1.7 | 1.7 | 0.868 | 0.868 | 95.1 | 93.1 | 90.2 | 45.5 |
| 6jo5 | F | 2.9 | 9853 | 0.05 | 1.12 | 165 | 1.6 | 1.6 | 0.952 | 0.952 | 97 | 86.2 | 88.5 | 66.4 |
| 6jo5 | L | 2.9 | 9853 | 0.05 | 1.12 | 118 | 2.2 | 2.2 | 0.887 | 0.887 | 94.9 | 82.1 | 84.7 | 74 |
| 6jo6 | 1 | 2.9 | 9854 | 0.05 | 1.12 | 194 | 1.8 | 1.8 | 0.937 | 0.937 | 95.4 | 90.3 | 86.1 | 82.6 |
| 6jo6 | 3 | 2.9 | 9854 | 0.05 | 1.12 | 202 | 1.7 | 1.7 | 0.947 | 0.947 | 96.5 | 87.7 | 83.2 | 56 |
| 6jo6 | 4 | 2.9 | 9854 | 0.05 | 1.12 | 203 | 3.6 | 3.6 | 0.894 | 0.894 | 91.6 | 67.2 | 71.9 | 60.3 |
| 6jo6 | 5 | 2.9 | 9854 | 0.05 | 1.12 | 223 | 2.7 | 2.7 | 0.930 | 0.930 | 96 | 80.4 | 65.9 | 42.9 |
| 6jo6 | 6 | 2.9 | 9854 | 0.05 | 1.12 | 229 | 2.3 | 2.3 | 0.931 | 0.931 | 94.8 | 75.6 | 78.2 | 30.7 |
| 6jo6 | 7 | 2.9 | 9854 | 0.05 | 1.12 | 212 | 2.1 | 2.1 | 0.928 | 0.928 | 94.3 | 86.5 | 89.2 | 57.1 |
| 6jo6 | 8 | 2.9 | 9854 | 0.05 | 1.12 | 217 | 2.1 | 2.1 | 0.954 | 0.954 | 97.2 | 86.3 | 88.5 | 77.6 |
| 6jo6 | C | 2.9 | 9854 | 0.05 | 1.12 | 80 | 1.7 | 1.7 | 0.889 | 0.889 | 90 | 94.4 | 68.8 | 40 |
| 6jo6 | D | 2.9 | 9854 | 0.05 | 1.12 | 144 | 4.2 | 4.2 | 0.877 | 0.877 | 91.7 | 78.8 | 94.4 | 74.3 |
| 6jo6 | E | 2.9 | 9854 | 0.05 | 1.12 | 61 | 1.4 | 1.4 | 0.879 | 0.879 | 96.7 | 96.6 | 59 | 8.3 |
| 6jo6 | F | 2.9 | 9854 | 0.05 | 1.12 | 165 | 2.4 | 2.4 | 0.922 | 0.922 | 93.9 | 83.2 | 90.9 | 91.3 |
| 6jo6 | G | 2.9 | 9854 | 0.05 | 1.12 | 68 | 7.5 | 5.7 | 0.663 | 0.677 | 76.5 | 46.2 | 86.8 | 81.4 |
| 6jo6 | L | 2.9 | 9854 | 0.05 | 1.12 | 118 | 2.4 | 2.4 | 0.902 | 0.902 | 94.1 | 82.9 | 81.4 | 77.1 |
| 6jxr | e | 3.7 | 9895 | 0.0291 | 1.057 | 123 | 7.7 | 4.4 | 0.832 | 0.839 | 87.8 | 20.4 | 83.7 | 11.7 |
| 6k0a | C | 4.6 | 30036 | 0.0206 | 1.09 | 226 | 18.7 | 17.0 | 0.358 | 0.358 | 58.4 | 5.3 | 49.1 | 7.2 |

| | | | | | | | | | | | | | | |
|---|---|---|---|---|---|---|---|---|---|---|---|---|---|---|
| 6k0a | D | 4.6 | 30036 | 0.0206 | 1.09 | 226 | 21.3 | 16.8 | 0.274 | 0.343 | 64.6 | 8.9 | 43.4 | 7.1 |
| 6k0a | I | 4.6 | 30036 | 0.0206 | 1.09 | 116 | 13.5 | 12.3 | 0.314 | 0.323 | 62.1 | 13.9 | 54.3 | 6.3 |
| 6k0a | J | 4.6 | 30036 | 0.0206 | 1.09 | 116 | 14.0 | 12.8 | 0.313 | 0.354 | 56.9 | 6.1 | 56 | 7.7 |
| 6k0b | C | 4.3 | 9900 | 0.02 | 1.32 | 231 | 9.2 | 8.4 | 0.621 | 0.672 | 81 | 33.7 | 67.5 | 8.3 |
| 6k0b | D | 4.3 | 9900 | 0.02 | 1.32 | 231 | 18.6 | 17.8 | 0.349 | 0.455 | 78.8 | 11.5 | 71 | 11 |
| 6k0b | I | 4.3 | 9900 | 0.02 | 1.32 | 116 | 14.3 | 12.5 | 0.297 | 0.334 | 69.8 | 11.1 | 54.3 | 15.9 |
| 6k0b | J | 4.3 | 9900 | 0.02 | 1.32 | 116 | 14.0 | 11.6 | 0.367 | 0.367 | 67.2 | 15.4 | 55.2 | 3.1 |
| 6k15 | G | 3.4 | 9905 | 0.02 | 1.07 | 246 | 14.5 | 13.0 | 0.716 | 0.723 | 85 | 21.1 | 79.7 | 15.3 |
| 6k15 | X | 3.4 | 9905 | 0.02 | 1.07 | 147 | 13.7 | 13.7 | 0.335 | 0.339 | 72.8 | 14 | 38.1 | 5.4 |
| 6k4y | A | 3.8 | 9916 | 0.0278 | 1.307 | 219 | 26.9 | 18.1 | 0.473 | 0.477 | 71.7 | 24.8 | 84.5 | 27 |
| 6k4y | B | 3.8 | 9916 | 0.0278 | 1.307 | 217 | 12.0 | 10.5 | 0.652 | 0.657 | 83.4 | 45.3 | 51.6 | 14.3 |
| 6kac | e | 2.7 | 9955 | 0.015 | 1.04 | 76 | 2.4 | 2.4 | 0.870 | 0.870 | 93.4 | 67.6 | 80.3 | 24.6 |
| 6kac | E | 2.7 | 9955 | 0.015 | 1.04 | 76 | 2.1 | 2.1 | 0.872 | 0.872 | 93.4 | 95.8 | 80.3 | 68.9 |
| 6kac | n | 2.7 | 9955 | 0.015 | 1.04 | 219 | 2.3 | 2.3 | 0.925 | 0.925 | 95 | 71.6 | 84.5 | 36.8 |
| 6kac | N | 2.7 | 9955 | 0.015 | 1.04 | 219 | 2.2 | 2.2 | 0.931 | 0.931 | 95.4 | 64.1 | 82.2 | 31.1 |
| 6kac | o | 2.7 | 9955 | 0.015 | 1.04 | 240 | 5.7 | 2.1 | 0.925 | 0.933 | 94.2 | 71.2 | 85 | 62.3 |
| 6kac | O | 2.7 | 9955 | 0.015 | 1.04 | 240 | 4.9 | 2.4 | 0.910 | 0.938 | 92.9 | 80.3 | 90.4 | 11.5 |
| 6kac | p | 2.7 | 9955 | 0.015 | 1.04 | 188 | 1.7 | 1.7 | 0.936 | 0.936 | 96.8 | 88.5 | 88.8 | 37.7 |
| 6kac | P | 2.7 | 9955 | 0.015 | 1.04 | 188 | 3.4 | 3.4 | 0.919 | 0.919 | 93.6 | 77.3 | 87.8 | 14.5 |
| 6kac | q | 2.7 | 9955 | 0.015 | 1.04 | 148 | 1.7 | 1.7 | 0.931 | 0.931 | 95.9 | 86.6 | 87.8 | 42.3 |
| 6kac | Q | 2.7 | 9955 | 0.015 | 1.04 | 148 | 1.2 | 1.2 | 0.952 | 0.952 | 97.3 | 97.2 | 88.5 | 34.4 |
| 6kac | y | 2.7 | 9955 | 0.015 | 1.04 | 221 | 2.3 | 2.3 | 0.916 | 0.916 | 94.6 | 78 | 87.8 | 68 |
| 6kac | Y | 2.7 | 9955 | 0.015 | 1.04 | 221 | 2.6 | 2.6 | 0.931 | 0.931 | 93.7 | 79.7 | 85.5 | 55 |
| 6kac | z | 2.7 | 9955 | 0.015 | 1.04 | 61 | 1.9 | 1.9 | 0.884 | 0.884 | 96.7 | 86.4 | 91.8 | 94.6 |
| 6kac | Z | 2.7 | 9955 | 0.015 | 1.04 | 61 | 2.2 | 2.2 | 0.862 | 0.862 | 93.4 | 91.2 | 93.4 | 47.4 |
| 6kiv | T | 4 | 9999 | 0.011 | 1.09 | 176 | 17.2 | 14.5 | 0.288 | 0.342 | 55.1 | 7.2 | 25.6 | 4.4 |
| 6kiz | T | 4.5 | 0695 | 0.005 | 1.1 | 176 | 18.2 | 16.2 | 0.307 | 0.307 | 40.9 | 11.1 | 9.7 | 5.9 |
| 6m7j | A | 4.4 | 9047 | 0.3 | 1.3 | 225 | 23.6 | 22.4 | 0.277 | 0.468 | 68.9 | 12.3 | 55.6 | 9.6 |
| 6m7j | B | 4.4 | 9047 | 0.3 | 1.3 | 237 | 27.2 | 19.0 | 0.333 | 0.340 | 74.3 | 5.7 | 30.4 | 5.6 |
| 6m7j | E | 4.4 | 9047 | 0.3 | 1.3 | 83 | 6.9 | 5.2 | 0.690 | 0.714 | 74.7 | 22.6 | 45.8 | 2.6 |
| 6-Mar | H | 4.5 | 9062 | 0.04 | 1.2 | 135 | 17.9 | 16.9 | 0.419 | 0.419 | 68.9 | 2.2 | 43.7 | 3.4 |
| 6-Mar | L | 4.5 | 9062 | 0.04 | 1.2 | 114 | 15.7 | 15.0 | 0.258 | 0.385 | 57.9 | 9.1 | 28.1 | 6.2 |
| 6-Mar | M | 4.5 | 9062 | 0.04 | 1.2 | 135 | 19.0 | 17.9 | 0.343 | 0.359 | 66.7 | 6.7 | 57.8 | 16.7 |
| 6-Mar | N | 4.5 | 9062 | 0.04 | 1.2 | 112 | 16.2 | 14.6 | 0.244 | 0.304 | 63.4 | 15.5 | 33.9 | 10.5 |
| 6mb3 | A | 3.4 | 9065 | 0.056 | 1.03 | 120 | 5.5 | 5.5 | 0.825 | 0.825 | 87.5 | 60 | 80 | 7.3 |
| 6mb3 | B | 3.4 | 9065 | 0.056 | 1.03 | 120 | 1.1 | 1.1 | 0.946 | 0.946 | 97.5 | 94 | 81.7 | 7.1 |
| 6mb3 | C | 3.4 | 9065 | 0.056 | 1.03 | 120 | 1.5 | 1.5 | 0.942 | 0.942 | 99.2 | 93.3 | 84.2 | 4 |

| | | | | | | | | | | | | | | |
|---|---|---|---|---|---|---|---|---|---|---|---|---|---|---|
| 6mb3 | D | 3.4 | 9065 | 0.056 | 1.03 | 120 | 3.4 | 3.4 | 0.883 | 0.883 | 93.3 | 68.8 | 78.3 | 17 |
| 6mb3 | F | 3.4 | 9065 | 0.056 | 1.03 | 120 | 1.9 | 1.9 | 0.911 | 0.911 | 95 | 87.7 | 84.2 | 8.9 |
| 6mb3 | G | 3.4 | 9065 | 0.056 | 1.03 | 120 | 1.6 | 1.6 | 0.925 | 0.925 | 97.5 | 89.7 | 95.8 | 13.9 |
| 6mb3 | H | 3.4 | 9065 | 0.056 | 1.03 | 120 | 13.6 | 9.9 | 0.751 | 0.760 | 85 | 56.9 | 80 | 9.4 |
| 6mb3 | I | 3.4 | 9065 | 0.056 | 1.03 | 120 | 15.0 | 12.7 | 0.441 | 0.618 | 79.2 | 9.5 | 56.7 | 8.8 |
| 6mb3 | J | 3.4 | 9065 | 0.056 | 1.03 | 120 | 15.9 | 13.0 | 0.357 | 0.506 | 80.8 | 6.2 | 61.7 | 8.1 |
| 6mb3 | K | 3.4 | 9065 | 0.056 | 1.03 | 112 | 13.3 | 10.2 | 0.385 | 0.708 | 81.2 | 29.7 | 61.6 | 8.7 |
| 6mb3 | L | 3.4 | 9065 | 0.056 | 1.03 | 112 | 15.7 | 14.1 | 0.365 | 0.514 | 80.4 | 6.7 | 36.6 | 7.3 |
| 6mb3 | M | 3.4 | 9065 | 0.056 | 1.03 | 112 | 2.0 | 2.0 | 0.865 | 0.865 | 93.8 | 71.4 | 77.7 | 18.4 |
| 6mb3 | N | 3.4 | 9065 | 0.056 | 1.03 | 112 | 2.4 | 2.4 | 0.892 | 0.892 | 95.5 | 66.4 | 84.8 | 12.6 |
| 6mb3 | O | 3.4 | 9065 | 0.056 | 1.03 | 112 | 14.2 | 10.2 | 0.592 | 0.719 | 86.6 | 10.3 | 75.9 | 9.4 |
| 6mb3 | P | 3.4 | 9065 | 0.056 | 1.03 | 112 | 2.5 | 2.5 | 0.894 | 0.894 | 92.9 | 86.5 | 83.9 | 29.8 |
| 6mb3 | Q | 3.4 | 9065 | 0.056 | 1.03 | 112 | 1.9 | 1.9 | 0.901 | 0.901 | 95.5 | 75.7 | 77.7 | 40.2 |
| 6mb3 | R | 3.4 | 9065 | 0.056 | 1.03 | 112 | 13.5 | 13.5 | 0.279 | 0.417 | 71.4 | 10 | 55.4 | 12.9 |
| 6mb3 | S | 3.4 | 9065 | 0.056 | 1.03 | 112 | 15.0 | 12.9 | 0.501 | 0.535 | 67.9 | 11.8 | 38.4 | 4.7 |
| 6mcb | C | 3.4 | 9066 | 0.28 | 0.83 | 123 | 6.4 | 4.5 | 0.763 | 0.823 | 79.7 | 34.7 | 74 | 5.5 |
| 6mcc | C | 3.9 | 9067 | 0.443 | 1.15 | 130 | 3.4 | 3.4 | 0.862 | 0.862 | 93.1 | 59.5 | 81.5 | 4.7 |
| 6mdo | F | 3.9 | 9102 | 0.0135 | 1.31 | 241 | 13.0 | 11.6 | 0.628 | 0.649 | 80.9 | 15.9 | 69.3 | 7.2 |
| 6mdp | F | 3.8 | 9103 | 0.0135 | 1.31 | 243 | 6.2 | 6.2 | 0.777 | 0.777 | 84.4 | 24.9 | 67.1 | 7.4 |
| 6meo | A | 3.9 | 9108 | 0.042 | 1.059 | 176 | 11.6 | 10.6 | 0.659 | 0.659 | 73.9 | 42.3 | 41.5 | 8.2 |
| 6mhg | A | 3.6 | 9114 | 0.81 | 1.03 | 120 | 16.1 | 12.5 | 0.459 | 0.544 | 81.7 | 39.8 | 67.5 | 6.2 |
| 6mhg | B | 3.6 | 9114 | 0.81 | 1.03 | 120 | 10.3 | 9.5 | 0.641 | 0.689 | 78.3 | 52.1 | 68.3 | 11 |
| 6mhg | C | 3.6 | 9114 | 0.81 | 1.03 | 120 | 5.6 | 4.1 | 0.809 | 0.894 | 91.7 | 82.7 | 70.8 | 12.9 |
| 6mhg | D | 3.6 | 9114 | 0.81 | 1.03 | 120 | 5.9 | 5.9 | 0.833 | 0.833 | 94.2 | 88.5 | 86.7 | 7.7 |
| 6mhg | F | 3.6 | 9114 | 0.81 | 1.03 | 120 | 1.2 | 1.2 | 0.934 | 0.934 | 98.3 | 99.2 | 95 | 5.3 |
| 6mhg | G | 3.6 | 9114 | 0.81 | 1.03 | 120 | 1.3 | 1.3 | 0.923 | 0.923 | 100 | 100 | 90.8 | 5.5 |
| 6mhg | H | 3.6 | 9114 | 0.81 | 1.03 | 120 | 15.3 | 12.7 | 0.283 | 0.358 | 56.7 | 11.8 | 30 | 5.6 |
| 6mhg | I | 3.6 | 9114 | 0.81 | 1.03 | 120 | 2.9 | 2.9 | 0.908 | 0.908 | 95 | 75.4 | 85.8 | 13.6 |
| 6mhg | J | 3.6 | 9114 | 0.81 | 1.03 | 120 | 4.8 | 2.7 | 0.808 | 0.891 | 92.5 | 82 | 80 | 21.9 |
| 6mhg | K | 3.6 | 9114 | 0.81 | 1.03 | 120 | 15.5 | 13.3 | 0.458 | 0.458 | 75.8 | 7.7 | 61.7 | 5.4 |
| 6mhg | N | 3.6 | 9114 | 0.81 | 1.03 | 112 | 16.0 | 12.5 | 0.290 | 0.441 | 72.3 | 11.1 | 31.2 | 5.7 |
| 6mhg | O | 3.6 | 9114 | 0.81 | 1.03 | 112 | 15.5 | 12.9 | 0.422 | 0.486 | 75.9 | 18.8 | 51.8 | 5.2 |
| 6mhg | P | 3.6 | 9114 | 0.81 | 1.03 | 112 | 8.1 | 5.6 | 0.785 | 0.822 | 84.8 | 37.9 | 49.1 | 3.6 |
| 6mhg | Q | 3.6 | 9114 | 0.81 | 1.03 | 112 | 13.0 | 7.5 | 0.717 | 0.721 | 90.2 | 7.9 | 66.1 | 9.5 |
| 6mhg | R | 3.6 | 9114 | 0.81 | 1.03 | 112 | 6.0 | 3.2 | 0.775 | 0.859 | 92.9 | 45.2 | 82.1 | 7.6 |
| 6mhg | S | 3.6 | 9114 | 0.81 | 1.03 | 112 | 2.5 | 2.5 | 0.868 | 0.868 | 92 | 60.2 | 89.3 | 11 |
| 6mhg | T | 3.6 | 9114 | 0.81 | 1.03 | 112 | 3.7 | 3.7 | 0.835 | 0.835 | 90.2 | 55.4 | 78.6 | 8 |

| | | | | | | | | | | | | | | |
|---|---|---|---|---|---|---|---|---|---|---|---|---|---|---|
| 6mhg | U | 3.6 | 9114 | 0.81 | 1.03 | 112 | 6.7 | 5.2 | 0.786 | 0.840 | 92 | 42.7 | 70.5 | 7.6 |
| 6mhg | V | 3.6 | 9114 | 0.81 | 1.03 | 112 | 13.1 | 12.7 | 0.438 | 0.448 | 72.3 | 14.8 | 51.8 | 10.3 |
| 6mhu | A | 4 | 9118 | 0.04 | 1.23 | 230 | 11.8 | 10.2 | 0.623 | 0.653 | 78.7 | 26 | 64.8 | 8.7 |
| 6mhu | B | 4 | 9118 | 0.04 | 1.23 | 230 | 18.1 | 14.1 | 0.525 | 0.528 | 83 | 12.6 | 69.6 | 6.2 |
| 6mhz | A | 4.1 | 9124 | 0.05 | 1.23 | 235 | 18.7 | 16.2 | 0.293 | 0.394 | 69.8 | 6.1 | 71.9 | 10.7 |
| 6mhz | B | 4.1 | 9124 | 0.05 | 1.23 | 235 | 20.5 | 14.4 | 0.282 | 0.420 | 75.3 | 7.9 | 57.4 | 5.9 |
| 6mi7 | A | 4.2 | 9125 | 0.085 | 1.23 | 230 | 9.9 | 9.9 | 0.697 | 0.697 | 83.5 | 42.7 | 77.4 | 5.6 |
| 6mi7 | B | 4.2 | 9125 | 0.085 | 1.23 | 230 | 9.0 | 5.9 | 0.684 | 0.833 | 81.7 | 41 | 72.6 | 9 |
| 6mi8 | A | 4.3 | 9126 | 0.045 | 1.06 | 235 | 15.7 | 15.7 | 0.325 | 0.569 | 68.1 | 10 | 72.8 | 16.4 |
| 6mi8 | B | 4.3 | 9126 | 0.045 | 1.06 | 235 | 18.9 | 17.5 | 0.316 | 0.541 | 80 | 8.5 | 78.7 | 7.6 |
| 6mjz | H | 4.3 | 9135 | 1.67 | 1.1 | 120 | 15.4 | 13.6 | 0.332 | 0.375 | 60 | 6.9 | 42.5 | 2 |
| 6mjz | L | 4.3 | 9135 | 1.67 | 1.1 | 108 | 13.2 | 12.9 | 0.287 | 0.509 | 66.7 | 4.2 | 55.6 | 3.3 |
| 6mlm | E | 3.5 | 9139 | 0.058 | 1.02 | 122 | 15.1 | 8.1 | 0.363 | 0.729 | 85.2 | 19.2 | 77.9 | 15.8 |
| 6mlm | F | 3.5 | 9139 | 0.058 | 1.02 | 122 | 13.4 | 12.6 | 0.505 | 0.670 | 82.8 | 5 | 77.9 | 11.6 |
| 6mlm | G | 3.5 | 9139 | 0.058 | 1.02 | 106 | 13.9 | 13.6 | 0.270 | 0.326 | 64.2 | 8.8 | 31.1 | 18.2 |
| 6mlm | I | 3.5 | 9139 | 0.058 | 1.02 | 106 | 13.9 | 12.4 | 0.270 | 0.314 | 65.1 | 11.6 | 38.7 | 4.9 |
| 6mlm | K | 3.5 | 9139 | 0.058 | 1.02 | 122 | 14.5 | 14.1 | 0.718 | 0.718 | 83.6 | 28.4 | 63.1 | 3.9 |
| 6mlm | L | 3.5 | 9139 | 0.058 | 1.02 | 106 | 15.4 | 13.6 | 0.321 | 0.380 | 67.9 | 9.7 | 47.2 | 6 |
| 6mph | 1 | 3.8 | 9189 | 1 | 1.06 | 118 | 16.8 | 14.6 | 0.298 | 0.306 | 61.9 | 9.6 | 41.5 | 2 |
| 6mph | 2 | 3.8 | 9189 | 1 | 1.06 | 112 | 15.1 | 13.3 | 0.313 | 0.313 | 62.5 | 11.4 | 30.4 | 5.9 |
| 6mph | 3 | 3.8 | 9189 | 1 | 1.06 | 118 | 16.1 | 15.2 | 0.299 | 0.299 | 61.9 | 9.6 | 27.1 | 3.1 |
| 6mph | 4 | 3.8 | 9189 | 1 | 1.06 | 112 | 16.2 | 14.7 | 0.272 | 0.358 | 58.9 | 7.6 | 25.9 | 17.2 |
| 6mph | 6 | 3.8 | 9189 | 1 | 1.06 | 132 | 6.2 | 4.7 | 0.752 | 0.779 | 81.1 | 29.9 | 72.7 | 5.2 |
| 6mph | a | 3.8 | 9189 | 1 | 1.06 | 103 | 14.9 | 13.1 | 0.245 | 0.317 | 69.9 | 4.2 | 32 | 3 |
| 6mph | D | 3.8 | 9189 | 1 | 1.06 | 132 | 5.5 | 5.5 | 0.764 | 0.764 | 81.1 | 35.5 | 70.5 | 5.4 |
| 6mph | E | 3.8 | 9189 | 1 | 1.06 | 132 | 5.8 | 5.8 | 0.768 | 0.768 | 80.3 | 50.9 | 65.9 | 9.2 |
| 6mph | f | 3.8 | 9189 | 1 | 1.06 | 129 | 7.0 | 5.8 | 0.798 | 0.806 | 85.3 | 60.9 | 79.1 | 4.9 |
| 6mph | g | 3.8 | 9189 | 1 | 1.06 | 129 | 3.0 | 3.0 | 0.900 | 0.900 | 95.3 | 57.7 | 90.7 | 10.3 |
| 6mph | h | 3.8 | 9189 | 1 | 1.06 | 102 | 10.9 | 10.9 | 0.676 | 0.682 | 79.4 | 38.3 | 81.4 | 7.2 |
| 6mph | H | 3.8 | 9189 | 1 | 1.06 | 118 | 14.7 | 14.7 | 0.258 | 0.292 | 61 | 1.4 | 28.8 | 5.9 |
| 6mph | i | 3.8 | 9189 | 1 | 1.06 | 102 | 5.4 | 3.0 | 0.834 | 0.834 | 92.2 | 45.7 | 75.5 | 6.5 |
| 6mph | L | 3.8 | 9189 | 1 | 1.06 | 112 | 14.6 | 13.6 | 0.246 | 0.302 | 65.2 | 2.7 | 25 | 17.9 |
| 6mph | M | 3.8 | 9189 | 1 | 1.06 | 132 | 16.1 | 16.1 | 0.280 | 0.306 | 71.2 | 4.3 | 26.5 | 5.7 |
| 6mph | N | 3.8 | 9189 | 1 | 1.06 | 103 | 13.5 | 12.8 | 0.310 | 0.327 | 74.8 | 6.5 | 30.1 | 0 |
| 6mph | Q | 3.8 | 9189 | 1 | 1.06 | 129 | 4.0 | 4.0 | 0.872 | 0.872 | 90.7 | 65 | 75.2 | 13.4 |
| 6mph | R | 3.8 | 9189 | 1 | 1.06 | 102 | 13.5 | 13.2 | 0.430 | 0.441 | 81.4 | 4.8 | 62.7 | 6.2 |
| 6mph | X | 3.8 | 9189 | 1 | 1.06 | 132 | 20.2 | 16.8 | 0.275 | 0.357 | 69.7 | 7.6 | 28.8 | 2.6 |

| | | | | | | | | | | | | | | |
|---|---|---|---|---|---|---|---|---|---|---|---|---|---|---|
| 6mph | Y | 3.8 | 9189 | 1 | 1.06 | 132 | 17.1 | 15.8 | 0.336 | 0.336 | 75 | 7.1 | 32.6 | 7 |
| 6mph | Z | 3.8 | 9189 | 1 | 1.06 | 103 | 16.0 | 15.1 | 0.255 | 0.385 | 72.8 | 6.7 | 37.9 | 5.1 |
| 6mpu | C | 3.4 | 9191 | 0.0595 | 1.15 | 293 | 24.6 | 23.6 | 0.294 | 0.475 | 81.6 | 6.7 | 65.5 | 9.4 |
| 6mpu | L | 3.4 | 9191 | 0.0595 | 1.15 | 189 | 16.3 | 16.1 | 0.319 | 0.371 | 69.3 | 7.6 | 54.5 | 3.9 |
| 6mur | D | 3.1 | 9253 | 0.025 | 1.089 | 287 | 3.4 | 3.4 | 0.921 | 0.921 | 93 | 57.7 | 83.6 | 29.6 |
| 6mur | E | 3.1 | 9253 | 0.025 | 1.089 | 286 | 1.4 | 1.4 | 0.959 | 0.959 | 97.6 | 93.5 | 87.1 | 41 |
| 6mus | D | 3.6 | 9254 | 0.02 | 1.089 | 287 | 15.5 | 15.5 | 0.731 | 0.731 | 80.5 | 42 | 71.4 | 20.5 |
| 6mus | E | 3.6 | 9254 | 0.02 | 1.089 | 279 | 7.5 | 7.5 | 0.872 | 0.879 | 88.2 | 57.7 | 83.9 | 18.8 |
| 6mus | K | 3.6 | 9254 | 0.02 | 1.089 | 275 | 10.9 | 10.9 | 0.726 | 0.726 | 80 | 28.6 | 62.5 | 12.2 |
| 6mut | C | 3.1 | 9255 | 0.02 | 1.089 | 276 | 3.3 | 3.3 | 0.903 | 0.903 | 91.3 | 61.9 | 84.8 | 47.4 |
| 6mut | D | 3.1 | 9255 | 0.02 | 1.089 | 267 | 9.1 | 4.4 | 0.878 | 0.894 | 89.5 | 42.3 | 70.4 | 21.3 |
| 6mut | E | 3.1 | 9255 | 0.02 | 1.089 | 286 | 1.4 | 1.4 | 0.964 | 0.964 | 97.2 | 91.4 | 89.5 | 35.9 |
| 6muu | C | 3 | 9256 | 0.02 | 1.089 | 276 | 4.0 | 4.0 | 0.930 | 0.930 | 93.1 | 68.1 | 88.4 | 40.2 |
| 6muu | D | 3 | 9256 | 0.02 | 1.089 | 283 | 3.7 | 3.7 | 0.911 | 0.911 | 91.2 | 70.5 | 77.7 | 37.7 |
| 6muu | E | 3 | 9256 | 0.02 | 1.089 | 285 | 2.2 | 2.2 | 0.959 | 0.959 | 97.2 | 88.1 | 92.3 | 58.6 |
| 6muv | A | 3.8 | 9257 | 6 | 1.31 | 246 | 2.7 | 2.7 | 0.903 | 0.903 | 91.9 | 77 | 89 | 22.8 |
| 6muv | B | 3.8 | 9257 | 6 | 1.31 | 224 | 2.6 | 2.6 | 0.902 | 0.902 | 93.3 | 63.6 | 85.3 | 30.9 |
| 6muv | C | 3.8 | 9257 | 6 | 1.31 | 240 | 3.6 | 3.6 | 0.907 | 0.907 | 93.3 | 60.3 | 80 | 13.5 |
| 6muv | D | 3.8 | 9257 | 6 | 1.31 | 227 | 4.7 | 4.7 | 0.866 | 0.866 | 88.1 | 56.5 | 70 | 18.2 |
| 6muv | E | 3.8 | 9257 | 6 | 1.31 | 227 | 3.9 | 3.9 | 0.849 | 0.849 | 86.8 | 34 | 84.6 | 25.5 |
| 6muv | F | 3.8 | 9257 | 6 | 1.31 | 236 | 2.6 | 2.6 | 0.903 | 0.903 | 90.7 | 65.4 | 80.9 | 8.4 |
| 6muv | K | 3.8 | 9257 | 6 | 1.31 | 195 | 3.6 | 3.6 | 0.888 | 0.888 | 90.8 | 67.8 | 84.1 | 6.7 |
| 6muv | O | 3.8 | 9257 | 6 | 1.31 | 246 | 3.3 | 3.3 | 0.888 | 0.888 | 91.5 | 53.3 | 88.6 | 22.5 |
| 6muv | P | 3.8 | 9257 | 6 | 1.31 | 224 | 16.9 | 15.7 | 0.448 | 0.457 | 87.5 | 20.4 | 85.3 | 28.8 |
| 6muv | Q | 3.8 | 9257 | 6 | 1.31 | 240 | 2.3 | 2.3 | 0.934 | 0.934 | 94.6 | 76.2 | 83.8 | 37.8 |
| 6muv | R | 3.8 | 9257 | 6 | 1.31 | 227 | 2.9 | 2.9 | 0.882 | 0.882 | 90.7 | 55.8 | 78.9 | 27.4 |
| 6muv | S | 3.8 | 9257 | 6 | 1.31 | 227 | 14.1 | 13.8 | 0.639 | 0.644 | 84.1 | 39.3 | 81.5 | 32.4 |
| 6muv | T | 3.8 | 9257 | 6 | 1.31 | 236 | 3.5 | 3.5 | 0.886 | 0.886 | 92.8 | 68 | 85.2 | 31.8 |
| 6muv | Y | 3.8 | 9257 | 6 | 1.31 | 195 | 7.2 | 3.1 | 0.867 | 0.896 | 90.8 | 61.6 | 85.6 | 6.6 |
| 6muw | A | 3.6 | 9258 | 5 | 1.31 | 251 | 5.0 | 3.8 | 0.889 | 0.919 | 88 | 75.1 | 81.3 | 38.2 |
| 6muw | B | 3.6 | 9258 | 5 | 1.31 | 229 | 2.1 | 2.1 | 0.923 | 0.923 | 93.9 | 71.2 | 84.3 | 37.8 |
| 6muw | C | 3.6 | 9258 | 5 | 1.31 | 241 | 1.8 | 1.8 | 0.953 | 0.953 | 96.7 | 86.3 | 89.6 | 32.9 |
| 6muw | D | 3.6 | 9258 | 5 | 1.31 | 233 | 3.0 | 3.0 | 0.932 | 0.932 | 95.3 | 70.7 | 78.5 | 45.4 |
| 6muw | E | 3.6 | 9258 | 5 | 1.31 | 243 | 2.7 | 2.7 | 0.918 | 0.918 | 92.6 | 76.4 | 79.4 | 28.5 |
| 6muw | F | 3.6 | 9258 | 5 | 1.31 | 236 | 3.0 | 3.0 | 0.902 | 0.902 | 91.1 | 56.3 | 80.9 | 51.3 |
| 6muw | K | 3.6 | 9258 | 5 | 1.31 | 195 | 2.0 | 2.0 | 0.932 | 0.932 | 96.9 | 81.5 | 86.7 | 26 |
| 6muw | O | 3.6 | 9258 | 5 | 1.31 | 251 | 3.7 | 3.7 | 0.912 | 0.912 | 92.8 | 62.2 | 79.7 | 4.5 |

| | | | | | | | | | | | | | | |
|---|---|---|---|---|---|---|---|---|---|---|---|---|---|---|
| 6muw | P | 3.6 | 9258 | 5 | 1.31 | 229 | 2.8 | 2.8 | 0.914 | 0.914 | 92.1 | 76.3 | 82.5 | 38.1 |
| 6muw | Q | 3.6 | 9258 | 5 | 1.31 | 241 | 1.9 | 1.9 | 0.932 | 0.932 | 94.6 | 77.2 | 86.3 | 27.9 |
| 6muw | R | 3.6 | 9258 | 5 | 1.31 | 233 | 2.3 | 2.3 | 0.912 | 0.912 | 93.1 | 77.4 | 73 | 32.9 |
| 6muw | S | 3.6 | 9258 | 5 | 1.31 | 243 | 3.9 | 3.9 | 0.894 | 0.894 | 90.1 | 58.4 | 71.2 | 24.9 |
| 6muw | T | 3.6 | 9258 | 5 | 1.31 | 236 | 3.4 | 3.4 | 0.905 | 0.909 | 92.4 | 52.3 | 79.2 | 28.3 |
| 6muw | Y | 3.6 | 9258 | 5 | 1.31 | 195 | 2.1 | 2.1 | 0.938 | 0.938 | 96.9 | 87.8 | 84.1 | 49.4 |
| 6mux | A | 3.9 | 9259 | 6 | 1.31 | 244 | 2.7 | 2.7 | 0.923 | 0.923 | 93 | 68.7 | 88.9 | 50.2 |
| 6mux | B | 3.9 | 9259 | 6 | 1.31 | 224 | 3.7 | 3.7 | 0.897 | 0.897 | 91.5 | 73.7 | 81.2 | 7.7 |
| 6mux | C | 3.9 | 9259 | 6 | 1.31 | 240 | 5.6 | 4.5 | 0.891 | 0.891 | 92.1 | 68.8 | 82.5 | 46.5 |
| 6mux | D | 3.9 | 9259 | 6 | 1.31 | 227 | 3.0 | 3.0 | 0.910 | 0.910 | 94.3 | 59.8 | 85.9 | 37.4 |
| 6mux | E | 3.9 | 9259 | 6 | 1.31 | 227 | 5.0 | 5.0 | 0.860 | 0.860 | 87.7 | 47.7 | 75.8 | 25 |
| 6mux | F | 3.9 | 9259 | 6 | 1.31 | 235 | 4.9 | 4.9 | 0.881 | 0.881 | 89.8 | 58.3 | 85.1 | 27 |
| 6mux | K | 3.9 | 9259 | 6 | 1.31 | 195 | 3.2 | 3.2 | 0.881 | 0.884 | 89.7 | 70.3 | 88.7 | 6.4 |
| 6mux | O | 3.9 | 9259 | 6 | 1.31 | 244 | 3.7 | 3.7 | 0.899 | 0.899 | 92.6 | 71.7 | 82.4 | 11.4 |
| 6mux | P | 3.9 | 9259 | 6 | 1.31 | 229 | 2.7 | 2.7 | 0.901 | 0.901 | 92.1 | 65.9 | 83.8 | 49.5 |
| 6mux | Q | 3.9 | 9259 | 6 | 1.31 | 245 | 2.9 | 2.9 | 0.925 | 0.925 | 94.7 | 73.7 | 84.5 | 28.5 |
| 6mux | R | 3.9 | 9259 | 6 | 1.31 | 233 | 6.9 | 6.9 | 0.868 | 0.881 | 87.1 | 74.9 | 80.3 | 18.7 |
| 6mux | S | 3.9 | 9259 | 6 | 1.31 | 228 | 5.1 | 5.1 | 0.852 | 0.852 | 87.7 | 45.5 | 86.4 | 11.2 |
| 6mux | T | 3.9 | 9259 | 6 | 1.31 | 234 | 2.6 | 2.6 | 0.906 | 0.906 | 93.6 | 70.8 | 81.6 | 46.6 |
| 6mux | Y | 3.9 | 9259 | 6 | 1.31 | 195 | 2.8 | 2.8 | 0.908 | 0.908 | 94.4 | 73.9 | 88.2 | 17.4 |
| 6myy | C | 3.8 | 9294 | 0.08 | 1.36 | 100 | 13.9 | 11.7 | 0.317 | 0.383 | 73 | 9.6 | 38 | 2.6 |
| 6myy | D | 3.8 | 9294 | 0.08 | 1.36 | 118 | 10.4 | 4.8 | 0.728 | 0.856 | 89 | 71.4 | 57.6 | 7.4 |
| 6myy | F | 3.8 | 9294 | 0.08 | 1.36 | 100 | 14.7 | 10.7 | 0.335 | 0.417 | 73 | 9.6 | 40 | 2.5 |
| 6myy | G | 3.8 | 9294 | 0.08 | 1.36 | 118 | 2.5 | 2.5 | 0.849 | 0.849 | 89.8 | 61.3 | 69.5 | 7.3 |
| 6myy | H | 3.8 | 9294 | 0.08 | 1.36 | 118 | 9.6 | 9.6 | 0.735 | 0.735 | 87.3 | 68 | 58.5 | 15.9 |
| 6myy | L | 3.8 | 9294 | 0.08 | 1.36 | 100 | 13.2 | 13.2 | 0.310 | 0.365 | 74 | 5.4 | 48 | 16.7 |
| 6mzc | E | 4.5 | 9298 | 0.06 | 1.32 | 102 | 4.1 | 4.1 | 0.643 | 0.647 | 73.5 | 25.3 | 71.6 | 6.8 |
| 6mzc | H | 4.5 | 9298 | 0.06 | 1.32 | 257 | 5.8 | 5.8 | 0.824 | 0.840 | 85.6 | 33.6 | 39.7 | 4.9 |
| 6mzj | C | 4.8 | 9303 | 0.065 | 1.36 | 100 | 15.0 | 12.6 | 0.277 | 0.301 | 66 | 10.6 | 30 | 3.3 |
| 6mzj | D | 4.8 | 9303 | 0.065 | 1.36 | 118 | 15.7 | 14.8 | 0.294 | 0.340 | 66.9 | 10.1 | 43.2 | 5.9 |
| 6mzj | H | 4.8 | 9303 | 0.065 | 1.36 | 118 | 14.5 | 14.3 | 0.408 | 0.408 | 69.5 | 12.2 | 38.1 | 8.9 |
| 6mzj | L | 4.8 | 9303 | 0.065 | 1.36 | 100 | 13.5 | 13.2 | 0.246 | 0.367 | 64 | 4.7 | 25 | 0 |
| 6n1v | 1 | 4 | 9319 | 0.73 | 1.12 | 224 | 18.4 | 18.4 | 0.267 | 0.383 | 56.7 | 6.3 | 32.6 | 4.1 |
| 6n1v | 2 | 4 | 9319 | 0.73 | 1.12 | 215 | 19.0 | 17.1 | 0.273 | 0.347 | 59.1 | 7.1 | 15.3 | 9.1 |
| 6n1v | 3 | 4 | 9319 | 0.73 | 1.12 | 224 | 20.5 | 18.8 | 0.337 | 0.360 | 59.8 | 8.2 | 18.8 | 7.1 |
| 6n1v | 4 | 4 | 9319 | 0.73 | 1.12 | 215 | 20.4 | 18.4 | 0.252 | 0.306 | 58.6 | 7.9 | 14.9 | 9.4 |
| 6n1v | a | 4 | 9319 | 0.73 | 1.12 | 103 | 15.5 | 12.9 | 0.277 | 0.317 | 71.8 | 12.2 | 45.6 | 6.4 |

| | | | | | | | | | | | | | | |
|---|---|---|---|---|---|---|---|---|---|---|---|---|---|---|
| 6n1v | D | 4 | 9319 | 0.73 | 1.12 | 132 | 4.3 | 4.3 | 0.782 | 0.782 | 84.8 | 50 | 75 | 33.3 |
| 6n1v | E | 4 | 9319 | 0.73 | 1.12 | 132 | 15.0 | 5.5 | 0.283 | 0.781 | 77.3 | 6.9 | 72 | 34.7 |
| 6n1v | F | 4 | 9319 | 0.73 | 1.12 | 132 | 5.6 | 5.3 | 0.754 | 0.778 | 80.3 | 27.4 | 76.5 | 5.9 |
| 6n1v | h | 4 | 9319 | 0.73 | 1.12 | 102 | 13.7 | 13.2 | 0.288 | 0.373 | 75.5 | 7.8 | 25.5 | 3.8 |
| 6n1v | H | 4 | 9319 | 0.73 | 1.12 | 224 | 18.7 | 18.2 | 0.248 | 0.358 | 54.9 | 10.6 | 22.3 | 8 |
| 6n1v | i | 4 | 9319 | 0.73 | 1.12 | 102 | 14.2 | 10.4 | 0.515 | 0.648 | 72.5 | 4.1 | 47.1 | 14.6 |
| 6n1v | L | 4 | 9319 | 0.73 | 1.12 | 215 | 18.0 | 17.1 | 0.255 | 0.338 | 60 | 5.4 | 18.1 | 5.1 |
| 6n1v | M | 4 | 9319 | 0.73 | 1.12 | 132 | 18.9 | 16.1 | 0.241 | 0.294 | 65.2 | 11.6 | 37.1 | 0 |
| 6n1v | N | 4 | 9319 | 0.73 | 1.12 | 103 | 15.1 | 14.3 | 0.241 | 0.273 | 72.8 | 6.7 | 54.4 | 12.5 |
| 6n1v | R | 4 | 9319 | 0.73 | 1.12 | 102 | 13.7 | 13.3 | 0.458 | 0.458 | 72.5 | 9.5 | 48 | 6.1 |
| 6n1v | X | 4 | 9319 | 0.73 | 1.12 | 132 | 15.6 | 15.6 | 0.447 | 0.449 | 72 | 12.6 | 29.5 | 5.1 |
| 6n1v | Y | 4 | 9319 | 0.73 | 1.12 | 132 | 17.0 | 17.0 | 0.241 | 0.344 | 68.2 | 7.8 | 44.7 | 3.4 |
| 6n1v | Z | 4 | 9319 | 0.73 | 1.12 | 103 | 16.2 | 13.2 | 0.322 | 0.455 | 70.9 | 4.1 | 47.6 | 8.2 |
| 6n1w | 5 | 4.2 | 9320 | 0.688 | 1.07 | 132 | 16.6 | 14.6 | 0.264 | 0.297 | 62.1 | 6.1 | 25.8 | 5.9 |
| 6n1w | 6 | 4.2 | 9320 | 0.688 | 1.07 | 105 | 16.5 | 13.6 | 0.283 | 0.295 | 67.6 | 5.6 | 43.8 | 4.3 |
| 6n1w | 7 | 4.2 | 9320 | 0.688 | 1.07 | 102 | 15.6 | 13.7 | 0.282 | 0.386 | 74.5 | 10.5 | 30.4 | 9.7 |
| 6n1w | 8 | 4.2 | 9320 | 0.688 | 1.07 | 128 | 18.2 | 14.7 | 0.277 | 0.404 | 68.8 | 4.5 | 43.8 | 3.6 |
| 6n1w | m | 4.2 | 9320 | 0.688 | 1.07 | 132 | 17.6 | 14.2 | 0.231 | 0.339 | 60.6 | 6.2 | 23.5 | 6.5 |
| 6n1w | M | 4.2 | 9320 | 0.688 | 1.07 | 132 | 15.5 | 15.5 | 0.283 | 0.289 | 62.9 | 7.2 | 27.3 | 8.3 |
| 6n1w | n | 4.2 | 9320 | 0.688 | 1.07 | 105 | 14.4 | 14.2 | 0.291 | 0.291 | 62.9 | 6.1 | 55.2 | 5.2 |
| 6n1w | N | 4.2 | 9320 | 0.688 | 1.07 | 105 | 13.8 | 13.8 | 0.235 | 0.309 | 73.3 | 13 | 33.3 | 5.7 |
| 6n1w | q | 4.2 | 9320 | 0.688 | 1.07 | 128 | 17.1 | 15.5 | 0.256 | 0.324 | 68.8 | 8 | 35.2 | 8.9 |
| 6n1w | Q | 4.2 | 9320 | 0.688 | 1.07 | 128 | 13.2 | 13.2 | 0.309 | 0.568 | 76.6 | 9.2 | 52.3 | 9 |
| 6n1w | r | 4.2 | 9320 | 0.688 | 1.07 | 102 | 13.2 | 13.0 | 0.302 | 0.378 | 73.5 | 12 | 33.3 | 0 |
| 6n1w | R | 4.2 | 9320 | 0.688 | 1.07 | 102 | 14.0 | 12.8 | 0.407 | 0.415 | 66.7 | 4.4 | 49 | 2 |
| 6n38 | A | 3.7 | 9341 | 0.03 | 1.37 | 299 | 1.9 | 1.9 | 0.954 | 0.954 | 97.3 | 82.8 | 88.3 | 32.6 |
| 6n38 | C | 3.7 | 9341 | 0.03 | 1.37 | 299 | 2.9 | 2.9 | 0.922 | 0.922 | 93.6 | 73.9 | 77.6 | 46.6 |
| 6n38 | D | 3.7 | 9341 | 0.03 | 1.37 | 299 | 2.4 | 2.4 | 0.934 | 0.934 | 93.3 | 83.5 | 89.3 | 35.6 |
| 6n3q | E | 3.7 | 0336 | 0.42 | 1.16 | 135 | 2.6 | 2.6 | 0.863 | 0.863 | 91.1 | 74.8 | 86.7 | 24.8 |
| 6n4b | S | 3 | 0339 | 0.085 | 0.86 | 232 | 15.1 | 11.9 | 0.606 | 0.667 | 84.5 | 38.8 | 87.9 | 72.5 |
| 6n51 | C | 4 | 0345 | 0.04 | 1.06 | 123 | 14.7 | 13.1 | 0.273 | 0.318 | 65 | 10 | 35.8 | 6.8 |
| 6n51 | D | 4 | 0345 | 0.04 | 1.06 | 123 | 16.0 | 15.4 | 0.293 | 0.295 | 69.1 | 3.5 | 28.5 | 2.9 |
| 6n7h | C | 3.6 | 0356 | 0.045 | 1.114 | 174 | 5.2 | 5.2 | 0.824 | 0.830 | 86.2 | 59.3 | 66.7 | 5.2 |
| 6n7i | A | 3.2 | 0357 | 0.025 | 1.07 | 271 | 23.0 | 16.2 | 0.341 | 0.422 | 75.6 | 8.8 | 63.1 | 7 |
| 6n7i | B | 3.2 | 0357 | 0.025 | 1.07 | 274 | 3.3 | 3.3 | 0.902 | 0.902 | 91.6 | 53.8 | 72.3 | 31.8 |
| 6n7i | C | 3.2 | 0357 | 0.025 | 1.07 | 276 | 20.3 | 16.1 | 0.616 | 0.685 | 87.7 | 36.8 | 80.8 | 24.7 |
| 6n7i | D | 3.2 | 0357 | 0.025 | 1.07 | 271 | 17.7 | 11.6 | 0.662 | 0.783 | 73.1 | 9.6 | 78.6 | 8 |

| | | | | | | | | | | | | | | |
|---|---|---|---|---|---|---|---|---|---|---|---|---|---|---|
| 6n7i | E | 3.2 | 0357 | 0.025 | 1.07 | 241 | 18.7 | 16.8 | 0.329 | 0.581 | 76.3 | 6.5 | 69.7 | 25 |
| 6n7n | B | 3.5 | 0359 | 0.026 | 1.07 | 274 | 18.6 | 17.6 | 0.286 | 0.304 | 60.2 | 7.3 | 25.9 | 23.9 |
| 6n7n | C | 3.5 | 0359 | 0.026 | 1.07 | 276 | 19.4 | 17.3 | 0.324 | 0.392 | 72.1 | 4.5 | 45.3 | 21.6 |
| 6n7p | C | 3.6 | 0360 | 0.065 | 1.36 | 132 | 15.6 | 15.3 | 0.350 | 0.580 | 77.3 | 19.6 | 65.2 | 9.3 |
| 6n7p | G | 3.6 | 0360 | 0.065 | 1.36 | 239 | 4.3 | 4.3 | 0.864 | 0.864 | 92.9 | 74.8 | 79.9 | 30.9 |
| 6n7p | I | 3.6 | 0360 | 0.065 | 1.36 | 192 | 8.4 | 8.0 | 0.644 | 0.644 | 77.6 | 33.6 | 66.7 | 19.5 |
| 6n7p | L | 3.6 | 0360 | 0.065 | 1.36 | 119 | 1.9 | 1.9 | 0.893 | 0.893 | 95 | 73.5 | 76.5 | 25.3 |
| 6n7p | M | 3.6 | 0360 | 0.065 | 1.36 | 107 | 1.9 | 1.9 | 0.910 | 0.910 | 95.3 | 90.2 | 82.2 | 47.7 |
| 6n7p | N | 3.6 | 0360 | 0.065 | 1.36 | 93 | 1.0 | 1.0 | 0.944 | 0.944 | 100 | 97.8 | 48.4 | 6.7 |
| 6n7p | O | 3.6 | 0360 | 0.065 | 1.36 | 74 | 4.3 | 4.3 | 0.794 | 0.794 | 86.5 | 62.5 | 89.2 | 47 |
| 6n7p | P | 3.6 | 0360 | 0.065 | 1.36 | 74 | 1.5 | 1.5 | 0.880 | 0.880 | 97.3 | 94.4 | 85.1 | 22.2 |
| 6n7p | Q | 3.6 | 0360 | 0.065 | 1.36 | 71 | 2.8 | 2.8 | 0.832 | 0.832 | 90.1 | 67.2 | 56.3 | 10 |
| 6n7r | C | 3.2 | 0361 | 0.05 | 1.36 | 133 | 4.4 | 4.4 | 0.778 | 0.778 | 80.5 | 56.1 | 70.7 | 26.6 |
| 6n7r | F | 3.2 | 0361 | 0.05 | 1.36 | 185 | 5.9 | 5.9 | 0.848 | 0.855 | 93.5 | 65.9 | 85.4 | 8.9 |
| 6n7r | G | 3.2 | 0361 | 0.05 | 1.36 | 232 | 4.4 | 4.4 | 0.884 | 0.885 | 93.1 | 87 | 86.2 | 44 |
| 6n7r | K | 3.2 | 0361 | 0.05 | 1.36 | 124 | 2.0 | 2.0 | 0.913 | 0.923 | 94.4 | 89.7 | 88.7 | 11.8 |
| 6n7r | L | 3.2 | 0361 | 0.05 | 1.36 | 119 | 2.5 | 2.5 | 0.899 | 0.899 | 95 | 74.3 | 83.2 | 12.1 |
| 6n7r | M | 3.2 | 0361 | 0.05 | 1.36 | 107 | 2.3 | 2.3 | 0.901 | 0.901 | 94.4 | 66.3 | 85 | 11 |
| 6n7r | N | 3.2 | 0361 | 0.05 | 1.36 | 91 | 3.3 | 3.3 | 0.871 | 0.871 | 90.1 | 62.2 | 51.6 | 10.6 |
| 6n7r | O | 3.2 | 0361 | 0.05 | 1.36 | 75 | 3.6 | 3.6 | 0.824 | 0.824 | 90.7 | 57.4 | 86.7 | 38.5 |
| 6n7r | P | 3.2 | 0361 | 0.05 | 1.36 | 74 | 2.2 | 2.2 | 0.884 | 0.884 | 94.6 | 62.9 | 90.5 | 4.5 |
| 6n7r | Q | 3.2 | 0361 | 0.05 | 1.36 | 71 | 2.2 | 2.2 | 0.904 | 0.904 | 95.8 | 58.8 | 80.3 | 7 |
| 6n7s | A | 4.6 | 0362 | 0.026 | 1.07 | 271 | 22.1 | 19.0 | 0.225 | 0.307 | 51.7 | 8.6 | 19.9 | 5.6 |
| 6n7s | C | 4.6 | 0362 | 0.026 | 1.07 | 275 | 22.0 | 21.0 | 0.231 | 0.302 | 54.9 | 9.3 | 32.4 | 5.6 |
| 6n7t | A | 3.9 | 0363 | 0.026 | 1.07 | 271 | 21.3 | 19.5 | 0.260 | 0.487 | 67.5 | 9.8 | 43.5 | 5.9 |
| 6n7t | B | 3.9 | 0363 | 0.026 | 1.07 | 274 | 20.4 | 16.6 | 0.235 | 0.286 | 67.9 | 5.9 | 49.3 | 7.4 |
| 6n7t | C | 3.9 | 0363 | 0.026 | 1.07 | 275 | 21.2 | 19.7 | 0.323 | 0.344 | 72.4 | 14.1 | 60.7 | 7.2 |
| 6n7t | D | 3.9 | 0363 | 0.026 | 1.07 | 271 | 21.3 | 16.2 | 0.291 | 0.503 | 73.1 | 6.1 | 77.1 | 8.1 |
| 6n7t | E | 3.9 | 0363 | 0.026 | 1.07 | 258 | 23.0 | 19.8 | 0.249 | 0.387 | 63.6 | 3.7 | 41.1 | 8.5 |
| 6n7v | A | 3.8 | 0364 | 0.05 | 1.72 | 271 | 20.9 | 18.8 | 0.420 | 0.420 | 64.9 | 6.2 | 41 | 7.2 |
| 6n7v | B | 3.8 | 0364 | 0.05 | 1.72 | 274 | 21.9 | 17.4 | 0.318 | 0.334 | 70.4 | 5.2 | 60.6 | 15.7 |
| 6n7v | C | 3.8 | 0364 | 0.05 | 1.72 | 276 | 23.3 | 14.7 | 0.297 | 0.439 | 65.9 | 8.8 | 61.6 | 16.5 |
| 6n7v | D | 3.8 | 0364 | 0.05 | 1.72 | 271 | 20.2 | 20.1 | 0.414 | 0.415 | 70.1 | 5.3 | 73.1 | 14.1 |
| 6n7v | E | 3.8 | 0364 | 0.05 | 1.72 | 259 | 22.3 | 19.6 | 0.451 | 0.551 | 68.7 | 11.2 | 69.1 | 6.7 |
| 6n7v | F | 3.8 | 0364 | 0.05 | 1.72 | 255 | 22.4 | 18.3 | 0.218 | 0.305 | 58.4 | 6 | 58.4 | 6.7 |
| 6n7w | I | 4.5 | 0365 | 0.07 | 1.72 | 106 | 13.1 | 12.4 | 0.283 | 0.329 | 67.9 | 5.6 | 53.8 | 7 |
| 6n9u | E | 3.7 | 0379 | 0.04 | 0.86 | 222 | 17.8 | 16.1 | 0.343 | 0.688 | 69.4 | 16.2 | 72.1 | 6.9 |

| | | | | | | | | | | | | | | |
|---|---|---|---|---|---|---|---|---|---|---|---|---|---|---|
| 6n9u | F | 3.7 | 0379 | 0.04 | 0.86 | 196 | 12.7 | 10.8 | 0.510 | 0.510 | 69.4 | 19.9 | 68.4 | 7.5 |
| 6n9v | A | 4 | 0380 | 0.5 | 0.86 | 253 | 22.7 | 12.7 | 0.367 | 0.760 | 76.7 | 8.2 | 79.8 | 17.3 |
| 6n9v | B | 4 | 0380 | 0.5 | 0.86 | 275 | 21.8 | 18.4 | 0.311 | 0.346 | 78.2 | 9.3 | 82.2 | 5.8 |
| 6n9v | C | 4 | 0380 | 0.5 | 0.86 | 272 | 15.6 | 11.0 | 0.576 | 0.740 | 78.3 | 25.4 | 82 | 8.5 |
| 6n9v | D | 4 | 0380 | 0.5 | 0.86 | 267 | 21.4 | 19.1 | 0.380 | 0.425 | 69.7 | 13.4 | 88 | 6.8 |
| 6n9w | C | 4 | 0381 | 0.4 | 0.86 | 276 | 21.4 | 16.3 | 0.469 | 0.469 | 68.5 | 18.5 | 68.5 | 5.3 |
| 6n9w | D | 4 | 0381 | 0.4 | 0.86 | 261 | 20.8 | 16.4 | 0.288 | 0.297 | 67.4 | 9.1 | 54 | 3.5 |
| 6n9w | E | 4 | 0381 | 0.4 | 0.86 | 241 | 19.1 | 16.8 | 0.272 | 0.307 | 56.8 | 7.3 | 33.6 | 4.9 |
| 6n9x | D | 4.1 | 0382 | 0.05 | 0.86 | 271 | 17.2 | 16.3 | 0.294 | 0.397 | 63.1 | 7.6 | 39.5 | 8.4 |
| 6nb3 | D | 3.5 | 0401 | 0.7 | 1.37 | 123 | 16.4 | 15.6 | 0.281 | 0.305 | 74 | 18.7 | 32.5 | 12.5 |
| 6nb3 | E | 3.5 | 0401 | 0.7 | 1.37 | 107 | 14.1 | 13.4 | 0.262 | 0.372 | 66.4 | 14.1 | 50.5 | 9.3 |
| 6nb3 | H | 3.5 | 0401 | 0.7 | 1.37 | 124 | 17.4 | 15.5 | 0.312 | 0.378 | 64.5 | 13.8 | 33.9 | 7.1 |
| 6nb3 | L | 3.5 | 0401 | 0.7 | 1.37 | 107 | 15.7 | 14.8 | 0.314 | 0.361 | 65.4 | 11.4 | 40.2 | 9.3 |
| 6nbc | A | 2.8 | 0407 | 0.04 | 0.558 | 140 | 1.7 | 1.7 | 0.931 | 0.931 | 96.4 | 87.4 | 91.4 | 96.1 |
| 6nbc | B | 2.8 | 0407 | 0.04 | 0.558 | 143 | 2.5 | 2.5 | 0.923 | 0.923 | 94.4 | 68.1 | 93 | 87.2 |
| 6nbc | C | 2.8 | 0407 | 0.04 | 0.558 | 140 | 1.8 | 1.8 | 0.934 | 0.934 | 97.1 | 89 | 89.3 | 98.4 |
| 6nbc | D | 2.8 | 0407 | 0.04 | 0.558 | 143 | 2.3 | 2.3 | 0.931 | 0.931 | 96.5 | 70.3 | 93 | 56.4 |
| 6nbd | A | 3.2 | 0408 | 0.045 | 0.5585 | 140 | 2.2 | 2.2 | 0.939 | 0.939 | 97.1 | 85.3 | 86.4 | 24 |
| 6nbd | B | 3.2 | 0408 | 0.045 | 0.5585 | 143 | 2.0 | 2.0 | 0.932 | 0.932 | 96.5 | 73.9 | 87.4 | 71.2 |
| 6nbd | D | 3.2 | 0408 | 0.045 | 0.5585 | 143 | 3.1 | 3.1 | 0.908 | 0.908 | 94.4 | 64.4 | 86.7 | 77.4 |
| 6nbi | N | 4 | 0412 | 0.014 | 1.014 | 126 | 16.8 | 14.7 | 0.590 | 0.611 | 77.8 | 9.2 | 68.3 | 5.8 |
| 6nbq | J | 3.1 | 0415 | 0.04 | 1.068 | 161 | 2.5 | 2.5 | 0.917 | 0.917 | 93.8 | 67.5 | 75.2 | 62.8 |
| 6nbq | M | 3.1 | 0415 | 0.04 | 1.068 | 111 | 2.1 | 2.1 | 0.923 | 0.923 | 96.4 | 88.8 | 92.8 | 53.4 |
| 6nbq | N | 3.1 | 0415 | 0.04 | 1.068 | 148 | 2.2 | 2.2 | 0.897 | 0.897 | 92.6 | 78.8 | 87.2 | 11.6 |
| 6nbq | O | 3.1 | 0415 | 0.04 | 1.068 | 69 | 2.3 | 2.3 | 0.858 | 0.858 | 92.8 | 56.2 | 55.1 | 5.3 |
| 6nbx | J | 3.5 | 0425 | 5.5 | 1.068 | 161 | 6.7 | 6.7 | 0.789 | 0.794 | 78.9 | 59.8 | 81.4 | 19.1 |
| 6nbx | M | 3.5 | 0425 | 5.5 | 1.068 | 111 | 2.2 | 2.2 | 0.896 | 0.896 | 94.6 | 65.7 | 84.7 | 60.6 |
| 6nbx | N | 3.5 | 0425 | 5.5 | 1.068 | 148 | 3.8 | 3.8 | 0.881 | 0.881 | 91.2 | 59.3 | 75.7 | 33.9 |
| 6nbx | O | 3.5 | 0425 | 5.5 | 1.068 | 69 | 3.4 | 3.4 | 0.836 | 0.836 | 91.3 | 44.4 | 82.6 | 15.8 |
| 6nc3 | H | 4.5 | 0434 | 0.9 | 1.03 | 119 | 15.1 | 15.1 | 0.272 | 0.272 | 59.7 | 8.5 | 28.6 | 0 |
| 6nc3 | L | 4.5 | 0434 | 0.9 | 1.03 | 106 | 14.1 | 13.6 | 0.309 | 0.314 | 64.2 | 5.9 | 29.2 | 9.7 |
| 6nc3 | O | 4.5 | 0434 | 0.9 | 1.03 | 119 | 13.8 | 13.1 | 0.282 | 0.286 | 54.6 | 4.6 | 32.8 | 7.7 |
| 6nc3 | P | 4.5 | 0434 | 0.9 | 1.03 | 119 | 17.0 | 14.6 | 0.287 | 0.287 | 60.5 | 2.8 | 26.1 | 9.7 |
| 6nc3 | Q | 4.5 | 0434 | 0.9 | 1.03 | 119 | 18.2 | 15.8 | 0.272 | 0.277 | 57.1 | 5.9 | 30.3 | 8.3 |
| 6nc3 | R | 4.5 | 0434 | 0.9 | 1.03 | 119 | 18.2 | 15.9 | 0.235 | 0.301 | 52.9 | 7.9 | 27.7 | 9.1 |
| 6nc3 | S | 4.5 | 0434 | 0.9 | 1.03 | 119 | 14.6 | 14.6 | 0.262 | 0.274 | 48.7 | 5.2 | 31.9 | 7.9 |
| 6nc3 | T | 4.5 | 0434 | 0.9 | 1.03 | 106 | 16.7 | 13.9 | 0.286 | 0.366 | 62.3 | 3 | 32.1 | 2.9 |

| | | | | | | | | | | | | | | |
|---|---|---|---|---|---|---|---|---|---|---|---|---|---|---|
| 6nc3 | U | 4.5 | 0434 | 0.9 | 1.03 | 106 | 14.7 | 13.4 | 0.288 | 0.314 | 57.5 | 9.8 | 29.2 | 6.5 |
| 6nc3 | V | 4.5 | 0434 | 0.9 | 1.03 | 106 | 14.8 | 13.6 | 0.297 | 0.297 | 56.6 | 8.3 | 34.9 | 2.7 |
| 6nc3 | W | 4.5 | 0434 | 0.9 | 1.03 | 106 | 15.6 | 14.2 | 0.282 | 0.297 | 63.2 | 7.5 | 25.5 | 0 |
| 6nc3 | X | 4.5 | 0434 | 0.9 | 1.03 | 106 | 15.8 | 13.8 | 0.216 | 0.299 | 53.8 | 5.3 | 29.2 | 19.4 |
| 6nd4 | e | 4.3 | 0441 | 0.0203 | 1.6 | 121 | 13.8 | 12.0 | 0.333 | 0.356 | 69.4 | 11.9 | 74.4 | 2.2 |
| 6nd4 | Z | 4.3 | 0441 | 0.0203 | 1.6 | 169 | 5.8 | 5.8 | 0.767 | 0.770 | 78.7 | 24.8 | 54.4 | 5.4 |
| 6nf2 | B | 3.7 | 9359 | 0.4 | 1.07325 | 132 | 12.7 | 6.4 | 0.676 | 0.751 | 74.2 | 46.9 | 69.7 | 4.3 |
| 6nf2 | C | 3.7 | 9359 | 0.4 | 1.07325 | 128 | 2.6 | 2.6 | 0.894 | 0.894 | 93.8 | 72.5 | 55.5 | 5.6 |
| 6nf2 | D | 3.7 | 9359 | 0.4 | 1.07325 | 102 | 15.3 | 12.8 | 0.565 | 0.578 | 82.4 | 6 | 73.5 | 4 |
| 6nf2 | E | 3.7 | 9359 | 0.4 | 1.07325 | 132 | 14.9 | 12.7 | 0.384 | 0.473 | 71.2 | 23.4 | 24.2 | 12.5 |
| 6nf2 | F | 3.7 | 9359 | 0.4 | 1.07325 | 105 | 16.4 | 13.5 | 0.647 | 0.676 | 76.2 | 2.5 | 58.1 | 1.6 |
| 6nf2 | I | 3.7 | 9359 | 0.4 | 1.07325 | 132 | 7.5 | 7.5 | 0.789 | 0.789 | 83.3 | 22.7 | 75 | 16.2 |
| 6nf2 | J | 3.7 | 9359 | 0.4 | 1.07325 | 132 | 15.6 | 15.3 | 0.455 | 0.468 | 71.2 | 9.6 | 28 | 21.6 |
| 6nf2 | K | 3.7 | 9359 | 0.4 | 1.07325 | 105 | 13.7 | 13.2 | 0.544 | 0.584 | 73.3 | 3.9 | 54.3 | 5.3 |
| 6nf2 | M | 3.7 | 9359 | 0.4 | 1.07325 | 102 | 8.7 | 6.5 | 0.666 | 0.766 | 83.3 | 45.9 | 70.6 | 2.8 |
| 6nf2 | N | 3.7 | 9359 | 0.4 | 1.07325 | 128 | 1.6 | 1.6 | 0.914 | 0.914 | 96.9 | 79 | 77.3 | 5.1 |
| 6nf2 | R | 3.7 | 9359 | 0.4 | 1.07325 | 132 | 11.9 | 4.6 | 0.684 | 0.806 | 70.5 | 31.2 | 63.6 | 57.1 |
| 6nf2 | S | 3.7 | 9359 | 0.4 | 1.07325 | 132 | 18.5 | 15.7 | 0.316 | 0.330 | 65.9 | 4.6 | 35.6 | 6.4 |
| 6nf2 | T | 3.7 | 9359 | 0.4 | 1.07325 | 105 | 15.6 | 10.8 | 0.475 | 0.618 | 74.3 | 10.3 | 40 | 4.8 |
| 6nf2 | U | 3.7 | 9359 | 0.4 | 1.07325 | 102 | 13.4 | 13.1 | 0.327 | 0.606 | 80.4 | 17.1 | 78.4 | 7.5 |
| 6nf2 | V | 3.7 | 9359 | 0.4 | 1.07325 | 128 | 7.7 | 5.5 | 0.804 | 0.824 | 85.9 | 60.9 | 64.1 | 8.5 |
| 6nf8 | a | 3.5 | 9362 | 0.15 | 1.07 | 289 | 15.9 | 10.9 | 0.578 | 0.686 | 71.3 | 20.4 | 65.7 | 6.8 |
| 6nf8 | B | 3.5 | 9362 | 0.15 | 1.07 | 217 | 6.4 | 6.3 | 0.828 | 0.836 | 84.8 | 40.2 | 74.7 | 18.5 |
| 6nht | C | 2.9 | 9373 | 0.9 | 1.06 | 168 | 1.9 | 1.9 | 0.927 | 0.927 | 94.6 | 76.1 | 77.4 | 26.9 |
| 6nht | H | 2.9 | 9373 | 0.9 | 1.06 | 118 | 2.1 | 2.1 | 0.916 | 0.916 | 94.9 | 77.7 | 89.8 | 64.2 |
| 6nht | J | 2.9 | 9373 | 0.9 | 1.06 | 168 | 1.4 | 1.4 | 0.956 | 0.956 | 97.6 | 92.7 | 78.6 | 97 |
| 6nht | L | 2.9 | 9373 | 0.9 | 1.06 | 168 | 4.4 | 4.4 | 0.857 | 0.857 | 92.9 | 78.2 | 80.4 | 92.6 |
| 6nht | P | 2.9 | 9373 | 0.9 | 1.06 | 118 | 1.6 | 1.6 | 0.926 | 0.926 | 97.5 | 91.3 | 93.2 | 82.7 |
| 6nht | Q | 2.9 | 9373 | 0.9 | 1.06 | 168 | 5.3 | 5.3 | 0.847 | 0.847 | 86.3 | 66.9 | 80.4 | 94.8 |
| 6nht | R | 2.9 | 9373 | 0.9 | 1.06 | 168 | 1.9 | 1.9 | 0.930 | 0.930 | 95.8 | 86.3 | 81.5 | 53.3 |
| 6nht | T | 2.9 | 9373 | 0.9 | 1.06 | 168 | 3.1 | 3.1 | 0.911 | 0.911 | 94 | 74.7 | 81.5 | 94.9 |
| 6ni2 | A | 4 | 9375 | 0.22 | 1.04 | 113 | 15.7 | 13.8 | 0.322 | 0.339 | 63.7 | 4.2 | 31 | 11.4 |
| 6ni2 | H | 4 | 9375 | 0.22 | 1.04 | 117 | 13.9 | 12.1 | 0.386 | 0.686 | 76.9 | 10 | 53.8 | 4.8 |
| 6ni2 | L | 4 | 9375 | 0.22 | 1.04 | 107 | 11.7 | 11.7 | 0.296 | 0.345 | 74.8 | 7.5 | 33.6 | 8.3 |
| 6ni3 | N | 3.8 | 9376 | 0.2 | 1.04 | 128 | 16.4 | 14.0 | 0.533 | 0.533 | 76.6 | 20.4 | 46.1 | 6.8 |
| 6ni3 | R | 3.8 | 9376 | 0.2 | 1.04 | 284 | 18.7 | 16.6 | 0.554 | 0.590 | 77.1 | 8.2 | 74.6 | 15.1 |
| 6nil | A | 3.9 | 9380 | 0.025 | 1.05 | 177 | 18.2 | 7.8 | 0.324 | 0.763 | 72.3 | 7.8 | 55.4 | 7.1 |

| | | | | | | | | | | | | | | |
|---|---|---|---|---|---|---|---|---|---|---|---|---|---|---|
| 6nil | B | 3.9 | 9380 | 0.025 | 1.05 | 140 | 17.3 | 15.1 | 0.271 | 0.313 | 70.7 | 15.2 | 45 | 50.8 |
| 6nil | C | 3.9 | 9380 | 0.025 | 1.05 | 133 | 16.9 | 11.1 | 0.285 | 0.504 | 64.7 | 10.5 | 62.4 | 37.3 |
| 6nil | D | 3.9 | 9380 | 0.025 | 1.05 | 177 | 17.5 | 14.5 | 0.343 | 0.494 | 76.3 | 14.1 | 61 | 10.2 |
| 6nil | E | 3.9 | 9380 | 0.025 | 1.05 | 140 | 15.2 | 15.2 | 0.281 | 0.394 | 75 | 11.4 | 40 | 1.8 |
| 6nil | F | 3.9 | 9380 | 0.025 | 1.05 | 133 | 15.2 | 13.7 | 0.368 | 0.511 | 66.2 | 5.7 | 66.2 | 28.4 |
| 6nil | G | 3.9 | 9380 | 0.025 | 1.05 | 177 | 12.2 | 8.8 | 0.649 | 0.698 | 79.7 | 36.9 | 67.8 | 5.8 |
| 6nil | H | 3.9 | 9380 | 0.025 | 1.05 | 140 | 15.6 | 15.6 | 0.283 | 0.402 | 73.6 | 10.7 | 25 | 5.7 |
| 6nil | I | 3.9 | 9380 | 0.025 | 1.05 | 133 | 10.8 | 7.3 | 0.687 | 0.689 | 71.4 | 25.3 | 52.6 | 4.3 |
| 6nil | J | 3.9 | 9380 | 0.025 | 1.05 | 177 | 19.0 | 14.0 | 0.295 | 0.342 | 80.2 | 7 | 60.5 | 4.7 |
| 6nil | K | 3.9 | 9380 | 0.025 | 1.05 | 140 | 16.3 | 15.8 | 0.280 | 0.394 | 71.4 | 10 | 42.9 | 5 |
| 6nil | L | 3.9 | 9380 | 0.025 | 1.05 | 133 | 17.6 | 10.4 | 0.423 | 0.658 | 70.7 | 6.4 | 59.4 | 2.5 |
| 6niy | N | 3.3 | 9382 | 0.015 | 1.06 | 128 | 3.3 | 3.3 | 0.856 | 0.856 | 93 | 57.1 | 73.4 | 9.6 |
| 6niy | R | 3.3 | 9382 | 0.015 | 1.06 | 261 | 4.7 | 4.7 | 0.823 | 0.834 | 82.4 | 42.3 | 76.6 | 28.5 |
| 6njp | G | 3.3 | 9391 | 0.05 | 1.02 | 64 | 5.1 | 4.7 | 0.670 | 0.670 | 75 | 10.4 | 65.6 | 2.4 |
| 6nm9 | A | 3.4 | 9398 | 0.0177 | 1.066 | 115 | 1.9 | 1.9 | 0.901 | 0.901 | 96.5 | 87.4 | 88.7 | 42.2 |
| 6nm9 | C | 3.4 | 9398 | 0.0177 | 1.066 | 115 | 1.9 | 1.9 | 0.888 | 0.888 | 93.9 | 82.4 | 86.1 | 34.3 |
| 6nma | A | 3.4 | 0445 | 0.0226 | 1.066 | 115 | 2.3 | 2.3 | 0.903 | 0.903 | 94.8 | 68.8 | 91.3 | 4.8 |
| 6nma | C | 3.4 | 0445 | 0.0226 | 1.066 | 114 | 13.1 | 11.4 | 0.408 | 0.429 | 68.4 | 33.3 | 50.9 | 43.1 |
| 6nmc | B | 4.2 | 0446 | 0.0169 | 1.066 | 156 | 6.7 | 3.8 | 0.803 | 0.803 | 82.1 | 46.9 | 76.3 | 5 |
| 6nmi | F | 3.7 | 0452 | 0.0172 | 1.15 | 263 | 7.6 | 5.1 | 0.828 | 0.828 | 84.4 | 46.8 | 77.6 | 14.2 |
| 6nmi | H | 3.7 | 0452 | 0.0172 | 1.15 | 210 | 9.5 | 8.3 | 0.665 | 0.726 | 73.8 | 23.9 | 27.6 | 6.9 |
| 6nn6 | L | 3.9 | 0458 | 0.03 | 1.08 | 74 | 10.8 | 8.6 | 0.347 | 0.347 | 67.6 | 8 | 48.6 | 5.6 |
| 6nog | L | 3.9 | 0468 | 0.0184 | 1.064 | 74 | 10.6 | 10.2 | 0.304 | 0.304 | 62.2 | 6.5 | 43.2 | 9.4 |
| 6npy | B | 3.8 | 0476 | 0.00926 | 0.84 | 167 | 16.4 | 13.7 | 0.286 | 0.381 | 66.5 | 4.5 | 32.9 | 1.8 |
| 6nqa | L | 3.5 | 0480 | 0.0166 | 1.064 | 75 | 9.2 | 9.1 | 0.367 | 0.406 | 77.3 | 6.9 | 57.3 | 4.7 |
| 6nqb | C | 3.8 | 0482 | 5.18 | 1.073 | 206 | 14.0 | 13.9 | 0.503 | 0.559 | 66 | 11 | 33.5 | 5.8 |
| 6nqb | D | 3.8 | 0482 | 5.18 | 1.073 | 205 | 15.0 | 7.0 | 0.610 | 0.767 | 76.1 | 14.7 | 31.7 | 6.2 |
| 6nsj | A | 2.7 | 0498 | 0.043 | 1.07 | 182 | 3.2 | 3.2 | 0.911 | 0.911 | 92.3 | 84.5 | 87.4 | 83 |
| 6nsj | B | 2.7 | 0498 | 0.043 | 1.07 | 182 | 4.6 | 4.6 | 0.867 | 0.867 | 89 | 71.6 | 79.7 | 84.8 |
| 6nsj | C | 2.7 | 0498 | 0.043 | 1.07 | 182 | 3.0 | 3.0 | 0.900 | 0.900 | 91.8 | 76 | 80.2 | 73.3 |
| 6nsj | D | 2.7 | 0498 | 0.043 | 1.07 | 182 | 4.3 | 4.3 | 0.879 | 0.879 | 90.1 | 80.5 | 85.2 | 81.9 |
| 6nsj | E | 2.7 | 0498 | 0.043 | 1.07 | 182 | 3.0 | 3.0 | 0.896 | 0.896 | 89.6 | 85.9 | 79.1 | 87.5 |
| 6nsj | F | 2.7 | 0498 | 0.043 | 1.07 | 182 | 2.8 | 2.8 | 0.907 | 0.907 | 92.9 | 82.2 | 79.1 | 94.4 |
| 6nsk | A | 2.7 | 0499 | 0.06 | 1.07 | 182 | 2.9 | 2.9 | 0.912 | 0.912 | 92.9 | 88.8 | 87.9 | 86.2 |
| 6nsk | B | 2.7 | 0499 | 0.06 | 1.07 | 182 | 4.0 | 4.0 | 0.906 | 0.906 | 91.8 | 76.6 | 86.8 | 65.8 |
| 6nsk | C | 2.7 | 0499 | 0.06 | 1.07 | 182 | 2.5 | 2.5 | 0.911 | 0.911 | 94 | 75.4 | 80.8 | 39.5 |
| 6nsk | D | 2.7 | 0499 | 0.06 | 1.07 | 182 | 3.1 | 3.1 | 0.908 | 0.908 | 91.8 | 74.9 | 87.9 | 60 |

| | | | | | | | | | | | | | | |
|---|---|---|---|---|---|---|---|---|---|---|---|---|---|---|
| 6nsk | E | 2.7 | 0499 | 0.06 | 1.07 | 182 | 2.6 | 2.6 | 0.916 | 0.916 | 93.4 | 85.3 | 84.1 | 64.1 |
| 6nsk | F | 2.7 | 0499 | 0.06 | 1.07 | 182 | 3.1 | 3.1 | 0.905 | 0.905 | 91.2 | 76.5 | 86.8 | 87.3 |
| 6nuw | C | 4.3 | 0523 | 0.03 | 1.23 | 194 | 5.0 | 5.0 | 0.753 | 0.775 | 84.5 | 57.3 | 75.8 | 5.4 |
| 6nyb | B | 4.1 | 0541 | 0.048 | 1.06 | 289 | 21.8 | 20.8 | 0.263 | 0.300 | 65.4 | 5.3 | 56.4 | 6.1 |
| 6nyb | C | 4.1 | 0541 | 0.048 | 1.06 | 229 | 3.2 | 3.2 | 0.893 | 0.893 | 89.5 | 54.1 | 91.7 | 7.1 |
| 6nyb | D | 4.1 | 0541 | 0.048 | 1.06 | 229 | 5.3 | 5.1 | 0.866 | 0.866 | 87.3 | 29 | 92.6 | 21.7 |
| 6nzd | F | 3.6 | 0554 | 0.16 | 1.149 | 227 | 6.7 | 6.7 | 0.742 | 0.742 | 81.1 | 34.2 | 84.1 | 3.7 |
| 6nzd | G | 3.6 | 0554 | 0.16 | 1.149 | 299 | 13.0 | 8.1 | 0.658 | 0.740 | 84.9 | 29.9 | 60.9 | 22.5 |
| 6nzo | S | 3.8 | 0559 | 0.012 | 1.04 | 233 | 18.1 | 17.3 | 0.283 | 0.288 | 59.2 | 2.9 | 12.4 | 3.4 |
| 6nzu | I | 3.2 | 0560 | 0.0425 | 1.086 | 118 | 2.5 | 2.5 | 0.914 | 0.914 | 94.9 | 73.2 | 89.8 | 31.1 |
| 6nzu | J | 3.2 | 0560 | 0.0425 | 1.086 | 118 | 1.4 | 1.4 | 0.939 | 0.939 | 97.5 | 91.3 | 85.6 | 8.9 |
| 6o1k | A | 3.1 | 0590 | 0.045 | 1.09 | 262 | 2.2 | 2.2 | 0.943 | 0.943 | 96.2 | 76.2 | 86.6 | 35.2 |
| 6o1k | B | 3.1 | 0590 | 0.045 | 1.09 | 262 | 1.7 | 1.7 | 0.953 | 0.953 | 97.3 | 83.1 | 89.7 | 30.2 |
| 6o1l | A | 3.4 | 0591 | 0.27 | 1.09 | 262 | 18.1 | 16.8 | 0.453 | 0.491 | 76 | 16.6 | 77.5 | 8.9 |
| 6o1l | B | 3.4 | 0591 | 0.27 | 1.09 | 262 | 16.6 | 14.4 | 0.345 | 0.475 | 73.3 | 5.7 | 75.2 | 32.5 |
| 6o1l | D | 3.4 | 0591 | 0.27 | 1.09 | 67 | 10.6 | 10.0 | 0.292 | 0.406 | 61.2 | 7.3 | 49.3 | 9.1 |
| 6o1l | F | 3.4 | 0591 | 0.27 | 1.09 | 65 | 11.1 | 11.1 | 0.248 | 0.379 | 61.5 | 12.5 | 58.5 | 7.9 |
| 6o1l | H | 3.4 | 0591 | 0.27 | 1.09 | 67 | 12.0 | 10.1 | 0.226 | 0.354 | 65.7 | 13.6 | 79.1 | 13.2 |
| 6o1l | J | 3.4 | 0591 | 0.27 | 1.09 | 65 | 9.6 | 7.2 | 0.618 | 0.618 | 78.5 | 29.4 | 80 | 1.9 |
| 6o1l | K | 3.4 | 0591 | 0.27 | 1.09 | 65 | 11.1 | 9.9 | 0.608 | 0.608 | 78.5 | 23.5 | 83.1 | 1.9 |
| 6o1l | L | 3.4 | 0591 | 0.27 | 1.09 | 66 | 11.4 | 7.6 | 0.553 | 0.553 | 72.7 | 25 | 53 | 11.4 |
| 6o1l | N | 3.4 | 0591 | 0.27 | 1.09 | 66 | 6.0 | 6.0 | 0.675 | 0.675 | 77.3 | 39.2 | 92.4 | 8.2 |
| 6o1l | R | 3.4 | 0591 | 0.27 | 1.09 | 262 | 18.2 | 15.9 | 0.350 | 0.359 | 58.4 | 9.8 | 6.1 | 18.8 |
| 6o1m | A | 3.2 | 0592 | 0.237 | 1.09 | 262 | 17.8 | 16.0 | 0.548 | 0.561 | 74.8 | 15.3 | 71.8 | 16.5 |
| 6o1m | B | 3.2 | 0592 | 0.237 | 1.09 | 262 | 19.2 | 16.3 | 0.376 | 0.427 | 66.8 | 7.4 | 76 | 25.1 |
| 6o1m | C | 3.2 | 0592 | 0.237 | 1.09 | 65 | 5.1 | 5.1 | 0.736 | 0.736 | 83.1 | 37 | 78.5 | 2 |
| 6o1m | D | 3.2 | 0592 | 0.237 | 1.09 | 67 | 3.6 | 3.6 | 0.777 | 0.777 | 86.6 | 63.8 | 80.6 | 5.6 |
| 6o1m | E | 3.2 | 0592 | 0.237 | 1.09 | 67 | 9.4 | 8.4 | 0.458 | 0.466 | 74.6 | 16 | 77.6 | 34.6 |
| 6o1m | F | 3.2 | 0592 | 0.237 | 1.09 | 65 | 9.3 | 9.3 | 0.536 | 0.543 | 73.8 | 33.3 | 76.9 | 6 |
| 6o1m | G | 3.2 | 0592 | 0.237 | 1.09 | 66 | 10.3 | 7.3 | 0.539 | 0.626 | 72.7 | 12.5 | 81.8 | 57.4 |
| 6o1m | H | 3.2 | 0592 | 0.237 | 1.09 | 67 | 6.9 | 5.1 | 0.603 | 0.748 | 73.1 | 2 | 80.6 | 48.1 |
| 6o1m | I | 3.2 | 0592 | 0.237 | 1.09 | 67 | 11.6 | 7.6 | 0.313 | 0.589 | 73.1 | 12.2 | 86.6 | 32.8 |
| 6o1m | J | 3.2 | 0592 | 0.237 | 1.09 | 65 | 8.4 | 7.3 | 0.591 | 0.606 | 72.3 | 21.3 | 75.4 | 6.1 |
| 6o1m | K | 3.2 | 0592 | 0.237 | 1.09 | 65 | 11.0 | 10.4 | 0.334 | 0.503 | 72.3 | 8.5 | 83.1 | 29.6 |
| 6o1m | L | 3.2 | 0592 | 0.237 | 1.09 | 66 | 7.5 | 5.2 | 0.679 | 0.696 | 77.3 | 31.4 | 84.8 | 12.5 |
| 6o1m | M | 3.2 | 0592 | 0.237 | 1.09 | 67 | 7.3 | 6.3 | 0.625 | 0.658 | 70.1 | 36.2 | 46.3 | 0 |
| 6o1m | N | 3.2 | 0592 | 0.237 | 1.09 | 66 | 11.1 | 9.9 | 0.529 | 0.529 | 72.7 | 14.6 | 89.4 | 27.1 |

| | | | | | | | | | | | | | | |
|---|---|---|---|---|---|---|---|---|---|---|---|---|---|---|
| 6o1m | R | 3.2 | 0592 | 0.237 | 1.09 | 262 | 19.0 | 16.7 | 0.272 | 0.336 | 53.1 | 5 | 32.1 | 3.6 |
| 6o1m | S | 3.2 | 0592 | 0.237 | 1.09 | 262 | 19.6 | 17.5 | 0.248 | 0.302 | 63 | 4.8 | 31.7 | 12 |
| 6o58 | A | 3.8 | 0625 | 0.017 | 0.84 | 273 | 10.6 | 6.9 | 0.694 | 0.747 | 83.2 | 41.4 | 65.9 | 9.4 |
| 6o58 | C | 3.8 | 0625 | 0.017 | 0.84 | 268 | 4.1 | 4.1 | 0.851 | 0.851 | 85.8 | 38.3 | 82.8 | 6.3 |
| 6o58 | E | 3.8 | 0625 | 0.017 | 0.84 | 270 | 8.8 | 5.3 | 0.856 | 0.856 | 85.9 | 36.6 | 31.5 | 11.8 |
| 6o58 | G | 3.8 | 0625 | 0.017 | 0.84 | 250 | 28.0 | 19.0 | 0.574 | 0.574 | 58.8 | 6.8 | 74 | 5.4 |
| 6o58 | I | 3.8 | 0625 | 0.017 | 0.84 | 273 | 8.4 | 8.3 | 0.739 | 0.739 | 79.5 | 29.5 | 32.2 | 4.5 |
| 6o58 | K | 3.8 | 0625 | 0.017 | 0.84 | 268 | 4.7 | 4.7 | 0.854 | 0.861 | 86.6 | 31 | 39.9 | 6.5 |
| 6o58 | M | 3.8 | 0625 | 0.017 | 0.84 | 270 | 5.9 | 4.3 | 0.813 | 0.858 | 82.6 | 33.2 | 77.4 | 26.8 |
| 6o58 | O | 3.8 | 0625 | 0.017 | 0.84 | 250 | 42.2 | 27.0 | 0.260 | 0.591 | 55.6 | 8.6 | 64.4 | 6.2 |
| 6o5b | A | 3.6 | 0626 | 0.01 | 0.84 | 273 | 3.9 | 3.9 | 0.893 | 0.893 | 92.3 | 66.7 | 77.7 | 29.7 |
| 6o5b | C | 3.6 | 0626 | 0.01 | 0.84 | 268 | 2.6 | 2.6 | 0.925 | 0.925 | 94 | 69 | 86.6 | 50.4 |
| 6o5b | E | 3.6 | 0626 | 0.01 | 0.84 | 270 | 4.3 | 4.3 | 0.905 | 0.905 | 91.9 | 49.2 | 88.9 | 16.2 |
| 6o5b | G | 3.6 | 0626 | 0.01 | 0.84 | 250 | 20.0 | 20.0 | 0.614 | 0.615 | 61.6 | 37.7 | 76.8 | 58.3 |
| 6o5b | I | 3.6 | 0626 | 0.01 | 0.84 | 91 | 13.9 | 11.5 | 0.288 | 0.351 | 74.7 | 19.1 | 74.7 | 5.9 |
| 6o5b | J | 3.6 | 0626 | 0.01 | 0.84 | 91 | 3.2 | 3.2 | 0.858 | 0.858 | 93.4 | 71.8 | 80.2 | 9.6 |
| 6o5b | K | 3.6 | 0626 | 0.01 | 0.84 | 91 | 5.0 | 4.5 | 0.799 | 0.810 | 87.9 | 32.5 | 78 | 4.2 |
| 6o6c | C | 3.1 | 0633 | 0.0509 | 1.16 | 270 | 2.8 | 2.8 | 0.933 | 0.933 | 94.1 | 80.7 | 84.8 | 15.7 |
| 6o6c | D | 3.1 | 0633 | 0.0509 | 1.16 | 215 | 7.1 | 7.1 | 0.840 | 0.840 | 85.6 | 65.8 | 77.7 | 18 |
| 6o6c | E | 3.1 | 0633 | 0.0509 | 1.16 | 81 | 1.7 | 1.7 | 0.870 | 0.870 | 93.8 | 84.2 | 84 | 35.3 |
| 6o6c | F | 3.1 | 0633 | 0.0509 | 1.16 | 146 | 7.9 | 7.9 | 0.755 | 0.755 | 78.8 | 51.3 | 63.7 | 5.4 |
| 6o6c | H | 3.1 | 0633 | 0.0509 | 1.16 | 65 | 1.5 | 1.5 | 0.848 | 0.848 | 93.8 | 91.8 | 83.1 | 0 |
| 6o6c | I | 3.1 | 0633 | 0.0509 | 1.16 | 111 | 6.8 | 4.7 | 0.838 | 0.864 | 88.3 | 78.6 | 82.9 | 43.5 |
| 6o7e | C | 3.2 | 0640 | 0.015 | 1.089 | 278 | 3.3 | 3.3 | 0.920 | 0.920 | 92.1 | 66 | 79.1 | 34.1 |
| 6o7e | D | 3.2 | 0640 | 0.015 | 1.089 | 284 | 7.2 | 7.2 | 0.847 | 0.853 | 88 | 76.4 | 78.5 | 50.7 |
| 6o7e | E | 3.2 | 0640 | 0.015 | 1.089 | 286 | 1.6 | 1.6 | 0.951 | 0.951 | 96.2 | 88 | 89.9 | 57.2 |
| 6o7h | D | 2.9 | 0641 | 0.015 | 1.089 | 285 | 2.9 | 2.9 | 0.927 | 0.927 | 94 | 76.9 | 84.2 | 40.8 |
| 6o7h | E | 2.9 | 0641 | 0.015 | 1.089 | 286 | 2.3 | 2.3 | 0.938 | 0.938 | 94.4 | 77.8 | 93.7 | 76.9 |
| 6o7i | D | 3.2 | 0642 | 0.011 | 1.089 | 272 | 3.9 | 3.9 | 0.931 | 0.931 | 94.1 | 64.8 | 83.8 | 46.1 |
| 6o7i | E | 3.2 | 0642 | 0.011 | 1.089 | 279 | 1.9 | 1.9 | 0.955 | 0.955 | 97.1 | 87.1 | 87.5 | 48 |
| 6o7i | K | 3.2 | 0642 | 0.011 | 1.089 | 275 | 23.8 | 11.3 | 0.324 | 0.781 | 76.7 | 11.4 | 61.1 | 17.3 |
| 6o7k | h | 4.2 | 0643 | 0.03 | 1.66 | 206 | 9.8 | 9.8 | 0.732 | 0.732 | 80.1 | 23 | 68 | 5 |
| 6o7k | j | 4.2 | 0643 | 0.03 | 1.66 | 218 | 12.4 | 12.4 | 0.534 | 0.543 | 71.1 | 21.9 | 69.3 | 4 |
| 6o7k | n | 4.2 | 0643 | 0.03 | 1.66 | 100 | 12.4 | 12.4 | 0.507 | 0.510 | 75 | 10.7 | 42 | 14.3 |
| 6oa9 | J | 3.9 | 0665 | 0.45 | 1.35 | 76 | 12.3 | 10.2 | 0.250 | 0.339 | 60.5 | 10.9 | 31.6 | 4.2 |
| 6oge | B | 4.4 | 20055 | 4.2 | 1.059 | 214 | 17.4 | 16.4 | 0.257 | 0.301 | 57.9 | 10.5 | 22.4 | 12.5 |
| 6oge | C | 4.4 | 20055 | 4.2 | 1.059 | 222 | 20.5 | 17.6 | 0.273 | 0.353 | 68.9 | 10.5 | 33.3 | 4.1 |

| | | | | | | | | | | | | | | |
|---|---|---|---|---|---|---|---|---|---|---|---|---|---|---|
| 6oge | D | 4.4 | 20055 | 4.2 | 1.059 | 214 | 18.8 | 17.6 | 0.238 | 0.299 | 54.2 | 5.2 | 17.8 | 7.9 |
| 6oge | E | 4.4 | 20055 | 4.2 | 1.059 | 220 | 18.7 | 17.7 | 0.320 | 0.350 | 54.1 | 11.8 | 22.3 | 6.1 |
| 6oij | H | 3.3 | 20078 | 0.01 | 1.06 | 236 | 20.2 | 15.5 | 0.538 | 0.659 | 79.2 | 11.8 | 52.1 | 8.1 |
| 6oij | R | 3.3 | 20078 | 0.01 | 1.06 | 293 | 4.4 | 4.4 | 0.900 | 0.900 | 90.1 | 71.6 | 79.9 | 20.1 |
| 6oik | H | 3.6 | 20079 | 0.01 | 1.06 | 236 | 16.8 | 16.2 | 0.494 | 0.501 | 83.1 | 13.3 | 70.3 | 19.3 |
| 6oik | R | 3.6 | 20079 | 0.01 | 1.06 | 278 | 15.8 | 10.4 | 0.440 | 0.700 | 80.2 | 4.9 | 68.7 | 6.8 |
| 6okp | M | 3.3 | 20100 | 0.022 | 1.09 | 131 | 18.2 | 16.6 | 0.311 | 0.354 | 75.6 | 14.1 | 48.1 | 6.3 |
| 6okp | N | 3.3 | 20100 | 0.022 | 1.09 | 104 | 14.2 | 14.1 | 0.300 | 0.433 | 72.1 | 5.3 | 70.2 | 13.7 |
| 6okp | O | 3.3 | 20100 | 0.022 | 1.09 | 131 | 16.1 | 15.3 | 0.445 | 0.471 | 75.6 | 8.1 | 48.1 | 14.3 |
| 6okp | P | 3.3 | 20100 | 0.022 | 1.09 | 104 | 14.7 | 13.8 | 0.316 | 0.331 | 66.3 | 5.8 | 43.3 | 6.7 |
| 6okp | Q | 3.3 | 20100 | 0.022 | 1.09 | 131 | 16.2 | 15.2 | 0.451 | 0.451 | 78.6 | 22.3 | 35.9 | 6.4 |
| 6okp | R | 3.3 | 20100 | 0.022 | 1.09 | 104 | 15.2 | 12.2 | 0.252 | 0.427 | 66.3 | 10.1 | 26.9 | 10.7 |
| 6olp | B | 4.2 | 20118 | 0.0218 | 1.03 | 143 | 3.2 | 3.2 | 0.839 | 0.839 | 88.8 | 40.9 | 85.3 | 4.9 |
| 6olp | G | 4.2 | 20118 | 0.0218 | 1.03 | 134 | 15.4 | 15.4 | 0.344 | 0.453 | 75.4 | 34.7 | 49.3 | 6.1 |
| 6olp | H | 4.2 | 20118 | 0.0218 | 1.03 | 135 | 13.4 | 13.4 | 0.511 | 0.540 | 78.5 | 30.2 | 31.9 | 9.3 |
| 6olp | I | 4.2 | 20118 | 0.0218 | 1.03 | 109 | 15.8 | 13.2 | 0.283 | 0.455 | 71.6 | 6.4 | 33.9 | 2.7 |
| 6olp | L | 4.2 | 20118 | 0.0218 | 1.03 | 111 | 16.1 | 15.0 | 0.278 | 0.344 | 69.4 | 5.2 | 39.6 | 13.6 |
| 6omf | A | 3.3 | 20090 | 0.45 | 1.3 | 233 | 2.7 | 2.7 | 0.904 | 0.910 | 90.6 | 76.3 | 78.1 | 31.9 |
| 6omf | B | 3.3 | 20090 | 0.45 | 1.3 | 223 | 11.0 | 10.1 | 0.668 | 0.707 | 89.2 | 46.7 | 58.3 | 10 |
| 6omf | E | 3.3 | 20090 | 0.45 | 1.3 | 79 | 4.3 | 4.3 | 0.856 | 0.856 | 92.4 | 31.5 | 89.9 | 54.9 |
| 6omf | F | 3.3 | 20090 | 0.45 | 1.3 | 278 | 3.1 | 3.1 | 0.918 | 0.918 | 92.1 | 57 | 75.2 | 26.8 |
| 6omf | J | 3.3 | 20090 | 0.45 | 1.3 | 127 | 15.8 | 14.9 | 0.294 | 0.453 | 67.7 | 8.1 | 51.2 | 6.2 |
| 6omv | A | 3.9 | 20132 | 0.01 | 1.066 | 115 | 13.0 | 12.1 | 0.304 | 0.484 | 68.7 | 10.1 | 45.2 | 3.8 |
| 6omv | C | 3.9 | 20132 | 0.01 | 1.066 | 114 | 14.2 | 12.2 | 0.312 | 0.441 | 64 | 15.1 | 52.6 | 10 |
| 6oo2 | M | 4.4 | 20142 | 0.011 | 1.09 | 50 | 4.4 | 4.4 | 0.438 | 0.438 | 78 | 10.3 | 66 | 9.1 |
| 6oo2 | O | 4.4 | 20142 | 0.011 | 1.09 | 50 | 4.0 | 4.0 | 0.453 | 0.453 | 74 | 10.8 | 68 | 20.6 |
| 6orn | A | 4.1 | 20175 | 0.725 | 1.436 | 133 | 16.1 | 14.9 | 0.254 | 0.309 | 75.9 | 12.9 | 27.8 | 10.8 |
| 6orn | B | 4.1 | 20175 | 0.725 | 1.436 | 107 | 10.2 | 10.2 | 0.714 | 0.714 | 77.6 | 7.2 | 45.8 | 6.1 |
| 6orn | C | 4.1 | 20175 | 0.725 | 1.436 | 124 | 8.2 | 8.2 | 0.745 | 0.745 | 78.2 | 45.4 | 85.5 | 43.4 |
| 6orn | F | 4.1 | 20175 | 0.725 | 1.436 | 133 | 16.1 | 15.6 | 0.421 | 0.421 | 65.4 | 5.7 | 33.1 | 15.9 |
| 6orn | I | 4.1 | 20175 | 0.725 | 1.436 | 107 | 16.5 | 14.3 | 0.310 | 0.509 | 79.4 | 7.1 | 68.2 | 11 |
| 6orn | J | 4.1 | 20175 | 0.725 | 1.436 | 124 | 7.4 | 7.4 | 0.742 | 0.773 | 78.2 | 29.9 | 70.2 | 8 |
| 6orn | O | 4.1 | 20175 | 0.725 | 1.436 | 133 | 19.2 | 17.1 | 0.286 | 0.387 | 68.4 | 7.7 | 37.6 | 6 |
| 6orn | P | 4.1 | 20175 | 0.725 | 1.436 | 107 | 15.5 | 11.5 | 0.464 | 0.692 | 79.4 | 8.2 | 79.4 | 7.1 |
| 6orn | Q | 4.1 | 20175 | 0.725 | 1.436 | 124 | 6.0 | 4.7 | 0.788 | 0.788 | 86.3 | 22.4 | 83.9 | 33.7 |
| 6oro | D | 3.9 | 20176 | 1.02 | 1.436 | 124 | 2.4 | 2.4 | 0.879 | 0.879 | 94.4 | 73.5 | 65.3 | 9.9 |
| 6oro | E | 3.9 | 20176 | 1.02 | 1.436 | 108 | 13.6 | 12.6 | 0.326 | 0.574 | 76.9 | 12 | 60.2 | 3.1 |

| | | | | | | | | | | | | | | |
|---|---|---|---|---|---|---|---|---|---|---|---|---|---|---|
| 6oro | H | 3.9 | 20176 | 1.02 | 1.436 | 124 | 10.8 | 10.3 | 0.684 | 0.693 | 77.4 | 42.7 | 58.1 | 2.8 |
| 6oro | J | 3.9 | 20176 | 1.02 | 1.436 | 124 | 6.2 | 6.0 | 0.758 | 0.829 | 91.9 | 66.7 | 62.1 | 6.5 |
| 6oro | K | 3.9 | 20176 | 1.02 | 1.436 | 108 | 12.8 | 12.5 | 0.578 | 0.578 | 89.8 | 48.5 | 59.3 | 3.1 |
| 6oro | L | 3.9 | 20176 | 1.02 | 1.436 | 108 | 13.9 | 10.3 | 0.361 | 0.610 | 84.3 | 8.8 | 57.4 | 4.8 |
| 6orp | J | 4.4 | 20177 | 1.02 | 1.436 | 124 | 17.4 | 14.0 | 0.285 | 0.384 | 71 | 10.2 | 30.6 | 7.9 |
| 6orq | D | 4.4 | 20178 | 1.02 | 1.436 | 122 | 17.1 | 13.7 | 0.278 | 0.312 | 68 | 7.2 | 55.7 | 7.4 |
| 6orq | E | 4.4 | 20178 | 1.02 | 1.436 | 111 | 16.8 | 14.3 | 0.338 | 0.430 | 69.4 | 13 | 28.8 | 3.1 |
| 6orq | H | 4.4 | 20178 | 1.02 | 1.436 | 122 | 17.8 | 15.0 | 0.376 | 0.414 | 68 | 8.4 | 50.8 | 12.9 |
| 6orq | J | 4.4 | 20178 | 1.02 | 1.436 | 122 | 17.0 | 14.5 | 0.413 | 0.413 | 70.5 | 7 | 50 | 9.8 |
| 6orq | K | 4.4 | 20178 | 1.02 | 1.436 | 111 | 16.6 | 15.7 | 0.356 | 0.356 | 65.8 | 9.6 | 52.3 | 6.9 |
| 6orq | L | 4.4 | 20178 | 1.02 | 1.436 | 111 | 14.8 | 12.2 | 0.275 | 0.370 | 69.4 | 6.5 | 33.3 | 2.7 |
| 6os9 | D | 3 | 20180 | 0.065 | 1.06 | 231 | 17.4 | 16.8 | 0.304 | 0.555 | 77.9 | 8.3 | 64.5 | 6.7 |
| 6osy | 5 | 4.3 | 20189 | 0.45 | 1.07325 | 132 | 20.2 | 17.2 | 0.341 | 0.348 | 62.9 | 6 | 43.2 | 8.8 |
| 6osy | 6 | 4.3 | 20189 | 0.45 | 1.07325 | 105 | 14.8 | 12.2 | 0.258 | 0.350 | 78.1 | 4.9 | 43.8 | 6.5 |
| 6osy | 7 | 4.3 | 20189 | 0.45 | 1.07325 | 102 | 14.3 | 13.3 | 0.251 | 0.342 | 73.5 | 9.3 | 20.6 | 0 |
| 6osy | 8 | 4.3 | 20189 | 0.45 | 1.07325 | 128 | 16.2 | 15.5 | 0.517 | 0.517 | 74.2 | 27.4 | 53.9 | 0 |
| 6osy | C | 4.3 | 20189 | 0.45 | 1.07325 | 132 | 17.1 | 15.8 | 0.295 | 0.300 | 58.3 | 31.2 | 19.7 | 0 |
| 6osy | D | 4.3 | 20189 | 0.45 | 1.07325 | 105 | 13.4 | 13.3 | 0.288 | 0.389 | 69.5 | 6.8 | 54.3 | 10.5 |
| 6osy | E | 4.3 | 20189 | 0.45 | 1.07325 | 102 | 11.2 | 11.2 | 0.278 | 0.332 | 69.6 | 8.5 | 34.3 | 2.9 |
| 6osy | F | 4.3 | 20189 | 0.45 | 1.07325 | 128 | 15.7 | 12.9 | 0.331 | 0.689 | 74.2 | 9.5 | 59.4 | 10.5 |
| 6osy | J | 4.3 | 20189 | 0.45 | 1.07325 | 104 | 13.4 | 12.9 | 0.250 | 0.287 | 59.6 | 6.5 | 26.9 | 3.6 |
| 6osy | L | 4.3 | 20189 | 0.45 | 1.07325 | 104 | 16.0 | 14.0 | 0.278 | 0.296 | 60.6 | 6.3 | 26 | 14.8 |
| 6osy | M | 4.3 | 20189 | 0.45 | 1.07325 | 132 | 18.3 | 14.3 | 0.261 | 0.345 | 59.1 | 7.7 | 19.7 | 11.5 |
| 6osy | N | 4.3 | 20189 | 0.45 | 1.07325 | 105 | 15.5 | 13.2 | 0.329 | 0.329 | 66.7 | 10 | 40 | 2.4 |
| 6osy | O | 4.3 | 20189 | 0.45 | 1.07325 | 102 | 12.1 | 11.9 | 0.289 | 0.327 | 70.6 | 9.7 | 38.2 | 2.6 |
| 6osy | P | 4.3 | 20189 | 0.45 | 1.07325 | 128 | 15.9 | 13.7 | 0.305 | 0.383 | 68.8 | 6.8 | 57.8 | 4.1 |
| 6osy | S | 4.3 | 20189 | 0.45 | 1.07325 | 104 | 13.2 | 12.8 | 0.312 | 0.317 | 63.5 | 6.1 | 41.3 | 7 |
| 6ot0 | H | 3.8 | 20190 | 0.33 | 1.07 | 126 | 17.3 | 16.3 | 0.250 | 0.322 | 59.5 | 12 | 41.3 | 7.7 |
| 6ot0 | L | 3.8 | 20190 | 0.33 | 1.07 | 104 | 15.2 | 9.9 | 0.330 | 0.654 | 76.9 | 13.8 | 60.6 | 7.9 |
| 6ot1 | B | 3.5 | 20191 | 0.4 | 1.07325 | 132 | 10.2 | 6.3 | 0.755 | 0.825 | 82.6 | 48.6 | 69.7 | 43.5 |
| 6ot1 | D | 3.5 | 20191 | 0.4 | 1.07325 | 132 | 4.2 | 4.2 | 0.849 | 0.849 | 89.4 | 83.9 | 65.2 | 32.6 |
| 6ot1 | F | 3.5 | 20191 | 0.4 | 1.07325 | 132 | 16.9 | 14.5 | 0.348 | 0.380 | 77.3 | 7.8 | 28.8 | 10.5 |
| 6ot1 | I | 3.5 | 20191 | 0.4 | 1.07325 | 105 | 13.2 | 4.6 | 0.291 | 0.781 | 81.9 | 11.6 | 55.2 | 6.9 |
| 6ot1 | J | 3.5 | 20191 | 0.4 | 1.07325 | 128 | 2.0 | 2.0 | 0.906 | 0.906 | 93.8 | 84.2 | 84.4 | 8.3 |
| 6ot1 | K | 3.5 | 20191 | 0.4 | 1.07325 | 102 | 10.0 | 7.9 | 0.754 | 0.755 | 92.2 | 24.5 | 58.8 | 8.3 |
| 6ot1 | m | 3.5 | 20191 | 0.4 | 1.07325 | 132 | 18.9 | 17.7 | 0.353 | 0.353 | 76.5 | 7.9 | 21.2 | 0 |
| 6ot1 | n | 3.5 | 20191 | 0.4 | 1.07325 | 105 | 14.7 | 11.5 | 0.571 | 0.754 | 81.9 | 5.8 | 45.7 | 8.3 |

| | | | | | | | | | | | | | | |
|---|---|---|---|---|---|---|---|---|---|---|---|---|---|---|
| 6ot1 | O | 3.5 | 20191 | 0.4 | 1.07325 | 132 | 7.7 | 6.8 | 0.751 | 0.772 | 79.5 | 33.3 | 69.7 | 35.9 |
| 6ot1 | q | 3.5 | 20191 | 0.4 | 1.07325 | 128 | 1.9 | 1.9 | 0.902 | 0.902 | 94.5 | 76 | 81.2 | 22.1 |
| 6ot1 | Q | 3.5 | 20191 | 0.4 | 1.07325 | 132 | 16.4 | 11.4 | 0.530 | 0.591 | 78 | 6.8 | 36.4 | 6.2 |
| 6ot1 | r | 3.5 | 20191 | 0.4 | 1.07325 | 102 | 2.3 | 2.3 | 0.870 | 0.870 | 94.1 | 55.2 | 82.4 | 13.1 |
| 6ot1 | R | 3.5 | 20191 | 0.4 | 1.07325 | 105 | 3.2 | 3.2 | 0.833 | 0.833 | 91.4 | 55.2 | 64.8 | 5.9 |
| 6ot1 | S | 3.5 | 20191 | 0.4 | 1.07325 | 128 | 2.5 | 2.5 | 0.900 | 0.900 | 93.8 | 66.7 | 84.4 | 7.4 |
| 6ot1 | T | 3.5 | 20191 | 0.4 | 1.07325 | 102 | 5.4 | 4.3 | 0.876 | 0.876 | 92.2 | 23.4 | 76.5 | 9 |
| 6owo | N | 3.2 | 20215 | 0.03 | 1.16 | 122 | 4.7 | 4.7 | 0.808 | 0.808 | 82.8 | 50.5 | 77 | 37.2 |
| 6owo | S | 3.2 | 20215 | 0.03 | 1.16 | 141 | 1.2 | 1.2 | 0.948 | 0.948 | 97.2 | 93.4 | 94.3 | 75.2 |
| 6oxl | N | 3.5 | 20220 | 1.1 | 1.16 | 124 | 12.8 | 9.4 | 0.700 | 0.721 | 81.5 | 54.5 | 69.4 | 33.7 |
| 6oxl | S | 3.5 | 20220 | 1.1 | 1.16 | 141 | 1.9 | 1.9 | 0.938 | 0.938 | 95.7 | 81.5 | 91.5 | 48.8 |
| 6oya | N | 3.3 | 20223 | 4 | 1.06 | 128 | 16.0 | 10.5 | 0.349 | 0.649 | 78.1 | 12 | 71.9 | 30.4 |
| 6p07 | A | 3.2 | 20226 | 1.12 | 1.15 | 270 | 6.8 | 6.8 | 0.780 | 0.798 | 84.4 | 58.3 | 87.8 | 19.4 |
| 6p18 | A | 3.5 | 20233 | 0.0286 | 1.066 | 230 | 3.4 | 3.4 | 0.932 | 0.932 | 94.3 | 58.5 | 81.7 | 25 |
| 6p18 | B | 3.5 | 20233 | 0.0286 | 1.066 | 228 | 12.9 | 11.7 | 0.584 | 0.587 | 86 | 48.5 | 71.5 | 14.7 |
| 6p18 | P | 3.5 | 20233 | 0.0286 | 1.066 | 145 | 17.4 | 7.7 | 0.298 | 0.654 | 75.2 | 9.2 | 48.3 | 5.7 |
| 6p19 | A | 3.8 | 20234 | 0.0204 | 1.024 | 230 | 11.6 | 11.6 | 0.542 | 0.690 | 83 | 29.3 | 79.6 | 10.4 |
| 6p19 | B | 3.8 | 20234 | 0.0204 | 1.024 | 228 | 14.7 | 13.9 | 0.624 | 0.624 | 82.9 | 29.1 | 56.6 | 7 |
| 6p1h | C | 3.2 | 20235 | 3.5 | 0.8549 | 111 | 8.6 | 4.4 | 0.748 | 0.768 | 82.9 | 29.3 | 75.7 | 3.6 |
| 6p4g | B | 3.1 | 20248 | 0.024 | 1.233 | 217 | 5.0 | 5.0 | 0.835 | 0.847 | 88.9 | 49.2 | 77 | 46.7 |
| 6p4g | C | 3.1 | 20248 | 0.024 | 1.233 | 213 | 3.9 | 3.9 | 0.908 | 0.908 | 93 | 47.5 | 69 | 6.8 |
| 6p4g | D | 3.1 | 20248 | 0.024 | 1.233 | 221 | 3.0 | 3.0 | 0.911 | 0.911 | 93.2 | 64.6 | 77.8 | 18.6 |
| 6p4h | B | 3.2 | 20249 | 0.024 | 1.233 | 217 | 11.6 | 11.2 | 0.763 | 0.763 | 83.9 | 55.5 | 74.2 | 13.7 |
| 6p4h | C | 3.2 | 20249 | 0.024 | 1.233 | 213 | 3.8 | 3.6 | 0.910 | 0.910 | 93.9 | 68 | 76.5 | 35.6 |
| 6p4h | L | 3.2 | 20249 | 0.024 | 1.233 | 96 | 3.4 | 3.4 | 0.790 | 0.792 | 86.5 | 55.4 | 71.9 | 29 |
| 6p4h | Z | 3.2 | 20249 | 0.024 | 1.233 | 124 | 3.6 | 3.6 | 0.855 | 0.855 | 92.7 | 58.3 | 87.9 | 25.7 |
| 6p62 | C | 3.6 | 20259 | 0.08 | 1.03 | 119 | 15.7 | 13.1 | 0.269 | 0.408 | 69.7 | 8.4 | 42 | 2 |
| 6p62 | D | 3.6 | 20259 | 0.08 | 1.03 | 108 | 9.0 | 9.0 | 0.468 | 0.468 | 80.6 | 6.9 | 74.1 | 6.2 |
| 6p62 | F | 3.6 | 20259 | 0.08 | 1.03 | 119 | 16.4 | 15.0 | 0.389 | 0.394 | 75.6 | 6.7 | 51.3 | 6.6 |
| 6p62 | G | 3.6 | 20259 | 0.08 | 1.03 | 108 | 10.1 | 9.9 | 0.783 | 0.783 | 84.3 | 38.5 | 73.1 | 8.9 |
| 6p62 | H | 3.6 | 20259 | 0.08 | 1.03 | 119 | 14.7 | 13.8 | 0.303 | 0.426 | 73.1 | 12.6 | 32.8 | 0 |
| 6p62 | L | 3.6 | 20259 | 0.08 | 1.03 | 108 | 15.3 | 9.3 | 0.311 | 0.498 | 75.9 | 8.5 | 74.1 | 6.2 |
| 6p65 | C | 3.9 | 20260 | 0.06 | 1.15 | 126 | 15.7 | 14.7 | 0.229 | 0.341 | 65.9 | 6 | 40.5 | 3.9 |
| 6p65 | D | 3.9 | 20260 | 0.06 | 1.15 | 110 | 16.9 | 15.1 | 0.264 | 0.398 | 70.9 | 7.7 | 29.1 | 12.5 |
| 6p65 | F | 3.9 | 20260 | 0.06 | 1.15 | 126 | 16.8 | 15.4 | 0.336 | 0.392 | 70.6 | 7.9 | 38.1 | 0 |
| 6p65 | G | 3.9 | 20260 | 0.06 | 1.15 | 110 | 15.6 | 14.3 | 0.267 | 0.358 | 70 | 13 | 32.7 | 11.1 |
| 6p65 | H | 3.9 | 20260 | 0.06 | 1.15 | 126 | 14.9 | 14.4 | 0.267 | 0.333 | 62.7 | 5.1 | 45.2 | 8.8 |

| | | | | | | | | | | | | | | |
|---|---|---|---|---|---|---|---|---|---|---|---|---|---|---|
| 6p65 | L | 3.9 | 20260 | 0.06 | 1.15 | 110 | 16.8 | 14.8 | 0.261 | 0.341 | 58.2 | 7.8 | 33.6 | 8.1 |
| 6p6w | A | 4 | 20265 | 3 | 1.056 | 229 | 13.5 | 5.8 | 0.637 | 0.779 | 81.2 | 11.8 | 58.1 | 15 |
| 6p6w | B | 4 | 20265 | 3 | 1.056 | 229 | 12.5 | 4.8 | 0.629 | 0.808 | 80.3 | 25 | 61.1 | 8.6 |
| 6p6w | D | 4 | 20265 | 3 | 1.056 | 229 | 8.8 | 5.6 | 0.762 | 0.762 | 76 | 24.7 | 76.4 | 7.4 |
| 6p7m | C | 3 | 20266 | 0.6 | 0.9 | 98 | 2.2 | 2.2 | 0.871 | 0.871 | 92.9 | 70.3 | 83.7 | 7.3 |
| 6pe2 | B | 4 | 20321 | 0.01 | 1.16 | 118 | 2.9 | 2.9 | 0.843 | 0.843 | 90.7 | 65.4 | 39 | 8.7 |
| 6pe2 | H | 4 | 20321 | 0.01 | 1.16 | 118 | 3.3 | 3.3 | 0.855 | 0.855 | 91.5 | 45.4 | 84.7 | 39 |
| 6pek | A | 4.2 | 20327 | 0.0448 | 1.056 | 283 | 22.4 | 21.8 | 0.366 | 0.506 | 76.3 | 9.7 | 66.8 | 9 |
| 6pek | B | 4.2 | 20327 | 0.0448 | 1.056 | 288 | 4.9 | 4.9 | 0.874 | 0.874 | 88.2 | 51.2 | 71.9 | 2.9 |
| 6pek | C | 4.2 | 20327 | 0.0448 | 1.056 | 288 | 22.1 | 17.6 | 0.501 | 0.679 | 75.7 | 7.8 | 41.3 | 8.4 |
| 6pek | D | 4.2 | 20327 | 0.0448 | 1.056 | 293 | 22.9 | 11.9 | 0.514 | 0.685 | 74.7 | 23.3 | 67.9 | 6.5 |
| 6peq | E | 3 | 20330 | 0.0299 | 1.066 | 132 | 15.4 | 6.6 | 0.477 | 0.673 | 81.8 | 29.6 | 88.6 | 88 |
| 6peq | G | 3 | 20330 | 0.0299 | 1.066 | 132 | 3.7 | 3.7 | 0.846 | 0.846 | 87.9 | 64.7 | 85.6 | 60.2 |
| 6ppd | 6 | 3.7 | 20433 | 0.02 | 1.03 | 294 | 3.9 | 3.9 | 0.910 | 0.910 | 91.2 | 65.3 | 49.7 | 9.6 |
| 6ppd | 7 | 3.7 | 20433 | 0.02 | 1.03 | 291 | 20.6 | 8.1 | 0.415 | 0.797 | 81.4 | 36.7 | 68.7 | 23 |
| 6ppf | Y | 3.4 | 20435 | 0.0732 | 1.073 | 65 | 2.7 | 2.7 | 0.831 | 0.831 | 93.8 | 55.7 | 80 | 34.6 |
| 6pph | 7 | 3.8 | 20436 | 0.022 | 1.03 | 294 | 25.9 | 22.5 | 0.576 | 0.576 | 90.8 | 30 | 88.1 | 29.3 |
| 6ppk | K | 4.4 | 20441 | 0.017 | 1.073 | 122 | 15.3 | 12.4 | 0.253 | 0.314 | 74.6 | 6.6 | 30.3 | 8.1 |
| 6ptj | A | 3.8 | 20471 | 0.034 | 1.074 | 208 | 5.8 | 4.0 | 0.855 | 0.855 | 87 | 35.4 | 76.9 | 26.2 |
| 6pv7 | F | 3.3 | 20487 | 0.0275 | 1.07 | 117 | 5.0 | 1.9 | 0.849 | 0.888 | 88.9 | 69.2 | 52.1 | 3.3 |
| 6pv7 | G | 3.3 | 20487 | 0.0275 | 1.07 | 105 | 2.4 | 2.4 | 0.895 | 0.895 | 95.2 | 78 | 59 | 8.1 |
| 6pv7 | H | 3.3 | 20487 | 0.0275 | 1.07 | 117 | 1.5 | 1.5 | 0.938 | 0.938 | 98.3 | 90.4 | 76.1 | 7.9 |
| 6pv7 | I | 3.3 | 20487 | 0.0275 | 1.07 | 105 | 1.6 | 1.6 | 0.942 | 0.942 | 98.1 | 80.6 | 87.6 | 25 |
| 6pvk | Y | 3.4 | 20491 | 0.076 | 1.073 | 63 | 3.6 | 3.6 | 0.838 | 0.838 | 92.1 | 36.2 | 92.1 | 48.3 |
| 6pwc | H | 4.9 | 20505 | 0.017 | 1.031 | 194 | 16.9 | 15.9 | 0.259 | 0.325 | 62.4 | 4.1 | 20.6 | 12.5 |
| 6pwc | L | 4.9 | 20505 | 0.017 | 1.031 | 183 | 19.1 | 17.8 | 0.276 | 0.391 | 61.2 | 11.6 | 26.2 | 2.1 |
| 6px3 | S | 4.1 | 20517 | 0.01 | 1.04 | 233 | 23.0 | 18.2 | 0.277 | 0.289 | 52.8 | 4.9 | 8.2 | 10.5 |
| 6pzw | E | 3 | 20538 | 0.985 | 1.03 | 108 | 6.6 | 6.6 | 0.784 | 0.784 | 91.7 | 73.7 | 93.5 | 36.6 |
| 6pzw | F | 3 | 20538 | 0.985 | 1.03 | 122 | 1.1 | 1.1 | 0.957 | 0.957 | 99.2 | 96.7 | 91.8 | 25 |
| 6pzw | G | 3 | 20538 | 0.985 | 1.03 | 108 | 2.9 | 2.9 | 0.887 | 0.887 | 95.4 | 63.1 | 93.5 | 65.3 |
| 6pzw | H | 3 | 20538 | 0.985 | 1.03 | 122 | 1.3 | 1.3 | 0.951 | 0.951 | 98.4 | 94.2 | 96.7 | 24.6 |
| 6pzw | I | 3 | 20538 | 0.985 | 1.03 | 108 | 2.4 | 2.4 | 0.905 | 0.905 | 95.4 | 80.6 | 92.6 | 49 |
| 6pzw | J | 3 | 20538 | 0.985 | 1.03 | 122 | 7.4 | 3.3 | 0.832 | 0.893 | 93.4 | 90.4 | 92.6 | 40.7 |
| 6pzw | K | 3 | 20538 | 0.985 | 1.03 | 122 | 7.0 | 5.7 | 0.787 | 0.895 | 90.2 | 64.5 | 91 | 47.7 |
| 6pzw | L | 3 | 20538 | 0.985 | 1.03 | 108 | 2.1 | 2.1 | 0.916 | 0.916 | 97.2 | 77.1 | 95.4 | 8.7 |
| 6pzy | C | 3.2 | 20540 | 0.488 | 1.03 | 129 | 2.7 | 2.7 | 0.902 | 0.902 | 94.6 | 61.5 | 89.1 | 9.6 |
| 6pzy | D | 3.2 | 20540 | 0.488 | 1.03 | 109 | 3.0 | 3.0 | 0.886 | 0.886 | 94.5 | 41.7 | 85.3 | 31.2 |

| | | | | | | | | | | | | | | |
|---|---|---|---|---|---|---|---|---|---|---|---|---|---|---|
| 6pzy | G | 3.2 | 20540 | 0.488 | 1.03 | 129 | 2.0 | 2.0 | 0.924 | 0.924 | 96.1 | 80.6 | 85.3 | 11.8 |
| 6pzy | H | 3.2 | 20540 | 0.488 | 1.03 | 129 | 1.6 | 1.6 | 0.921 | 0.921 | 96.1 | 87.1 | 96.1 | 10.5 |
| 6pzy | I | 3.2 | 20540 | 0.488 | 1.03 | 129 | 3.4 | 3.4 | 0.927 | 0.927 | 96.9 | 69.6 | 79.8 | 7.8 |
| 6pzy | J | 3.2 | 20540 | 0.488 | 1.03 | 109 | 3.8 | 3.8 | 0.868 | 0.868 | 92.7 | 51.5 | 83.5 | 7.7 |
| 6pzy | K | 3.2 | 20540 | 0.488 | 1.03 | 109 | 2.0 | 2.0 | 0.915 | 0.915 | 96.3 | 75.2 | 59.6 | 9.2 |
| 6pzy | L | 3.2 | 20540 | 0.488 | 1.03 | 109 | 2.3 | 2.3 | 0.900 | 0.900 | 96.3 | 67.6 | 93.6 | 11.8 |
| 6pzz | E | 3.6 | 20541 | 0.641 | 1.03 | 108 | 5.7 | 5.7 | 0.827 | 0.827 | 90.7 | 42.9 | 81.5 | 12.5 |
| 6pzz | F | 3.6 | 20541 | 0.641 | 1.03 | 120 | 15.4 | 13.4 | 0.310 | 0.366 | 72.5 | 9.2 | 61.7 | 6.8 |
| 6pzz | G | 3.6 | 20541 | 0.641 | 1.03 | 108 | 3.4 | 3.4 | 0.831 | 0.831 | 88 | 65.3 | 79.6 | 16.3 |
| 6pzz | H | 3.6 | 20541 | 0.641 | 1.03 | 120 | 13.7 | 13.7 | 0.559 | 0.559 | 76.7 | 31.5 | 35.8 | 7 |
| 6pzz | I | 3.6 | 20541 | 0.641 | 1.03 | 120 | 14.7 | 12.7 | 0.408 | 0.498 | 77.5 | 6.5 | 55 | 7.6 |
| 6pzz | J | 3.6 | 20541 | 0.641 | 1.03 | 108 | 9.8 | 3.4 | 0.700 | 0.849 | 86.1 | 48.4 | 83.3 | 5.6 |
| 6pzz | K | 3.6 | 20541 | 0.641 | 1.03 | 120 | 13.9 | 13.9 | 0.335 | 0.438 | 81.7 | 8.2 | 65.8 | 8.9 |
| 6pzz | L | 3.6 | 20541 | 0.641 | 1.03 | 108 | 3.3 | 3.3 | 0.858 | 0.858 | 93.5 | 54.5 | 78.7 | 12.9 |
| 6q0j | A | 4.9 | 20550 | 0.0285 | 1.11 | 280 | 15.6 | 13.5 | 0.608 | 0.608 | 63.6 | 11.2 | 38.2 | 1.9 |
| 6q0j | B | 4.9 | 20550 | 0.0285 | 1.11 | 281 | 14.3 | 11.7 | 0.647 | 0.660 | 68.3 | 15.1 | 60.9 | 11.7 |
| 6q0j | X | 4.9 | 20550 | 0.0285 | 1.11 | 229 | 11.3 | 10.2 | 0.428 | 0.620 | 69.4 | 13.8 | 49.8 | 7 |
| 6q0j | Y | 4.9 | 20550 | 0.0285 | 1.11 | 229 | 18.5 | 13.0 | 0.396 | 0.571 | 68.1 | 7.1 | 38.4 | 2.3 |
| 6q2n | C | 4.4 | 20575 | 0.03 | 1.07 | 200 | 18.4 | 14.6 | 0.276 | 0.315 | 55.5 | 5.4 | 26.5 | 5.7 |
| 6q2n | D | 4.4 | 20575 | 0.03 | 1.07 | 200 | 17.9 | 12.6 | 0.316 | 0.323 | 56 | 3.6 | 41.5 | 8.4 |
| 6q2r | C | 4.3 | 20578 | 0.015 | 1.07 | 280 | 21.4 | 19.7 | 0.298 | 0.361 | 59.3 | 6 | 45.4 | 8.7 |
| 6q2r | D | 4.3 | 20578 | 0.015 | 1.07 | 280 | 23.3 | 20.5 | 0.320 | 0.429 | 53.9 | 4.6 | 17.9 | 6 |
| 6q2r | W | 4.3 | 20578 | 0.015 | 1.07 | 280 | 21.4 | 19.4 | 0.227 | 0.276 | 61.1 | 3.5 | 23.9 | 3 |
| 6q2s | C | 3.8 | 20579 | 0.036 | 1.07 | 199 | 17.1 | 8.2 | 0.464 | 0.782 | 68.8 | 5.8 | 67.8 | 7.4 |
| 6q2s | D | 3.8 | 20579 | 0.036 | 1.07 | 199 | 15.1 | 13.7 | 0.377 | 0.428 | 67.3 | 9 | 65.3 | 9.2 |
| 6qee | B | 3.9 | 4536 | 0.02 | 0.84 | 220 | 12.1 | 10.2 | 0.656 | 0.731 | 88.2 | 28.4 | 65.5 | 3.5 |
| 6qee | C | 3.9 | 4536 | 0.02 | 0.84 | 225 | 9.8 | 9.8 | 0.725 | 0.725 | 84.9 | 48.7 | 64.9 | 15.1 |
| 6qel | G | 3.9 | 4537 | 0.05 | 1.07 | 229 | 14.2 | 11.8 | 0.533 | 0.633 | 83.8 | 12.5 | 79 | 6.1 |
| 6qel | H | 3.9 | 4537 | 0.05 | 1.07 | 237 | 16.6 | 7.0 | 0.524 | 0.856 | 82.3 | 27.7 | 76.8 | 7.1 |
| 6qel | I | 3.9 | 4537 | 0.05 | 1.07 | 237 | 15.8 | 4.8 | 0.470 | 0.832 | 84.8 | 31.3 | 76.4 | 14.4 |
| 6qel | J | 3.9 | 4537 | 0.05 | 1.07 | 237 | 3.2 | 3.2 | 0.900 | 0.900 | 92 | 69.3 | 80.2 | 36.3 |
| 6qel | K | 3.9 | 4537 | 0.05 | 1.07 | 237 | 12.5 | 12.5 | 0.686 | 0.697 | 80.2 | 24.2 | 73.4 | 8 |
| 6qel | L | 3.9 | 4537 | 0.05 | 1.07 | 237 | 18.4 | 13.9 | 0.453 | 0.453 | 68.8 | 4.9 | 54.4 | 2.3 |
| 6qem | G | 3.4 | 4538 | 0.05 | 1.047 | 239 | 2.6 | 2.6 | 0.911 | 0.911 | 92.9 | 73.9 | 85.4 | 40.2 |
| 6qem | H | 3.4 | 4538 | 0.05 | 1.047 | 239 | 2.4 | 2.4 | 0.939 | 0.939 | 95.4 | 74.6 | 80.8 | 6.2 |
| 6qem | I | 3.4 | 4538 | 0.05 | 1.047 | 239 | 2.9 | 2.9 | 0.893 | 0.893 | 92.1 | 60.5 | 75.7 | 28.7 |
| 6qem | J | 3.4 | 4538 | 0.05 | 1.047 | 239 | 9.3 | 8.1 | 0.690 | 0.760 | 82.8 | 40.9 | 68.6 | 26.2 |

| | | | | | | | | | | | | | | |
|---|---|---|---|---|---|---|---|---|---|---|---|---|---|---|
| 6qem | L | 3.4 | 4538 | 0.05 | 1.047 | 239 | 13.3 | 13.3 | 0.618 | 0.709 | 83.7 | 26.5 | 65.3 | 7.1 |
| 6qex | B | 3.6 | 4539 | 0.008 | 0.84 | 220 | 7.2 | 7.2 | 0.843 | 0.843 | 86.4 | 57.9 | 62.3 | 6.6 |
| 6qex | C | 3.6 | 4539 | 0.008 | 0.84 | 225 | 14.3 | 8.2 | 0.682 | 0.848 | 88.4 | 61.3 | 69.8 | 11.5 |
| 6qi9 | D | 4.6 | 4553 | 0.01 | 1.07 | 268 | 25.5 | 17.5 | 0.244 | 0.382 | 63.4 | 5.9 | 50.4 | 5.2 |
| 6qld | O | 4.2 | 4579 | 0.017 | 1.09 | 205 | 8.0 | 6.6 | 0.772 | 0.805 | 88.3 | 49.7 | 74.6 | 29.4 |
| 6qld | Y | 4.2 | 4579 | 0.017 | 1.09 | 223 | 4.6 | 4.6 | 0.812 | 0.819 | 83 | 24.9 | 81.2 | 5.5 |
| 6qld | Z | 4.2 | 4579 | 0.017 | 1.09 | 140 | 18.5 | 4.0 | 0.426 | 0.759 | 82.9 | 6.9 | 40.7 | 7 |
| 6qle | Z | 3.6 | 4580 | 0.026 | 1.09 | 140 | 9.1 | 4.4 | 0.574 | 0.816 | 83.6 | 45.3 | 82.9 | 44 |
| 6qm7 | A | 2.8 | 4590 | 0.0667 | 1.07 | 244 | 1.2 | 1.2 | 0.970 | 0.970 | 98.8 | 92.1 | 95.9 | 99.1 |
| 6qm7 | B | 2.8 | 4590 | 0.0667 | 1.07 | 229 | 1.7 | 1.7 | 0.957 | 0.957 | 97.4 | 87.4 | 89.5 | 77.6 |
| 6qm7 | C | 2.8 | 4590 | 0.0667 | 1.07 | 276 | 1.7 | 1.7 | 0.957 | 0.957 | 96.7 | 88 | 89.1 | 78 |
| 6qm7 | D | 2.8 | 4590 | 0.0667 | 1.07 | 239 | 2.3 | 2.3 | 0.940 | 0.940 | 94.6 | 81.9 | 85.4 | 84.8 |
| 6qm7 | E | 2.8 | 4590 | 0.0667 | 1.07 | 229 | 2.7 | 2.7 | 0.939 | 0.939 | 95.2 | 76.1 | 88.2 | 66.3 |
| 6qm7 | F | 2.8 | 4590 | 0.0667 | 1.07 | 238 | 2.0 | 2.0 | 0.950 | 0.950 | 96.2 | 82.1 | 90.3 | 76.7 |
| 6qm7 | K | 2.8 | 4590 | 0.0667 | 1.07 | 206 | 1.8 | 1.8 | 0.947 | 0.947 | 95.1 | 81.1 | 93.7 | 69.4 |
| 6qm7 | O | 2.8 | 4590 | 0.0667 | 1.07 | 244 | 1.8 | 1.8 | 0.960 | 0.960 | 98 | 78.7 | 94.3 | 90 |
| 6qm7 | P | 2.8 | 4590 | 0.0667 | 1.07 | 229 | 1.8 | 1.8 | 0.951 | 0.951 | 96.5 | 85.1 | 89.5 | 79.5 |
| 6qm7 | Q | 2.8 | 4590 | 0.0667 | 1.07 | 276 | 1.8 | 1.8 | 0.954 | 0.954 | 95.7 | 86.7 | 47.1 | 6.2 |
| 6qm7 | R | 2.8 | 4590 | 0.0667 | 1.07 | 239 | 1.9 | 1.9 | 0.945 | 0.945 | 96.2 | 79.1 | 90.8 | 78.3 |
| 6qm7 | S | 2.8 | 4590 | 0.0667 | 1.07 | 229 | 2.8 | 2.8 | 0.942 | 0.942 | 96.1 | 81.8 | 90 | 63.1 |
| 6qm7 | T | 2.8 | 4590 | 0.0667 | 1.07 | 238 | 2.0 | 2.0 | 0.956 | 0.956 | 97.9 | 89.3 | 91.2 | 77 |
| 6qm7 | Y | 2.8 | 4590 | 0.0667 | 1.07 | 206 | 2.4 | 2.4 | 0.947 | 0.947 | 96.6 | 73.9 | 94.2 | 91.2 |
| 6qm8 | A | 3.3 | 4591 | 0.0545 | 1.07 | 244 | 1.5 | 1.5 | 0.954 | 0.954 | 97.5 | 86.6 | 89.8 | 67.1 |
| 6qm8 | B | 3.3 | 4591 | 0.0545 | 1.07 | 229 | 1.6 | 1.6 | 0.957 | 0.957 | 97.8 | 86.6 | 86.9 | 46.7 |
| 6qm8 | C | 3.3 | 4591 | 0.0545 | 1.07 | 276 | 2.0 | 2.0 | 0.943 | 0.943 | 95.3 | 81.7 | 86.2 | 31.1 |
| 6qm8 | D | 3.3 | 4591 | 0.0545 | 1.07 | 239 | 1.6 | 1.6 | 0.952 | 0.952 | 97.1 | 87.1 | 83.7 | 50 |
| 6qm8 | E | 3.3 | 4591 | 0.0545 | 1.07 | 229 | 1.9 | 1.9 | 0.942 | 0.942 | 96.5 | 86.9 | 84.3 | 45.6 |
| 6qm8 | F | 3.3 | 4591 | 0.0545 | 1.07 | 238 | 2.3 | 2.3 | 0.928 | 0.928 | 94.1 | 76.3 | 78.2 | 36.6 |
| 6qm8 | K | 3.3 | 4591 | 0.0545 | 1.07 | 206 | 3.5 | 3.5 | 0.938 | 0.938 | 95.6 | 67.5 | 88.8 | 47.5 |
| 6qm8 | O | 3.3 | 4591 | 0.0545 | 1.07 | 244 | 1.5 | 1.5 | 0.954 | 0.954 | 97.1 | 89 | 79.1 | 41.5 |
| 6qm8 | P | 3.3 | 4591 | 0.0545 | 1.07 | 229 | 1.8 | 1.8 | 0.951 | 0.951 | 97.4 | 81.2 | 89.1 | 47.1 |
| 6qm8 | Q | 3.3 | 4591 | 0.0545 | 1.07 | 276 | 3.3 | 3.3 | 0.925 | 0.925 | 94.2 | 78.5 | 88.8 | 49.4 |
| 6qm8 | R | 3.3 | 4591 | 0.0545 | 1.07 | 239 | 2.0 | 2.0 | 0.946 | 0.946 | 95.8 | 83.4 | 82.4 | 41.6 |
| 6qm8 | S | 3.3 | 4591 | 0.0545 | 1.07 | 229 | 2.4 | 2.4 | 0.932 | 0.932 | 94.8 | 70 | 88.2 | 26.7 |
| 6qm8 | T | 3.3 | 4591 | 0.0545 | 1.07 | 238 | 2.7 | 2.7 | 0.921 | 0.921 | 94.1 | 70.1 | 80.7 | 63 |
| 6qm8 | Y | 3.3 | 4591 | 0.0545 | 1.07 | 206 | 2.2 | 2.2 | 0.937 | 0.937 | 96.1 | 79.8 | 94.2 | 22.7 |
| 6qno | H | 4.4 | 4598 | 0.03 | 0.83 | 221 | 19.5 | 19.2 | 0.445 | 0.447 | 71.9 | 10.1 | 48.9 | 7.4 |

| | | | | | | | | | | | | | | |
|---|---|---|---|---|---|---|---|---|---|---|---|---|---|---|
| 6qno | L | 4.4 | 4598 | 0.03 | 0.83 | 217 | 17.6 | 17.3 | 0.366 | 0.416 | 67.3 | 8.9 | 31.8 | 4.3 |
| 6qul | P | 3 | 4638 | 0.1 | 1.079 | 117 | 2.3 | 2.3 | 0.896 | 0.896 | 94 | 69.1 | 90.6 | 51.9 |
| 6qul | W | 3 | 4638 | 0.1 | 1.079 | 94 | 3.9 | 3.9 | 0.803 | 0.803 | 87.2 | 57.3 | 85.1 | 22.5 |
| 6qum | G | 3.3 | 4640 | 0.035 | 1.085 | 207 | 7.3 | 5.0 | 0.854 | 0.869 | 87.4 | 61.9 | 76.8 | 25.8 |
| 6qum | K | 3.3 | 4640 | 0.035 | 1.085 | 99 | 5.9 | 5.9 | 0.751 | 0.751 | 83.8 | 10.8 | 68.7 | 10.3 |
| 6qum | L | 3.3 | 4640 | 0.035 | 1.085 | 187 | 6.6 | 6.6 | 0.754 | 0.754 | 84.5 | 38 | 80.2 | 44.7 |
| 6qx3 | O | 3.8 | 4661 | 0.03 | 1.08 | 121 | 15.7 | 12.8 | 0.369 | 0.401 | 78.5 | 24.2 | 33.9 | 2.4 |
| 6qxf | A | 3.6 | 4668 | 0.03 | 1.085 | 218 | 2.7 | 2.7 | 0.913 | 0.913 | 94.5 | 74.8 | 79.4 | 7.5 |
| 6qxf | B | 3.6 | 4668 | 0.03 | 1.085 | 219 | 5.7 | 5.7 | 0.812 | 0.823 | 93.2 | 63.7 | 91.8 | 43.3 |
| 6qxf | C | 3.6 | 4668 | 0.03 | 1.085 | 218 | 4.7 | 4.7 | 0.863 | 0.863 | 90.4 | 48.2 | 89.4 | 11.8 |
| 6qxf | D | 3.6 | 4668 | 0.03 | 1.085 | 219 | 2.7 | 2.7 | 0.913 | 0.913 | 96.3 | 76.3 | 96.3 | 6.6 |
| 6qxf | E | 3.6 | 4668 | 0.03 | 1.085 | 218 | 2.1 | 2.1 | 0.934 | 0.934 | 95.4 | 73.1 | 86.2 | 32.4 |
| 6qxf | F | 3.6 | 4668 | 0.03 | 1.085 | 219 | 4.7 | 3.4 | 0.901 | 0.901 | 91.8 | 66.2 | 92.2 | 50.5 |
| 6qxf | G | 3.6 | 4668 | 0.03 | 1.085 | 218 | 4.4 | 4.4 | 0.860 | 0.860 | 89.9 | 51.5 | 94 | 45.9 |
| 6qxf | H | 3.6 | 4668 | 0.03 | 1.085 | 219 | 2.1 | 2.1 | 0.917 | 0.917 | 94.1 | 74.3 | 94.1 | 18 |
| 6qxf | I | 3.6 | 4668 | 0.03 | 1.085 | 287 | 20.7 | 17.3 | 0.472 | 0.472 | 67.9 | 14.4 | 38 | 6.4 |
| 6qxf | J | 3.6 | 4668 | 0.03 | 1.085 | 287 | 22.5 | 15.2 | 0.291 | 0.518 | 72.8 | 7.7 | 48.4 | 3.6 |
| 6qxf | L | 3.6 | 4668 | 0.03 | 1.085 | 287 | 19.6 | 18.9 | 0.366 | 0.449 | 70.7 | 7.4 | 40.4 | 7.8 |
| 6qxf | M | 3.6 | 4668 | 0.03 | 1.085 | 287 | 18.8 | 17.5 | 0.416 | 0.497 | 72.8 | 14.4 | 30 | 8.1 |
| 6qxf | O | 3.6 | 4668 | 0.03 | 1.085 | 287 | 20.0 | 20.0 | 0.362 | 0.471 | 66.6 | 6.8 | 23.7 | 7.4 |
| 6qxf | P | 3.6 | 4668 | 0.03 | 1.085 | 287 | 14.8 | 13.7 | 0.415 | 0.478 | 68.6 | 8.1 | 50.5 | 4.1 |
| 6qxf | R | 3.6 | 4668 | 0.03 | 1.085 | 287 | 19.8 | 19.6 | 0.252 | 0.432 | 66.6 | 6.8 | 40.4 | 5.2 |
| 6qxf | S | 3.6 | 4668 | 0.03 | 1.085 | 287 | 20.0 | 19.4 | 0.520 | 0.522 | 72.8 | 7.7 | 55.7 | 5 |
| 6r0c | D | 4.2 | 4692 | 0.05 | 1.09 | 93 | 1.5 | 1.5 | 0.897 | 0.897 | 95.7 | 96.6 | 96.8 | 100 |
| 6r0c | H | 4.2 | 4692 | 0.05 | 1.09 | 93 | 2.5 | 2.5 | 0.879 | 0.879 | 93.5 | 75.9 | 91.4 | 56.5 |
| 6r0w | G | 3.6 | 4699 | 0.035 | 1.085 | 207 | 7.5 | 7.5 | 0.827 | 0.827 | 82.6 | 56.7 | 71 | 10.2 |
| 6r0w | H | 3.6 | 4699 | 0.035 | 1.085 | 104 | 4.8 | 4.8 | 0.674 | 0.674 | 76 | 29.1 | 47.1 | 2 |
| 6r0w | O | 3.6 | 4699 | 0.035 | 1.085 | 73 | 6.5 | 4.3 | 0.396 | 0.574 | 72.6 | 9.4 | 78.1 | 14 |
| 6r0w | P | 3.6 | 4699 | 0.035 | 1.085 | 73 | 7.0 | 4.4 | 0.400 | 0.562 | 78.1 | 5.3 | 60.3 | 40.9 |
| 6r0w | Q | 3.6 | 4699 | 0.035 | 1.085 | 74 | 8.3 | 8.3 | 0.398 | 0.555 | 75.7 | 16.1 | 60.8 | 15.6 |
| 6r0w | R | 3.6 | 4699 | 0.035 | 1.085 | 73 | 5.3 | 3.5 | 0.575 | 0.588 | 76.7 | 21.4 | 65.8 | 14.6 |
| 6r0w | S | 3.6 | 4699 | 0.035 | 1.085 | 73 | 7.3 | 5.5 | 0.420 | 0.548 | 65.8 | 6.2 | 58.9 | 14 |
| 6r0w | T | 3.6 | 4699 | 0.035 | 1.085 | 73 | 3.2 | 3.2 | 0.597 | 0.597 | 72.6 | 41.5 | 64.4 | 4.3 |
| 6r0w | U | 3.6 | 4699 | 0.035 | 1.085 | 73 | 5.0 | 5.0 | 0.582 | 0.582 | 75.3 | 23.6 | 58.9 | 14 |
| 6r0w | V | 3.6 | 4699 | 0.035 | 1.085 | 73 | 6.0 | 3.6 | 0.423 | 0.585 | 76.7 | 8.9 | 63 | 10.9 |
| 6r0w | W | 3.6 | 4699 | 0.035 | 1.085 | 73 | 6.3 | 4.7 | 0.408 | 0.627 | 76.7 | 14.3 | 72.6 | 15.1 |
| 6r0w | X | 3.6 | 4699 | 0.035 | 1.085 | 72 | 7.1 | 3.7 | 0.369 | 0.581 | 70.8 | 13.7 | 86.1 | 16.1 |

| | | | | | | | | | | | | | | |
|---|---|---|---|---|---|---|---|---|---|---|---|---|---|---|
| 6r0w | Y | 3.6 | 4699 | 0.035 | 1.085 | 73 | 6.4 | 4.7 | 0.461 | 0.551 | 65.8 | 10.4 | 64.4 | 14.9 |
| 6r0w | Z | 3.6 | 4699 | 0.035 | 1.085 | 73 | 9.3 | 3.9 | 0.419 | 0.604 | 83.6 | 18 | 63 | 15.2 |
| 6r0z | G | 3.8 | 4702 | 0.033 | 1.085 | 210 | 12.2 | 8.9 | 0.777 | 0.802 | 77.6 | 31.3 | 70.5 | 6.8 |
| 6r0z | L | 3.8 | 4702 | 0.033 | 1.085 | 186 | 11.0 | 9.6 | 0.643 | 0.643 | 77.4 | 19.4 | 68.3 | 12.6 |
| 6r0z | O | 3.8 | 4702 | 0.033 | 1.085 | 73 | 3.9 | 3.9 | 0.582 | 0.597 | 78.1 | 47.4 | 60.3 | 13.6 |
| 6r0z | P | 3.8 | 4702 | 0.033 | 1.085 | 73 | 5.7 | 5.7 | 0.548 | 0.548 | 71.2 | 19.2 | 56.2 | 14.6 |
| 6r0z | Q | 3.8 | 4702 | 0.033 | 1.085 | 74 | 10.3 | 5.8 | 0.403 | 0.595 | 85.1 | 15.9 | 63.5 | 14.9 |
| 6r0z | R | 3.8 | 4702 | 0.033 | 1.085 | 73 | 6.5 | 4.6 | 0.564 | 0.593 | 78.1 | 22.8 | 65.8 | 50 |
| 6r0z | S | 3.8 | 4702 | 0.033 | 1.085 | 73 | 5.1 | 5.1 | 0.580 | 0.580 | 82.2 | 21.7 | 58.9 | 7 |
| 6r0z | T | 3.8 | 4702 | 0.033 | 1.085 | 73 | 9.1 | 4.7 | 0.511 | 0.511 | 69.9 | 23.5 | 57.5 | 21.4 |
| 6r0z | U | 3.8 | 4702 | 0.033 | 1.085 | 73 | 4.5 | 4.5 | 0.585 | 0.585 | 83.6 | 31.1 | 64.4 | 21.3 |
| 6r0z | V | 3.8 | 4702 | 0.033 | 1.085 | 73 | 7.7 | 5.0 | 0.392 | 0.531 | 71.2 | 11.5 | 45.2 | 24.2 |
| 6r0z | W | 3.8 | 4702 | 0.033 | 1.085 | 73 | 6.9 | 4.3 | 0.387 | 0.553 | 74 | 9.3 | 60.3 | 18.2 |
| 6r0z | X | 3.8 | 4702 | 0.033 | 1.085 | 72 | 4.1 | 4.1 | 0.558 | 0.558 | 79.2 | 24.6 | 63.9 | 15.2 |
| 6r0z | Y | 3.8 | 4702 | 0.033 | 1.085 | 73 | 3.9 | 3.9 | 0.619 | 0.619 | 79.5 | 25.9 | 69.9 | 11.8 |
| 6r10 | G | 4.3 | 4703 | 0.0215 | 1.085 | 207 | 15.1 | 12.2 | 0.694 | 0.755 | 72.9 | 15.9 | 69.1 | 4.2 |
| 6r10 | I | 4.3 | 4703 | 0.0215 | 1.085 | 103 | 7.2 | 5.0 | 0.518 | 0.611 | 72.8 | 16 | 77.7 | 12.5 |
| 6r10 | J | 4.3 | 4703 | 0.0215 | 1.085 | 186 | 10.4 | 7.9 | 0.600 | 0.625 | 72.6 | 21.5 | 71 | 11.4 |
| 6r10 | L | 4.3 | 4703 | 0.0215 | 1.085 | 187 | 11.6 | 9.3 | 0.626 | 0.626 | 72.7 | 18.4 | 47.6 | 9 |
| 6r69 | G | 3.7 | 4733 | 0.018 | 0.85 | 89 | 5.4 | 4.0 | 0.857 | 0.857 | 92.1 | 25.6 | 95.5 | 51.8 |
| 6r69 | H | 3.7 | 4733 | 0.018 | 0.85 | 89 | 2.5 | 2.5 | 0.859 | 0.859 | 93.3 | 63.9 | 93.3 | 4.8 |
| 6r69 | I | 3.7 | 4733 | 0.018 | 0.85 | 89 | 2.8 | 2.8 | 0.852 | 0.852 | 93.3 | 68.7 | 87.6 | 38.5 |
| 6r69 | J | 3.7 | 4733 | 0.018 | 0.85 | 89 | 6.0 | 3.5 | 0.790 | 0.810 | 85.4 | 26.3 | 86.5 | 28.6 |
| 6r70 | A | 3.5 | 4738 | 0.025 | 1.015 | 230 | 5.5 | 5.5 | 0.875 | 0.875 | 89.6 | 60.2 | 80.9 | 40.9 |
| 6r70 | B | 3.5 | 4738 | 0.025 | 1.015 | 248 | 7.3 | 4.4 | 0.827 | 0.827 | 83.1 | 42.7 | 65.7 | 33.7 |
| 6r70 | D | 3.5 | 4738 | 0.025 | 1.015 | 233 | 20.1 | 19.4 | 0.326 | 0.356 | 65.2 | 5.9 | 65.7 | 28.1 |
| 6r70 | E | 3.5 | 4738 | 0.025 | 1.015 | 233 | 4.8 | 4.8 | 0.851 | 0.851 | 85 | 61.1 | 76 | 38.4 |
| 6r70 | G | 3.5 | 4738 | 0.025 | 1.015 | 241 | 3.5 | 3.5 | 0.870 | 0.870 | 86.7 | 57.9 | 60.2 | 44.8 |
| 6r70 | J | 3.5 | 4738 | 0.025 | 1.015 | 195 | 15.8 | 12.9 | 0.354 | 0.595 | 69.7 | 6.6 | 50.8 | 8.1 |
| 6r70 | O | 3.5 | 4738 | 0.025 | 1.015 | 230 | 20.8 | 19.8 | 0.515 | 0.549 | 83.9 | 42 | 83.9 | 34.2 |
| 6r70 | P | 3.5 | 4738 | 0.025 | 1.015 | 247 | 15.2 | 10.4 | 0.714 | 0.714 | 74.9 | 46.5 | 69.6 | 39 |
| 6r70 | R | 3.5 | 4738 | 0.025 | 1.015 | 233 | 21.2 | 16.6 | 0.309 | 0.543 | 73 | 12.9 | 60.1 | 22.1 |
| 6r70 | S | 3.5 | 4738 | 0.025 | 1.015 | 237 | 4.8 | 4.8 | 0.853 | 0.853 | 86.9 | 40.8 | 71.3 | 9.5 |
| 6r70 | U | 3.5 | 4738 | 0.025 | 1.015 | 232 | 5.1 | 5.1 | 0.837 | 0.837 | 84.1 | 51.3 | 70.3 | 17.2 |
| 6r70 | X | 3.5 | 4738 | 0.025 | 1.015 | 195 | 8.6 | 8.6 | 0.726 | 0.726 | 77.9 | 32.2 | 42.1 | 2.4 |
| 6r8f | A | 3.8 | 4759 | 0.055 | 1.0651 | 238 | 7.1 | 5.9 | 0.803 | 0.861 | 91.6 | 61 | 78.6 | 3.2 |
| 6r8f | C | 3.8 | 4759 | 0.055 | 1.0651 | 238 | 11.3 | 4.6 | 0.678 | 0.862 | 84.9 | 50.5 | 76.5 | 4.9 |

| | | | | | | | | | | | | | | |
|---|---|---|---|---|---|---|---|---|---|---|---|---|---|---|
| 6raf | C | 3.8 | 4773 | 0.57 | 1.077 | 121 | 14.7 | 13.5 | 0.317 | 0.368 | 62.8 | 2.6 | 25.6 | 6.5 |
| 6rag | C | 4.2 | 4774 | 0.5 | 1.077 | 121 | 14.8 | 14.3 | 0.240 | 0.297 | 59.5 | 6.9 | 33.1 | 7.5 |
| 6rah | C | 2.8 | 4775 | 0.03 | 1.077 | 121 | 14.2 | 6.9 | 0.547 | 0.811 | 79.3 | 11.5 | 80.2 | 3.1 |
| 6rai | C | 2.9 | 4776 | 0.03 | 1.077 | 121 | 6.3 | 6.3 | 0.821 | 0.821 | 90.1 | 74.3 | 69.4 | 7.1 |
| 6raj | C | 3.5 | 4777 | 0.6 | 1.077 | 121 | 14.6 | 11.0 | 0.274 | 0.404 | 77.7 | 6.4 | 60.3 | 5.5 |
| 6rak | C | 3.3 | 4778 | 0.6 | 1.077 | 121 | 13.6 | 12.7 | 0.378 | 0.429 | 82.6 | 6 | 57.9 | 8.6 |
| 6ral | C | 3.5 | 4779 | 0.6 | 1.077 | 121 | 14.2 | 12.3 | 0.392 | 0.671 | 78.5 | 5.3 | 38 | 6.5 |
| 6ram | C | 3.8 | 4780 | 0.5 | 1.077 | 121 | 15.8 | 14.7 | 0.253 | 0.341 | 66.9 | 8.6 | 44.6 | 13 |
| 6ran | C | 4.2 | 4781 | 0.02 | 1.077 | 121 | 14.1 | 14.1 | 0.301 | 0.307 | 62.8 | 6.6 | 22.3 | 11.1 |
| 6rbd | A | 3.5 | 4792 | 0.075 | 1.067 | 206 | 18.6 | 8.7 | 0.367 | 0.664 | 75.7 | 19.2 | 64.1 | 28 |
| 6rbd | B | 3.5 | 4792 | 0.075 | 1.067 | 214 | 19.1 | 19.1 | 0.386 | 0.404 | 68.2 | 7.5 | 51.9 | 0.9 |
| 6rbd | C | 3.5 | 4792 | 0.075 | 1.067 | 217 | 21.5 | 15.5 | 0.315 | 0.526 | 72.4 | 8.9 | 76 | 18.8 |
| 6rbd | E | 3.5 | 4792 | 0.075 | 1.067 | 260 | 8.5 | 4.3 | 0.860 | 0.860 | 86.5 | 52 | 76.2 | 6.1 |
| 6rbd | I | 3.5 | 4792 | 0.075 | 1.067 | 188 | 17.5 | 6.3 | 0.488 | 0.814 | 86.7 | 24.5 | 58.5 | 5.5 |
| 6rbd | J | 3.5 | 4792 | 0.075 | 1.067 | 185 | 4.6 | 4.0 | 0.823 | 0.845 | 86.5 | 61.2 | 65.9 | 6.6 |
| 6rbd | L | 3.5 | 4792 | 0.075 | 1.067 | 140 | 4.4 | 4.4 | 0.877 | 0.877 | 92.9 | 41.5 | 72.1 | 5.9 |
| 6rbd | W | 3.5 | 4792 | 0.075 | 1.067 | 129 | 6.5 | 6.4 | 0.781 | 0.781 | 80.6 | 46.2 | 79.8 | 5.8 |
| 6rbd | Y | 3.5 | 4792 | 0.075 | 1.067 | 134 | 11.3 | 7.9 | 0.558 | 0.763 | 83.6 | 15.2 | 72.4 | 15.5 |
| 6rd7 | 3 | 2.7 | 4808 | 0.04 | 1.053 | 245 | 3.0 | 3.0 | 0.929 | 0.929 | 94.3 | 90 | 83.7 | 76.1 |
| 6rd7 | 9 | 2.7 | 4808 | 0.04 | 1.053 | 96 | 1.8 | 1.8 | 0.915 | 0.915 | 95.8 | 85.9 | 87.5 | 20.2 |
| 6rd7 | C | 2.7 | 4808 | 0.04 | 1.053 | 74 | 1.7 | 1.7 | 0.878 | 0.878 | 94.6 | 80 | 97.3 | 100 |
| 6rd7 | F | 2.7 | 4808 | 0.04 | 1.053 | 74 | 2.4 | 2.4 | 0.885 | 0.885 | 94.6 | 57.1 | 97.3 | 100 |
| 6rd7 | I | 2.7 | 4808 | 0.04 | 1.053 | 74 | 1.3 | 1.3 | 0.918 | 0.918 | 98.6 | 93.2 | 98.6 | 98.6 |
| 6rd8 | 3 | 3.1 | 4809 | 0.04 | 1.053 | 245 | 3.2 | 3.2 | 0.929 | 0.929 | 93.1 | 81.1 | 79.6 | 83.1 |
| 6rd8 | 9 | 3.1 | 4809 | 0.04 | 1.053 | 96 | 2.9 | 2.9 | 0.886 | 0.886 | 93.8 | 86.7 | 90.6 | 64.4 |
| 6rd8 | J | 3.1 | 4809 | 0.04 | 1.053 | 74 | 1.1 | 1.1 | 0.916 | 0.916 | 97.3 | 97.2 | 100 | 97.3 |
| 6rd9 | 3 | 3 | 4810 | 0.04 | 1.053 | 245 | 2.8 | 2.8 | 0.919 | 0.919 | 93.1 | 76.8 | 77.6 | 62.6 |
| 6rd9 | 9 | 3 | 4810 | 0.04 | 1.053 | 97 | 2.6 | 2.6 | 0.863 | 0.863 | 91.8 | 80.9 | 84.5 | 53.7 |
| 6rd9 | R | 3 | 4810 | 0.04 | 1.053 | 177 | 10.4 | 10.4 | 0.734 | 0.735 | 77.4 | 22.6 | 60.5 | 30.8 |
| 6rd9 | S | 3 | 4810 | 0.04 | 1.053 | 277 | 8.2 | 3.5 | 0.816 | 0.919 | 92.4 | 66.4 | 76.9 | 65.3 |
| 6rda | 3 | 3 | 4811 | 0.04 | 1.053 | 245 | 5.0 | 5.0 | 0.890 | 0.890 | 89.8 | 81.8 | 78 | 62.8 |
| 6rda | 9 | 3 | 4811 | 0.04 | 1.053 | 97 | 2.1 | 2.1 | 0.863 | 0.863 | 91.8 | 84.3 | 86.6 | 58.3 |
| 6rdb | P | 2.8 | 4812 | 0.04 | 1.053 | 113 | 1.9 | 1.9 | 0.917 | 0.917 | 96.5 | 78.9 | 94.7 | 100 |
| 6rdb | R | 2.8 | 4812 | 0.04 | 1.053 | 177 | 15.6 | 9.2 | 0.249 | 0.680 | 68.9 | 7.4 | 49.2 | 34.5 |
| 6rdb | S | 2.8 | 4812 | 0.04 | 1.053 | 277 | 18.6 | 18.3 | 0.527 | 0.707 | 75.1 | 26.4 | 80.5 | 20.6 |
| 6rdc | 3 | 3.2 | 4813 | 0.04 | 1.053 | 245 | 2.5 | 2.5 | 0.915 | 0.915 | 92.7 | 83.7 | 76.7 | 56.4 |
| 6rdc | R | 3.2 | 4813 | 0.04 | 1.053 | 177 | 4.4 | 4.0 | 0.882 | 0.882 | 89.8 | 67.3 | 70.1 | 27.4 |

| | | | | | | | | | | | | | | |
|---|---|---|---|---|---|---|---|---|---|---|---|---|---|---|
| 6rdc | S | 3.2 | 4813 | 0.04 | 1.053 | 277 | 3.8 | 3.8 | 0.868 | 0.868 | 91.7 | 57.5 | 83.4 | 49.8 |
| 6rdd | 3 | 3.2 | 4814 | 0.04 | 1.053 | 245 | 3.5 | 3.5 | 0.914 | 0.914 | 93.5 | 85.6 | 76.3 | 38 |
| 6rde | P | 2.9 | 4815 | 0.04 | 1.053 | 113 | 1.3 | 1.3 | 0.935 | 0.935 | 99.1 | 92.9 | 91.2 | 74.8 |
| 6rde | R | 2.9 | 4815 | 0.04 | 1.053 | 177 | 5.7 | 4.1 | 0.792 | 0.810 | 81.4 | 43.1 | 51.4 | 14.3 |
| 6rde | S | 2.9 | 4815 | 0.04 | 1.053 | 277 | 2.1 | 2.1 | 0.944 | 0.944 | 95.3 | 77.7 | 84.8 | 35.3 |
| 6rdf | 3 | 3.2 | 4816 | 0.04 | 1.053 | 245 | 3.4 | 3.4 | 0.904 | 0.904 | 91.4 | 62.9 | 77.1 | 77.8 |
| 6rdg | P | 2.9 | 4817 | 0.04 | 1.053 | 113 | 2.0 | 2.0 | 0.939 | 0.939 | 98.2 | 74.8 | 93.8 | 92.5 |
| 6rdg | Q | 2.9 | 4817 | 0.04 | 1.053 | 72 | 3.4 | 3.4 | 0.766 | 0.766 | 86.1 | 50 | 72.2 | 46.2 |
| 6rdg | R | 2.9 | 4817 | 0.04 | 1.053 | 177 | 8.6 | 7.2 | 0.654 | 0.795 | 81.4 | 40.3 | 78 | 38.4 |
| 6rdg | S | 2.9 | 4817 | 0.04 | 1.053 | 277 | 3.8 | 3.8 | 0.925 | 0.925 | 93.9 | 52.3 | 84.5 | 50 |
| 6rdh | 3 | 3 | 4818 | 0.04 | 1.053 | 245 | 2.6 | 2.6 | 0.908 | 0.908 | 90.6 | 77.9 | 78.4 | 66.1 |
| 6rdh | Q | 3 | 4818 | 0.04 | 1.053 | 72 | 2.9 | 2.9 | 0.814 | 0.814 | 90.3 | 70.8 | 72.2 | 42.3 |
| 6rdh | R | 3 | 4818 | 0.04 | 1.053 | 177 | 3.6 | 3.6 | 0.855 | 0.855 | 87 | 55.8 | 66.1 | 31.6 |
| 6rdh | S | 3 | 4818 | 0.04 | 1.053 | 277 | 3.0 | 3.0 | 0.939 | 0.939 | 95.3 | 67.8 | 85.9 | 30.7 |
| 6rdi | 3 | 3.2 | 4819 | 0.04 | 1.053 | 245 | 2.7 | 2.7 | 0.910 | 0.910 | 91.8 | 73.8 | 73.1 | 43 |
| 6rdi | R | 3.2 | 4819 | 0.04 | 1.053 | 177 | 6.0 | 6.0 | 0.813 | 0.813 | 83.1 | 47.6 | 66.1 | 14.5 |
| 6rdi | S | 3.2 | 4819 | 0.04 | 1.053 | 277 | 3.3 | 3.3 | 0.922 | 0.922 | 93.5 | 59.8 | 83 | 14.8 |
| 6rdj | P | 2.9 | 4820 | 0.04 | 1.053 | 114 | 1.2 | 1.2 | 0.938 | 0.938 | 97.4 | 96.4 | 91.2 | 78.8 |
| 6rdj | R | 2.9 | 4820 | 0.04 | 1.053 | 177 | 6.1 | 6.1 | 0.816 | 0.816 | 83.6 | 38.5 | 73.4 | 21.5 |
| 6rdj | S | 2.9 | 4820 | 0.04 | 1.053 | 277 | 1.9 | 1.9 | 0.952 | 0.952 | 96.4 | 81.6 | 86.3 | 36 |
| 6rdk | 3 | 3.7 | 4821 | 0.04 | 1.053 | 245 | 3.4 | 3.4 | 0.873 | 0.873 | 87.3 | 64 | 71.8 | 51.1 |
| 6rdk | S | 3.7 | 4821 | 0.04 | 1.053 | 277 | 8.7 | 8.6 | 0.785 | 0.787 | 88.8 | 54.9 | 79.4 | 20.9 |
| 6rdl | 3 | 3.7 | 4822 | 0.04 | 1.053 | 245 | 7.6 | 4.2 | 0.769 | 0.879 | 86.5 | 52.4 | 76.3 | 53.5 |
| 6rdl | S | 3.7 | 4822 | 0.04 | 1.053 | 277 | 16.4 | 16.4 | 0.688 | 0.688 | 74 | 17.1 | 70.8 | 8.7 |
| 6rdm | P | 3.4 | 4823 | 0.04 | 1.053 | 114 | 3.3 | 3.3 | 0.891 | 0.891 | 93.9 | 58.9 | 89.5 | 5.9 |
| 6rdm | S | 3.4 | 4823 | 0.04 | 1.053 | 277 | 11.9 | 9.9 | 0.677 | 0.683 | 80.9 | 27.2 | 61.7 | 9.9 |
| 6rdn | 3 | 3.2 | 4824 | 0.04 | 1.053 | 245 | 5.2 | 5.2 | 0.868 | 0.868 | 86.9 | 60.6 | 79.2 | 57.2 |
| 6rdn | 9 | 3.2 | 4824 | 0.04 | 1.053 | 97 | 3.3 | 3.3 | 0.847 | 0.847 | 89.7 | 65.5 | 84.5 | 40.2 |
| 6rdn | S | 3.2 | 4824 | 0.04 | 1.053 | 277 | 9.3 | 8.6 | 0.764 | 0.821 | 85.9 | 57.1 | 61.7 | 42.1 |
| 6rdo | 3 | 3.1 | 4825 | 0.04 | 1.053 | 245 | 2.8 | 2.8 | 0.915 | 0.915 | 92.2 | 70.8 | 79.6 | 53.8 |
| 6rdo | 9 | 3.1 | 4825 | 0.04 | 1.053 | 97 | 2.6 | 2.6 | 0.855 | 0.855 | 91.8 | 69.7 | 82.5 | 55 |
| 6rdo | S | 3.1 | 4825 | 0.04 | 1.053 | 277 | 9.7 | 9.7 | 0.758 | 0.767 | 85.6 | 48.5 | 70.8 | 56.6 |
| 6rdp | P | 2.8 | 4826 | 0.04 | 1.053 | 114 | 1.5 | 1.5 | 0.937 | 0.937 | 98.2 | 87.5 | 95.6 | 89.9 |
| 6rdp | S | 2.8 | 4826 | 0.04 | 1.053 | 277 | 10.0 | 10.0 | 0.715 | 0.715 | 77.3 | 44.4 | 79.1 | 47 |
| 6rdq | 3 | 4 | 4827 | 0.04 | 1.053 | 245 | 6.0 | 5.4 | 0.832 | 0.832 | 83.7 | 24.9 | 79.2 | 38.7 |
| 6rdr | 3 | 4.1 | 4828 | 0.04 | 1.053 | 245 | 12.1 | 10.8 | 0.776 | 0.776 | 79.2 | 20.1 | 78.4 | 5.7 |
| 6rds | P | 3.8 | 4829 | 0.04 | 1.053 | 114 | 2.8 | 2.8 | 0.853 | 0.853 | 90.4 | 66 | 93.9 | 7.5 |

| | | | | | | | | | | | | | | |
|---|---|---|---|---|---|---|---|---|---|---|---|---|---|---|
| 6rdt | 3 | 3.4 | 4830 | 0.04 | 1.053 | 245 | 2.9 | 2.9 | 0.890 | 0.890 | 90.2 | 63.3 | 76.7 | 61.7 |
| 6rdt | S | 3.4 | 4830 | 0.04 | 1.053 | 277 | 11.9 | 11.8 | 0.746 | 0.746 | 81.2 | 33.8 | 70.8 | 44.9 |
| 6rdu | 3 | 3.5 | 4831 | 0.025 | 1.053 | 245 | 3.4 | 3.4 | 0.890 | 0.890 | 89 | 59.2 | 77.6 | 55.8 |
| 6rdv | P | 3.1 | 4832 | 0.025 | 1.053 | 114 | 2.4 | 2.4 | 0.922 | 0.922 | 97.4 | 71.2 | 90.4 | 47.6 |
| 6rdv | S | 3.1 | 4832 | 0.025 | 1.053 | 277 | 12.5 | 10.4 | 0.662 | 0.748 | 82.3 | 28.9 | 68.2 | 35.4 |
| 6rdw | 3 | 3.8 | 4833 | 0.04 | 1.053 | 245 | 5.7 | 3.6 | 0.824 | 0.872 | 83.7 | 49.3 | 74.3 | 47.3 |
| 6rdw | S | 3.8 | 4833 | 0.04 | 1.053 | 277 | 4.9 | 4.9 | 0.875 | 0.875 | 93.5 | 43.2 | 84.8 | 24.7 |
| 6rdx | 3 | 3.9 | 4834 | 0.04 | 1.053 | 245 | 3.2 | 3.2 | 0.875 | 0.875 | 87.8 | 55.8 | 76.3 | 26.2 |
| 6rdx | S | 3.9 | 4834 | 0.04 | 1.053 | 277 | 8.5 | 8.4 | 0.850 | 0.850 | 86.3 | 21.3 | 71.1 | 26.4 |
| 6rdy | P | 3.6 | 4835 | 0.04 | 1.053 | 114 | 3.6 | 3.6 | 0.851 | 0.859 | 90.4 | 49.5 | 88.6 | 46.5 |
| 6rdy | S | 3.6 | 4835 | 0.04 | 1.053 | 277 | 23.6 | 18.3 | 0.386 | 0.510 | 77.3 | 10.7 | 83 | 27 |
| 6rdz | 3 | 3.5 | 4836 | 0.04 | 1.053 | 245 | 3.2 | 3.2 | 0.884 | 0.884 | 87.3 | 79.4 | 76.3 | 5.3 |
| 6rdz | S | 3.5 | 4836 | 0.04 | 1.053 | 277 | 12.7 | 10.8 | 0.685 | 0.721 | 81.2 | 36 | 62.1 | 30.8 |
| 6re0 | 3 | 3.6 | 4837 | 0.04 | 1.053 | 245 | 3.1 | 3.1 | 0.893 | 0.893 | 92.2 | 70.4 | 79.2 | 44.3 |
| 6re1 | P | 3.2 | 4838 | 0.04 | 1.053 | 114 | 1.9 | 1.9 | 0.887 | 0.887 | 94.7 | 81.5 | 92.1 | 73.3 |
| 6re1 | S | 3.2 | 4838 | 0.04 | 1.053 | 277 | 12.1 | 12.0 | 0.647 | 0.690 | 75.8 | 31.4 | 61.4 | 40 |
| 6re2 | 3 | 3.2 | 4839 | 0.04 | 1.053 | 245 | 5.6 | 4.2 | 0.801 | 0.872 | 91.4 | 61.2 | 73.9 | 53 |
| 6re2 | R | 3.2 | 4839 | 0.04 | 1.053 | 177 | 2.1 | 2.1 | 0.907 | 0.907 | 93.8 | 72.3 | 83.6 | 8.8 |
| 6re2 | S | 3.2 | 4839 | 0.04 | 1.053 | 277 | 2.1 | 2.1 | 0.940 | 0.940 | 94.9 | 82.1 | 84.5 | 54.3 |
| 6re3 | 3 | 3.3 | 4840 | 0.04 | 1.053 | 245 | 7.6 | 4.2 | 0.866 | 0.891 | 88.2 | 69.9 | 75.1 | 38 |
| 6re3 | 9 | 3.3 | 4840 | 0.04 | 1.053 | 97 | 3.9 | 3.9 | 0.802 | 0.802 | 86.6 | 40.5 | 82.5 | 38.8 |
| 6re3 | R | 3.3 | 4840 | 0.04 | 1.053 | 177 | 3.8 | 3.8 | 0.881 | 0.881 | 91 | 70.8 | 65 | 10.4 |
| 6re3 | S | 3.3 | 4840 | 0.04 | 1.053 | 277 | 2.6 | 2.6 | 0.949 | 0.949 | 96.4 | 72.3 | 85.6 | 46.8 |
| 6re4 | P | 3 | 4841 | 0.04 | 1.053 | 114 | 1.9 | 1.9 | 0.911 | 0.911 | 94.7 | 71.3 | 96.5 | 99.1 |
| 6re4 | Q | 3 | 4841 | 0.04 | 1.053 | 72 | 2.3 | 2.3 | 0.815 | 0.815 | 90.3 | 64.6 | 75 | 7.4 |
| 6re4 | R | 3 | 4841 | 0.04 | 1.053 | 177 | 3.3 | 3.3 | 0.869 | 0.869 | 90.4 | 60 | 78 | 27.5 |
| 6re4 | S | 3 | 4841 | 0.04 | 1.053 | 277 | 2.8 | 2.8 | 0.931 | 0.931 | 94.6 | 71.8 | 84.5 | 44.4 |
| 6re5 | 3 | 3.2 | 4842 | 0.04 | 1.053 | 245 | 4.0 | 3.6 | 0.891 | 0.906 | 90.2 | 64.3 | 72.2 | 36.2 |
| 6re5 | Q | 3.2 | 4842 | 0.04 | 1.053 | 72 | 4.7 | 4.7 | 0.708 | 0.708 | 79.2 | 47.4 | 72.2 | 46.2 |
| 6re5 | R | 3.2 | 4842 | 0.04 | 1.053 | 177 | 16.8 | 14.9 | 0.705 | 0.716 | 72.3 | 16.4 | 80.8 | 9.8 |
| 6re5 | S | 3.2 | 4842 | 0.04 | 1.053 | 277 | 3.3 | 3.3 | 0.935 | 0.935 | 95.3 | 80.3 | 88.1 | 28.3 |
| 6re6 | 3 | 3.4 | 4843 | 0.04 | 1.053 | 245 | 5.7 | 4.0 | 0.837 | 0.890 | 90.6 | 60.8 | 78 | 73.3 |
| 6re6 | S | 3.4 | 4843 | 0.04 | 1.053 | 277 | 5.5 | 3.3 | 0.901 | 0.901 | 91.3 | 46.6 | 85.2 | 49.2 |
| 6re7 | P | 3.1 | 4844 | 0.04 | 1.053 | 114 | 2.0 | 2.0 | 0.921 | 0.921 | 96.5 | 82.7 | 93 | 67 |
| 6re7 | S | 3.1 | 4844 | 0.04 | 1.053 | 277 | 3.1 | 3.1 | 0.916 | 0.930 | 92.8 | 59.1 | 84.1 | 54.5 |
| 6re8 | 3 | 3.8 | 4845 | 0.04 | 1.053 | 245 | 5.9 | 3.6 | 0.768 | 0.868 | 86.1 | 28.4 | 48.2 | 6.8 |
| 6re8 | S | 3.8 | 4845 | 0.04 | 1.053 | 277 | 7.5 | 7.5 | 0.825 | 0.831 | 84.5 | 38 | 75.5 | 28.2 |

| | | | | | | | | | | | | | | |
|---|---|---|---|---|---|---|---|---|---|---|---|---|---|---|
| 6re9 | 3 | 3.9 | 4846 | 0.04 | 1.053 | 245 | 4.2 | 4.2 | 0.863 | 0.863 | 85.7 | 41 | 75.1 | 7.1 |
| 6rea | P | 3.6 | 4847 | 0.04 | 1.053 | 114 | 2.7 | 2.7 | 0.879 | 0.879 | 93 | 57.5 | 90.4 | 43.7 |
| 6rea | S | 3.6 | 4847 | 0.04 | 1.053 | 277 | 11.4 | 7.8 | 0.756 | 0.803 | 83 | 20.9 | 74.7 | 26.6 |
| 6reb | 3 | 3.2 | 4848 | 0.04 | 1.053 | 245 | 3.2 | 3.2 | 0.899 | 0.899 | 91.8 | 64.9 | 76.7 | 60.1 |
| 6reb | R | 3.2 | 4848 | 0.04 | 1.053 | 177 | 3.1 | 3.1 | 0.877 | 0.877 | 91.5 | 71.6 | 74.6 | 34.1 |
| 6reb | S | 3.2 | 4848 | 0.04 | 1.053 | 277 | 2.4 | 2.4 | 0.938 | 0.938 | 94.9 | 77.2 | 84.1 | 50.6 |
| 6rec | 3 | 3.3 | 4849 | 0.04 | 1.053 | 245 | 2.9 | 2.9 | 0.900 | 0.900 | 89.8 | 65.9 | 76.3 | 51.9 |
| 6rec | R | 3.3 | 4849 | 0.04 | 1.053 | 177 | 5.3 | 5.3 | 0.813 | 0.818 | 83.6 | 54.1 | 70.6 | 42.4 |
| 6rec | S | 3.3 | 4849 | 0.04 | 1.053 | 277 | 3.7 | 3.7 | 0.903 | 0.903 | 91 | 46.4 | 83 | 26.5 |
| 6ree | 3 | 3.1 | 4851 | 0.04 | 1.053 | 245 | 4.0 | 4.0 | 0.898 | 0.898 | 92.2 | 75.2 | 80 | 49.5 |
| 6ree | Q | 3.1 | 4851 | 0.04 | 1.053 | 72 | 3.3 | 3.3 | 0.782 | 0.782 | 86.1 | 53.2 | 70.8 | 9.8 |
| 6ree | R | 3.1 | 4851 | 0.04 | 1.053 | 177 | 8.2 | 3.4 | 0.825 | 0.872 | 88.1 | 44.9 | 75.7 | 24.6 |
| 6ree | S | 3.1 | 4851 | 0.04 | 1.053 | 277 | 8.3 | 3.8 | 0.831 | 0.933 | 94.6 | 48.9 | 87 | 73 |
| 6ref | 3 | 3.3 | 4852 | 0.04 | 1.053 | 245 | 2.4 | 2.4 | 0.917 | 0.917 | 92.2 | 77.4 | 78.8 | 74.6 |
| 6ref | R | 3.3 | 4852 | 0.04 | 1.053 | 177 | 3.6 | 3.6 | 0.853 | 0.854 | 87 | 53.9 | 60.5 | 21.5 |
| 6ref | S | 3.3 | 4852 | 0.04 | 1.053 | 277 | 9.3 | 5.9 | 0.826 | 0.879 | 92.4 | 64.1 | 83.8 | 48.7 |
| 6rep | 3 | 3.1 | 4853 | 0.04 | 1.053 | 245 | 2.3 | 2.3 | 0.928 | 0.928 | 93.9 | 82.6 | 70.2 | 64.5 |
| 6rep | Q | 3.1 | 4853 | 0.04 | 1.053 | 72 | 3.1 | 3.1 | 0.807 | 0.807 | 87.5 | 76.2 | 69.4 | 10 |
| 6rep | R | 3.1 | 4853 | 0.04 | 1.053 | 177 | 2.5 | 2.5 | 0.887 | 0.887 | 91.5 | 71.6 | 57.6 | 24.5 |
| 6rep | S | 3.1 | 4853 | 0.04 | 1.053 | 277 | 2.6 | 2.6 | 0.929 | 0.929 | 93.5 | 63.3 | 87 | 46.9 |
| 6rer | P | 2.9 | 4854 | 0.04 | 1.053 | 114 | 2.1 | 2.1 | 0.921 | 0.921 | 98.2 | 85.7 | 96.5 | 91.8 |
| 6rer | S | 2.9 | 4854 | 0.04 | 1.053 | 277 | 2.9 | 2.9 | 0.942 | 0.942 | 96 | 60.2 | 85.6 | 24.1 |
| 6res | 3 | 4.3 | 4855 | 0.04 | 1.053 | 245 | 6.0 | 5.5 | 0.806 | 0.806 | 82.4 | 40.1 | 72.7 | 6.2 |
| 6res | S | 4.3 | 4855 | 0.04 | 1.053 | 277 | 14.6 | 14.2 | 0.595 | 0.652 | 72.2 | 21.5 | 66.4 | 7.6 |
| 6ret | 3 | 4.3 | 4856 | 0.04 | 1.053 | 245 | 10.7 | 6.4 | 0.701 | 0.805 | 77.6 | 21.1 | 70.6 | 13.3 |
| 6rey | A | 3 | 4860 | 5.7 | 0.81 | 234 | 2.0 | 2.0 | 0.949 | 0.949 | 97.4 | 74.6 | 91.5 | 46.7 |
| 6rey | B | 3 | 4860 | 5.7 | 0.81 | 214 | 3.1 | 3.1 | 0.913 | 0.913 | 92.1 | 69.5 | 92.5 | 62.6 |
| 6rey | C | 3 | 4860 | 5.7 | 0.81 | 207 | 9.5 | 9.5 | 0.771 | 0.771 | 87.4 | 47 | 85 | 27.8 |
| 6rey | D | 3 | 4860 | 5.7 | 0.81 | 210 | 2.1 | 2.1 | 0.930 | 0.930 | 95.2 | 76.5 | 89.5 | 27.1 |
| 6rey | E | 3 | 4860 | 5.7 | 0.81 | 227 | 1.9 | 1.9 | 0.942 | 0.942 | 95.6 | 77.9 | 85 | 33.7 |
| 6rey | F | 3 | 4860 | 5.7 | 0.81 | 236 | 1.6 | 1.6 | 0.956 | 0.956 | 98.3 | 87.5 | 88.1 | 42.8 |
| 6rey | K | 3 | 4860 | 5.7 | 0.81 | 196 | 1.2 | 1.2 | 0.956 | 0.956 | 98 | 96.4 | 92.9 | 80.2 |
| 6rey | O | 3 | 4860 | 5.7 | 0.81 | 234 | 2.5 | 2.5 | 0.927 | 0.927 | 93.6 | 66.7 | 91.9 | 72.6 |
| 6rey | P | 3 | 4860 | 5.7 | 0.81 | 214 | 2.8 | 2.8 | 0.941 | 0.941 | 95.3 | 88.2 | 86 | 45.7 |
| 6rey | Q | 3 | 4860 | 5.7 | 0.81 | 207 | 8.1 | 8.1 | 0.850 | 0.850 | 87.9 | 48.9 | 88.9 | 19.6 |
| 6rey | R | 3 | 4860 | 5.7 | 0.81 | 210 | 3.7 | 3.7 | 0.898 | 0.898 | 91.4 | 56.2 | 91.9 | 31.1 |
| 6rey | S | 3 | 4860 | 5.7 | 0.81 | 227 | 2.2 | 2.2 | 0.940 | 0.940 | 93.8 | 79.8 | 85.9 | 28.2 |

| | | | | | | | | | | | | | | |
|---|---|---|---|---|---|---|---|---|---|---|---|---|---|---|
| 6rey | T | 3 | 4860 | 5.7 | 0.81 | 236 | 2.3 | 2.3 | 0.932 | 0.932 | 94.1 | 78.4 | 91.5 | 33.3 |
| 6rey | Y | 3 | 4860 | 5.7 | 0.81 | 196 | 1.7 | 1.7 | 0.955 | 0.955 | 97.4 | 82.7 | 93.4 | 68.3 |
| 6rfl | E | 2.8 | 4868 | 0.05 | 1.0635 | 184 | 1.7 | 1.7 | 0.946 | 0.946 | 96.2 | 84.2 | 91.3 | 56 |
| 6rfl | G | 2.8 | 4868 | 0.05 | 1.0635 | 153 | 1.5 | 1.5 | 0.951 | 0.951 | 98 | 86.7 | 82.4 | 11.9 |
| 6rfl | Q | 2.8 | 4868 | 0.05 | 1.0635 | 124 | 2.2 | 2.2 | 0.910 | 0.910 | 96 | 72.3 | 82.3 | 31.4 |
| 6rfl | R | 2.8 | 4868 | 0.05 | 1.0635 | 129 | 2.0 | 2.0 | 0.909 | 0.909 | 94.6 | 80.3 | 83.7 | 20.4 |
| 6rfq | b | 3.3 | 4872 | 0.021 | 1.077 | 64 | 1.5 | 1.5 | 0.897 | 0.897 | 96.9 | 88.7 | 98.4 | 100 |
| 6rfq | E | 3.3 | 4872 | 0.021 | 1.077 | 263 | 14.2 | 12.4 | 0.731 | 0.811 | 83.7 | 43.6 | 75.7 | 17.1 |
| 6rfq | f | 3.3 | 4872 | 0.021 | 1.077 | 79 | 4.2 | 4.2 | 0.748 | 0.748 | 87.3 | 52.2 | 84.8 | 7.5 |
| 6rfq | P | 3.3 | 4872 | 0.021 | 1.077 | 123 | 3.3 | 3.3 | 0.886 | 0.886 | 92.7 | 75.4 | 87 | 49.5 |
| 6rfq | Q | 3.3 | 4872 | 0.021 | 1.077 | 85 | 5.5 | 3.4 | 0.861 | 0.861 | 92.9 | 41.8 | 84.7 | 11.1 |
| 6rfq | X | 3.3 | 4872 | 0.021 | 1.077 | 168 | 1.1 | 1.1 | 0.957 | 0.957 | 98.2 | 97 | 85.1 | 53.1 |
| 6rfs | b | 4 | 4874 | 0.014 | 1.09 | 64 | 2.8 | 2.8 | 0.798 | 0.798 | 87.5 | 64.3 | 96.9 | 8.1 |
| 6rfs | f | 4 | 4874 | 0.014 | 1.09 | 80 | 13.6 | 8.2 | 0.394 | 0.611 | 83.8 | 11.9 | 43.8 | 11.4 |
| 6rfs | F | 4 | 4874 | 0.014 | 1.09 | 121 | 4.1 | 4.1 | 0.800 | 0.800 | 85.1 | 25.2 | 90.1 | 9.2 |
| 6rfs | H | 4 | 4874 | 0.014 | 1.09 | 213 | 10.0 | 8.6 | 0.664 | 0.699 | 80.3 | 31.6 | 42.7 | 5.5 |
| 6rfs | M | 4 | 4874 | 0.014 | 1.09 | 117 | 12.8 | 11.3 | 0.342 | 0.696 | 79.5 | 12.9 | 70.9 | 7.2 |
| 6rfs | O | 4 | 4874 | 0.014 | 1.09 | 77 | 10.4 | 5.1 | 0.294 | 0.493 | 64.9 | 10 | 72.7 | 8.9 |
| 6rfs | P | 4 | 4874 | 0.014 | 1.09 | 123 | 5.2 | 5.2 | 0.777 | 0.777 | 87 | 32.7 | 76.4 | 4.3 |
| 6rfs | Q | 4 | 4874 | 0.014 | 1.09 | 85 | 8.6 | 4.4 | 0.707 | 0.744 | 76.5 | 10.8 | 55.3 | 8.5 |
| 6rfs | U | 4 | 4874 | 0.014 | 1.09 | 171 | 7.3 | 3.8 | 0.833 | 0.834 | 87.7 | 30.7 | 67.3 | 7.8 |
| 6rfs | X | 4 | 4874 | 0.014 | 1.09 | 167 | 3.0 | 3.0 | 0.886 | 0.886 | 91 | 53.3 | 71.3 | 10.9 |
| 6rgq | A | 2.6 | 4877 | 2.6 | 0.81 | 240 | 1.5 | 1.5 | 0.958 | 0.958 | 96.7 | 89.7 | 86.2 | 71.5 |
| 6rgq | B | 2.6 | 4877 | 2.6 | 0.81 | 229 | 1.0 | 1.0 | 0.972 | 0.972 | 98.7 | 97.8 | 92.6 | 60.8 |
| 6rgq | C | 2.6 | 4877 | 2.6 | 0.81 | 247 | 2.0 | 2.0 | 0.939 | 0.939 | 94.7 | 77.4 | 83.8 | 66.7 |
| 6rgq | D | 2.6 | 4877 | 2.6 | 0.81 | 232 | 2.3 | 2.3 | 0.928 | 0.928 | 94.8 | 71.8 | 85.8 | 50.8 |
| 6rgq | E | 2.6 | 4877 | 2.6 | 0.81 | 233 | 3.1 | 3.1 | 0.931 | 0.934 | 94 | 65.3 | 85.4 | 39.2 |
| 6rgq | F | 2.6 | 4877 | 2.6 | 0.81 | 233 | 1.8 | 1.8 | 0.943 | 0.943 | 96.1 | 80.4 | 90.1 | 82.4 |
| 6rgq | K | 2.6 | 4877 | 2.6 | 0.81 | 196 | 1.3 | 1.3 | 0.963 | 0.963 | 98.5 | 93.8 | 94.9 | 84.9 |
| 6rgq | O | 2.6 | 4877 | 2.6 | 0.81 | 240 | 2.1 | 2.1 | 0.951 | 0.951 | 96.2 | 83.1 | 90 | 67.6 |
| 6rgq | P | 2.6 | 4877 | 2.6 | 0.81 | 229 | 1.3 | 1.3 | 0.962 | 0.962 | 98.3 | 89.8 | 91.7 | 69.5 |
| 6rgq | Q | 2.6 | 4877 | 2.6 | 0.81 | 247 | 2.7 | 2.7 | 0.925 | 0.925 | 93.9 | 60.8 | 87.4 | 55.1 |
| 6rgq | R | 2.6 | 4877 | 2.6 | 0.81 | 232 | 2.1 | 2.1 | 0.934 | 0.934 | 95.3 | 85.5 | 82.3 | 61.3 |
| 6rgq | S | 2.6 | 4877 | 2.6 | 0.81 | 233 | 2.0 | 2.0 | 0.956 | 0.956 | 97 | 77 | 82.8 | 21.8 |
| 6rgq | T | 2.6 | 4877 | 2.6 | 0.81 | 233 | 3.1 | 3.1 | 0.953 | 0.953 | 97 | 69.9 | 89.3 | 48.6 |
| 6rgq | Y | 2.6 | 4877 | 2.6 | 0.81 | 196 | 1.6 | 1.6 | 0.953 | 0.953 | 96.9 | 84.7 | 94.4 | 82.7 |
| 6rh3 | A | 3.6 | 4882 | 0.35 | 1.042 | 228 | 2.4 | 2.4 | 0.937 | 0.937 | 96.1 | 79 | 92.1 | 25.2 |

| | | | | | | | | | | | | | | |
|---|---|---|---|---|---|---|---|---|---|---|---|---|---|---|
| 6rh3 | B | 3.6 | 4882 | 0.35 | 1.042 | 229 | 3.1 | 3.1 | 0.887 | 0.887 | 89.5 | 60.5 | 76.9 | 17.6 |
| 6rh3 | E | 3.6 | 4882 | 0.35 | 1.042 | 73 | 2.2 | 2.2 | 0.893 | 0.893 | 95.9 | 78.6 | 93.2 | 41.2 |
| 6ri7 | A | 3.9 | 4885 | 0.5 | 1.039 | 228 | 3.0 | 3.0 | 0.907 | 0.907 | 93 | 71.2 | 86.4 | 7.6 |
| 6ri7 | B | 3.9 | 4885 | 0.5 | 1.039 | 229 | 14.5 | 13.8 | 0.682 | 0.706 | 76.4 | 26.9 | 66.4 | 5.9 |
| 6ri7 | E | 3.9 | 4885 | 0.5 | 1.039 | 73 | 2.4 | 2.4 | 0.824 | 0.824 | 91.8 | 65.7 | 53.4 | 7.7 |
| 6ri7 | F | 3.9 | 4885 | 0.5 | 1.039 | 156 | 9.4 | 7.9 | 0.698 | 0.698 | 78.8 | 24.4 | 76.3 | 4.2 |
| 6ri9 | A | 3.7 | 4886 | 0.5 | 1.067 | 228 | 3.6 | 3.6 | 0.922 | 0.922 | 94.3 | 54.4 | 87.7 | 21 |
| 6ri9 | B | 3.7 | 4886 | 0.5 | 1.067 | 229 | 13.3 | 12.6 | 0.618 | 0.702 | 83.4 | 37.7 | 39.3 | 12.2 |
| 6ri9 | E | 3.7 | 4886 | 0.5 | 1.067 | 73 | 3.4 | 3.4 | 0.792 | 0.792 | 89 | 40 | 86.3 | 4.8 |
| 6ric | E | 2.8 | 4888 | 0.0275 | 0.81 | 184 | 1.7 | 1.7 | 0.936 | 0.936 | 96.2 | 88.7 | 89.7 | 64.2 |
| 6ric | F | 2.8 | 4888 | 0.0275 | 0.81 | 103 | 1.4 | 1.4 | 0.925 | 0.925 | 97.1 | 90 | 92.2 | 87.4 |
| 6ric | G | 2.8 | 4888 | 0.0275 | 0.81 | 131 | 7.6 | 7.6 | 0.749 | 0.749 | 84 | 57.3 | 75.6 | 35.4 |
| 6rid | E | 2.9 | 4889 | 0.028 | 1.05 | 184 | 2.5 | 2.5 | 0.938 | 0.938 | 96.2 | 82.5 | 90.2 | 7.2 |
| 6rid | F | 2.9 | 4889 | 0.028 | 1.05 | 103 | 2.0 | 2.0 | 0.926 | 0.926 | 96.1 | 85.9 | 91.3 | 73.4 |
| 6rie | E | 3.1 | 4890 | 0.028 | 1.05 | 184 | 3.4 | 3.4 | 0.927 | 0.927 | 95.1 | 81.1 | 90.2 | 39.8 |
| 6rie | F | 3.1 | 4890 | 0.028 | 1.05 | 103 | 1.4 | 1.4 | 0.933 | 0.933 | 96.1 | 94.9 | 91.3 | 34 |
| 6rie | G | 3.1 | 4890 | 0.028 | 1.05 | 131 | 11.9 | 7.2 | 0.642 | 0.778 | 80.9 | 49.1 | 45 | 8.5 |
| 6rie | L | 3.1 | 4890 | 0.028 | 1.05 | 287 | 22.7 | 21.8 | 0.422 | 0.527 | 84.3 | 24.8 | 69 | 16.7 |
| 6rin | A | 3.7 | 4892 | 0.4 | 1.1 | 228 | 2.4 | 2.4 | 0.913 | 0.913 | 93.9 | 62.6 | 68 | 9 |
| 6rin | B | 3.7 | 4892 | 0.4 | 1.1 | 229 | 14.6 | 12.6 | 0.579 | 0.586 | 77.7 | 28.1 | 44.5 | 6.9 |
| 6rin | E | 3.7 | 4892 | 0.4 | 1.1 | 73 | 1.1 | 1.1 | 0.898 | 0.898 | 98.6 | 94.4 | 93.2 | 45.6 |
| 6rin | F | 3.7 | 4892 | 0.4 | 1.1 | 156 | 3.2 | 3.2 | 0.873 | 0.873 | 92.9 | 71 | 82.1 | 8.6 |
| 6rip | A | 3.4 | 4893 | 0.6 | 1.067 | 228 | 2.3 | 2.3 | 0.951 | 0.951 | 96.9 | 83.3 | 90.4 | 42.2 |
| 6rip | B | 3.4 | 4893 | 0.6 | 1.067 | 229 | 11.7 | 9.4 | 0.746 | 0.753 | 87.8 | 39.8 | 69.4 | 50.3 |
| 6rip | E | 3.4 | 4893 | 0.6 | 1.067 | 73 | 2.0 | 2.0 | 0.892 | 0.892 | 95.9 | 84.3 | 87.7 | 50 |
| 6rj9 | A | 3.2 | 4900 | 0.05 | 0.889 | 183 | 2.0 | 2.0 | 0.942 | 0.942 | 96.7 | 84.2 | 90.7 | 49.4 |
| 6rj9 | B | 3.2 | 4900 | 0.05 | 0.889 | 183 | 5.5 | 3.2 | 0.818 | 0.875 | 89.6 | 72.6 | 73.2 | 39.6 |
| 6rja | A | 3 | 4901 | 0.0625 | 0.889 | 183 | 1.9 | 1.9 | 0.946 | 0.946 | 97.3 | 86 | 91.3 | 74.3 |
| 6rja | B | 3 | 4901 | 0.0625 | 0.889 | 183 | 2.8 | 2.8 | 0.908 | 0.908 | 91.8 | 65.5 | 91.8 | 60.7 |
| 6rjg | A | 3.2 | 4904 | 0.05 | 0.889 | 183 | 2.3 | 2.3 | 0.915 | 0.915 | 94.5 | 70.5 | 84.7 | 40.6 |
| 6rks | B | 4 | 4909 | 0.0263 | 0.88 | 200 | 9.2 | 9.2 | 0.761 | 0.761 | 78.5 | 45.2 | 71.5 | 18.2 |
| 6rks | D | 4 | 4909 | 0.0263 | 0.88 | 200 | 9.6 | 8.7 | 0.669 | 0.716 | 75 | 26 | 71 | 7.7 |
| 6rlb | G | 4.5 | 4918 | 0.0318 | 1.39 | 93 | 13.4 | 11.4 | 0.345 | 0.449 | 73.1 | 14.7 | 47.3 | 9.1 |
| 6rlb | H | 4.5 | 4918 | 0.0318 | 1.39 | 93 | 14.3 | 8.8 | 0.255 | 0.567 | 68.8 | 6.2 | 53.8 | 8 |
| 6rld | A | 2.9 | 4919 | 0.0259 | 1.0635 | 265 | 2.2 | 2.2 | 0.946 | 0.946 | 96.6 | 82.8 | 82.3 | 62.8 |
| 6rld | B | 2.9 | 4919 | 0.0259 | 1.0635 | 265 | 1.6 | 1.6 | 0.956 | 0.956 | 97 | 86.8 | 84.2 | 78.5 |
| 6rld | C | 2.9 | 4919 | 0.0259 | 1.0635 | 265 | 2.0 | 2.0 | 0.954 | 0.954 | 96.6 | 83.2 | 82.3 | 49.5 |

| | | | | | | | | | | | | | | |
|---|---|---|---|---|---|---|---|---|---|---|---|---|---|---|
| 6rld | D | 2.9 | 4919 | 0.0259 | 1.0635 | 265 | 2.1 | 2.1 | 0.941 | 0.941 | 95.8 | 71.7 | 84.2 | 74 |
| 6rld | E | 2.9 | 4919 | 0.0259 | 1.0635 | 265 | 2.5 | 2.5 | 0.936 | 0.936 | 94.7 | 67.3 | 82.3 | 54.6 |
| 6rld | F | 2.9 | 4919 | 0.0259 | 1.0635 | 265 | 1.9 | 1.9 | 0.946 | 0.946 | 95.5 | 78.3 | 84.2 | 78.5 |
| 6rld | G | 2.9 | 4919 | 0.0259 | 1.0635 | 265 | 2.3 | 2.3 | 0.940 | 0.940 | 95.5 | 79.8 | 82.3 | 49.1 |
| 6rmg | B | 3.4 | 4936 | 0.016 | 0.8141 | 150 | 6.6 | 6.6 | 0.682 | 0.790 | 87.3 | 49.6 | 82 | 9.8 |
| 6ro4 | D | 3.5 | 4970 | 0.015 | 1.05 | 280 | 13.0 | 10.8 | 0.665 | 0.669 | 82.1 | 26.1 | 70.7 | 7.1 |
| 6ro4 | E | 3.5 | 4970 | 0.015 | 1.05 | 237 | 13.5 | 10.6 | 0.582 | 0.704 | 87.8 | 53.8 | 86.1 | 10.8 |
| 6rqf | A | 3.6 | 4981 | 0.0144 | 1.065 | 215 | 2.4 | 2.4 | 0.935 | 0.935 | 95.8 | 78.2 | 89.8 | 62.7 |
| 6rqf | C | 3.6 | 4981 | 0.0144 | 1.065 | 285 | 7.8 | 7.2 | 0.857 | 0.857 | 88.8 | 51.4 | 61.1 | 24.1 |
| 6rqf | D | 3.6 | 4981 | 0.0144 | 1.065 | 179 | 8.2 | 8.2 | 0.809 | 0.816 | 81 | 53.8 | 69.8 | 26.4 |
| 6rqf | I | 3.6 | 4981 | 0.0144 | 1.065 | 215 | 2.3 | 2.3 | 0.934 | 0.934 | 94.9 | 81.9 | 89.3 | 64.1 |
| 6rqf | K | 3.6 | 4981 | 0.0144 | 1.065 | 285 | 14.3 | 14.3 | 0.669 | 0.670 | 87 | 38.3 | 66.3 | 25.4 |
| 6rqf | L | 3.6 | 4981 | 0.0144 | 1.065 | 179 | 15.3 | 7.6 | 0.748 | 0.808 | 77.7 | 21.6 | 73.2 | 26.7 |
| 6rqh | E | 3.7 | 4982 | 0.025 | 1.041 | 215 | 5.2 | 5.2 | 0.842 | 0.842 | 84.7 | 35.7 | 71.6 | 16.9 |
| 6rqh | H | 3.7 | 4982 | 0.025 | 1.041 | 134 | 16.4 | 14.7 | 0.436 | 0.545 | 78.4 | 8.6 | 35.1 | 4.3 |
| 6rql | E | 2.9 | 4984 | 0.04 | 1.04 | 215 | 4.5 | 4.5 | 0.828 | 0.832 | 92.6 | 74.4 | 80.9 | 29.3 |
| 6rql | F | 2.9 | 4984 | 0.04 | 1.04 | 100 | 1.6 | 1.6 | 0.906 | 0.906 | 95 | 85.3 | 86 | 57 |
| 6rql | H | 2.9 | 4984 | 0.04 | 1.04 | 134 | 2.5 | 2.5 | 0.880 | 0.880 | 91.8 | 71.5 | 82.1 | 3.6 |
| 6rqt | H | 4 | 4985 | 0.05 | 1.3268 | 134 | 14.9 | 14.3 | 0.249 | 0.397 | 59.7 | 10 | 29.9 | 12.5 |
| 6rrd | E | 3.1 | 4987 | 0.035 | 1.041 | 215 | 4.5 | 4.5 | 0.830 | 0.830 | 93 | 72.5 | 85.6 | 40.2 |
| 6rrd | F | 3.1 | 4987 | 0.035 | 1.041 | 100 | 1.7 | 1.7 | 0.935 | 0.935 | 97 | 80.4 | 95 | 66.3 |
| 6rrd | H | 3.1 | 4987 | 0.035 | 1.041 | 134 | 2.9 | 2.9 | 0.904 | 0.904 | 93.3 | 74.4 | 91 | 15.6 |
| 6rui | E | 2.7 | 10006 | 0.04 | 1.04 | 215 | 3.6 | 3.6 | 0.893 | 0.893 | 92.1 | 63.6 | 80.9 | 56.3 |
| 6rui | F | 2.7 | 10006 | 0.04 | 1.04 | 100 | 1.6 | 1.6 | 0.928 | 0.928 | 97 | 85.6 | 93 | 63.4 |
| 6rui | H | 2.7 | 10006 | 0.04 | 1.04 | 134 | 10.8 | 10.8 | 0.855 | 0.855 | 89.6 | 25.8 | 55.2 | 9.5 |
| 6rui | J | 2.7 | 10006 | 0.04 | 1.04 | 69 | 1.2 | 1.2 | 0.901 | 0.901 | 97.1 | 98.5 | 89.9 | 6.5 |
| 6rui | K | 2.7 | 10006 | 0.04 | 1.04 | 103 | 3.8 | 3.8 | 0.901 | 0.901 | 92.2 | 63.2 | 90.3 | 74.2 |
| 6rzb | C | 4.1 | 10061 | 0.013 | 1.04 | 149 | 15.9 | 14.3 | 0.396 | 0.452 | 66.4 | 7.1 | 43.6 | 10.8 |
| 6s01 | C | 3.2 | 10069 | 0.05 | 1.05 | 107 | 1.6 | 1.6 | 0.938 | 0.938 | 97.2 | 83.7 | 91.6 | 65.3 |
| 6s01 | D | 3.2 | 10069 | 0.05 | 1.05 | 94 | 1.0 | 1.0 | 0.944 | 0.944 | 98.9 | 98.9 | 92.6 | 81.6 |
| 6s01 | G | 3.2 | 10069 | 0.05 | 1.05 | 107 | 2.3 | 2.3 | 0.910 | 0.910 | 95.3 | 69.6 | 87.9 | 55.3 |
| 6s01 | H | 3.2 | 10069 | 0.05 | 1.05 | 94 | 1.5 | 1.5 | 0.937 | 0.937 | 96.8 | 90.1 | 93.6 | 61.4 |
| 6s0k | a | 3.1 | 10073 | 0.82 | 1.39 | 58 | 1.9 | 1.9 | 0.856 | 0.856 | 93.1 | 83.3 | 93.1 | 50 |
| 6s0k | C | 3.1 | 10073 | 0.82 | 1.39 | 272 | 4.1 | 4.1 | 0.889 | 0.889 | 91.9 | 77.9 | 82.3 | 11.2 |
| 6s0k | D | 3.1 | 10073 | 0.82 | 1.39 | 209 | 2.6 | 2.6 | 0.915 | 0.915 | 93.3 | 69.2 | 79.9 | 24.6 |
| 6s0k | E | 3.1 | 10073 | 0.82 | 1.39 | 201 | 3.3 | 3.3 | 0.900 | 0.905 | 95 | 80.1 | 85.6 | 48.8 |
| 6s0k | F | 3.1 | 10073 | 0.82 | 1.39 | 178 | 3.0 | 3.0 | 0.859 | 0.859 | 90.4 | 47.5 | 67.8 | 12.5 |

| | | | | | | | | | | | | | | |
|---|---|---|---|---|---|---|---|---|---|---|---|---|---|---|
| 6s0k | G | 3.1 | 10073 | 0.82 | 1.39 | 176 | 9.2 | 6.4 | 0.797 | 0.804 | 85.8 | 41.1 | 70.5 | 8.9 |
| 6s0k | K | 3.1 | 10073 | 0.82 | 1.39 | 142 | 2.6 | 2.6 | 0.898 | 0.898 | 93 | 74.2 | 84.5 | 30 |
| 6s0k | L | 3.1 | 10073 | 0.82 | 1.39 | 123 | 2.5 | 2.5 | 0.889 | 0.889 | 93.5 | 73.9 | 52.8 | 7.7 |
| 6s0k | N | 3.1 | 10073 | 0.82 | 1.39 | 136 | 1.1 | 1.1 | 0.949 | 0.949 | 99.3 | 99.3 | 94.9 | 5.4 |
| 6s0k | O | 3.1 | 10073 | 0.82 | 1.39 | 125 | 1.7 | 1.7 | 0.921 | 0.921 | 96.8 | 90.9 | 84.8 | 41.5 |
| 6s0k | P | 3.1 | 10073 | 0.82 | 1.39 | 117 | 4.1 | 4.1 | 0.852 | 0.861 | 91.5 | 45.8 | 83.8 | 28.6 |
| 6s0k | Q | 3.1 | 10073 | 0.82 | 1.39 | 114 | 2.8 | 2.8 | 0.887 | 0.887 | 92.1 | 82.9 | 78.9 | 5.6 |
| 6s0k | S | 3.1 | 10073 | 0.82 | 1.39 | 103 | 2.6 | 2.6 | 0.877 | 0.877 | 94.2 | 69.1 | 68 | 28.6 |
| 6s0k | T | 3.1 | 10073 | 0.82 | 1.39 | 110 | 3.7 | 3.7 | 0.908 | 0.908 | 94.5 | 69.2 | 93.6 | 31.1 |
| 6s0k | U | 3.1 | 10073 | 0.82 | 1.39 | 92 | 2.5 | 2.5 | 0.865 | 0.865 | 91.3 | 72.6 | 82.6 | 6.6 |
| 6s0k | V | 3.1 | 10073 | 0.82 | 1.39 | 103 | 3.5 | 3.5 | 0.799 | 0.799 | 86.3 | 50 | 84.3 | 7 |
| 6s0k | W | 3.1 | 10073 | 0.82 | 1.39 | 94 | 2.6 | 2.6 | 0.844 | 0.844 | 89.4 | 65.5 | 58.5 | 5.5 |
| 6s0k | Y | 3.1 | 10073 | 0.82 | 1.39 | 77 | 1.8 | 1.8 | 0.879 | 0.879 | 96.1 | 78.4 | 85.7 | 50 |
| 6s0k | Z | 3.1 | 10073 | 0.82 | 1.39 | 62 | 3.5 | 3.5 | 0.804 | 0.804 | 88.7 | 49.1 | 88.7 | 52.7 |
| 6s1m | D | 4.3 | 10080 | 0.015 | 1.08 | 66 | 3.5 | 3.5 | 0.668 | 0.668 | 81.8 | 51.9 | 62.1 | 34.1 |
| 6s1m | E | 4.3 | 10080 | 0.015 | 1.08 | 251 | 20.7 | 17.3 | 0.258 | 0.268 | 54.2 | 4.4 | 24.3 | 3.3 |
| 6s1m | F | 4.3 | 10080 | 0.015 | 1.08 | 249 | 19.0 | 17.2 | 0.310 | 0.385 | 59.8 | 12.1 | 44.2 | 7.3 |
| 6s1m | G | 4.3 | 10080 | 0.015 | 1.08 | 249 | 23.2 | 17.5 | 0.237 | 0.267 | 60.6 | 6.6 | 28.9 | 4.2 |
| 6s1n | D | 4.9 | 10081 | 0.014 | 1.08 | 66 | 4.2 | 4.2 | 0.549 | 0.549 | 74.2 | 42.9 | 48.5 | 12.5 |
| 6s1n | E | 4.9 | 10081 | 0.014 | 1.08 | 251 | 22.0 | 18.7 | 0.244 | 0.282 | 53 | 6 | 31.1 | 10.3 |
| 6s1n | F | 4.9 | 10081 | 0.014 | 1.08 | 249 | 21.6 | 19.2 | 0.249 | 0.303 | 59 | 5.4 | 26.5 | 9.1 |
| 6s1n | G | 4.9 | 10081 | 0.014 | 1.08 | 249 | 19.6 | 18.6 | 0.245 | 0.293 | 51 | 10.2 | 26.1 | 6.2 |
| 6s8g | A | 3.5 | 10121 | 0.05 | 1.014 | 238 | 2.2 | 2.2 | 0.933 | 0.933 | 94.5 | 75.1 | 89.1 | 17 |
| 6s8g | B | 3.5 | 10121 | 0.05 | 1.014 | 238 | 3.0 | 3.0 | 0.920 | 0.920 | 94.6 | 79.2 | 88.7 | 42.5 |
| 6s8h | A | 3.7 | 10122 | 0.05 | 1.014 | 239 | 3.1 | 3.1 | 0.927 | 0.927 | 95.4 | 74.6 | 86.2 | 6.8 |
| 6s8h | B | 3.7 | 10122 | 0.05 | 1.014 | 238 | 5.7 | 5.7 | 0.887 | 0.887 | 92.9 | 73.8 | 75.2 | 32.4 |
| 6s8h | F | 3.7 | 10122 | 0.05 | 1.014 | 239 | 22.5 | 22.2 | 0.315 | 0.345 | 53.1 | 5.5 | 82.4 | 48.7 |
| 6s8h | G | 3.7 | 10122 | 0.05 | 1.014 | 243 | 22.3 | 18.9 | 0.369 | 0.403 | 64.6 | 29.3 | 82.3 | 42 |
| 6s8n | A | 3.1 | 10125 | 0.03 | 1.014 | 239 | 1.6 | 1.6 | 0.956 | 0.956 | 97.1 | 89.2 | 94.1 | 83.6 |
| 6s8n | B | 3.1 | 10125 | 0.03 | 1.014 | 238 | 1.7 | 1.7 | 0.944 | 0.944 | 96.2 | 80.3 | 91.2 | 47.9 |
| 6se6 | D | 3.5 | 10152 | 0.0134 | 0.65 | 92 | 2.7 | 2.7 | 0.879 | 0.879 | 92.4 | 80 | 93.5 | 66.3 |
| 6se6 | H | 3.5 | 10152 | 0.0134 | 0.65 | 90 | 2.2 | 2.2 | 0.901 | 0.901 | 95.6 | 82.6 | 94.4 | 84.7 |
| 6see | H | 4.2 | 10153 | 0.012 | 1.3 | 90 | 3.9 | 3.9 | 0.745 | 0.745 | 82.2 | 45.9 | 44.4 | 7.5 |
| 6sef | D | 3.7 | 10154 | 0.012 | 1.3 | 92 | 3.7 | 3.5 | 0.838 | 0.838 | 91.3 | 51.2 | 90.2 | 1.2 |
| 6sef | H | 3.7 | 10154 | 0.012 | 1.3 | 90 | 2.2 | 2.2 | 0.881 | 0.881 | 94.4 | 71.8 | 88.9 | 2.5 |
| 6sj7 | C | 3.5 | 10213 | 0.006 | 0.86 | 79 | 12.7 | 10.3 | 0.570 | 0.570 | 70.9 | 16.1 | 53.2 | 4.8 |
| 6sof | D | 4.3 | 10273 | 0.015 | 1.06 | 162 | 22.5 | 15.8 | 0.276 | 0.295 | 57.4 | 4.3 | 33.3 | 7.4 |

| | | | | | | | | | | | | | | |
|---|---|---|---|---|---|---|---|---|---|---|---|---|---|---|
| 6spb | T | 2.8 | 10280 | 0.0166 | 1.1 | 92 | 1.6 | 1.6 | 0.906 | 0.906 | 96.7 | 83.1 | 46.7 | 7 |
| 6spb | Y | 2.8 | 10280 | 0.0166 | 1.1 | 60 | 1.5 | 1.5 | 0.892 | 0.892 | 96.7 | 82.8 | 95 | 93 |
| 6spc | b | 3 | 10281 | 0.00936 | 1.1 | 221 | 4.3 | 4.3 | 0.866 | 0.875 | 87.8 | 60.8 | 77.8 | 29.7 |
| 6spc | f | 3 | 10281 | 0.00936 | 1.1 | 100 | 2.8 | 2.8 | 0.843 | 0.859 | 92 | 83.7 | 93 | 8.6 |
| 6spc | o | 3 | 10281 | 0.00936 | 1.1 | 86 | 0.7 | 0.7 | 0.958 | 0.958 | 100 | 100 | 89.5 | 70.1 |
| 6spd | G | 3.3 | 10282 | 0.0566 | 1.1 | 173 | 9.7 | 6.3 | 0.770 | 0.860 | 83.8 | 57.9 | 78 | 26.7 |
| 6spd | V | 3.3 | 10282 | 0.0566 | 1.1 | 188 | 3.9 | 3.9 | 0.894 | 0.894 | 92 | 53.8 | 44.7 | 2.4 |
| 6spe | c | 3.6 | 10283 | 0.028 | 1.1 | 205 | 3.3 | 3.3 | 0.892 | 0.892 | 91.7 | 52.7 | 80 | 6.1 |
| 6spe | d | 3.6 | 10283 | 0.028 | 1.1 | 205 | 3.1 | 3.1 | 0.907 | 0.907 | 94.1 | 56.5 | 71.7 | 27.9 |
| 6spe | e | 3.6 | 10283 | 0.028 | 1.1 | 156 | 4.8 | 4.8 | 0.846 | 0.846 | 90.4 | 66 | 84.6 | 9.8 |
| 6spe | f | 3.6 | 10283 | 0.028 | 1.1 | 105 | 2.5 | 2.5 | 0.839 | 0.839 | 92.4 | 60.8 | 56.2 | 8.5 |
| 6spe | g | 3.6 | 10283 | 0.028 | 1.1 | 154 | 4.5 | 4.5 | 0.832 | 0.832 | 85.1 | 35.9 | 75.3 | 11.2 |
| 6spe | h | 3.6 | 10283 | 0.028 | 1.1 | 129 | 2.3 | 2.3 | 0.900 | 0.900 | 93 | 82.5 | 89.1 | 5.2 |
| 6spe | o | 3.6 | 10283 | 0.028 | 1.1 | 86 | 1.9 | 1.9 | 0.858 | 0.858 | 91.9 | 84.8 | 81.4 | 50 |
| 6spe | p | 3.6 | 10283 | 0.028 | 1.1 | 78 | 1.7 | 1.7 | 0.868 | 0.868 | 94.9 | 83.8 | 73.1 | 7 |
| 6spe | q | 3.6 | 10283 | 0.028 | 1.1 | 76 | 2.9 | 2.9 | 0.838 | 0.838 | 90.8 | 60.9 | 92.1 | 25.7 |
| 6spe | r | 3.6 | 10283 | 0.028 | 1.1 | 71 | 3.8 | 3.8 | 0.783 | 0.783 | 90.1 | 51.6 | 67.6 | 43.8 |
| 6swy | 4 | 3.2 | 10333 | 0.0306 | 1.09 | 217 | 3.7 | 3.7 | 0.895 | 0.895 | 90.3 | 63.8 | 87.6 | 38.4 |
| 6tdv | d | 2.8 | 10468 | 0.025 | 1.05 | 186 | 1.5 | 1.5 | 0.944 | 0.944 | 96.2 | 89.4 | 91.4 | 27.6 |
| 6tdv | D | 2.8 | 10468 | 0.025 | 1.05 | 186 | 1.4 | 1.4 | 0.962 | 0.962 | 97.8 | 90.1 | 88.2 | 48.2 |
| 6tdv | e | 2.8 | 10468 | 0.025 | 1.05 | 96 | 1.5 | 1.5 | 0.937 | 0.937 | 97.9 | 88.3 | 95.8 | 94.6 |
| 6tdv | E | 2.8 | 10468 | 0.025 | 1.05 | 96 | 1.2 | 1.2 | 0.938 | 0.938 | 97.9 | 92.6 | 91.7 | 85.2 |
| 6tdv | f | 2.8 | 10468 | 0.025 | 1.05 | 274 | 1.3 | 1.3 | 0.967 | 0.967 | 98.5 | 91.9 | 94.2 | 66.7 |
| 6tdv | F | 2.8 | 10468 | 0.025 | 1.05 | 273 | 1.7 | 1.7 | 0.962 | 0.962 | 97.1 | 85 | 92 | 80.2 |
| 6tdv | i | 2.8 | 10468 | 0.025 | 1.05 | 97 | 1.4 | 1.4 | 0.914 | 0.914 | 94.8 | 94.6 | 94.8 | 82.6 |
| 6tdv | I | 2.8 | 10468 | 0.025 | 1.05 | 97 | 1.7 | 1.7 | 0.916 | 0.916 | 96.9 | 89.4 | 91.8 | 74.2 |
| 6tdv | j | 2.8 | 10468 | 0.025 | 1.05 | 103 | 1.3 | 1.3 | 0.944 | 0.944 | 98.1 | 93.1 | 90.3 | 77.4 |
| 6tdv | J | 2.8 | 10468 | 0.025 | 1.05 | 103 | 1.0 | 1.0 | 0.953 | 0.953 | 98.1 | 97 | 88.3 | 61.5 |
| 6tdv | p | 2.8 | 10468 | 0.025 | 1.05 | 114 | 2.6 | 2.6 | 0.921 | 0.921 | 94.7 | 96.3 | 90.4 | 81.6 |
| 6tdv | q | 2.8 | 10468 | 0.025 | 1.05 | 89 | 1.7 | 1.7 | 0.902 | 0.902 | 95.5 | 83.5 | 89.9 | 62.5 |
| 6tdv | Q | 2.8 | 10468 | 0.025 | 1.05 | 89 | 2.4 | 2.4 | 0.900 | 0.900 | 95.5 | 58.8 | 91 | 48.1 |
| 6tdw | C | 3.8 | 10469 | 0.02 | 1.05 | 157 | 3.6 | 3.6 | 0.831 | 0.839 | 86 | 57 | 82.8 | 31.5 |
| 6tdx | G | 3.3 | 10470 | 0.025 | 1.05 | 276 | 1.7 | 1.7 | 0.955 | 0.955 | 96.4 | 85 | 88.4 | 55.7 |
| 6tdy | M | 3 | 10471 | 0.025 | 1.05 | 243 | 7.0 | 6.8 | 0.755 | 0.761 | 87.7 | 49.8 | 68.7 | 25.1 |
| 6tdz | H | 3.1 | 10472 | 0.025 | 1.05 | 160 | 2.5 | 2.5 | 0.900 | 0.900 | 93.8 | 82.7 | 85 | 5.1 |
| 6tdz | I | 3.1 | 10472 | 0.025 | 1.05 | 66 | 1.7 | 1.7 | 0.858 | 0.858 | 97 | 84.4 | 87.9 | 70.7 |
| 6tdz | J | 3.1 | 10472 | 0.025 | 1.05 | 170 | 1.5 | 1.5 | 0.951 | 0.951 | 98.2 | 91.6 | 92.9 | 65.2 |

| | | | | | | | | | | | | | | |
|---|---|---|---|---|---|---|---|---|---|---|---|---|---|---|
| 6tdz | K | 3.1 | 10472 | 0.025 | 1.05 | 170 | 1.3 | 1.3 | 0.950 | 0.950 | 97.1 | 94.5 | 90.6 | 40.9 |
| 6tdz | L | 3.1 | 10472 | 0.025 | 1.05 | 170 | 1.1 | 1.1 | 0.961 | 0.961 | 98.2 | 97.6 | 88.2 | 89.3 |
| 6tdz | M | 3.1 | 10472 | 0.025 | 1.05 | 243 | 3.1 | 3.1 | 0.900 | 0.900 | 90.9 | 62.4 | 61.7 | 23.3 |
| 6te0 | J | 3.9 | 10473 | 0.017 | 1.05 | 170 | 5.1 | 5.1 | 0.862 | 0.862 | 89.4 | 49.3 | 84.7 | 29.9 |
| 6te0 | K | 3.9 | 10473 | 0.017 | 1.05 | 170 | 4.4 | 4.4 | 0.809 | 0.809 | 83.5 | 31 | 73.5 | 16 |
| 6te0 | L | 3.9 | 10473 | 0.017 | 1.05 | 170 | 2.4 | 2.4 | 0.888 | 0.888 | 93.5 | 72.3 | 83.5 | 32.4 |
| 6te0 | M | 3.9 | 10473 | 0.017 | 1.05 | 243 | 15.8 | 7.7 | 0.494 | 0.717 | 82.3 | 18.5 | 58.4 | 5.6 |
| 6tny | C | 3.1 | 10539 | 0.0095 | 0.87 | 143 | 8.9 | 7.0 | 0.764 | 0.773 | 81.1 | 47.4 | 70.6 | 5.9 |
| 6tny | D | 3.1 | 10539 | 0.0095 | 0.87 | 66 | 2.4 | 2.4 | 0.840 | 0.840 | 92.4 | 67.2 | 87.9 | 72.4 |
| 6tny | F | 3.1 | 10539 | 0.0095 | 0.87 | 249 | 17.5 | 15.2 | 0.687 | 0.737 | 73.9 | 14.7 | 70.3 | 2.3 |
| 6tnz | C | 4.1 | 10540 | 0.0073 | 0.87 | 143 | 17.2 | 10.9 | 0.412 | 0.585 | 64.3 | 9.8 | 58.7 | 7.1 |
| 6tnz | D | 4.1 | 10540 | 0.0073 | 0.87 | 66 | 14.0 | 3.8 | 0.280 | 0.695 | 83.3 | 3.6 | 65.2 | 0 |
| 6tnz | E | 4.1 | 10540 | 0.0073 | 0.87 | 251 | 19.9 | 18.6 | 0.235 | 0.288 | 55 | 5.1 | 33.9 | 1.2 |
| 6tnz | F | 4.1 | 10540 | 0.0073 | 0.87 | 249 | 22.4 | 19.0 | 0.369 | 0.387 | 71.1 | 7.9 | 26.1 | 6.2 |
| 6tnz | G | 4.1 | 10540 | 0.0073 | 0.87 | 249 | 19.4 | 18.0 | 0.244 | 0.331 | 59 | 7.5 | 43 | 4.7 |
| 6tys | C | 3.5 | 20584 | 0.9 | 1.37 | 120 | 1.7 | 1.7 | 0.918 | 0.918 | 95.8 | 88.7 | 90 | 13 |
| 6tys | D | 3.5 | 20584 | 0.9 | 1.37 | 107 | 2.1 | 2.1 | 0.906 | 0.906 | 95.3 | 74.5 | 86 | 12 |
| 6tys | F | 3.5 | 20584 | 0.9 | 1.37 | 120 | 2.2 | 2.2 | 0.904 | 0.904 | 96.7 | 78.4 | 79.2 | 11.6 |
| 6tys | G | 3.5 | 20584 | 0.9 | 1.37 | 107 | 1.4 | 1.4 | 0.916 | 0.916 | 98.1 | 93.3 | 86 | 7.6 |
| 6tys | H | 3.5 | 20584 | 0.9 | 1.37 | 120 | 4.3 | 4.0 | 0.914 | 0.914 | 96.7 | 72.4 | 82.5 | 8.1 |
| 6tys | L | 3.5 | 20584 | 0.9 | 1.37 | 107 | 5.2 | 3.0 | 0.777 | 0.878 | 95.3 | 72.5 | 90.7 | 10.3 |
| 6u0l | D | 3.3 | 20605 | 0.028 | 1.057 | 97 | 8.7 | 8.7 | 0.426 | 0.713 | 86.6 | 14.3 | 68 | 4.5 |
| 6u0l | E | 3.3 | 20605 | 0.028 | 1.057 | 97 | 14.2 | 14.1 | 0.517 | 0.655 | 79.4 | 11.7 | 38.1 | 8.1 |
| 6u0l | F | 3.3 | 20605 | 0.028 | 1.057 | 97 | 13.4 | 11.8 | 0.465 | 0.479 | 84.5 | 11 | 66 | 10.9 |
| 6u0l | H | 3.3 | 20605 | 0.028 | 1.057 | 130 | 17.5 | 16.1 | 0.283 | 0.354 | 66.9 | 4.6 | 35.4 | 8.7 |
| 6u0l | I | 3.3 | 20605 | 0.028 | 1.057 | 130 | 17.8 | 15.0 | 0.219 | 0.365 | 64.6 | 7.1 | 35.4 | 4.3 |
| 6u0l | J | 3.3 | 20605 | 0.028 | 1.057 | 110 | 13.9 | 13.8 | 0.278 | 0.320 | 66.4 | 5.5 | 26.4 | 10.3 |
| 6u0l | L | 3.3 | 20605 | 0.028 | 1.057 | 110 | 15.2 | 13.9 | 0.368 | 0.368 | 65.5 | 13.9 | 32.7 | 5.6 |
| 6u0l | P | 3.3 | 20605 | 0.028 | 1.057 | 130 | 16.1 | 15.5 | 0.259 | 0.349 | 60 | 14.1 | 38.5 | 8 |
| 6u0l | Q | 3.3 | 20605 | 0.028 | 1.057 | 110 | 12.6 | 12.5 | 0.377 | 0.378 | 70 | 3.9 | 31.8 | 2.9 |
| 6u0l | X | 3.3 | 20605 | 0.028 | 1.057 | 133 | 23.3 | 18.3 | 0.244 | 0.646 | 83.5 | 4.5 | 78.9 | 6.7 |
| 6u0l | Z | 3.3 | 20605 | 0.028 | 1.057 | 121 | 19.1 | 18.1 | 0.275 | 0.661 | 86.8 | 4.8 | 61.2 | 8.1 |
| 6u0n | D | 3.5 | 20608 | 0.02 | 1.057 | 97 | 12.7 | 10.4 | 0.329 | 0.664 | 81.4 | 5.1 | 48.5 | 10.6 |
| 6u0n | E | 3.5 | 20608 | 0.02 | 1.057 | 97 | 9.6 | 9.6 | 0.584 | 0.584 | 75.3 | 37 | 57.7 | 10.7 |
| 6u0n | F | 3.5 | 20608 | 0.02 | 1.057 | 97 | 6.7 | 4.9 | 0.816 | 0.816 | 89.7 | 29.9 | 60.8 | 8.5 |
| 6u0n | H | 3.5 | 20608 | 0.02 | 1.057 | 130 | 12.4 | 12.4 | 0.601 | 0.605 | 73.8 | 8.3 | 32.3 | 4.8 |
| 6u0n | I | 3.5 | 20608 | 0.02 | 1.057 | 130 | 18.0 | 15.9 | 0.324 | 0.463 | 80 | 9.6 | 49.2 | 9.4 |

| | | | | | | | | | | | | | | |
|---|---|---|---|---|---|---|---|---|---|---|---|---|---|---|
| 6u0n | J | 3.5 | 20608 | 0.02 | 1.057 | 110 | 15.0 | 13.5 | 0.406 | 0.469 | 65.5 | 6.9 | 38.2 | 2.4 |
| 6u0n | L | 3.5 | 20608 | 0.02 | 1.057 | 110 | 15.1 | 13.2 | 0.239 | 0.460 | 79.1 | 4.6 | 45.5 | 10 |
| 6u0n | P | 3.5 | 20608 | 0.02 | 1.057 | 130 | 17.4 | 15.0 | 0.281 | 0.440 | 68.5 | 3.4 | 26.2 | 5.9 |
| 6u0n | Q | 3.5 | 20608 | 0.02 | 1.057 | 110 | 17.0 | 14.7 | 0.277 | 0.278 | 67.3 | 1.4 | 12.7 | 7.1 |
| 6u0n | X | 3.5 | 20608 | 0.02 | 1.057 | 129 | 8.1 | 8.1 | 0.617 | 0.624 | 86 | 57.7 | 79.1 | 36.3 |
| 6u0n | Z | 3.5 | 20608 | 0.02 | 1.057 | 124 | 19.3 | 16.7 | 0.663 | 0.716 | 82.3 | 28.4 | 68.5 | 42.4 |
| 6u62 | B | 3.2 | 20660 | 0.01 | 1.059 | 298 | 2.0 | 2.0 | 0.953 | 0.953 | 96.3 | 81.9 | 94.3 | 65.1 |
| 6u62 | C | 3.2 | 20660 | 0.01 | 1.059 | 281 | 2.9 | 2.9 | 0.930 | 0.930 | 91.5 | 84 | 90.7 | 72.5 |
| 6u62 | E | 3.2 | 20660 | 0.01 | 1.059 | 124 | 3.2 | 3.2 | 0.872 | 0.872 | 91.9 | 45.6 | 81.5 | 51.5 |
| 6u62 | F | 3.2 | 20660 | 0.01 | 1.059 | 120 | 13.1 | 8.0 | 0.706 | 0.732 | 80.8 | 19.6 | 77.5 | 30.1 |
| 6uan | A | 3.9 | 20708 | 0.014 | 1.06 | 231 | 5.0 | 5.0 | 0.840 | 0.840 | 85.7 | 34.3 | 89.6 | 27.5 |
| 6uan | B | 3.9 | 20708 | 0.014 | 1.06 | 260 | 13.1 | 11.1 | 0.683 | 0.697 | 73.5 | 28.3 | 66.2 | 4.1 |
| 6uan | C | 3.9 | 20708 | 0.014 | 1.06 | 284 | 12.5 | 12.5 | 0.575 | 0.601 | 72.2 | 11.2 | 64.1 | 5.5 |
| 6uan | Z | 3.9 | 20708 | 0.014 | 1.06 | 230 | 8.2 | 6.4 | 0.556 | 0.822 | 72.6 | 25.7 | 88.3 | 41.9 |
| 6ud8 | E | 3.2 | 20734 | 0.0218 | 1.066 | 132 | 5.6 | 5.6 | 0.796 | 0.806 | 82.6 | 51.4 | 82.6 | 33 |
| 6ud8 | F | 3.2 | 20734 | 0.0218 | 1.066 | 134 | 5.5 | 5.5 | 0.782 | 0.782 | 81.3 | 56.9 | 81.3 | 40.4 |
| 6ud8 | G | 3.2 | 20734 | 0.0218 | 1.066 | 132 | 5.7 | 5.7 | 0.789 | 0.789 | 83.3 | 58.2 | 72.7 | 51 |
| 6ud8 | H | 3.2 | 20734 | 0.0218 | 1.066 | 134 | 3.9 | 3.9 | 0.851 | 0.851 | 90.3 | 66.1 | 81.3 | 65.1 |
| 6uda | C | 4.2 | 20735 | 0.7 | 1.08 | 220 | 20.3 | 18.7 | 0.430 | 0.470 | 65.9 | 8.3 | 45.9 | 7.9 |
| 6uda | D | 4.2 | 20735 | 0.7 | 1.08 | 210 | 16.5 | 16.4 | 0.303 | 0.340 | 61.9 | 10 | 31.9 | 10.4 |
| 6uda | F | 4.2 | 20735 | 0.7 | 1.08 | 126 | 5.6 | 5.6 | 0.721 | 0.721 | 77.8 | 21.4 | 72.2 | 33 |
| 6uda | G | 4.2 | 20735 | 0.7 | 1.08 | 220 | 21.3 | 19.4 | 0.458 | 0.458 | 60.5 | 10.5 | 34.1 | 6.7 |
| 6uda | H | 4.2 | 20735 | 0.7 | 1.08 | 210 | 18.0 | 18.0 | 0.269 | 0.280 | 65.2 | 9.5 | 29.5 | 4.8 |
| 6uda | J | 4.2 | 20735 | 0.7 | 1.08 | 126 | 15.5 | 5.0 | 0.306 | 0.756 | 74.6 | 10.6 | 48.4 | 3.3 |
| 6uda | K | 4.2 | 20735 | 0.7 | 1.08 | 220 | 21.3 | 19.8 | 0.298 | 0.501 | 64.5 | 9.9 | 37.7 | 7.2 |
| 6uda | L | 4.2 | 20735 | 0.7 | 1.08 | 210 | 16.7 | 15.1 | 0.286 | 0.321 | 56.7 | 6.7 | 18.1 | 2.6 |
| 6uda | Z | 4.2 | 20735 | 0.7 | 1.08 | 126 | 4.8 | 4.8 | 0.781 | 0.781 | 84.9 | 29 | 78.6 | 12.1 |
| 6uh5 | A | 3.5 | 20767 | 0.009 | 1.056 | 98 | 1.1 | 1.1 | 0.940 | 0.940 | 98 | 95.8 | 93.9 | 73.9 |
| 6uh5 | B | 3.5 | 20767 | 0.009 | 1.056 | 82 | 3.6 | 3.5 | 0.851 | 0.851 | 92.7 | 90.8 | 79.3 | 80 |
| 6uh5 | C | 3.5 | 20767 | 0.009 | 1.056 | 107 | 1.8 | 1.8 | 0.911 | 0.911 | 96.3 | 79.6 | 73.8 | 44.3 |
| 6uh5 | D | 3.5 | 20767 | 0.009 | 1.056 | 93 | 1.9 | 1.9 | 0.897 | 0.897 | 94.6 | 83 | 90.3 | 81 |
| 6uh5 | H | 3.5 | 20767 | 0.009 | 1.056 | 93 | 1.3 | 1.3 | 0.920 | 0.920 | 96.8 | 90 | 88.2 | 86.6 |
| 6uiv | A | 3.3 | 20788 | 0.0105 | 1.026 | 287 | 17.8 | 5.6 | 0.745 | 0.860 | 85 | 40.6 | 61.7 | 27.1 |
| 6uiv | B | 3.3 | 20788 | 0.0105 | 1.026 | 287 | 5.4 | 5.4 | 0.856 | 0.856 | 85.7 | 51.6 | 55.1 | 52.5 |
| 6uiv | C | 3.3 | 20788 | 0.0105 | 1.026 | 287 | 6.4 | 5.9 | 0.783 | 0.824 | 82.2 | 36.4 | 63.4 | 54.4 |
| 6uiv | D | 3.3 | 20788 | 0.0105 | 1.026 | 287 | 6.3 | 3.9 | 0.861 | 0.876 | 86.1 | 53.8 | 59.2 | 44.7 |
| 6uiv | E | 3.3 | 20788 | 0.0105 | 1.026 | 287 | 4.9 | 4.9 | 0.863 | 0.876 | 86.4 | 52.4 | 22.6 | 7.7 |

| | | | | | | | | | | | | | | |
|---|---|---|---|---|---|---|---|---|---|---|---|---|---|---|
| 6uiv | F | 3.3 | 20788 | 0.0105 | 1.026 | 287 | 4.2 | 4.2 | 0.884 | 0.889 | 91.6 | 59.3 | 43.2 | 5.6 |
| 6uiv | G | 3.3 | 20788 | 0.0105 | 1.026 | 287 | 5.4 | 5.3 | 0.883 | 0.883 | 89.2 | 40.2 | 55.7 | 5.6 |
| 6uiv | H | 3.3 | 20788 | 0.0105 | 1.026 | 287 | 8.4 | 5.4 | 0.851 | 0.872 | 84.3 | 38.4 | 27.9 | 5 |
| 6uiv | I | 3.3 | 20788 | 0.0105 | 1.026 | 287 | 5.9 | 4.0 | 0.867 | 0.882 | 86.1 | 50.6 | 64.5 | 5.9 |
| 6uiv | J | 3.3 | 20788 | 0.0105 | 1.026 | 287 | 5.7 | 5.7 | 0.860 | 0.860 | 85.4 | 40 | 64.1 | 29.9 |
| 6uiv | K | 3.3 | 20788 | 0.0105 | 1.026 | 287 | 16.2 | 5.2 | 0.674 | 0.876 | 87.8 | 49.6 | 59.6 | 5.8 |
| 6uiw | A | 2.7 | 20789 | 0.012 | 1.026 | 294 | 6.3 | 4.6 | 0.845 | 0.888 | 87.1 | 55.5 | 73.5 | 31.5 |
| 6uiw | B | 2.7 | 20789 | 0.012 | 1.026 | 294 | 3.9 | 3.9 | 0.923 | 0.923 | 93.2 | 68.6 | 69.7 | 48.3 |
| 6uiw | C | 2.7 | 20789 | 0.012 | 1.026 | 294 | 3.2 | 3.2 | 0.917 | 0.917 | 91.8 | 71.9 | 76.9 | 41.6 |
| 6uiw | D | 2.7 | 20789 | 0.012 | 1.026 | 294 | 4.2 | 4.2 | 0.921 | 0.921 | 94.2 | 76.9 | 72.8 | 50 |
| 6uiw | E | 2.7 | 20789 | 0.012 | 1.026 | 294 | 3.2 | 3.2 | 0.919 | 0.919 | 92.2 | 73.8 | 72.4 | 49.8 |
| 6uiw | F | 2.7 | 20789 | 0.012 | 1.026 | 294 | 5.2 | 5.2 | 0.909 | 0.909 | 92.2 | 64.9 | 67 | 49.7 |
| 6uiw | G | 2.7 | 20789 | 0.012 | 1.026 | 294 | 4.8 | 4.8 | 0.912 | 0.912 | 93.9 | 76.8 | 69.7 | 49.8 |
| 6uiw | H | 2.7 | 20789 | 0.012 | 1.026 | 294 | 4.1 | 3.7 | 0.918 | 0.918 | 92.2 | 75.3 | 77.2 | 14.1 |
| 6uiw | I | 2.7 | 20789 | 0.012 | 1.026 | 294 | 4.1 | 4.1 | 0.924 | 0.924 | 92.9 | 75.8 | 64.6 | 41.1 |
| 6uiw | J | 2.7 | 20789 | 0.012 | 1.026 | 294 | 5.8 | 5.0 | 0.901 | 0.901 | 90.8 | 68.2 | 67 | 45.2 |
| 6uiw | K | 2.7 | 20789 | 0.012 | 1.026 | 294 | 3.3 | 3.3 | 0.918 | 0.918 | 92.5 | 71 | 72.4 | 51.6 |
| 6uix | A | 3.5 | 20790 | 0.0097 | 1.026 | 275 | 4.8 | 4.8 | 0.855 | 0.855 | 86.5 | 60.5 | 58.2 | 8.8 |
| 6uix | B | 3.5 | 20790 | 0.0097 | 1.026 | 275 | 4.7 | 4.5 | 0.867 | 0.867 | 88 | 44.2 | 61.5 | 10.7 |
| 6uix | C | 3.5 | 20790 | 0.0097 | 1.026 | 275 | 4.3 | 4.3 | 0.877 | 0.877 | 89.8 | 57.1 | 53.8 | 4.7 |
| 6uix | D | 3.5 | 20790 | 0.0097 | 1.026 | 275 | 3.6 | 3.6 | 0.867 | 0.867 | 85.8 | 54.2 | 58.9 | 19.8 |
| 6uix | E | 3.5 | 20790 | 0.0097 | 1.026 | 275 | 3.7 | 3.7 | 0.888 | 0.888 | 89.8 | 54.7 | 58.2 | 3.1 |
| 6uix | F | 3.5 | 20790 | 0.0097 | 1.026 | 275 | 5.4 | 4.8 | 0.879 | 0.886 | 88 | 36.4 | 21.1 | 1.7 |
| 6uix | G | 3.5 | 20790 | 0.0097 | 1.026 | 275 | 3.8 | 3.8 | 0.866 | 0.866 | 88.4 | 43.2 | 63.6 | 5.1 |
| 6uix | H | 3.5 | 20790 | 0.0097 | 1.026 | 275 | 6.2 | 5.1 | 0.875 | 0.875 | 88.7 | 52 | 52.4 | 6.9 |
| 6uix | I | 3.5 | 20790 | 0.0097 | 1.026 | 275 | 3.5 | 3.5 | 0.877 | 0.877 | 87.6 | 53.5 | 57.8 | 34 |
| 6uix | J | 3.5 | 20790 | 0.0097 | 1.026 | 275 | 4.8 | 4.8 | 0.836 | 0.836 | 86.2 | 46.4 | 25.8 | 4.2 |
| 6uix | K | 3.5 | 20790 | 0.0097 | 1.026 | 275 | 3.1 | 3.1 | 0.878 | 0.878 | 89.8 | 54.3 | 57.8 | 5 |
| 6uix | L | 3.5 | 20790 | 0.0097 | 1.026 | 275 | 5.6 | 4.0 | 0.848 | 0.855 | 84.7 | 28.8 | 49.1 | 6.7 |
| 6uix | M | 3.5 | 20790 | 0.0097 | 1.026 | 275 | 4.2 | 4.2 | 0.876 | 0.876 | 88.7 | 49.2 | 50.5 | 8.6 |
| 6uix | N | 3.5 | 20790 | 0.0097 | 1.026 | 275 | 4.1 | 4.1 | 0.871 | 0.871 | 88.4 | 51.9 | 58.9 | 6.8 |
| 6uix | O | 3.5 | 20790 | 0.0097 | 1.026 | 275 | 6.3 | 3.6 | 0.820 | 0.875 | 87.6 | 57.7 | 52 | 11.9 |
| 6uix | P | 3.5 | 20790 | 0.0097 | 1.026 | 275 | 7.0 | 4.7 | 0.814 | 0.874 | 86.2 | 30.8 | 36.7 | 2 |
| 6uix | Q | 3.5 | 20790 | 0.0097 | 1.026 | 275 | 10.6 | 4.3 | 0.622 | 0.862 | 79.6 | 34.7 | 25.8 | 4.2 |
| 6uix | R | 3.5 | 20790 | 0.0097 | 1.026 | 275 | 4.0 | 4.0 | 0.855 | 0.855 | 85.1 | 47 | 56.4 | 6.5 |
| 6uix | S | 3.5 | 20790 | 0.0097 | 1.026 | 275 | 6.4 | 6.4 | 0.814 | 0.814 | 84.4 | 49.1 | 56.4 | 9 |
| 6uix | T | 3.5 | 20790 | 0.0097 | 1.026 | 275 | 4.0 | 4.0 | 0.870 | 0.870 | 85.8 | 53.4 | 49.1 | 10.4 |

| | | | | | | | | | | | | | | |
|---|---|---|---|---|---|---|---|---|---|---|---|---|---|---|
| 6uix | U | 3.5 | 20790 | 0.0097 | 1.026 | 275 | 4.6 | 4.6 | 0.849 | 0.849 | 84.7 | 44.6 | 57.8 | 26.4 |
| 6uix | V | 3.5 | 20790 | 0.0097 | 1.026 | 275 | 11.1 | 6.4 | 0.786 | 0.853 | 83.3 | 34.5 | 55.3 | 5.9 |
| 6ukj | H | 3.3 | 20806 | 8 | 1.035 | 127 | 4.9 | 4.0 | 0.852 | 0.865 | 89 | 52.2 | 89.8 | 28.9 |
| 6ukj | L | 3.3 | 20806 | 8 | 1.035 | 105 | 1.5 | 1.5 | 0.922 | 0.922 | 98.1 | 83.5 | 71.4 | 9.3 |
| 6ulg | A | 3.3 | 20814 | 0.002 | 1.04 | 121 | 2.6 | 2.6 | 0.879 | 0.879 | 90.9 | 66.4 | 83.5 | 7.9 |
| 6ulg | B | 3.3 | 20814 | 0.002 | 1.04 | 125 | 2.5 | 2.5 | 0.876 | 0.876 | 92 | 80 | 84 | 39 |
| 6ulg | C | 3.3 | 20814 | 0.002 | 1.04 | 91 | 11.9 | 10.9 | 0.484 | 0.582 | 79.1 | 27.8 | 75.8 | 4.3 |
| 6ulg | F | 3.3 | 20814 | 0.002 | 1.04 | 273 | 4.8 | 4.6 | 0.855 | 0.918 | 90.5 | 68.4 | 86.8 | 44.3 |
| 6um5 | C | 4.2 | 20817 | 0.7 | 1.08 | 126 | 17.9 | 15.6 | 0.319 | 0.411 | 61.1 | 3.9 | 38.9 | 10.2 |
| 6um5 | D | 4.2 | 20817 | 0.7 | 1.08 | 110 | 17.2 | 13.3 | 0.303 | 0.338 | 61.8 | 17.6 | 36.4 | 5 |
| 6um5 | G | 4.2 | 20817 | 0.7 | 1.08 | 126 | 16.4 | 16.2 | 0.235 | 0.346 | 63.5 | 3.8 | 46 | 5.2 |
| 6um5 | H | 4.2 | 20817 | 0.7 | 1.08 | 110 | 15.4 | 15.0 | 0.338 | 0.361 | 67.3 | 12.2 | 37.3 | 9.8 |
| 6um5 | K | 4.2 | 20817 | 0.7 | 1.08 | 126 | 14.4 | 14.4 | 0.372 | 0.407 | 66.7 | 10.7 | 44.4 | 8.9 |
| 6um5 | L | 4.2 | 20817 | 0.7 | 1.08 | 110 | 13.8 | 13.8 | 0.248 | 0.262 | 60.9 | 4.5 | 35.5 | 10.3 |
| 6um6 | C | 4.3 | 20818 | 0.7 | 1.08 | 126 | 18.9 | 14.6 | 0.288 | 0.378 | 68.3 | 7 | 50 | 9.5 |
| 6um6 | D | 4.3 | 20818 | 0.7 | 1.08 | 108 | 14.9 | 13.9 | 0.255 | 0.289 | 64.8 | 10 | 25.9 | 7.1 |
| 6um6 | G | 4.3 | 20818 | 0.7 | 1.08 | 126 | 15.7 | 15.1 | 0.310 | 0.467 | 69.8 | 4.5 | 50 | 9.5 |
| 6um6 | H | 4.3 | 20818 | 0.7 | 1.08 | 108 | 15.3 | 12.0 | 0.292 | 0.298 | 70.4 | 2.6 | 31.5 | 5.9 |
| 6um6 | K | 4.3 | 20818 | 0.7 | 1.08 | 126 | 15.6 | 14.2 | 0.302 | 0.437 | 72.2 | 5.5 | 31.7 | 5 |
| 6um6 | L | 4.3 | 20818 | 0.7 | 1.08 | 108 | 14.6 | 13.2 | 0.339 | 0.339 | 63 | 4.4 | 32.4 | 5.7 |
| 6um7 | B | 3.5 | 20819 | 1.18 | 1.07 | 126 | 16.6 | 15.0 | 0.286 | 0.444 | 69 | 9.2 | 54.8 | 10.1 |
| 6um7 | F | 3.5 | 20819 | 1.18 | 1.07 | 126 | 17.0 | 15.8 | 0.400 | 0.465 | 73 | 5.4 | 35.7 | 6.7 |
| 6um7 | G | 3.5 | 20819 | 1.18 | 1.07 | 110 | 15.5 | 11.5 | 0.402 | 0.530 | 71.8 | 10.1 | 46.4 | 9.8 |
| 6um7 | J | 3.5 | 20819 | 1.18 | 1.07 | 126 | 14.7 | 14.0 | 0.283 | 0.324 | 70.6 | 5.6 | 46 | 5.2 |
| 6um7 | K | 3.5 | 20819 | 1.18 | 1.07 | 110 | 10.6 | 8.7 | 0.431 | 0.721 | 75.5 | 12 | 46.4 | 9.8 |
| 6uph | A | 2.7 | 20839 | 0.16 | 0.849 | 93 | 1.4 | 1.4 | 0.946 | 0.946 | 98.9 | 91.3 | 88.2 | 58.5 |
| 6uph | B | 2.7 | 20839 | 0.16 | 0.849 | 79 | 2.2 | 2.2 | 0.902 | 0.902 | 93.7 | 85.1 | 89.9 | 77.5 |
| 6uph | C | 2.7 | 20839 | 0.16 | 0.849 | 97 | 2.2 | 2.2 | 0.927 | 0.927 | 96.9 | 86.2 | 94.8 | 82.6 |
| 6uph | D | 2.7 | 20839 | 0.16 | 0.849 | 96 | 3.3 | 3.3 | 0.907 | 0.907 | 94.8 | 80.2 | 91.7 | 70.5 |
| 6uph | E | 2.7 | 20839 | 0.16 | 0.849 | 97 | 1.7 | 1.7 | 0.932 | 0.932 | 97.9 | 94.7 | 93.8 | 95.6 |
| 6uph | F | 2.7 | 20839 | 0.16 | 0.849 | 79 | 1.6 | 1.6 | 0.926 | 0.926 | 97.5 | 93.5 | 86.1 | 100 |
| 6uph | G | 2.7 | 20839 | 0.16 | 0.849 | 99 | 2.7 | 2.7 | 0.898 | 0.898 | 96 | 76.8 | 92.9 | 55.4 |
| 6uph | H | 2.7 | 20839 | 0.16 | 0.849 | 92 | 1.4 | 1.4 | 0.923 | 0.923 | 95.7 | 85.2 | 97.8 | 100 |
| 6uzz | B | 3.1 | 20965 | 5 | 1.03 | 144 | 2.2 | 2.2 | 0.927 | 0.927 | 95.8 | 89.9 | 89.6 | 61.2 |
| 6uzz | D | 3.1 | 20965 | 5 | 1.03 | 144 | 2.3 | 2.3 | 0.900 | 0.900 | 92.4 | 66.9 | 83.3 | 7.5 |
| 6uzz | F | 3.1 | 20965 | 5 | 1.03 | 144 | 1.8 | 1.8 | 0.927 | 0.927 | 96.5 | 86.3 | 81.2 | 7.7 |
| 6uzz | H | 3.1 | 20965 | 5 | 1.03 | 144 | 1.9 | 1.9 | 0.906 | 0.906 | 94.4 | 84.6 | 78.5 | 8.8 |

| | | | | | | | | | | | | | | |
|---|---|---|---|---|---|---|---|---|---|---|---|---|---|---|
| 6v00 | B | 3.1 | 20966 | 6 | 1.03 | 144 | 1.3 | 1.3 | 0.945 | 0.945 | 97.9 | 95 | 82.6 | 59.7 |
| 6v00 | E | 3.1 | 20966 | 6 | 1.03 | 144 | 1.4 | 1.4 | 0.928 | 0.928 | 96.5 | 89.2 | 81.9 | 37.3 |
| 6v00 | H | 3.1 | 20966 | 6 | 1.03 | 144 | 1.9 | 1.9 | 0.926 | 0.926 | 94.4 | 83.8 | 84 | 6.6 |
| 6v00 | K | 3.1 | 20966 | 6 | 1.03 | 144 | 2.1 | 2.1 | 0.926 | 0.926 | 95.8 | 82.6 | 84 | 15.7 |
| 6v22 | E | 3.2 | 21025 | 6.19 | 1.04 | 199 | 11.5 | 6.8 | 0.720 | 0.852 | 87.4 | 41.4 | 85.4 | 44.1 |
| 6v22 | F | 3.2 | 21025 | 6.19 | 1.04 | 199 | 12.3 | 3.7 | 0.838 | 0.875 | 89.9 | 62 | 70.4 | 40.7 |
| 6v22 | G | 3.2 | 21025 | 6.19 | 1.04 | 199 | 13.1 | 11.5 | 0.782 | 0.782 | 83.4 | 47 | 78.9 | 45.9 |
| 6v22 | H | 3.2 | 21025 | 6.19 | 1.04 | 199 | 8.1 | 4.0 | 0.881 | 0.881 | 91 | 47.5 | 81.4 | 45.7 |
| 6v35 | E | 3.5 | 21028 | 6.7 | 1.3 | 198 | 13.4 | 13.4 | 0.351 | 0.607 | 78.8 | 13.5 | 79.8 | 24.7 |
| 6v35 | F | 3.5 | 21028 | 6.7 | 1.3 | 198 | 15.5 | 13.8 | 0.642 | 0.642 | 81.8 | 30.2 | 80.3 | 21.4 |
| 6v35 | G | 3.5 | 21028 | 6.7 | 1.3 | 198 | 12.4 | 12.2 | 0.715 | 0.715 | 83.3 | 34.5 | 78.3 | 21.9 |
| 6v35 | H | 3.5 | 21028 | 6.7 | 1.3 | 198 | 16.5 | 14.7 | 0.567 | 0.599 | 74.7 | 12.8 | 73.7 | 4.8 |
| | | | | | | Average | 9.8 | 8.4 | 0.648 | 0.694 | 81.7 | 41.0 | 66.4 | 22.1 |
