## Supplementary Table 5 for "Full-length *de novo* protein structure determination from cryo-EM maps using deep learning"

| SCOPe code |
| --- |
| d1acoa1 |
| d1c3ga1 |
| d1cfza_ |
| d1ctfa_ |
| d1dvka_ |
| d1e9ga_ |
| d1eaic_ |
| d1ecwa_ |
| d1ehia2 |
| d1em8b_ |
| d1f3va_ |
| d1f7ca_ |
| d1gm5a1 |
| d1gu7a2 |
| d1hbnc_ |
| d1hf2a2 |
| d1hw6a_ |
| d1hyha2 |
| d1i2ta_ |
| d1ijya_ |
| d1jnra3 |
| d1kbla2 |
| d1knya2 |
| d1kp8a3 |
| d1lama2 |
| d1ljoa1 |
| d1m1eb_ |
| d1m9sa2 |
| d1mb3a_ |
| d1oisa_ |
| d1okca_ |
| d1poca_ |
| d1q0qa1 |
| d1qtwa_ |
| d1r0va1 |
| d1rqpa1 |
| d1rtra_ |
| d1sg1x1 |
| d1shux_ |
| d1slua_ |
| d1syxb_ |
| d1thfd_ |
| d1un8a1 |
| d1ut7a_ |
| d1uura1 |

d1v8da_
d1vk5a_
d1vq8q1
d1vq8z1
d1w96a1
d1w96a2
d1wb9a3
d1wpna_
d1wpxb1
d1yqea1
d1zd0a1
d1zt2b1
d2auwa2
d2b9da1
d2dpla2
d2f1na1
d2f4mb2
d2fdna_
d2fhza1
d2gyka1
d2i15a1
d2j32a1
d2liga_
d2p1ma2
d2p61a1
d2pk8a_
d2qam31
d2r7511
d2v8qb1
d2vv5a2
d2wkxa1
d2z37a_
d2za4b_
d3ccda_
d3cx5e2
d3cx5g_
d3ezqb_
d3hola3
d3ieza_
d3ivva1
d3ka1a_
d3ldqb1
d3m7va2
d3mksb1
d3nfta_
d3o0gd_

d3rlff1
d3s2ra_
d3t8xa1
d3ziah1
d4eada3
d4gq4a1
d4gt9a1
d4peoa_
d4sgbi_
