## Supplementary Table 6 for "Full-length *de novo* protein structure determination from cryo-EM maps using deep learning"

| PDB id | Chain | Resolution (Å) | EMDB id | Recommended contour level | Pixel spacing (Å) |
|---|---|---|---|---|---|
| 6gyb | a | 3.3 | 0089 | 0.035 | 1.06 |
| 6gyb | b | 3.3 | 0089 | 0.035 | 1.06 |
| 6gyb | c | 3.3 | 0089 | 0.035 | 1.06 |
| 6h3i | B | 3.5 | 0133 | 0.072 | 0.82 |
| 6hu7 | B | 2.8 | 0273 | 0.04 | 1.056 |
| 6n3q | F | 3.7 | 0336 | 0.42 | 1.16 |
| 6nt5 | A | 4.1 | 0502 | 0.0233 | 0.84 |
| 6nt6 | A | 4 | 0503 | 0.024 | 0.84 |
| 6odi | b | 3.8 | 20020 | 0.027 | 1.64 |
| 6of4 | A | 3.2 | 20042 | 4.5 | 1.08 |
| 6ois | B | 3.6 | 20080 | 0.05 | 1.07 |
| 6ois | D | 3.6 | 20080 | 0.05 | 1.07 |
| 6oks | A | 4.2 | 20103 | 0.035 | 1.45 |
| 6os9 | A | 3 | 20180 | 0.065 | 1.06 |
| 6owt | M | 3.8 | 20217 | 0.485 | 1.149 |
| 6p7w | E | 4.1 | 20271 | 0.013 | 1.0605 |
| 6ptj | 2 | 3.8 | 20471 | 0.034 | 1.074 |
| 6ptj | 3 | 3.8 | 20471 | 0.034 | 1.074 |
| 6ptj | 4 | 3.8 | 20471 | 0.034 | 1.074 |
| 6ptj | 5 | 3.8 | 20471 | 0.034 | 1.074 |
| 6ptj | 6 | 3.8 | 20471 | 0.034 | 1.074 |
| 6ptj | B | 3.8 | 20471 | 0.034 | 1.074 |
| 6ptj | D | 3.8 | 20471 | 0.034 | 1.074 |
| 6pyh | B | 4.3 | 20524 | 0.018 | 1.07 |
| 6umm | A | 3.7 | 20820 | 0.004 | 0.82 |
| 5fn2 | C | 4.2 | 3237 | 0.04 | 1.4 |
| 5fn3 | B | 4.1 | 3238 | 0.04 | 1.4 |
| 5g06 | G | 4.2 | 3366 | 0.02 | 1.30654 |
| 5g06 | H | 4.2 | 3366 | 0.02 | 1.30654 |
| 5g06 | B | 4.2 | 3366 | 0.02 | 1.30654 |
| 5g06 | D | 4.2 | 3366 | 0.02 | 1.30654 |
| 5g06 | E | 4.2 | 3366 | 0.02 | 1.30654 |
| 5g06 | F | 4.2 | 3366 | 0.02 | 1.30654 |
| 5lcw | L | 4.2 | 4037 | 0.08 | 1.36 |
| 5lcw | Z | 4.2 | 4037 | 0.08 | 1.36 |
| 5lij | P | 4.2 | 4054 | 3 | 1.38 |
| 5ljo | D | 4.9 | 4061 | 0.5 | 1.04 |
| 5ljo | E | 4.9 | 4061 | 0.5 | 1.04 |
| 5lzp | N | 3.5 | 4128 | 0.045 | 1.4 |
| 6f1v | m | 3.4 | 4170 | 0.1 | 1.34 |
| 6f2d | A | 4.2 | 4173 | 0.07 | 0.86 |
| 6f2d | F | 4.2 | 4173 | 0.07 | 0.86 |
| 6f36 | M | 3.7 | 4176 | 0.03 | 1.105 |
| 6f36 | N | 3.7 | 4176 | 0.03 | 1.105 |

| | | | | | |
|---|---|---|---|---|---|
| 6fe8 | C | 4.1 | 4241 | 0.0664 | 1.06 |
| 6i7t | I | 4.6 | 4428 | 0.045 | 1.34 |
| 6i7t | K | 4.6 | 4428 | 0.045 | 1.34 |
| 6q6g | H | 3.2 | 4465 | 0.0085 | 1.047 |
| 6qlf | L | 3.5 | 4581 | 0.023 | 1.09 |
| 6qlf | P | 3.5 | 4581 | 0.023 | 1.09 |
| 6qm8 | b | 3.3 | 4591 | 0.0545 | 1.07 |
| 6qm8 | J | 3.3 | 4591 | 0.0545 | 1.07 |
| 6qnt | B | 3.5 | 4608 | 0.07 | 1.067 |
| 6qnt | D | 3.5 | 4608 | 0.07 | 1.067 |
| 6r23 | A | 4.9 | 4708 | 2.5 | 1.08 |
| 6r6b | A | 3.5 | 4734 | 0.01 | 0.822 |
| 6r6b | F | 3.5 | 4734 | 0.01 | 0.822 |
| 6r6b | G | 3.5 | 4734 | 0.01 | 0.822 |
| 6r70 | a | 3.5 | 4738 | 0.025 | 1.015 |
| 6r70 | b | 3.5 | 4738 | 0.025 | 1.015 |
| 6rd6 | 4 | 2.8 | 4807 | 0.04 | 1.053 |
| 6rd6 | P | 2.8 | 4807 | 0.04 | 1.053 |
| 6riu | A | 3.9 | 4898 | 0.01 | 1.058 |
| 6rlb | I | 4.5 | 4918 | 0.0318 | 1.39 |
| 3j9i | 1 | 3.3 | 5623 | 0.25 | 1.2156 |
| 3jc5 | C | 4.7 | 6535 | 0.018 | 1.01 |
| 5yi5 | A | 3 | 6830 | 0.3 | 1.3973 |
| 6bgo | J | 4.2 | 7098 | 0.00952 | 1.09 |
| 6bx3 | E | 4.3 | 7303 | 0.16 | 1 |
| 6bx3 | F | 4.3 | 7303 | 0.16 | 1 |
| 6bx3 | K | 4.3 | 7303 | 0.16 | 1 |
| 6c14 | A | 4.5 | 7328 | 7 | 1.04 |
| 6c6l | H | 3.5 | 7348 | 0.035 | 1.23 |
| 6coz | A | 3.4 | 7545 | 0.01 | 1.03 |
| 6d7l | A | 4 | 7823 | 0.0448 | 1.096 |
| 6dkf | A | 3.7 | 7952 | 0.54 | 1.06 |
| 5tcp | 0 | 4.3 | 8398 | 0.06 | 1.71 |
| 5tcp | 1 | 4.3 | 8398 | 0.06 | 1.71 |
| 5vfo | p | 3.5 | 8662 | 0.01 | 0.75 |
| 5vfo | s | 3.5 | 8662 | 0.01 | 0.75 |
| 5vot | F | 4.9 | 8721 | 0.09 | 1.72 |
| 5wc3 | 0 | 3.5 | 8795 | 0.045 | 1.35 |
| 6e0f | A | 3.7 | 8946 | 0.0341 | 1 |
| 6e14 | H | 4 | 8953 | 0.047 | 1.09 |
| 6ebm | B | 4 | 9026 | 0.02 | 0.835 |
| 6mdr | a | 3.5 | 9104 | 0.05 | 1.1 |
| 6mlu | A | 4 | 9146 | 0.033 | 1.31 |
| 6mrt | A | 2.8 | 9212 | 0.07 | 1.07 |
| 6muv | b | 3.8 | 9257 | 6 | 1.31 |
| 6muv | f | 3.8 | 9257 | 6 | 1.31 |

| | | | | | |
|---|---|---|---|---|---|
| 6muv | L | 3.8 | 9257 | 6 | 1.31 |
| 6muw | a | 3.6 | 9258 | 5 | 1.31 |
| 6muw | G | 3.6 | 9258 | 5 | 1.31 |
| 6muw | H | 3.6 | 9258 | 5 | 1.31 |
| 6muw | I | 3.6 | 9258 | 5 | 1.31 |
| 6muw | J | 3.6 | 9258 | 5 | 1.31 |
| 6ip1 | E | 3.9 | 9697 | 0.065 | 1.30654 |
| 6ixh | A | 4 | 9747 | 0.03 | 1.32 |
| 6jlx | A | 4.6 | 9841 | 0.0104 | 1.47 |
| 6jxr | n | 3.7 | 9895 | 0.0291 | 1.057 |
